# *Cutibacterium* adaptation to life on humans provides a novel biomarker of *C. acnes* infections

**DOI:** 10.1101/2024.09.18.613542

**Authors:** Md Shafiuddin, Gabriel William Prather, Wen Chi Huang, Jeffrey Ryan Anton, Andrew Lawrence Martin, Sydney Brianna Sillart, Jonathan Z. Tang, Michael R. Vittori, Michael J. Prinsen, Jessica Jane Ninneman, Chandrashekhara Manithody, Jeffrey P. Henderson, Alexander W. Aleem, Ma Xenia Garcia Ilagan, William H. McCoy

## Abstract

The domestication of cattle provided Propionibacteriaceae the opportunity to adapt to human skin. These bacteria constitute a distinct genus (*Cutibacterium*), and a single species within that genus (*C. acnes*) dominates 25% of human skin. *C. acnes* protects humans from pathogen colonization, but it can also infect indwelling medical devices inserted through human skin. Proteins that help Cutibacteria live on our skin may also act as virulence factors during an opportunistic infection, like a shoulder periprosthetic joint infection (PJI). To better understand the evolution of this commensal and opportunistic pathogen, we sought to extensively characterize one of these proteins, RoxP. This secreted protein is only found in the *Cutibacterium* genus, helps *C. acnes* grow in oxic environments, and is required for *C. acnes* to colonize human skin. Structure-based sequence analysis of twenty-one RoxP orthologs (71-100% identity to *C. acnes* strain KPA171202 RoxP_1) revealed a high-degree of molecular surface conservation and helped identify a potential heme-binding interface. Biophysical evaluation of a subset of seven RoxP orthologs (71-100% identity) demonstrated that heme-binding is conserved. Computational modeling of these orthologs suggests that RoxP heme-binding is mediated by an invariant molecular surface composed of a surface-exposed tryptophan (W66), adjacent cationic pocket, and nearby potential heme axial ligands. Further, these orthologs were found to undergo heme-dependent oligomerization. To further probe the role of this protein in *C. acnes* biology, we developed four monoclonal anti-RoxP antibodies, assessed the binding of those antibodies to a subset of ten RoxP orthologs (71-100% identity), developed an anti-RoxP sandwich ELISA (sELISA) with sub-nanogram sensitivity, and adapted that sELISA to quantitate RoxP in human biofluids that can be infected by *C. acnes* (serum, synovial fluid, cerebrospinal fluid). This study expands our understanding of how an environmental bacterium evolved to live on humans, and the assays developed in this work can now be used to identify this organism when it gains access to sterile sites to cause opportunistic infections.

**Author Summary:** The longer humans live, the more they require internal “replacement parts,” like prosthetic joints. Increased placement of these and other medical devices has increased their complications, which frequently are infections caused by microbes that live on humans. One of these microbes is *Cutibacterium acnes*, which dominates 25% of human skin. It appears that when humans domesticated cattle, a *C. acnes* ancestor adapted from living in cows to living on people. One of these adaptations was RoxP, a protein only found in *Cutibacterium* and carried by all *C. acnes*. Here, we describe our extensive characterization of RoxP. We found that distantly related RoxP conserve high stability at the low pH found on human skin. They also conserve the ability to bind heme, a source of iron used by microbes when they infect humans. As a part of this work, we developed tests that measure RoxP to identify *C. acnes* growth. In a clinic or hospital, these tests could allow a doctor to rapidly identify *C. acnes* infections, which would improve patient outcomes and lower healthcare costs. This work has helped us better understand how *C. acnes* adapted to live on humans and to identify *C. acnes* infections of medical devices.

## INTRODUCTION

Propionibacteriaceae grow in numerous environments (e.g., animals, water, sewage, soil) [1]. When humans domesticated cows, it appears that a bovine Propionibacteriaceae family member adapted to human skin [2]. *Cutibacterium* (formerly known as *Propionibacterium* [2]) *acnes* is the progeny of that microbial immigrant, and it evolved to dominate 25% of adult human skin. *C. acnes* is a lipophilic, gram-positive, aerotolerant anaerobe that accounts for 92% of the microbes that live on human sebaceous skin [3,4] (e.g., head, neck, upper trunk). This skin contains a high density of pilosebaceous units (PSUs) whose sebaceous glands produce sebum, a lipid-rich secretion that moisturizes human skin [5,6]. *C. acnes* catabolizes sebum triglycerides forming free fatty acids (FFAs) [7], and it significantly contributes to the low pH found on human skin [8]. The major *C. acnes* FFA, propionic acid, impedes pathogen colonization (e.g., *Staphylococcus aureus*) [9] while also improving skin barrier function by inducing epidermal lipid synthesis [10].

*C. acnes* is part of the normal human skin flora [11–13], yet it also contributes to human disease. *C. acnes* contributes to acne vulgaris [14], which impacts nearly every human [15]. In the United States each year, acne incurs $1.3 billion dollars in healthcare costs [16] and leads to >2 million office visits [15]. At these visits, patients are prescribed topical antimicrobial therapies and 5 million oral antibiotic prescriptions/year [17]. Extensive prior exposure to these anti-microbial treatments likely explains in part why *C. acnes* is not effectively reduced by pre-surgical skin sterilization or prophylactic antibiotics [18–20]. The effectiveness of these strategies to reduce surgical site infections by other bacteria (e.g., staphylococci) has likely contributed to *C. acnes* becoming an emerging pathogen of indwelling medical devices (IMDs) inserted through human skin [21].

*C. acnes*’ survival in the face of these measures may in part stem from its propensity to form biofilms [22–27]. *C. acnes* have been observed as macrocolonies that are consistent with biofilms in both its native PSU microenvironment [28] and on medical devices, like cerebrospinal fluid (CSF) shunt catheters [23] and hip prostheses [29]. With infection prophylaxis significantly reducing the burden of other microbes (e.g., staphylococci), *C. acnes* has emerged as the most common cause of shoulder prosthesis infection [30]. *C. acnes* is also tied with *S. aureus* as the most common cause of sternal wound infections [31], and it causes 4-15% of CSF shunt infections [32–34]. Further, the sonication of hip prostheses to dislodge microbial biofilms suggests that *C. acnes* is the most common cause (62%) of hip periprosthetic joint infection (PJI) [35].

*C. acnes* IMD infection treatment normally requires one or more revision surgeries and long-term antibiotics [36]. This approach exacts a significant toll on a patient’s quality of life, and it is very costly. For shoulder *C. acnes* PJI alone, these costs are projected to be >$250 million per year in the USA by 2030 [37–39]. Unfortunately, some patients also succumb to their *C. acnes* infections, as occurs for 15-27% of *C. acnes* endocarditis patients [40,41]. A significant contributor to these poor outcomes and high costs are routine delays in the identification of true *C. acnes* infections. These delays are due to *C. acnes*’ slow anaerobic growth (5-14 days) [42] and ubiquitous nature.

During routine clinical microbiology cultures, *C. acnes* is frequently dismissed as an environmental contaminant, particularly when more than one organism is isolated and the alternative is a microbe more commonly associated with infection (e.g., staphylococci). Yet, polymicrobial infections of coagulase-negative staphylococci (coNS) and *C. acnes* have been reported [35,29,30,43,31], which suggests that culturing staphylococci does not preclude *C. acnes* contributing to an infection. Strategies are needed to help identify true *C. acnes* infections. These strategies would allow for accurate assessments of *C. acnes* infection incidence and the optimization of *C. acnes* infection treatments. A better understanding of *C. acnes* biology is necessary to develop these strategies. Genomic studies have revealed that *C. acnes* lost 26 genes and acquired 108 genes as it moved from cows to humans [2], but those evolutionary adaptations have not been well characterized.

*RoxP* (Radical oxygenase of *Propionibacterium acnes*) is one of the genes that Cutibacteria acquired during their evolution to live on humans. This gene is only found in Cutibacteria, and it encodes a highly conserved, secreted, heme-binding protein [44]. Since RoxP is often the most highly secreted protein of *C. acnes* [45,46], it has been suggested that RoxP plays a significant functional role in host-microbe interactions. RoxP is essential for *C. acnes* colonization of human skin *ex vivo* and survival under oxic environments [44]. RoxP is remarkably conserved among all *C. acnes* phylotypes, including several clinical isolates [44]. A previous analysis of 106 *Cutibacterium* genomes showed that 86% of RoxP sequences share ≥99% identity and 55% of RoxP sequences share 100% identity [44]. Due to its high degree of conservation among *Cutibacterium* genus members and its potential to counteract oxic environments, it is possible that RoxP played an important role in *C. acnes* adaptation to human skin. Further, it is a potential *Cutibacterium*-specific growth biomarker.

RoxP’s poorly understood role in *C. acnes’* adaptation to life on humans led our group to perform an in-depth RoxP sequence analysis. This work led to biochemical investigations of several recombinant RoxP proteins, which demonstrated that ligand-dependent RoxP oligomerization is highly conserved in Cutibacteria. The high degree of RoxP sequence and functional conservation suggested to our group that anti-RoxP antibodies could help investigate RoxP’s role in *C. acnes* biology. We then generated four anti-RoxP antibodies and used them to develop *Cutibacterium*-specific immunoassays that can assess RoxP secretion in culture medium and human biofluids.

## RESULTS

### Assessment of RoxP Sequence Space

A BlastP search using the RoxP protein sequence from *C. acnes* strain KPA171202 (RoxP_1) identified 21 RoxP orthologs (RoxP_1-21, 71-99% identity to RoxP_1, **SI Table 1**). All sequence identities are relative to RoxP_1. RoxP_1 was chosen as the query sequence, because it was the first *roxP* gene evaluated [47]. These RoxP ortholog sequences were used to query the National Center for Biotechnology Information (NCBI) Identical Protein Groups (IPG) to identify all available RoxP sequences. A total of 946 IPG RoxP entries were identified, and 91% were from *C. acnes* (**Fig 1a**). Overlap in RoxP orthologs at the *C. acnes* subspecies level was identified by assessing (**Fig 1b**) protein identification numbers and (**Fig 1c**) IPG entries. The latter IPG assessment suggests that RoxP_6 is unique to *C. acnes* subsp. *defendens*. RoxP_1 was found to be the most common RoxP ortholog amongst all RoxP sequences (67%), *Cutibacterium* spp. (68%), *C. acnes* (71%), and *C. acnes* subsp. *acnes* (93%) (**Fig 1c**). Strain level RoxP analysis was possible for 96% of entries and showed similar RoxP ortholog profiles to the entire RoxP sequence space (**Fig 1c**,**d**).

**Figure 1.**
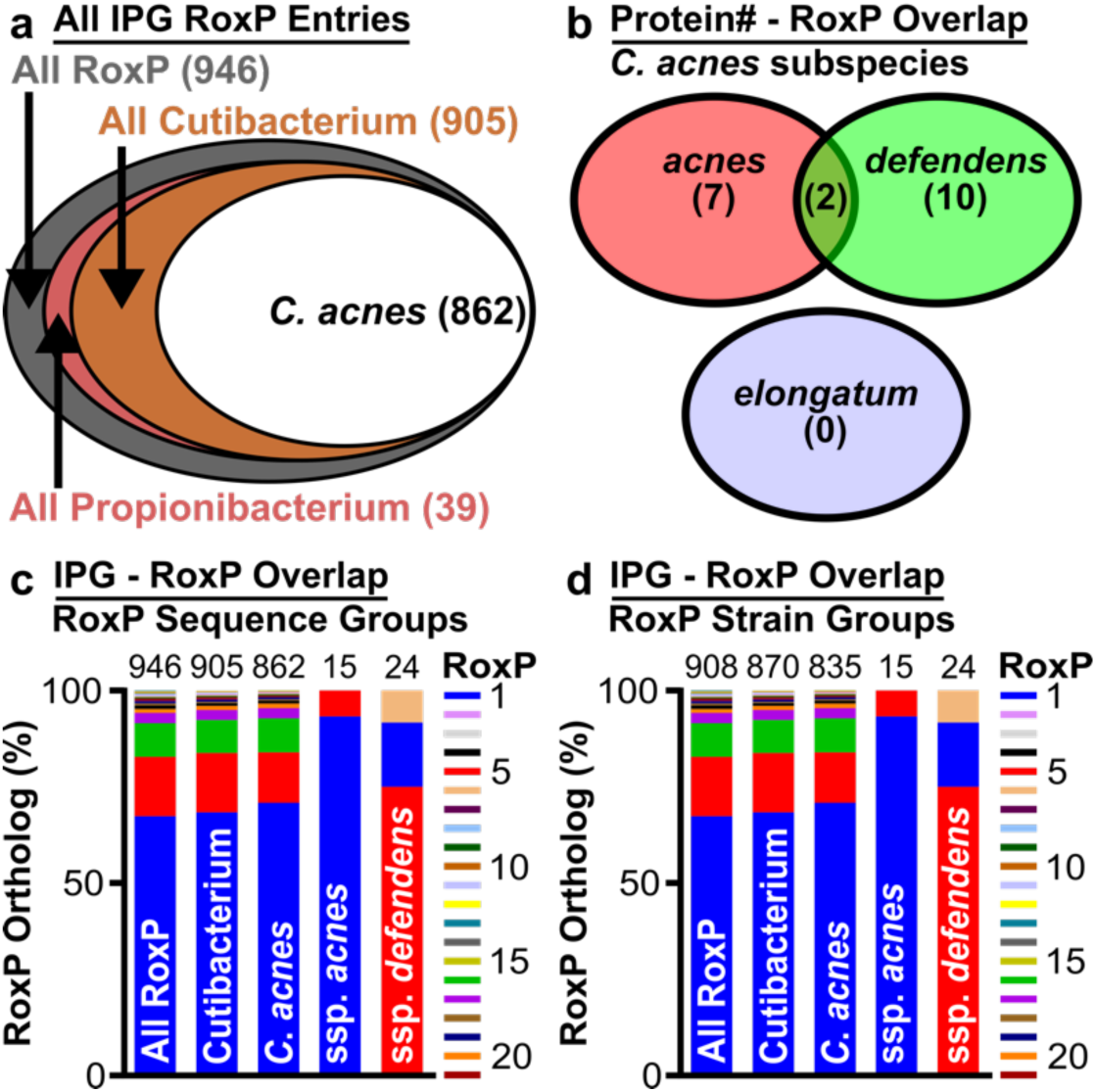
RoxP Sequence Space Analysis. RoxP sequence space was assessed by identifying the Identical Protein Group (IPG) entries for each of the 21 RoxP orthologs, and then assessing the number of entries shared at the (**a**) genus, (**b**,**c**) species, and (**d**) strain level. Species level RoxP ortholog assessments were performed (**b**) first by direct comparison of the unique protein identification numbers listed for each IPG entry and (**c**) then by an assessment of IPG entries alone, which revealed that multiple “unique” entries in (**b**) collapsed to two orthologs (RoxP_1,5) shared by *C. acnes* subsp. *acnes* and *defendens* and one ortholog (RoxP_6) that is unique to *C. acnes* subsp. *defendens*. (**c**) A comparison of *C. acnes* subsp. *acnes* and *defendens* RoxP ortholog carriage also revealed subspecies-specific ortholog profiles (*C. acnes* subsp. *acnes:* RoxP_1; *defendens*: RoxP_5).

To expand upon this analysis, we then assessed *roxP* gene carriage in publicly-available *Cutibacterium* genomes from named strains that included species-level information (N=189). By assessing *roxP* gene carriage by each strain in the context of *C. acnes* phylotypes (**SI Table 2**), we determined that RoxP_1 is present in type I and II phylotypes, but it was not found in *C. acnes* phylotype III. Other RoxP orthologs were found primarily in *C. acnes* whole genome sequence (WGS) clades IB-1 (RoxP_17, 99% identity), IB-2 (RoxP_16, 99% identity), II (RoxP_5, 82% identity; RoxP_6, 81% identity), and III (RoxP_2, 98% identity). RoxP_5 was also found in *C. modestum* (formerly known as *Propionibacterium humerusii* [43]), while RoxP_7 and _8 were exclusively found in *C. namnetense.* This analysis demonstrated that some *Cutibacterium* species utilized specific RoxP orthologs, but it was not yet clear if *C. acnes* subspecies-specific RoxP orthologs existed beyond RoxP_6.

While our initial assessment of RoxP sequence space did not identify a *C. acnes* subsp. *elongatum* RoxP (**Fig 1b**,**c**), the use of phylotype III as a surrogate marker of this subspecies [48] in **SI Table 2** entries that were only annotated as *C. acnes* (N=123) suggests that RoxP_2 is unique to *C. acnes* subsp. *elongatum* (N=3). Extending that strategy to *C. acnes* subsp. *defendens* based on phylotype II [49] expanded our RoxP ortholog assessment of this subspecies from 5 annotated strains (80% RoxP_5, 20% RoxP_6) to 31 total strains (61% RoxP_5; 26% RoxP_1; 3% RoxP_4,6,18,21). Applying this strategy to *C. acnes* subsp. *acnes* (phylotype I) expanded our RoxP ortholog assessment of this subspecies from 2 annotated strains (100% RoxP_1) to 75 total strains (76% RoxP_1; 11% RoxP_16; 7% RoxP_4; 4% RoxP_17; 1% RoxP_10,19). IPG analysis also identified *C. acnes* subsp. *acnes* with RoxP_5 (**Fig 1c**), though that was not observed in our genome analysis. Subdividing this phylotype I search into IA/B/C revealed a similar ortholog profile for all three types (IA, N=50: RoxP_1>16>17>19 [77>15>6>2%]; IB, N=27: RoxP_1>16>17>10 [67>19>11>4%]; IC, N=1, RoxP_1). Interestingly, RoxP_17 appeared in strains that were labeled as IA/IB suggesting this ortholog may be unique to phylotype I strains that may be typed as IA or IB depending on the phylotyping strategy. This *roxP* gene carriage assessment clearly shows that there are specific RoxP ortholog profiles found in *C. acnes* subspecies, and relatively, three orthologs segregate *C. acnes* subspecies *acnes* (RoxP_1); *defendens* (RoxP_5,6); and *elongatum* (RoxP_2).

The literature was then reviewed to determine which RoxP ortholog sequences had been previously experimentally evaluated (**SI Table 3**). While 43% of orthologs had been included in prior multiple sequence alignments, there had been minimal evaluation of transcription (14%), secretion (29%), function (10%), or structure (5%). Analysis of the RoxP secretion literature (**SI Table 4**) revealed that the secretion of six RoxP orthologs (RoxP_1,2,5,16,17,19; sequence identity 82-100%) had been previously observed, but only one of these studies directly investigated RoxP [47]. All studies were performed under anoxic conditions at 37°C, and most studies were performed during exponential growth phase in planktonic growth mode. All evaluated RoxP orthologs were found to be secreted into the media (cell-free). A subset of five orthologs (RoxP_1,2,5,16,17) were also found to be cell surface-associated (cell surface/wall) [50,51]. Theis analysis of prior literature suggested that RoxP secretion and localization (cell-free, surface-associated) is conserved across orthologs.

### Assessment of RoxP Sequence Variation

To explore RoxP sequence variation and attempt to correlate it with the prior literature, a multiple sequence alignment (MSA) of all 21 RoxP orthologs was generated, and then the predicted per residue solvent-accessible surface area (SASA) was calculated using the nuclear magnetic resonance (NMR) structure of apo-RoxP_1 [52] (**SI Fig 1**). This information was used to evaluate conservation of molecular surface features like amino acid side chains, electrostatics, and potential functional residues (**Fig 2**). ConSurf analysis (**Fig 2a**) and identification of non-conservative amino acid variants (**Fig 2b-c**) identified a hydrophobic patch (**Fig 2d**) surrounded by potential heme axial ligands (**Fig 2e**).

**Figure 2.**
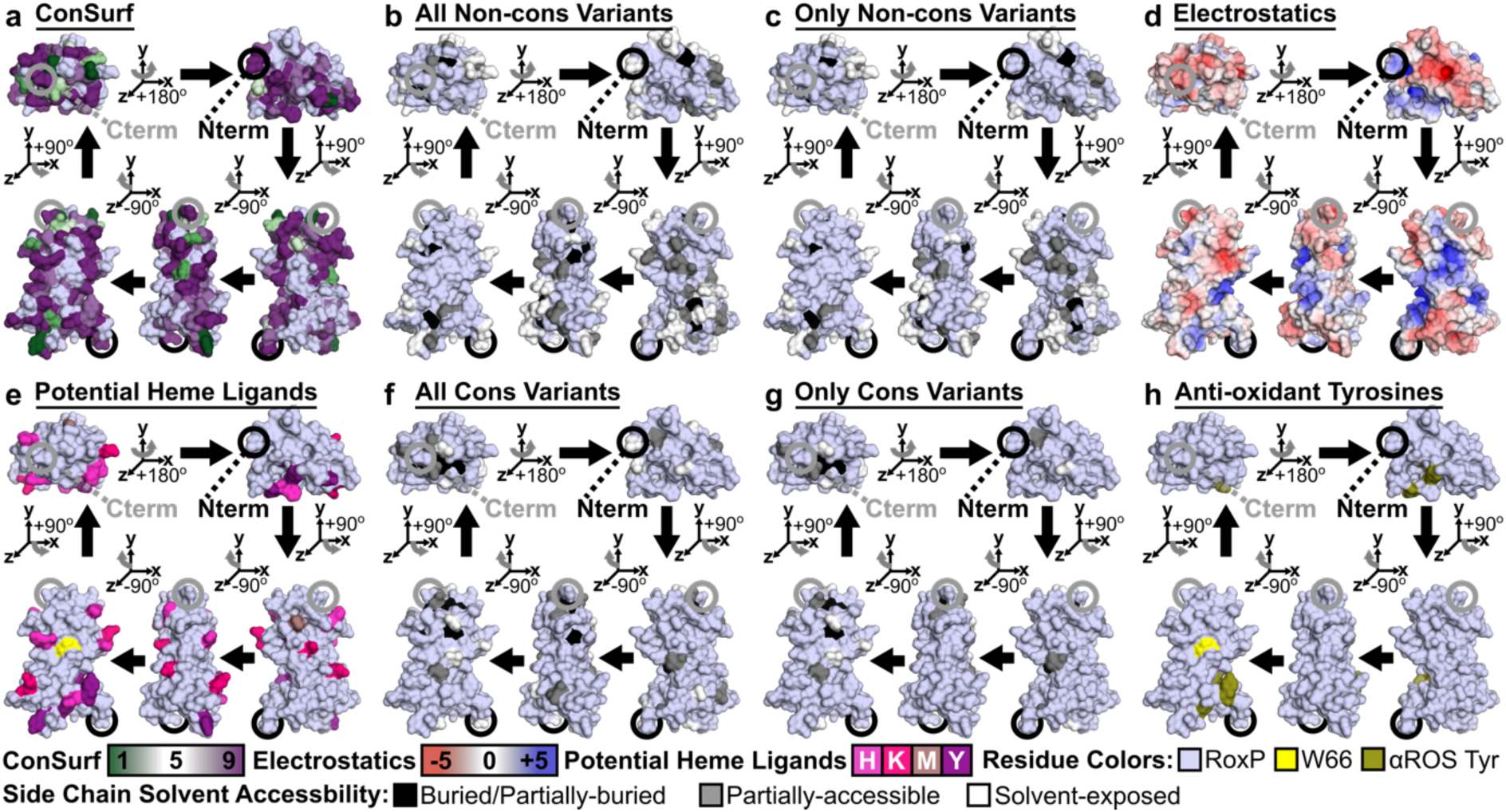
RoxP Molecular Surface Conservation. RoxP molecular surface conservation, chemical nature, and amino acid exposure were evaluated using a RoxP surface generated from PDB 7bcj (NMR state 1). This RoxP surface was colored blue-white except where noted in the legend below panels (**a**-**h**). Polypeptide termini (amino(N)-/carboxyl(C)-termini = N/Cterm) and model rotations are shown. (**a**-**c**,**f**,**g**) Surface conservation analysis is based on a multiple sequence alignment (MSA) of 21 RoxP orthologs (**SI Fig 1**). (**b**,**c**,**f**,**g**) Coloring of variable side chains based on solvent accessibility is the same as that described in **SI Fig 1** except that buried and partially-buried were combined. Buried (≤1%) residues did not contribute to any colored surface in these panels. Conservative in these panels is abbreviated “Cons.” (**a**) The ConSurf web server (http://consurf.tau.ac.il) used the RoxP MSA and PDB 7bcj to assess evolutionary conservation of amino acids (e.g., low = 1, high = 9). The color of residue positions with insufficient variation to be assessed (e.g., invariant positions) remained blue-white. (**b**,**c**) Non-conservative variants (e.g., a charge reversal like E to K) are shown (**b**) in total and (**c**) only when exclusively non-conservative variation was present. See **SI File 2** for a complete definition of non-conservative/conservative substitutions and analysis of all RoxP residue variants. (**d**) An electrostatic surface was generated in PyMOL to identify positive (blue), uncharged (white), and negative (red) RoxP surfaces. (**e**) All potential heme axial ligands (Cys, His, Lys, Met, Tyr) were colored shades of magenta and the partially-buried tryptophan (Trp66, W66) was colored yellow. RoxP’s two cysteines are not visible, as they form a buried, disulfide bond. (**f**,**g**) Conservative variants (e.g., a small hydrophobic substitution like L to V) are shown (**f**) in total and (**g**) only when exclusively non-conservative variation was present. (**h**) All Tyr residues previously shown to be able to reduce free radicals (αROS: anti-reactive oxygen species) were colored deep-olive and partially-buried W66 was colored yellow.

This surface also contained an invariant, partially surface-exposed (12.6% exposed compared to random coil) tryptophan (RoxP_1 W66) (**SI Fig 1**). Surface-exposed tryptophan side chains have been reported to contribute to heme-binding [53–55]. Evaluation of conservative amino acid variation (**Fig 2f-g**) also supported conservation of this unique interface with W66 at its center. This hydrophobic surface is distinct from the previously identified loop containing three invariant tyrosines (Y110,113,116) that mediate RoxP’s non-enzymatic, anti-oxidant activity (**Fig 2h**) [52].

Conservation of this surface suggests that it could be a conserved RoxP heme-binding site. A previous study demonstrated RoxP_1 heme-binding using Soret peak analysis, hemin-agarose (HA) pulldown, and NATIVE gel heme-stain [47], while a subsequent study by the same group on RoxP_1 concluded that their Soret peak analysis did not demonstrate binding to either heme or protoporphyrin IX [52]. Due to this inconsistency in experimental outcomes, we chose to conduct an in-depth biochemical analysis of RoxP ligand binding.

### Biochemical Characterization of Apo-RoxP

Before assessing RoxP ligand binding, we biochemically characterized the ligand-free (apo) form of RoxP. We chose to work with the RoxP_1 ortholog (RefSeq ID: WP_002515361.1), because it was (1) the most commonly observed ortholog (**Fig 1c**,**d**), (2) encoded in the genome of the first sequenced *C. acnes* strain (KPA171202) [56], and (3) previously assessed for heme-binding in the literature [47,52]. RoxP_1 was cloned downstream of an 8 histidine (8His) tag followed by a Tobacco Etch Virus (TEV) cleavage site. This construct was then recombinantly expressed in *Escherichia coli*, 8His-TEV-RoxP_1 was purified from *E. coli* lysate using immobilized-metal affinity chromatography (IMAC), and tagless RoxP protein was produced via TEV digest followed by IMAC capture of tagged proteins (**Fig 3a**). Size-exclusion chromatography (SEC) showed that 8His_TEV_RoxP_1 elutes predominantly as a single, monodispersed peak at a SEC-estimated molecular weight (MW) of 21 kilodaltons (kDa) (**Fig 3b**,**c**). This MW is consistent with the monomer MW (1) predicted after PelB signal peptide cleavage (17.2 kDa) and (2) observed by sodium dodecyl-sulfate polyacrylamide gel electrophoresis (SDS-PAGE) for tagged RoxP (**Fig 3 a**,**c**; MW ∼19 kDa, +His), as well as tagless-RoxP (-His) produced after TEV proteolysis (**Fig 3 a**; MW 14.9 kDa).

**Figure 3.**
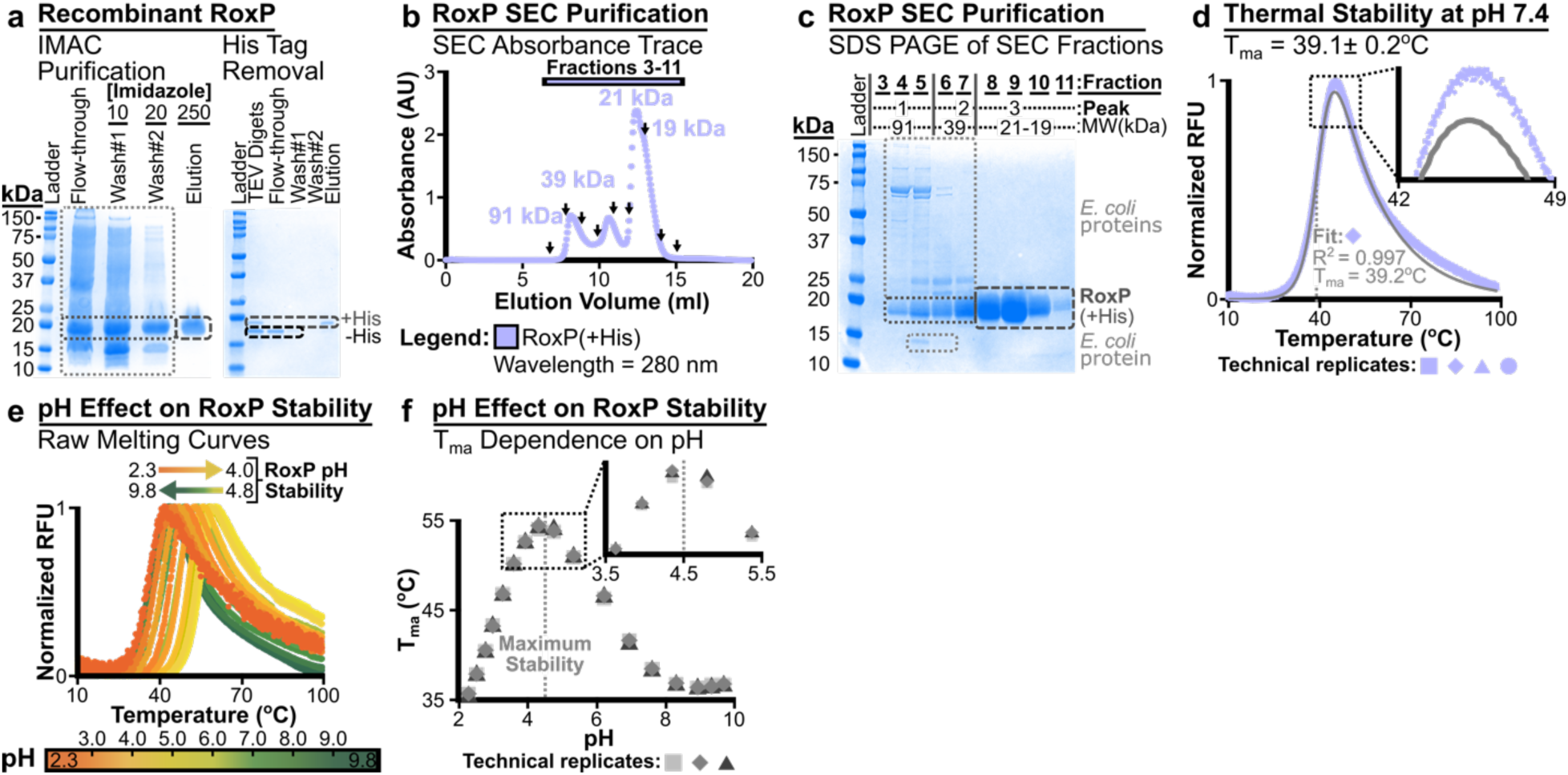
RoxP Biochemical Evaluation. Recombinant RoxP_1 was (**a**) purified, and then its (**b-c**) molecular size/weight, (**d**) thermal stability, (**e-f**) and pH stability were assessed. (**a**) RoxP_1 purified from *Escherichia coli* lysate using immobilized metal affinity chromatography (IMAC) was >95% pure, and TEV digest of this protein followed by IMAC capture of His-tagged proteins produced tagless RoxP_1. High MW protein ladder bands (75, 150 kDa) are indicated by a **black** tick mark. Short-dashed boxes: **light grey** (*E. coli* proteins and lysozyme [15 kDa band]); **dark grey** (RoxP_1). Long-dashed boxes (>95% pure RoxP_1): **dark grey** (+His tag); **black** (-His tag = tagless). IMAC experiment details: left gel (6.0 L culture, 0.5 mL Ni-NTA, gravity-flow, single application of lysate, 22.0 mg yield of RoxP_1 ^a^), right gel (22.0 mg RoxP_1 cut with 12 µg TEV at 4°C for 72 hours ^b^, ∼75% tag cleavage, 2.0 mL Ni-NTA, gravity-flow, 25.0 mg yield of tagless RoxP_1 ^c^). Imidazole concentration is in mM. (**b**) Size-exclusion chromatography (SEC) was used to evaluate IMAC-purified RoxP_1 (recovery run #2 ^a^). The major peak/shoulder have predicted MWs consistent with monomeric RoxP_1. Peak 1 is likely >91 kDa, as the fractionation range of this column (Superdex^TM^ 75 Increase 10/300 GL) is 70-3 kDa. SEC profile evaluated by SDS PAGE (**c**) is indicated with individual fractions marked (,). Absorbance saturation at ∼12.5 mL occurred due to application of more sample (0.6 mL of 100 mg/mL) than normally recommended for this column (e.g., ≤50 mg/mL, ≤500 µL). See **SI Fig 5a** for SEC MW calibration curve. (**c**) SDS PAGE of SEC fractions in (**b**) indicated with (,) is shown with boxes as described in (**a**). Peaks and their associated MWs that were assessed by specific fractions are indicated above the gel. An equal volume of each fraction was loaded per lane. (**d-f**) Differential scanning fluorimetry (DSF) was used to ascertain the apparent melting temperature (T_ma_) of RoxP_1 in (**d**) HEPES Sizing Buffer (HSB: 20 mM HEPES pH 7.4, 150 mM NaCl, 0.01% azide) and (**e**,**f**) as a function of pH. (**d**) Data from four replicates and a fit for one replicate are shown. (**e**,**f**) RoxP appears most stable at pH ∼4.5 based on both (**e**) raw DSF melting curves and (**f**) T_ma_ values from fitting the raw data. **^a^ Two subsequent IMAC purifications (recovery runs #2 and #3) over 6.0 mL of Ni-NTA (three applications of sample) using this lysate captured 282 mg and then 195 mg of RoxP_1 protein, respectively. Thus, the total RoxP_1 protein yield was ∼500 mg, and ∼83 mg of RoxP_1 protein was produced per liter of culture.** **^b^ TEV digest was optimized (e.g., lower protein mass ratio, shorter incubation time) after this experiment.** **^c^ Pre-/post-TEV digest RoxP_1 yield discrepancy is most likely due to 10 mM imidazole present in the protein sample used for the TEV digest. As such, the final RoxP_1-tagless yield was ∼95 mg per liter of *E. coli* culture.**

Recombinant RoxP_1 did not appear to be bound to an endogenous *E. coli* porphyrin, as absorbance scans from 220-800 nm did not identify peaks beyond the 280 nm protein peak. Therefore, RoxP_1 appeared to be purified in its apo-form. Apo-RoxP_1-tagless could then be concentrated to ∼282 mg/mL, which suggested that it was a highly stable protein. Surprisingly, differential scanning fluorimetry (DSF) assessment of the apo-RoxP_1-tagless apparent melting temperature (T_ma_) in SEC buffer (HEPES Sizing Buffer [HSB]: 20 mM HEPES pH 7.4, 150 mM NaCl, 0.01% azide) revealed that it has very low T_ma_ of 39.1±0.2°C (**Fig 3d**). Since human skin structures (e.g., PSU) exist across a vertical temperature gradient (deep: 37°C; surface: 31.6°C [57]) that also varies regionally across the face (30.4-36.9°C [57–59]), we determined if RoxP might be more stable under conditions more like those found in the PSU. We examined the effect of pH on apo-RoxP_1-tagless and found that (1) low pH stabilized this protein’s T_ma_ by >15°C (**Fig 3e**,**f**) and (2) it appeared to be most stable at pH ∼4.5, which is consistent with the pH 4.0-5.5 found on forehead skin [60,61].

### Assessment of RoxP Ligand Binding

After thoroughly characterizing recombinant apo-RoxP_1-tagless, we moved on to assess RoxP’s ability to bind heme. First, heme-binding was demonstrated for apo-RoxP_1-tagless by HA pulldown (**Fig 4a**). A negative control protein of similar MW (hSCNkk) was not significantly pulled down by HA. Pre-incubation with hemin did not prevent RoxP_1 pulldown, which may support hemin exchange during the experiment, as was observed for hemoglobin (data now shown). Soret peak analysis then demonstrated RoxP:hemin binding reached equilibrium by 2.3 hours in HSB at room temperature (RT). Soret peak analysis also revealed evidence of both penta-(370 nm) and hexa-coordinated complexes (412 nm) [62] (**Fig 4b**). In comparison, heme-binding control proteins bovine serum albumin (BSA) and human hemopexin (Hmp) reach equilibrium in this assay at 1.8 and 0.3 hr, respectively (**SI Fig 2 a**-**d**). The time required for BSA/Hmp:hemin to reach equilibrium has previously been attributed to slow dissociation of hemin dimers into monomers in aqueous solutions [63]. Heme preparation (hemin in DMSO vs. hematin in 1.4 N NaOH) was also found to produce similar RoxP Soret peaks (**Fig 4b**, **SI Fig 2 e**,**g**,**h**).

**Figure 4.**
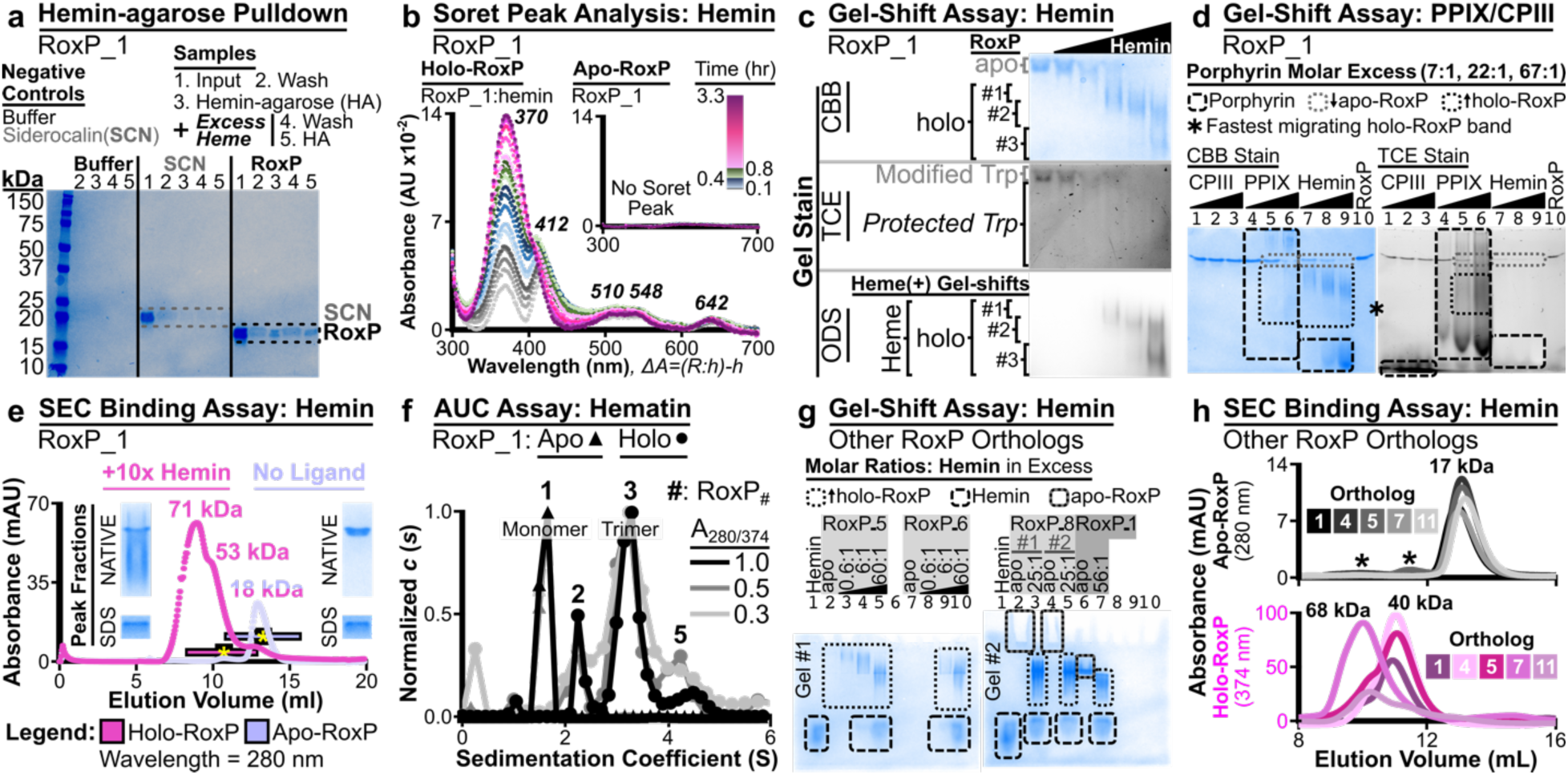
Assessment of RoxP:Porphyrin Binding. Using tagless, recombinant RoxP proteins, porphyrin-binding assays were performed for (**a-h**) RoxP_1 and (**g**,**h**) other RoxP orthologs. (**a**) Heme-agarose (HA) pulled down RoxP_1 (RoxP, lane 3), but it did not significantly pull down a similarly-sized, tagless, negative control protein (NCP) involved in human nutritional immunity (SCN, lane 3). Pre-incubation with excess of hemin did not prevent HA pulldown of RoxP_1 (RoxP, lane 5), but it eliminated non-specific, HA-pulldown of NCP (SCN, lane 5). SCN and RoxP bands are outlined by dashed, **grey** and **black** lines, respectively. (**b**) Soret peak analysis of holo-RoxP (RoxP_1:hemin, 10:1 molar ratio) is compared to an inset of the same analysis performed on apo-RoxP (RoxP_1). Scans were taken over 3.3 hr. Soret peaks and Q-bands are shown in ***bold italics***. Incubation time prior to each scan is shown in the legend (top-right) as a color gradient. Color gradient heights were scaled to the time span examined. From bottom-to-top, the color gradient and number of included traces for each incubation time span are grey-scale (0-240 s, 5 traces), blue (300-1600 s, 5 traces), green (1560-2760 s, 3 traces), and magenta (2820-11819 s, 6 traces). All spectra were blank-subtracted. See x-axis label for difference spectra calculation. <u>Abbreviations</u>: absorbance (A), RoxP (R), hemin (h), RoxP:hemin (R:h). (**c**,**d**) NATIVE gel-shift assessment demonstrated RoxP_1 porphyrin binding of (**c**) hemin and (**d**) protoporphyrin IX (PPIX) but not coproporphyrin III (CPIII). (**c**) RoxP:heme binding was shown by titratable, hemin-dependent (i) decrease in apo-RoxP, (ii) increase in holo-RoxP, (iii) protection of RoxP W66 by heme-binding, and (iv) the co-migration of heme with three distinct holo-RoxP species. <u>Abbreviations</u>: CBB (Coomassie brilliant blue), 2,2,2-TCE (trichloroethanol = W protection assay), ODS (O-dianisidine = heme stain). See **SI Fig 3** for images of entire gels that include control lanes (e.g., hemin alone, NCPs). (**d**) RoxP:PPIX binding was shown by titratable, PPIX-dependent (i) decrease in apo-RoxP and (ii) increase in holo-RoxP visualized by both CBB and TCE. As opposed to hemin and CPIII that migrate to the bottom of the gel (**Fig 4d; SI Fig 4d,e**), PPIX alone at 375 µM (67:1 molar PPIX concentration) displays distinct CBB and TCE staining patterns that extend throughout the lane but are distinct from holo-RoxP bands (**SI Fig 4a,b**, lane 2). (**e**) Size-exclusion chromatography (SEC) was performed on apo-RoxP_1 alone and pre-incubated with 10-fold molar excess of hemin. Holo-RoxP eluted significantly earlier than apo-RoxP with a higher 280 nm absorbance (A_280_) and NATIVE gel-shift bands consistent with holo-RoxP. This SEC peak contained (**SI Fig 5 c-f**) NATIVE gel-shift bands that exhibited tryptophan protection (TCE negative) and heme co-migration (ODS positive). (**SI Fig 5 g**) Comparison of apo-/holo-RoxP peak SDS PAGE gel samples suggests that the increased holo-RoxP A_280_ (∼36 mAU) is due to hemin, as the apo-RoxP peak fraction contained 10-fold more protein than the holo-RoxP peak fraction. See **SI Fig 5a** for SEC MW calibration curve and **SI Fig 5d/g** uncropped gel images (NATIVE, SDS PAGE) shown in this panel. The cropped gel images in panel € are from the fractions marked with a yellow asterisk (*), and all fractions examined in **SI Fig 5 c-g** are shown as a black-outlined bar with the same color as the A_280_ trace. (**f**) Analytical ultracentrifugation (AUC) was performed on apo-RoxP_1 and SEC-purified, holo-RoxP_1 (RoxP:hematin) to assess heme-dependent oligomerization. See **SI Fig 5b** for MW calibration curve of SEC used to prepare holo-RoxP_1. (**g**) NATIVE gel-shift assessments demonstrated RoxP_5,6,8 all bind heme. (***Gel #1***) Apo-RoxP_5/6 (pI 9.56) did not enter the gel run with forward polarity, while (***Gel #2***) apo-RoxP_8 (pI 8.98) slightly entered the borders of the wells (lanes 2,4) and disappeared upon heme-binding (lanes 3,5). Apo-RoxP_1 (pI 6.19) entered the gel #2 (lane 6), as seen in **Fig 4c,d**. Single biologic replicates are shown for RoxP_5/6, and two biologic replicates are shown for RoxP_8. Unlabeled lanes were loaded with blank samples. (**h**) SEC analysis of RoxP_1,4,5,7,11 demonstrated a heme-dependent increase in MW (+23-51 kDa) for all assessed orthologs. Absorbance at 374 nm (A_374_) was used to monitor heme-bound RoxP (holo-RoxP). Holo-RoxP A_280_ peaks coincided with A_374_ peaks due to both protein and hemin absorbing at 280 nm. Apo-RoxP and hemin alone did not contain any A_374_ SEC trace peaks. Asterisks mark small A_280_ peaks (<1 mAU) that eluted prior to (RoxP_4: 70 kDa; RoxP_1,5: 35 kDa) the main apo-RoxP peak (17 kDa).

The conditions assessed by this spectroscopy work were used to develop a NATIVE gel-shift assay that revealed heme-titration-dependent (1) appearance of new, faster migrating bands (holo-RoxP) by Coomassie brilliant blue (CBB) staining; (2) protection of RoxP’s partially-buried tryptophan (W66) from 2,2,2-trichloroethanol (TCE) modification [64]; and (3) heme-staining of the new, faster migrating bands (**Fig 4c**, **SI Fig 3**). NATIVE gel-shift analysis of two *C. acnes* porphyrins[65,66] revealed weaker binding to PPIX but no detectable binding to coproporphyrin III (CPIII) (**Fig 4d**) even after extended incubation (0.5 hr to overnight, **SI Fig 4a-e**). Like hemin, PPIX protected apo-RoxP_1 W66 from TCE modification (**Fig 4d**, **SI Fig 4b**). Unlike hemin though, PPIX also produced a distinct TCE-positive gel-shift coincident with the RoxP_1:PPIX CBB gel-shift, likely due to TCE modification of metal-free porphyrins (**SI Fig 4b** – PPIX, **SI Fig 4d** – CPIII).

Weak/absent RoxP binding of *C. acnes* porphyrins may be due to the assayed condition (HSB, pH 7.4) being quite different from the low pH, lipid-rich PSU that *C. acnes* normally inhabits. This hypothesis is supported by an observation that we made during the development of our RoxP-hemin NATIVE gel-shift assay. A minor increase in running buffer pH from 8.3 to 8.6 decreased RoxP:hemin gel-shifts suggesting a decrease in affinity (**SI Fig 4f**,**g).**

While NATIVE gel-shift band migration is not always an indicator of a change in oligomeric state, two observations from this work suggested that might occur with RoxP:heme binding. First, the appearance of multiple new bands with hemin titration (**Fig 4c**, **SI Fig 3**) suggested there were multiple RoxP:heme species present. Second, extended incubation (0.5 hr to overnight) favored the appearance of a distinct, faster migrating holo-RoxP species (**SI Fig 4e**,**g**,**h**). These observations led us to assess the holo-RoxP oligomeric state using SEC and analytical ultracentrifugation (AUC). SEC suggested that heme-binding increased RoxP’s MW (**Fig 4e**), while AUC demonstrated several RoxP:heme oligomeric states (trimer >> dimer > pentamer)(**Fig 4f**). Heme-binding after SEC was confirmed by NATIVE gel-shift analysis (**Fig 4e**, **SI Fig 5c-f**), as well as heme-specific absorbance at 374 nm **(Fig 4f**). These findings revealed that RoxP_1 undergoes heme-dependent oligomerization, which had not been previously identified.

**Figure 5.**
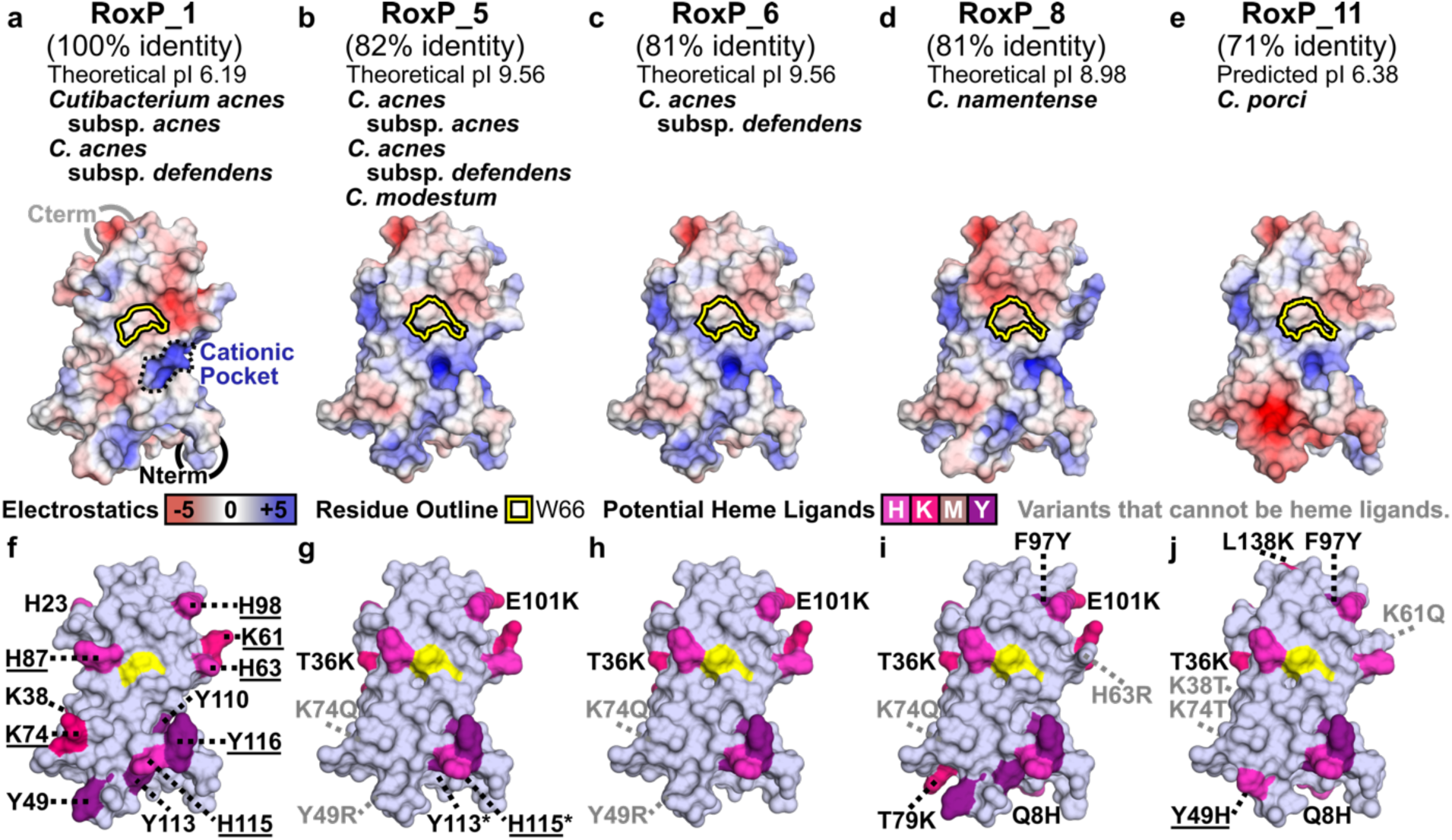
Molecular Modeling to Characterize the RoxP Heme-Binding Site. For each heme-binding RoxP ortholog (**SI Table 5**), sequence identity relative to RoxP_1, theoretical pI, and *Cutibacterium* species/subspecies that encode the ortholog are shown. (**a**-**e**) Electrostatic surfaces and (**f**-**j**) potential heme ligands are shown for (**a**,**f**) the NMR structure of RoxP_1 (PDB 7bcj) and (**b**-**e**, **g**-**j**) three-dimensional models of four RoxP orthologs (RoxP_5, 6, 8, 11). The partially-buried tryptophan surface is (**a**-**e**) outlined or (**f**-**j**) colored yellow (see Fig 2) on each molecular surface, and the (**a**) **cationic pocket** is outlined by a **black**, dashed line. (**f**-**j**) Potential RoxP heme axial ligands are labeled in **black,** <u>underlined</u> if in close proximity to W66, and colored according to Fig 2e. (**g**-**j**) Ortholog positions lacking a potential heme axial ligand found in RoxP_1 are labeled in **grey**. (**f**) Three potential RoxP_1 heme axial ligands (Y105, K130, M133) located on the molecular face opposite that containing the partially-exposed tryptophan residue were not labeled. (**g**) Asterisks mark side chains that flipped during modeling of RoxP_4,5,6,11, which might make it difficult to identify two potential, invariant heme ligands (Y113, H115). See **SI Fig 7** for this analysis of two additional heme-binding orthologs (RoxP_4, 7).

### Evolutionary Conservation of RoxP Biochemistry and Ligand Binding

Due to RoxP’s high sequence conservation (**Fig 1**), we next sought to determine if heme-binding or oligomerization is conserved across RoxP orthologs. Three RoxP orthologs (RoxP_5,6,8) with 81-82% sequence identity to RoxP_1 were initially shown (**Fig 4g**) to bind hemin via NATIVE gel-shift assays. RoxP_8 also demonstrated heme-binding by Soret peak analysis (**SI Fig 2f**).These findings led us to extend our SEC heme-binding assessment to include three additional RoxP orthologs (RoxP_4,7,11), which (**Fig 4h**) demonstrated that heme-binding is conserved across RoxP orthologs with sequence identity from 71-100%. Further, (**Fig 4h**) the increased MW of the holo-RoxP_4/5/7/11 complexes suggest that heme-dependent oligomerization is also conserved. Due to the conservation of heme-binding and heme-dependent oligomerization across seven RoxP orthologs (**SI Table 5**), we extended our DSF pH stability analysis and found that low pH stabilization was also conserved (**SI Fig 6**) for RoxP_5 (82% identity) and RoxP_11 (71% identity), as both were most stable at pH 4.8 and 4.5, respectively. These results suggested that RoxP function at low pH is highly conserved across RoxP orthologs.

**Figure 6.**
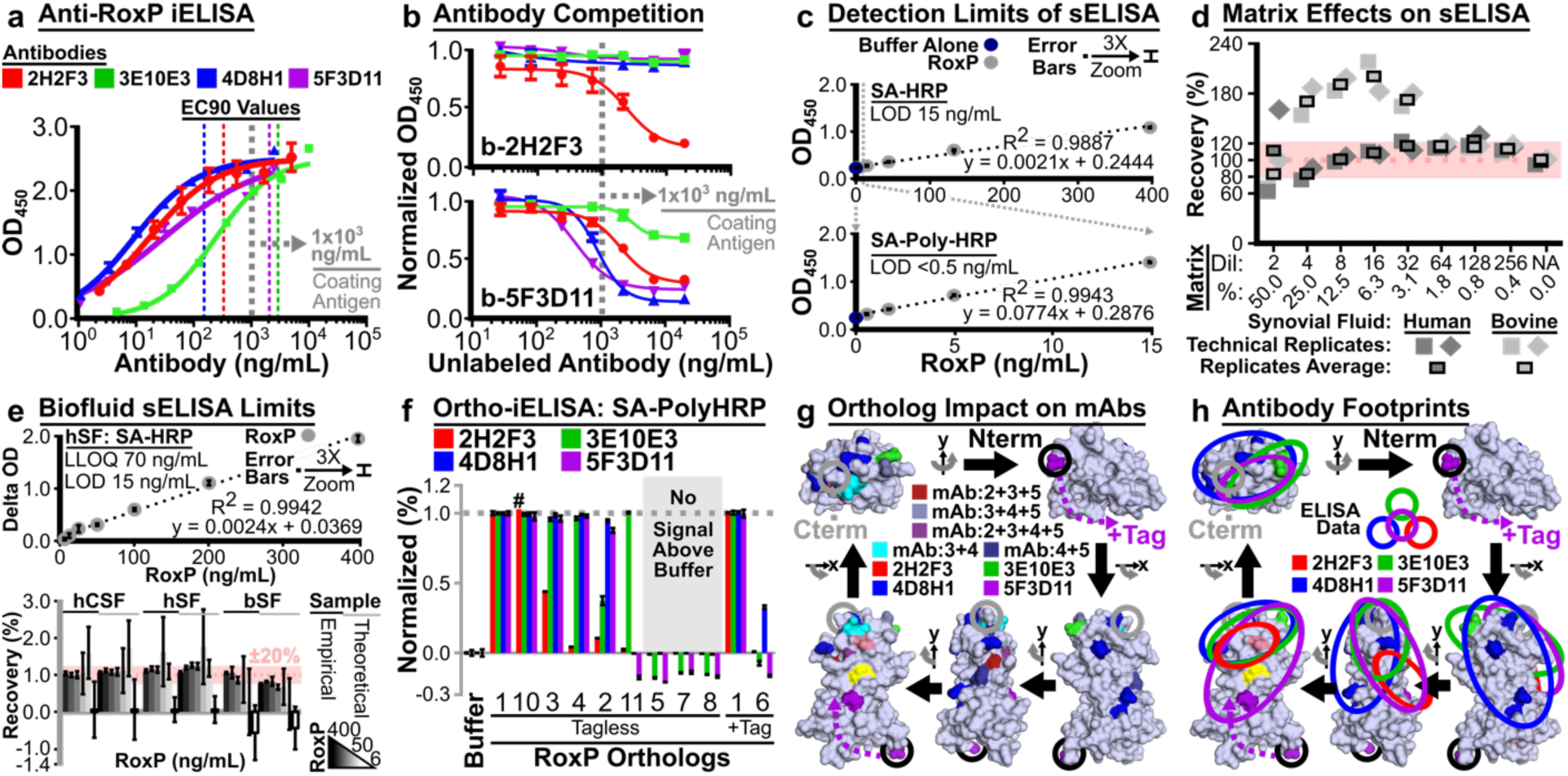
Anti-RoxP Immunoassay Development. (**a**) Indirect ELISA (iELISA) curves showing binding of recombinant apo-RoxP_1-tagless by antibodies 2H2F3, 3E10E3, 4D8H1, and 5F3D11. Two-layer detection: (1) biotinylated, goat anti-murine IgG, (2) streptavidin-horseradish peroxidase (SA-HRP). (**b**) Competition iELISA (ciELISA) curves showing RoxP detection by 2H2F3, 3E10E3, 4D8H1, and 5F3D11 antibodies (see **a** for mAb colors) in the presence of (*top panel*) biotinylated 2H2F3 (b-2H2F3) and (*bottom panel*) b-5F3D11. (**c**) Linear range of anti-RoxP sandwich ELISA (sELISA) using 4D8H1 (capture mAb), b-2H2F3 (detection mAb), and two versions of SA-HRP: SA-HRP, SA-Poly-HRP. All samples run with three technical replicates. (**d**,**e**) Horizontal, dotted red line: 100% recovery. Pink box: ± 20% recovery. (**d**) The spike-and-recovery sELISA (sar-sELISA) approach was used to determine the matrix effect of synovial fluid on the anti-RoxP sELISA. Matrix in each sample is shown as dilution (Dil) and (v/v) percentage (%). (**e**) After sample dilution (8-fold: hSF/CSF, 4-fold: bSF), a 2-fold RoxP dilution series (400-6.25 ng/mL) was used to determine the linearity (*top:* hSF) and limit of detection (*bottom:* hCSF, hSF, bSF) for the anti-RoxP sELISA in complex samples. (**f**) Ten recombinant RoxP orthologs were evaluated by iELISA (coating concentration: 10 µg/mL; terminal detection reagent: SA-Poly-HRP) in technical triplicate for binding by 2H2F3, 3E10E3, 4D8H1, and 5F3D11. See **SI Results** for explanation of negative signal. ^#^Marks a single RoxP_10-2H2F3 replicate due to two miss-seated micropipettor tips, but 100% 2H2F3 binding was observed for this ortholog under all other conditions (**SI Fig 15**). (**g**) Based on **SI Table 7**, RoxP ortholog variant positions that impact antibody binding were mapped to PDB 7bcj and colored according to which antibodies were impacted by a variant. For example, cyan residues (28-30) only affected 3E10E3 and 4D8H1 (3+4), while the violet-purple residue 89 affected all mAbs (2+3+4+5). (**h**) Antibody footprints were mapped onto RoxP_1 using variant positions (i) unique to each antibody (**g**,**h**) and (ii) all variants impacting an antibody (**SI Fig 16**). (**g**,**h**) Model x/y-axis rotations, RoxP_1/W66 surface coloring, and N-/C-termini circles are the same as in Fig 2. RoxP P91 is not surface exposed, so it’s location and effect on 2H2F3 binding is indicated by a red oval with 30% transparency in the bottom-left model. The proposed extension of the N-terminal tag causing partial inhibition of 5F3D11 (**6f**, **SI Fig 15**) is shown as “+Tag.” (**a**-**c**,**e**,**f**) Error bars represent Standard Deviation (SD).

### Molecular Modeling to Identify RoxP Heme-Binding Site

The high degree of RoxP functional conservation led us to assess what conserved structural determinants might support these functions. Phyre2 [67] was used to generate three-dimensional models of six RoxP orthologs (RoxP_4-8,11) that bind heme (**SI Table 5**). Electrostatic analysis in PyMOL [68] of these models (**Fig 5a**-**e**; **SI Fig 7a-c**,**8**) identified a cationic pocket immediately below the invariant tryptophan (RoxP_1 W66). This pocket contains three arginines (R56,121,123) and two tyrosines (Y110,116) that are invariant across all 21 RoxP orthologs. The positive charge in this pocket could provide an ideal environment to bind one of heme’s two negatively-charged propionic acid side chains at positions C(2) and C(18) of the porphyrin ring[69]. Unlike this area of charge conservation, there were also differences in charged surfaces leading to a range of predicted isoelectric points (pI 6.19-9.56), though these differences were primarily on the RoxP molecular surface opposite that of the conserved heme-binding site (**SI Fig 8**).

Next, we sought to identify RoxP’s heme axial ligand(s). Five different amino acids (C, H, K, M, Y) can serve as axial ligands to heme iron, though histidine is the dominant residue (∼80%) in both heme b and heme c types [70]. Several potential heme axial ligands were easily eliminated from consideration based on their (1) sequestration within a disulfide bond (C43,111) or (2) location (**Fig 2e**, **5f**) on the molecular face distant from of W66 (H23, K38, Y105, K130, M133). Further, several ortholog variants that introduced new potential heme ligands (**Fig 5g**-**j**, **SI Fig 7e,f**) could be eliminated because they were (1) not found in RoxP_1 or (2) >25 Å from W66. Then, an assessment of conservation of potential ligands across heme-binding orthologs (**Fig 5f**-**j**, **SI Fig 7d-f**) and the distance of these residues to W66 (**SI Table 6**) was able to reduce eight potential ligands to four probable ligands (H87>Y116,H98>H115). These residues encircle W66 at clock positions 1(H98), 5(H115,Y116), and 10(H87)-o’clock. Since the heme iron’s location is unknown, any of these four residues could serve as heme axial ligands. This analysis suggested that RoxP function requires a highly conserved molecular surface that could be targeted diagnostically to better (1) study *C. acnes* adaptation to human skin and (2) identify *C. acnes* infections of humans.

### Development of Anti-RoxP Immunoassays

#### Indirect Enzyme Linked Immunosorbent Assay (iELISA)

To further investigate RoxP function, we generated four murine, IgG, anti-RoxP antibodies (2H2F3, 3E10E3, 4D8H1, 5F3D11) against RoxP_1-tagless, and then validated their specificity using an anti-RoxP iELISA (**Fig 6a**). This assay revealed that three antibodies had a higher affinity for RoxP_1 (4D8H1>2H2F3>5F3D11 >>3E10E3). To simplify anti-RoxP immunoassay development, these antibodies were biotinylated and then validated by iELISA (**SI Fig 9**), which revealed that biotinylated-3E10E3 (b-3E10E3) is not able to detect RoxP_1. Sequencing of the fragment variable (Fv) regions of these antibodies (**SI File 5**) suggested that b-3E10E3 was biotinylated at a complementarity determining region (CDR) lysine required for antigen binding.

#### Competition iELISA (ciELISA)

Biotinylated antibodies (b-2H2F3, b-4D8H1, b-5F3D11) were then used in ciELISAs against a titration of all four unlabeled antibodies (**Fig 6b**; **SI Fig 10**). While multiple antibodies could compete against b-5F3D11, b-2H2F3 and b-4D8H1 were only out competed by their unlabeled counterparts. The 5F3D11 competition data suggested that the binding site of this antibody sterically overlaps with the other three antibodies, while the lack of competition between 2H2F3 and 4D8H1 suggested that these antibodies bind non-overlapping epitopes.

#### Sandwich ELISA (sELISA)

These non-competitive antibodies (4D8H1, 2H2F3) were then evaluated in both orientations to develop an anti-RoxP sandwich ELISA (sELISA). The best orientation (capture: 4D8H1; detection: biotinylated-2H2F3 [b-2H2F3]) was shown to be linear and to have a limit of detection (LOD) that could be enhanced >30-fold from 15 to <0.5 ng/mL RoxP_1 (**Fig 6c**). To determine if this sELISA could be used to identify *C. acnes* infections, we performed spike-and-recovery sELISA (sar-sELISA) in human synovial fluid (hSF), cerebrospinal fluid (CSF), serum; bovine synovial fluid (bSF); and *C. acnes* growth media (**Fig 6d, SI Fig 11**). Unlike hSF, the bSF matrix effect increased between sar-sELISA experiments (#1: **SI Fig 11d**; #2: **Fig 6d**), which suggests the intervening biofluid storage at -20°C affected bSF but not hSF. Matrix effects identified by sar-sELISA demonstrated that all complex fluids required dilution (1/4 or 1/8) to recover 100±20% of the spiked in RoxP_1 with stable recovery upon further dilution.

Using this dilution information, sELISA linearity of dilution, LOD, and lower limit of quantitation (LLOQ) were assessed in all six complex fluids (**Fig 6e**, **SI Fig 12**). Using the SA-HRP sELISA approach, the assessed LOD ∼15 ng/mL and LLOQ ∼70 ng/mL were consistent with the percent recovery limit of 50-100 ng/mL, which would be expected to be enhanced >30-fold to 2-3 ng/mL with the use of SA-Poly-HRP. The anti-RoxP sELISA was then used to assess endogenous RoxP secretion from mid-log RCM cultures (n=5) of two *C. acnes* reference strains (KPA171202: 65.52+26.09 ng/mL, ATCC 6919 23.58+6.67 ng/mL). Prior literature noted that stationary phase culture of KPA171202 contained ∼10 μg/mL RoxP [47], which suggests that RoxP accumulates during culture and may be expressed at higher levels during stationary phase. As such, this sELISA appears to be able to assess RoxP secretion both *in vitro* and *in vivo*.

#### Ortholog Specificity of Anti-RoxP Immunoassays

Though RoxP is highly conserved, our analysis of non-conservative amino acid solvent exposure on RoxP (**Fig 2b**) suggested that sequence variation might inhibit anti-RoxP antibody recognition of some clinically-relevant RoxP orthologs (e.g., **SI Table 2/3**: RoxP_2 – Asn12, RoxP_5 – ATCC 11828). Ten RoxP orthologs (RoxP_2-11) that represented a wide range of variation compared to RoxP_1 (71-99% identity, 1-39 amino acid differences, **SI Fig 13**) were selected to determine the ortholog specificity of our anti-RoxP antibodies. Tagless, recombinant RoxP was prepared using a small-scale protocol (**SI Fig 14**) and then assessed by anti-RoxP iELISA (**Fig 6f; SI Fig 15**). While RoxP orthologs with >90% identity to RoxP_1 were bound by many antibodies, even a small number of variant positions could impede antibody binding. For example, RoxP_2 has 3 variant positions, which result in significant inhibition of both 2H2F3 and 3E10E3 binding. With greater sequence divergence from RoxP_1, the success rate of ortholog detection by anti-RoxP antibodies decreased, but not all antibodies were impeded by significant differences in ortholog sequence identity to RoxP_1 (e.g., RoxP_6, 81%; RoxP_11, 71%).

#### Anti-RoxP Antibody Footprinting Analysis

Variable ortholog recognition gave us the opportunity to perform antibody footprinting to localize anti-RoxP epitopes. Antibody binders were compared to partial-/non-binders to identify variants near/within antibody footprints (**Fig 6f**, **SI Fig 15**, **SI Results**). Those ortholog variant positions (**SI Table 7**) were mapped to the apo-RoxP_1 NMR structure (PBD 7bcj) (**Fig 6g**, **SI Fig 16**) and used to identify potential antibody binding footprints (**Fig 6h**, **SI Results**). These antibody footprints largely reproduced our ciELISA findings (**Fig 6b**, **SI Fig 10**) and are supported by CDR conservation between antibodies (**SI Results, SI File 5-12**). Interestingly, all four antibodies were affected by G89, which effectively pinned their footprints to a single location near the C-terminus (**Fig 6g**). While three of these antibodies have footprints that appear to overlap the C-terminus (3E10E3/4D8H1>5F3D11), the 5F3D11 footprint appears to extend away from the C-terminus towards the anti-oxidant tyrosine loop. The antibody footprint locations relative to W66 and the conserved cationic pocket also suggest that heme-binding may affect 5F3D11>2H2F3>3E10E3>4D8H1 (**Fig 6h**, **SI Fig 16**).

Since an antibody typically buries ∼500 Å^2^ of antigen SASA [71,72], it is quite remarkable that these four antibodies have distinct footprints on a protein as small as RoxP (SASA 8,836.82 Å^2^). Even more striking is the fact that two of these antibodies could be used to develop an anti-RoxP sELISA. Further, the arrangement of these four antibodies on RoxP now provide a unique opportunity for future *in vitro* studies of RoxP function, as well as the development of *Cutibacterium*-specific clinical diagnostics and antibody-based therapeutics.

## DISCUSSION

RoxP appears to be a Cutibacteria adaptation to human skin that arose after they had immigrated from cows to humans but before *C. acnes* emerged to dominate human sebaceous skin [2,47]. As such, RoxP may have played a key role in establishing this microbiome and helped guide host-microbe co-evolution during the transition of humans from hunter-gather to agrarian communities. Since no other organism encodes a RoxP homolog, our group decided to perform an in-depth analysis of RoxP sequence space, biochemistry, and function. We then developed immunoassays to measure RoxP in culture media and mammalian biofluids, so that we can investigate this protein’s role in *C. acnes* biology and aid in the identification of *C. acnes* infections. Through this work, we identified high conservation of RoxP solvent-exposed surfaces, ligand-dependent oligomerization, and low pH stability. This analysis has begun to shed light on RoxP’s function and helps us to understand the unique needs of a PSU microbe. Notably, RoxP low pH thermal stability also helps to suggest how this protein has guided host-microbe co-evolution.

### Life on Humans Favors RoxP Conservation

Cutibacteria evolution [2] appears to have preceded from older species that do not encode *roxP* (*C. granulosum*/*avidum*) to intermediary ancestors with high pI RoxP (*C. modestum/namnetense* [2,73,74]). They then evolved into *C. acnes* and split into subspecies with high pI RoxP (*C. acnes* subsp. *defendens*) or low pI RoxP (*C. acnes* subsp. *acnes/elongatum*). *C. acnes* significantly contributes to the low pH of human skin [8], and the dominant RoxP ortholog on human skin has a much lower pI (RoxP_1, pI 6.19) than many other orthologs. Based on our analysis of RoxP orthologs (**Fig 1c**, **SI Table 2**), 93% of *C. acnes* subsp. *acnes* RoxP orthologs (N=90) have this pI, while 76% of *C. acnes* subsp. *defendens* (N=55) have a pI >8.7. *C. acnes* subsp. *elongatum* has an even lower pI of 5.90, though it is not associated with acne and seems to preferentially inhabit the less acne-prone skin of the lower back [75,76,48].

Strikingly, a comparison of *C. acnes* phylotypes in acne- and healthy-skin [77] suggests that decreasing the burden of *C. acnes* that encode low pI RoxP is correlated with healthy skin. Thus, the low pH environment created by *C. acnes* may have supported the adaptation of a pathobiont subspecies (*C. acnes* subsp. *acnes*) that contributes to the human disease acne vulgaris. Interestingly, the only known acne vulgaris cure (isotretinoin) raises skin pH by >0.5 pH [78], which may support the expansion of healthy *C. acnes* (phylotype II, *C. acnes* subsp. *defendens* [77]) that encode high pI RoxP. In support of this hypothesis, 67% of *C. acnes* subsp. *defendens* encode RoxP_5, which is more stable at neutral pH (T_ma_ +8.1°C) than RoxP_1 (**Fig 4d**-**f**, **SI Fig 6**).

While it has been established that *C. acne*s requires RoxP to colonize human skin [47], it may have also contributed to *C. acnes* dominance of sebaceous skin. *C. acnes* constitutes ∼50% of skin surface microbes [11,13] and ∼90% of PSU microbes [12,79]. Thus, this unique microbial tool may have given *C. acnes* a survival advantage that helped it secure its place as the dominant member of the sebaceous skin microbiome.

### RoxP Conservation is a C. acnes Achilles’ Heel

RoxP sequence space (946 entries) is dominated by RoxP_1 (67% of entries) and a limited number of other orthologs with similar sequence identity (**Fig 1**). In fact, 80% of RoxP (RoxP_1,16,17,20) are >99% identical. Further, all *C. acnes* phylotypes have been observed to secrete RoxP (**SI Table 4**), which suggests that this protein is a *Cutibacterium*-specific growth biomarker that can be targeted diagnostically and therapeutically. For such a strategy to work, it would be ideal for anti-RoxP reagents to bind a diversity of RoxP orthologs.

Based on **Fig 6f**, five RoxP orthologs (RoxP_1/10>3>2>4; 90-100% identity) should be detected by our sELISA antibody pair 2H2F3/4D8H1. The sELISA detection limits for each ortholog will likely vary based on their 2H2F3 affinity, so optimization may be necessary for low-affinity orthologs (e.g., RoxP_2-4). RoxP_17 and RoxP_20 are also likely to be recognized by the sELISA due to the antibody-binding permissibility of their variants G89E and T34I (**SI Table 7**), respectively. On the other hand, RoxP_16’s single variant position S20F likely precludes sELISA recognition, as it appears that RoxP_11 is not recognized by 4D8H1 due at least in part to S20I.

RoxP orthologs with low 2H2F3/4D8H1 affinity (RoxP_2-4,6; ∼1% of RoxP entries) could also be readily assessed by converting the sELISA into a competition sELISA where the standard curve would be a fixed concentration of RoxP_1 titrated against recombinant RoxP_2-4,6. For this reason, we conclude that some version of our sELISA should be able to detect 69% of all RoxP and 72% of *C. acnes* specific-RoxP IPG sequences (RoxP_1-4,6,10). Further, the reported sELISA should detect 67% of all RoxP, 71% of *C. acnes*-specific RoxP, and 93% of *C. acnes* subsp. *acnes*-specific-RoxP IPG sequences (RoxP_1/10).

The development of this type of assay is critically needed, because *C. acnes* is an emerging pathogen that causes the majority of shoulder PJIs, as well as many other IMD infections. Yet, *C. acnes* infections are frequently not identified due to an inability of current diagnostics to quickly and clearly differentiate a “true” *C. acnes* infection from *C. acnes* contaminants. The latter concern arises from *C. acnes* ubiquitous nature, as a false-positive culture result might arise from a patient’s own skin, the skin of a close contact (e.g., treating surgeon), or the environment. Recently, two groups have analyzed the *C. acnes* genomes from PJIs (shoulder, elbow, hip, knee; 17 patients; 27 isolates) [80,81], and our analysis of *roxP* in their genomes (**SI Table 2**) revealed that 80-92% of *C. acnes* PJIs include a strain that encodes RoxP_1. Of note, two closely-related orthologs (RoxP_5/21; 99% identical; ∼82% identity to RoxP_1) that are not recognized by our antibodies were also present in 40% of “infection-likely” *C. acnes* PJIs (5 patients,15 isolates) [81]. Regardless, this genomic analysis indicates that the ability of our anti-RoxP sELISA to identify *C. acnes* IMD infections is likely higher than that predicted by our evaluation of the total RoxP sequence space.

### Other C. acnes Antibodies Are Not Amendable to Clinical Care

Though *C. acnes* antibodies have been previously described (**SI Table 8**) [82–86], there is no *C. acnes* immunoassay used in routine clinical practice due to several limitations. For example, *C. acnes* antibody PAB is commercially-available, but it recognizes a cell-surface antigen that has limited its use to immunohistochemistry (IHC) [86]. Further, PAB has only been used to stain acid-fast bodies in sarcoid lymph nodes, and no group has reported using it to identify infection. Another *C. acnes* antibody (QUBPa4) [85] binds the secreted protein Christie–Atkins–Munch– Petersen (CAMP) factor 1 [85,87,88], but CAMP factor 1 does not appear to be secreted by all *C. acnes* strains [89]. There is also significant inter-strain variation in the expression of *C. acnes* CAMP factors 1-5 [85]. In comparison, RoxP secretion has been observed to occur in all *C. acnes* phylotypes [90] and all assessed *C. acnes* strains (**SI Table 4**).

Recently, a Luminex assay using anti-*C. acnes* polyclonal antibody (pAb) was used to assess 94 *C. acnes*-culture positive hSF samples [91]. *C. acnes* antigen was detected (signal/cutoff >1) in the majority of these samples (65%). Further, this assay only detected *C. acnes* antigen in a small percentage of the two negative-control groups: (1) infected by other bacteria (5.8%, N=103), (2) low probability of infection (0.28%, N=1050). Interestingly, the immunogen for this polyclonal antibody was *C. acnes* strain ATCC 11827, which encodes RoxP_1 (**SI Table 2**). Unfortunately, the nature of this immunization (e.g., intact cells, supernatants, lysate) was not reported, so it is unclear if one of the pAb targets is RoxP_1.

Another group has generated an anti-RoxP pAb and used it to develop a competition ELISA with a LOD of >80 ng/mL in PBS. This assay can detect RoxP_1 recombinantly produced by *Pichia pastoris* [92] and RoxP_1 endogenously secreted by *C. acnes* strain KPA171202 [93]. Due to the nature of a polyclonal antibody response, this assay may be able to detect a range of RoxP orthologs, though the authors did not evaluate any other RoxP orthologs. Further, pAb assays can suffer from batch-to-batch pAb variability, which can impede their use. In comparison to this pAb anti-RoxP assay, the monoclonal anti-RoxP sELISA described in this manuscript (1) has >150-fold higher sensitivity, (2) can detect RoxP in multiple human biofluids, and (3) has been evaluated against a diverse array of ten RoxP orthologs (71-100% identity). As such, our assay appears to be more suitable as a clinical diagnostic where the RoxP concentration and ortholog are unknown.

### Other RoxP Assays Are Not Amendable to Clinical Care

RoxP detection has also been reported using capacitive [94] and surface plasmon resonance (SPR) [93] biosensors that were created using a molecular surface imprinting technology. Though the SPR assay is quite sensitive (3.68 ng/mL), these approaches are not yet clinically feasible due to methodological limitations (e.g., severe mass transport in complex samples) and the absence of biosensor equipment in clinical labs. In comparison, ELISAs are used in clinical laboratories worldwide, which makes our sELISA easily implementable in established clinical workflows. Further, our sELISA is (1) >7-fold more sensitive than the RoxP SPR assay and (2) could be converted to a lateral flow immunoassay to create a point-of-care (POC) assay for *C. acnes* infection detection.

### Future Impact of RoxP Immunoassays

This manuscript’s immunoassays target a protein secreted only by Cutibacteria, which provides us the unique opportunity to improve healthcare and our understanding of microbe evolution. Use of these assays in clinics would help to address widespread confirmation bias where *C. acnes* cultures are discounted due to the (1) growth of a “more likely” infectious organism (e.g., staphylococci) or (2) assumption that *C. acnes* must be a contaminant. Further, identification of *C. acnes* infections using these assays would lead to appropriate treatment far sooner than awaiting the slow growth of a *C. acnes* culture (5-14 days) [42]. Notably, such treatment might include targeting both *C. acnes* and staphylococci, as these microbes can live in close proximity to one another in the PSU [28] and have been co-cultured from IMD infections [35,82,30,43,31].

Beyond the potential impact of these assays on healthcare, their use in future studies will help us understand RoxP’s role in *C. acnes* (1) biology and (2) interactions with other microbes. Such work will expand our understanding of host-microbe co-evolution and thus characterize the ecological shift(s) that occurred in a human microbiome upon *C. acnes* arrival to the PSU. To support this future work, our group is already developing reagents that recognize RoxP orthologs that escape detection by our current assays, so that growth by *C. acnes* subsp. *defendens*/*elongatum*, *C. modestum*, and *C. namnetense* can be assessed both *in vitro* and in humans. In summary, future work utilizing this paper’s anti-RoxP immunoassays will shed light on the burden of *C. acnes* infections, what role RoxP plays in human disease, and how *C. acnes* arose to dominate human sebaceous skin microbiomes.

## CONCLUSIONS

Herein, we have begun to dissect the molecular determinants of the protein RoxP. This protein helped dairy Propionibacteria move from simply adapting to live on human skin (*C. avidum*/*granulosum*) to occupying 25% of the skin and dominating the PSU. Reagents developed as a part of this study have created a highly sensitive sELISA that should be able to detect ∼90% of *C. acnes* infections of human biofluids. This initial RoxP work has led to a better understanding of commensal microbe adaptation, while providing tools to investigate pathobiont evolution and identify human *C. acnes* infections.

## ACKNOWLEDGEMENTS AND FUNDING

Acknowledgements

We thank William E. Phillips, D.O. (laboratory technician) for project support including preparation of RoxP_1 protein stocks and training of other lab members to express/purify RoxP_1.

## Core/Center Support

Research reported in this publication was supported by the following university core/center facilities and the indicated core/center personnel:

1. University of Connecticut (UCONN) Center for Open Research Resources and Equipment (COR2E) – Biophysics Core. Experiments performed and analyzed by the Biophysics Core Director (Heidi Erlandsen, Ph.D.).
2. WUSM High Throughput Screening Center (HTSC). Experiments designed/performed by Michael J. Prinsen and designed/analyzed by the HTSC Director (Ma Xenia Garcia Ilagan, Ph.D.).

## Funding

Research reported in this publication was supported by the following sources. The content is solely the responsibility of the authors and does not necessarily represent the official views of the National Institutes of Health (NIH), Department of Health and Human Services (DHHS), National Library of Medicine, or any of the other funding agencies listed below. The funding organizations had no role in design and conduct of the study; collection, management, analysis, and interpretation of the data; preparation, review, or approval of the manuscript; and decision to submit the manuscript for publication.

### WHM

National Center for Advancing Translational Sciences (NCATS) of the NIH under Award Number KL2 TR002346; National Institute of Arthritis, Musculoskeletal and Skin Diseases (NIAMS) of the NIH under Award Number 5K08AR076464; 2019 Longer Life Foundation Grant

### JZT, MRV

Washington University in St. Louis School of Medicine, Office of Medical Student Research, Dean’s Funding for Summer Research Program (SRP).

## METHODS

Previously described protocols were adapted to express recombinant proteins using autoinduction [95]; prepare previously described recombinant proteins (6His-TEV[96], hSCNkk[97]); assess protein sequence conservation [98–101]; use protein sequences to predict protein biochemical characteristics [102–104]; analyze and predict protein structures [105–107,100,108,68]; and perform biophysical experiments (AUC [109–111], DSF [112–114]).

Please see **SI Methods** for further details on these experiments.

## Supplementary Information (SI) Results

### Detailed Anti-RoxP Antibody Footprinting Analysis

RoxP ortholog variant positions (**SI Fig 13**) from ten soluble, recombinant RoxP proteins (tagless: RoxP_1-5,7,8,10,11; +Tag: RoxP_1,6) that were tested in anti-RoxP indirect enzyme-linked immunosorbent assay (iELISA: **Fig 6f**, **SI Fig 15**) were mapped to the apo-RoxP_1 NMR structure (PBD 7bcj). Then, variant positions near/within antibody footprints were identified by comparing antibody binders to partial-/non-binders. The effect of RoxP ortholog variant positions on antibody binding are summarized in **SI Table 7** and shown in **SI Fig 16**.

For example, RoxP_10 demonstrated that T69A had a modest effect on 5F3D11 binding (**SI Fig 15b**). T69A is a conservative variant shown in grey (partially SASA accessible) in the middle of the bottom-left model of **Fig 2f**. The effect RoxP_10 T69A on 5F3D11 binding was similar to the effect of including the N-terminal tag (**SI Fig 15b**), which is on the same side of RoxP (bottom-right of RoxP in the bottom-left view of all **Fig 2** RoxP models). Since the N-terminal tag is predicted to be random coil and long enough (e.g., 20 amino acids can extend ∼72Å) to potentially interact with T69 and adjacent residues, it seems possible that the tag’s effect on this antibody is direct antibody inhibition, rather than an effect on RoxP coating to the ELISA plate. The latter situation may explain the lower signal for all four antibodies for RoxP_1+Tag observed in **SI Fig 15a**, though experimental variability in the measurement of protein concentration (e.g., variation within ∼2.5-fold) might also explain those results. Regardless, 5F3D11’s binding for RoxP_1+Tag is still significantly lower in **SI Fig 15a**, which supports an effect of the tag on that antibody’s ability to bind RoxP.

Then, a comparison of 4D8H1 binding of RoxP_2 and RoxP_3 suggested that S31 is required for 4D8H1 binding, as it nearly returns iELISA signal to RoxP_1 for RoxP_2 (**SI Fig 15a**). Further, these two orthologs have three variant positions relative to RoxP_1, and they share two of those positions (G89E, S100N). Since RoxP_2 has the T34A variant, antibody 4D8H1 seems to be insensitive to this substitution. That suggests that RoxP_2/3 G89E and S100N may also contribute to reduced 4D8H1 signal in **SI Fig 15a**, though it appears that those variants have a lesser impact on 4D8H1 binding than S31N. It is unclear why the 4D8H1 signal for RoxP_3 increases to a greater extent than RoxP_2 from **SI Fig 15a** to **SI Fig 15b**. This assessment supports S31>G89/S100 as near/within the 4D8H1 antibody footprint.

On the other hand, 5F3D11 binding was impeded more than 4D8H1 binding by RoxP_2/3 variant positions (**SI Fig 15a**), which suggests that RoxP_2/3 variants G89E/S100N have a greater effect on 5F3D11 binding than 4D8H1 binding. 5F3D11 also bound RoxP_2 to a greater extent than RoxP_3 (**SI Fig 15a**,**b**), which suggests that (1) S31 contributes to 5F3D11 binding and (2) T34A may modestly inhibit 5F3D11 binding. With those observations in mind, a comparison of 4D8H1 non-binders RoxP_5,7,8,11; very weak binder RoxP_6; and binders RoxP_1-4,10 identified positions RoxP_5/6 S20P; RoxP_11 S20I; RoxP_7/8 A28T; RoxP_5/6 A28S, A29T, S30A; RoxP_5-8,11 T36K; RoxP_4-6,11 M133T; RoxP_7/8 M133A, D135N; RoxP 4-6 L138I; RoxP_7/8 L138A; and RoxP_11 L138K as potentially impeding 4D8H1 binding (**Fig 6f**). Of note, a comparison of 4D8H1 binding to RoxP_6 to the non-binder RoxP_5 suggests that RoxP variants G12D and Q14R enhance 4D8H1 binding, as those are the only differences between these orthologs.

A comparison of 3E10E3 non-binders RoxP_5-8, weak binders RoxP_3/4>2, and strong binders RoxP_1,10,11 identified RoxP_5/6 A28S, A29T, S30A; RoxP_2 T34A; RoxP_2/3 G89E, S100N; RoxP_4-6 S100G; RoxP_7/8 G89D, S100D; and RoxP_4-8 E101K as likely impeding 3E10E3 binding (**Fig 6f**, **SI Fig 15**). 3E10E3 footprinting was greatly aided by 3E10E3 displaying equivalent binding for RoxP_1 and RoxP_11. RoxP_11 is the most divergent RoxP ortholog (71% identity), yet it has significant amino acid similarity with RoxP_1 residues 28-30, 34, 100-101. Further, 3E10E3 binds RoxP_11 despite RoxP_11’s variant T36K, which is also found in RoxP_5-8 that do not bind 3E10E3.

With the above assessment in mind, a comparison of 2H2F3 non-binders RoxP_5-8; very weak binders RoxP_2>4>>11; and binders RoxP_1,10>>3 identified RoxP_2 T34A; RoxP_2/3/7/8 G89D/E; RoxP_4-8,11 P91A as likely impeding 2H2F3 binding (**Fig 6f**, **SI Fig 15**). The P91A variant may extend the RoxP helix 1 by one full turn impeding 2H2F3 binding. Then, a comparison of **SI Fig 15b** with **Fig 6f** for 2H2F3 and 3E10E3 binding suggests that RoxP_2 T34A impedes 2H2F3 > 3E10E3 binding.

While all variant positions that affect anti-RoxP antibody binding were considered when localizing potential antibody binding footprints (**Fig 6h**), some residues were given greater weight in initially centering a predicted footprint based on their side-chain surface exposure, the nature of the change in the chemical landscape (e.g., a charge reversal mutation, like G89D/E), likelihood of a change impacting protein secondary structure (e.g., substitution of a proline residue, like P91A), or greater incidence/severity of a variant in non-binders than weak/partial binders and exclusion from binders. Further, the ability of antibodies to compete for RoxP binding (**Fig 6b**; **SI Fig 10**) was used to ascertain relative orientation to one another on the RoxP 7bcj surface. For example, the 2H2F3 antigen-binding footprint (**Fig 6h**) was localized using non-conservative variation of G89 (**Fig 2b,c** – upper-left, lower-left, and lower-middle models; **SI Fig 16a** – from top-to-bottom:1^st^,4^th^, and 5^th^ models) and P91 (buried residue, variation changes surface contour; **SI Fig 16a** – bottom two models). The 3E10E3 antigen-binding footprint (**Fig 6h**) was localized using non-conservative variation of S100 and E101 (**Fig 2b,c** – upper-left, lower-left, and lower-right models; **SI Fig 16b** – top, middle, and bottom models). The 4D8H1 antigen-binding footprint (**Fig 6h**) was localized using non-conservative variation of S20 (**Fig 2b,c** – upper-left, lower-left, and lower-middle models; **SI Fig 16c** – top and bottom two models) and mixed variation of L138 (**Fig 2b,f** – upper-left, lower-middle, and lower-right models; **SI Fig 16c** – from top-to-bottom: 1^st^, 3^rd^, and 4^th^ models), as well as antibody-specific variants at positions 12, 14, 133, and 135. The 5F3D11 antigen-binding footprint (**Fig 6h**) was localized using mixed variation of T34 (**Fig 2b,f** – upper-left, lower-left, and lower-middle models; **SI Fig 16d** – top and bottom two models) and the unique inhibitory effect of T69A and the N-terminal tag on 5F3D11 binding (**SI Fig 16d** – from top-to-bottom: 2^nd^, 4^th^, and 5^th^ models), which rotated the footprint down from the cluster of shared antibody-inhibiting variant positions (e.g., S31, T34, T36, G89, S100) shared with the three antibodies that it competes with by ciELISA (**Fig 6b**; **SI Fig 10**). After initial antibody footprint placement, each footprint was adjusted (e.g., size, rotation) to capture all implicated variant positions (**SI Table 7**, **SI Fig 16**).

### Comparison of Anti-RoxP Antibody CDR Use

Multiple sequence alignment (MSA) using ClustalW[2] in UGENE[3] for each of the six complementarity-determining region (CDR) gene segments (**SI File 5-12**) identified several similarities that may contribute to overlapping RoxP epitopes. Heavy chain CDR1 (hc-CDR1) was highly conserved between 3E10E3/5F3D11 (57% identity, 86% similarity) and partially conserved between 2H2F3/4D8H1 (40% identity), while light chain CDR1 (lc-CDR1) was highly conserved between 2H2F3/5F3D11 (69% identity, 75% similarity) but only weakly conserved for 3E10E3/4D8H1 (27% identity). Low identity/similarity (e.g., 3E10E3/4D8H1 lc-CDR1) is likely due to use of the same parent CDR1 sequence without necessarily linking to recognition of the same RoxP epitope feature(s). CDR2 alignments also revealed high conservation for hc-CDR2 3E10E3/5F3D11 (59% identity, 65% similarity) and lc-CDR2 2H2F3/5F3D11 (100% identity). On the other hand, an alignment of CDR3 sequences only revealed high conservation of lc-CDR3 2H2F3/5F3D11 (56% identity, 67% similarity), though hc-CDR3 2H2F3/5F3D11 also likely arose from the same parent hc-CDR3 sequence, as they were 27% identical. While this findings might only be due to a restricted set of CDR sequences that expanded during apo-RoxP_1-tagless antigen priming, the high degree of similarity for antibodies that compete with one another (2H2F3/5F3D11 – 3/6 CDRs, all 3 lc-CDRs; 3E10E3/5F3D11 – 2/6 CDRs, hc-CDR1/2) and low similarity for non-competitive antibodies (2H2F3/4D8H1 – hc-CDR1) suggest that CDR conservation is linked to the recognition of specific RoxP epitope features.

### RoxP Orthologs with Decreased Anti-RoxP Antibody Binding Compared to Buffer Alone

Several high theoretical isoelectric point (pI) RoxP orthologs (RoxP_5-8) displayed signal for 3E10E3 and 5F3D11 that was lower than buffer-alone (**Fig 6f**, **SI Fig 15b**). A high pI protein would be positively charged (see unique blue surface on the middle row models of **SI Fig 8c**-**f** as compared to the middle row model of **SI Fig 8a**) at neutral pH used in PBS. An assessment of the complementarity-determining regions (CDRs) of all anti-RoxP antibodies (**SI File 5**) suggests that this phenomenon may be due to positive-positive charge repulsion from three potential antibody sources. First, hc-CDR2s of antibodies 3E10E3 and 5F3D11 are 63% identical with conservation of three positively-charged residues (H,K,K). Second, these two antibodies each have two CDRs with ≥3 positively-charged residues, while antibodies 2H2F3 and 4D8H1 have ≤2 positively-charged residues in their CDRs. Third, antibodies 3E10E3 and 5F3D11 have side-by-side positively-charged residues within their CDRs (3E10E3 lc-CDR3 HH, 5F3D11 hc-CDR2 KH). Beyond RoxP_5-8 potentially repelling antibodies 3E10E3 and 5F3D11 through charge-charge interactions, these orthologs may also have directly impeded SA-HRP binding, as HRP has an experimentally determined pI of 8.7-9[1] (i.e., positively-charged at neutral pH) and the signal for 3E10E3 and 5F3D11 that was lower than buffer-alone was exacerbated by the use of SA-Poly-HRP (compare **SI Fig 15b** to **Fig 6f**).

## Supplementary Information (SI) Methods

### General Information

A table of key resources is provided as **SI Table 9**. All centrifugation steps are listed as “xg”, which corresponds to the relative centrifugal force (RCF) used on each centrifuge. For percentage concentrations, “m/v” refers to mass concentration (e.g., reagent mass divided total volume), while “v/v” refers to volume concentration (e.g., reagent volume divided by total volume). Unless otherwise noted, all solutions/stocks were aqueous and prepared using double-distilled H_2_O (ddH_2_O).

### Abbreviations

A_280_ (absorbance at 280 nm); diH_2_O (distilled water); ɛ280 (extinction coefficient at 280 nm); EtOH (ethanol); g (gram); hr (hour); in. (inch); mAU (milli-absorbance unit); min (minute); PES (polyethersulfone); rpm (revolutions per minute); RT (room temperature, ∼22°C); #x (numerical concentration factor for a solution)

### Microbial Media, Reagents, and Consumables

Lysogeny broth (LB) – Lennox formulation (5 g/L NaCl) was prepared per the manufacturer’s recommendations (SIGMA: 20.0 g LB in 1 L of distilled water). ZYM-5052-Carb media was prepared as described previously [1]. Brain Heart Infusion (BHI) media (Becton Dickinson) was prepared by dissolving 37.0 g powder in 1.0 L water. Reinforced Clostridial Medium (RCM) was prepared by dissolving 10.0 g of peptone, 10.0 g of beef extract, 3.0 g of yeast extract, 5.0 g of dextrose, 5.0 g of sodium chloride, 1.0 g of soluble starch, 3.0 g of sodium acetate, 0.5 g of l-cysteine in 1.0 L of deionized distilled water. The RCM broth was adjusted to a final pH of 6.8. Overnight 5.0 mL cultures were grown in 14 mL Falcon tubes (Corning # 352059).

## SOLUTIONS

### General Information – Solution Assembly

The general assembly of essential solutions for this work is outlined below. Unless otherwise stated, all solutions in this work were assembled using double-distilled H_2_O (ddH_2_O), which refers to Type I – Ultrapure Water. Type I water is defined by the American Society for Testing and Materials (ASTM) as >18 MΩ-cm, conductivity <0.056 µS/cm, and <50 ppb of Total Organic Carbons (TOC). All dry chemical additions from polystyrene weigh boats included a ddH_2_O wash bottle rinse of the weigh boat into the solution. Flick-to-mix refers to using a finger to flick a microcentrifuge tube to gently mix its contents without introducing shear forces, like those introduced by vortexing, that can denature proteins and other sensitive reagents.

#### Assembly of 1.0 L of 1x Solution from a 10x Solution

Add 0.9 L ddH_2_O to a 1.0 L graduated cylinder, use a 0.1 L graduate cylinder to add 0.1 L of 10x Solution to the 1.0 L graduated cylinder, cover the top of the cylinder with parafilm (≥3 layers), firmly press a palm onto the parafilm to seal the cylinder, invert the cylinder ten times create a homogeneous solution, remove the parafilm from the cylinder.

#### Assembly of 1.0 L of 1x Solution from a 20x Solution

Add 0.95 L ddH_2_O to a 1.0 L graduated cylinder, use a 0.1 L graduate cylinder to add 0.05 L of 20x Solution to the 1.0 L graduated cylinder, cover the top of the cylinder with parafilm (≥3 layers), firmly press a palm onto the parafilm to seal the cylinder, invert the cylinder ten times create a homogeneous solution, remove the parafilm from the cylinder.

#### Assembly of 2.0 L of 1x Solution from a 20x Solution

Add 1.90 L ddH_2_O to a 2.0 L graduated cylinder, use a 0.1 L graduate cylinder to add 0.1 L of 20x Solution to the 2.0 L graduated cylinder, cover the top of the cylinder with parafilm (≥3 layers), firmly press a palm onto the parafilm to seal the cylinder, invert the cylinder ten times create a homogeneous solution, remove the parafilm from the cylinder.

### Immobilized Metal Affinity Chromatography (IMAC) Solutions

#### Gravity Flow Chromatography – IMAC Solutions

##### 10x Nickel-NTA (NiNTA) LYSIS Buffer (NiNTA-LB)

0.5 M Na phosphate, 3.0 M NaCl, 0.1 M Imidazole, pH 8.0

#### Assembly of 1.0 L of 10x NiNTA-LB

Add 0.8 L ddH_2_O to a 1.0 L flask; add a 2 in. stir bar; place on a stir plate (slight vortex without frothing, heat set to 100°C); add 4.86 g NaH_2_PO_4_·H_2_O (MW 137.99 g/mol); add 65.97 g Na_2_HPO_4_ (MW 141.96 g/mol); add 175.4 g NaCl (MW 58.44 g/mol); add 6.81 g Imidazole (MW 68.08 g/mol); adjust volume to ∼0.95 L with ddH_2_O; stir for 30 min; turn off stir plate heat (flask should not feel warm); check pH and add NaOH/HCl to adjust pH to 8.0; remove stir bar; adjust volume to 1.0 L with a 1.0 L graduated cylinder; pour solution back into 1.0 L flask with 2 in. stir bar on stir plate (slight vortex without frothing, no heat); check pH and adjust to 8.0 (if necessary); filter-sterilize (0.2 µm PES, 500 mL bottle-top filter) into a clean, dust-free, 1.0 L borosilicate bottle; and store at RT.

##### 1x NiNTA LYSIS Buffer (NiNTA-LB)

0.05 M Na phosphate, 0.3 M NaCl, 0.010 M Imidazole, pH 8.0*

1. ***Use <u>10x Stock Solution</u> to Assemble 1.0 L of 1x NiNTA-LB\**.** Assembly protocol (10x to 1x) was followed with the following modifications at the end of the protocol: adjust to pH 8.0* by drop-wise addition of 1.0 N HCl/NaOH; filter-sterilize (0.2 µm PES, 500 mL bottle-top filter) into a clean, dust-free, 1.0 L borosilicate bottle; store solution at RT. **\*Later work with a similar <u>10x Stock Solution</u> buffer (10x IMAC Buffer BASE) suggests that there would be a decrease in pH upon dilution from this <u>10x Stock Solution</u>, though we did not check the pH of 1x NiNTA-LB after dilution or adjust it after dilution when this stock was used for this paper’s experiments.**
2. ***Use <u>Reagents</u> to Assemble 1.0 L of 1x NiNTA-LB\**.** Add 0.8 L ddH_2_O to a 1.0 L flask; add a 2 in. stir bar; place on a stir plate (slight vortex without frothing, heat set to 100°C); add 6.90 g NaH_2_PO_4_·H_2_O (MW 137.99 g/mol); add 17.54 g NaCl (MW 58.44 g/mol); add 0.68 g Imidazole (MW 68.08 g/mol); adjust volume to ∼0.95 L with ddH_2_O; stir for 15 min; turn off stir plate heat (flask should not feel warm); check pH and add NaOH/HCl to adjust the starting pH from pH ∼6.4 to pH 8.0***\****; remove stir bar; adjust volume to 1.0 L with a 1.0 L graduated cylinder; cover the top of the cylinder with parafilm (≥3 layers); firmly press a palm onto the parafilm to seal the cylinder; invert the cylinder ten times create a homogeneous solution; remove the parafilm from the cylinder; filter-sterilize (0.2 µm PES, 500 mL bottle-top filter) into a clean, dust-free, 1.0 L borosilicate bottle; and store at RT. **\*Adjustment of pH normally required ∼1.5 mL 10 N NaOH, so the final solution contained ∼15.0 mN NaOH.**

##### 1x NiNTA WASH Buffer (NiNTA-WB)

0.05 M Na phosphate, 0.3 M NaCl, 0.020 M Imidazole, pH 8.0*

#### Use Reagents to Assemble 1.0 L of 1x NiNTA-WB*

Add 0.8 L ddH_2_O to a 1.0 L flask; add a 2 in. stir bar; place on a stir plate (slight vortex without frothing, heat set to 100°C); add 6.90 g NaH_2_PO_4_·H_2_O (MW 137.99 g/mol); add 17.54 g NaCl (MW 58.44 g/mol); add 1.36 g Imidazole (MW 68.08 g/mol); adjust volume to ∼0.95 L with ddH_2_O; stir for 15 min; turn off stir plate heat (flask should not feel warm); check pH and add NaOH/HCl to adjust the starting pH to pH 8.0***\****; remove stir bar; adjust volume to 1.0 L with a 1.0 L graduated cylinder; cover the top of the cylinder with parafilm (≥3 layers); firmly press a palm onto the parafilm to seal the cylinder; invert the cylinder ten times create a homogeneous solution; remove the parafilm from the cylinder; filter-sterilize (0.2 µm PES, 500 mL bottle-top filter) into a clean, dust-free, 1.0 L borosilicate bottle; and store at RT.

**\*Adjustment of pH normally required ∼7.25 mL 10 N NaOH, so the final solution contained ∼72.5 mN NaOH.**

##### 1x NiNTA ELUTION Buffer (NiNTA-EB)

0.05 M Na phosphate, 0.3 M NaCl, 0.250 M Imidazole, pH 8.0*\**

#### Use Reagents to Assemble 1.0 L of 1x NiNTA-EB

Add 0.8 L ddH_2_O to a 1.0 L flask; add a 2 in. stir bar; place on a stir plate (slight vortex without frothing, heat set to 100°C); add 6.90 g NaH_2_PO_4_·H_2_O (MW 137.99 g/mol); add 17.54 g NaCl (MW 58.44 g/mol); add 17.02 g Imidazole (MW 68.08 g/mol); adjust volume to ∼0.95 L with ddH_2_O; stir for 15 min; turn off stir plate heat (flask should not feel warm); check pH and add NaOH/HCl to adjust the starting pH to pH 8.0***\****; remove stir bar; adjust volume to 1.0 L with a 1.0 L graduated cylinder; cover the top of the cylinder with parafilm (≥3 layers); firmly press a palm onto the parafilm to seal the cylinder; invert the cylinder ten times create a homogeneous solution; remove the parafilm from the cylinder; filter-sterilize (0.2 µm PES, 500 mL bottle-top filter) into a clean, dust-free, 1.0 L borosilicate bottle; and store at RT.

**\*Adjustment of pH normally required ∼5.85 mL 10 N NaOH, so the final solution contained ∼58.5 mN NaOH.**

#### High Pressure Liquid Chromatography (HPLC) – IMAC Solutions

##### 10x IMAC Buffer BASE (IMAC-BB)

0.5 M Na phosphate, 3.0 M NaCl, pH 8.0*

#### Use Reagents to Assemble 2.0 L of 10x NiNTA-BB*

Add 1.2 L ddH_2_O to a 2.0 L flask; add a 2 in. stir bar; place on a stir plate (slight vortex without frothing, heat set to 100°C); add 8.46 g NaH_2_PO_4_ (MW 119.98 g/mol); add 249.17 g Na_2_HPO_4_·7H_2_O (MW 268.07 g/mol); add 350.64 g NaCl (MW 58.44 g/mol); adjust volume to ∼1.9 L with ddH_2_O; stir for 30 min; turn off stir plate heat (flask should not feel warm); check pH and add NaOH/HCl to adjust pH to 8.0*; remove stir bar; adjust volume to 2.0 L with a 2.0 L graduated cylinder; pour solution back into 2.0 L flask with 2 in. stir bar on stir plate (slight vortex without frothing, no heat), check pH and adjust to 8.0 (if necessary); filter-sterilize (0.2 µm PES, 500 mL bottle-top filter) into a clean, dust-free, 2.0 L borosilicate bottle; and store at RT.

**\*∼0.03 N NaOH added to adjust pH to 8.0**

##### 5x IMAC Imidazole Buffer (IMAC-IB)

5.0 M Imidazole, 0.05 M Na phosphate, 0.3 M NaCl, pH 8.0

#### Use Reagents to Assemble 1.0 L of 5x IMAC-IB

Add 0.3 L ddH_2_O to a 2.0 L borosilicate beaker; add a 2 in. stir bar; place on a stir plate (slight vortex without frothing, heat set to 100°C); add 0.1 L of 10x IMAC Buffer Base; slowly* add 340.4 g Imidazole (MW 68.08 g/mol, density 1.23 g/cm³); adjust volume to ∼0.95 L with ddH_2_O; stir for 30 min; turn off stir plate heat (flask should not feel warm); check pH and add NaOH/HCl to adjust pH to 8.0; remove stir bar; adjust volume to 1.0 L with a 1.0 L graduated cylinder; pour solution back into 1.0 L flask with 2 in. stir bar on stir plate (slight vortex without frothing, no heat); check pH and adjust to pH 8.0 by drop-wise addition of 1.0 N HCl/NaOH; filter-sterilize (0.2 µm PES, 500 mL bottle-top filter) into a new, dust-free, brown (low actinic**), 1.0 L borosilicate bottle; and store at 4°C.

**\*Add 10-20 g of imidazole, wait until it is mostly dissolved into solution, and then add additional imidazole.**

**\*\*If a low actinic bottle was not available, then the entire bottle was wrapped in aluminum foil.**

##### 1x HPLC IMAC Buffer A

0.05 M Na phosphate, 0.3 M NaCl, pH 8.0*

#### Use 10x Stock Solution (10x IMAC-BB) to Assemble 1.0 L of 1x HPLC IMAC Buffer A

Assembly protocol (10x to 1x) was followed with the following modifications at the end of the protocol: adjust to pH 8.0* by drop-wise addition of 1.0 N HCl/NaOH; filter-sterilize (0.2 µm PES, 500 mL bottle-top filter) into a clean, dust-free, 1.0 L borosilicate bottle; degas under house vacuum at RT for 30 minutes; store at 4°C.

**\*A decrease in pH was observed upon dilution from a 10x Solution.**

##### 1x HPLC IMAC Buffer B

0.05 M Na phosphate, 0.3 M NaCl, 1.0 M Imidazole, pH 8.0*

#### Use 10x and 5x Stock Solutions to Assemble 1.0 L of 1x HPLC IMAC Buffer B

Add 0.6 L ddH_2_O to a 1.0 L graduated cylinder; add 0.08 L of 10x IMAC-BB; add 0.2 L of 5x IMAC-IB, adjust volume to ∼0.95 L with ddH_2_O; adjust to pH 8.0* by drop-wise addition of 1.0 N HCl/NaOH; adjust volume to 1.0 L with ddH_2_O; cover the top of the cylinder with parafilm (≥3 layers); firmly press a palm onto the parafilm to seal the cylinder; invert the cylinder ten times create a homogeneous solution; remove the parafilm from the cylinder; filter-sterilize (0.2 µm PES, 500 mL bottle-top filter) into a new, dust-free, brown (low actinic**), 1.0 L borosilicate bottle; degas at RT for 30 minutes, and store at 4°C.

**\*A decrease in pH was observed upon dilution from a 10x Solution.**

**\*\*If a low actinic bottle was not available, then the entire bottle was wrapped in aluminum foil.**

### HPLC – Size-Exclusion Chromatography (SEC) Solutions

#### 20x HEPES Sizing Buffer (HSB)

0.4 M HEPES, 3.0 M NaCl, 0.2% Na Azide, 123 mM NaOH*, pH 7.4*

**\*Added to adjust pH to 7.4.**

#### Use Reagents to Assemble 2.0 L of 20x HSB

Add 1.2 L ddH_2_O to a 2.0 L flask; add a 2 in. stir bar; place on a stir plate (slight vortex without frothing, heat set to 50°C); add 9.80 g NaOH (39.997 g/mol); add 190.6 g HEPES (MW 238.30 g/mol); add 4.0 g Na azide (MW 65.01 g/mol); add 350.64 g NaCl (58.44 g/mol); adjust volume to ∼1.9 L with ddH_2_O; stir for 30 min; turn off stir plate heat (flask should not feel warm); check pH and add NaOH/HCl to adjust pH to 7.4; remove stir bar; adjust volume to 2.0 L with a 2.0 L graduated cylinder; pour solution back into 2.0 L flask with 2 in. stir bar on stir plate (slight vortex without frothing, no heat); check pH and adjust to 7.4 (if necessary); filter-sterilize (0.2 µm PES, 500 mL bottle-top filter) into a clean, dust-free, 2.0 L borosilicate bottle; and store at RT.

##### 1x HEPES Sizing Buffer (HSB)

0.02 M HEPES, 0.15 M NaCl, 0.01% Na Azide, pH 7.4*

#### Use 20x Stock Solution to Assemble 2.0 L of 1x HSB

Assembly protocol (20x to 1x, 2.0 L final volume) was followed with the following modifications: filter-sterilize (0.2 µm PES, 500 mL bottle-top filter) into a clean, dust-free, 2.0 L borosilicate bottle; degas under house vacuum at RT for 30 minutes; store at 4°C.

**\*The pH of this solution (pH 7.4) does not change upon dilution of the 20x stock to a 1x stock.**

### Gel Electrophoresis Solutions

#### Sodium Dodecyl-Sulfate (SDS) Polyacrylamide Gel Electrophoresis (PAGE) Solutions

##### 5x SDS PAGE Sample Buffer (SDS-SB)

*<u>5.26x SDS-SB(-)BME</u>:* 263.2 mM Tris (pH 7.0), 52.6% (m/v) glycerol, 10.5% (m/v) SDS, 0.053% (m/v) bromophenol blue (BPB)

<u>5x SDS-SB(+)BME:</u> 250 mM Tris (pH 7.0), 50% (m/v) glycerol, 10% SDS (m/v), 0.05% (m/v) BPB, 5% (v/v) beta-mercaptoethanol (BME)

#### Assembly of 45 mL of 5x SDS-SB

Mass 22.5 g glycerol into a 50 mL conical tube, add 22.5 mg BPB, add 4.5 g SDS, add 11.25 mL 1.0 M Tris (pH 7.0), add ddH_2_O to 42.75 mL, cap tube and secure the lid with parafilm (≥3 layers), vortex into solution (e.g., vortex for 3 min, invert five times, vortex for 3 min), pulse spin tube (Eppendorf 5920 R, rotor S-4x1000, 1000 xg, 1 min, RT), use a P1000 filter-tip to aliquot 475 µL into ninety microcentrifuge tubes, store these (-)BME tubes at - 20°C, use a P200 filter-tip to add 25 µL of 100% BME to aliquot(s) to be used immediately as 5x SDS-SB(+)BME, vortex BME into solution, pulse spin (miniature microcentrifuge, 3 sec pulse, RT), and store at RT for ≤30 days to avoid losing reductive capacity due to BME evaporation.

10x SDS PAGE Running Buffer (SDS-RB a.k.a., Tris/Glycine/SDS Buffer)

250 mM Tris, 1.92 M Glycine, 1.0% (m/v) SDS, pH 8.3*

***\*∼0.084 N HCl added to adjust pH to 8.3*.**

#### Assembly of 2.0 L of 10x SDS-RB

Add 1.2 L ddH_2_O to a 2.0 L flask; add a 2 in. stir bar; place on a stir plate (slight vortex without frothing, plate heat set to 70°C); add 288.27 g glycine (75.07 g/mol); add 60.55 g Tris-base (121.14 g/mol); add 20.0 g SDS; add 16.8 mL 10 N HCl; adjust volume to ∼1.9 L with ddH_2_O; stir until all reagents are in solution; turn off stir plate heat (flask should not feel warm); check pH and add HCl/NaOH to adjust pH to 8.3; remove stir bar; adjust volume to 2.0 L with a 2.0 L graduated cylinder; pour solution back into 2.0 L flask with 2 in. stir bar on stir plate (slight vortex without frothing, no heat); check pH and adjust to 8.3 (if necessary); filter-sterilize (0.2 µm PES, 500 mL bottle-top filter) into a clean, dust-free, 2.0 L borosilicate bottle; and store at RT.

### NATIVE PAGE Solutions

#### 2x NATIVE PAGE Sample Buffer (NATIVE-SB)

62.5 mM Tris, 40%(m/v) glycerol, 0.01%(m/v) BPB, pH 6.8

#### Assembly of 50 mL of 2x NATIVE-SB

Mass 20.0 g glycerol into a 50 mL conical tube, add 3.125 mL 1.0 M Tris (pH 7.0), add ddH_2_O to 45.0 mL, cap tube and secure the lid with parafilm (≥3 layers), vortex into solution, pulse spin tube (Eppendorf 5920 R, rotor S-4x1000, 1000 xg, 1 min, RT), check pH and add HCl/NaOH to adjust pH to 6.8, add 5.0 mg BPB, cap tube and secure the lid with parafilm (≥3 layers), vortex into solution, pulse spin tube (Eppendorf 5920 R, rotor S-4x1000, 1000 xg, 1 min, RT), adjust volume to 50 mL with ddH_2_O, cap tube and secure the lid with parafilm (≥3 layers), vortex into solution, pulse spin tube (Eppendorf 5920 R, rotor S-4x1000, 1000 xg, 1 min, RT), filter-sterilize (0.2 µm PES syringe filter, ≥25 mm diameter) into microcentrifuge tubes as 1.0 mL aliquots, and store at -20°C.

### 4x NATIVE PAGE Sample Buffer (NATIVE-SB)

125 mM Tris, 80%(m/v) glycerol, 0.02%(m/v) BPB, 0.01% azide*, pH 6.8

#### Assembly of 50 mL of 4x NATIVE-SB

Mass 40.0 g glycerol into a 50 mL conical tube, add 6.25 mL 1.0 M Tris (pH 7.0), add ddH_2_O to 45 mL, cap tube and secure the lid with parafilm (≥3 layers), vortex into solution, pulse spin tube (Eppendorf 5920 R, rotor S-4x1000, 1000 xg, 1 min, RT), check pH and add HCl/NaOH to adjust pH to 6.8, add 10.0 mg BPB, add 50 µL of 10% (w/v) sodium azide (1000x stock), cap tube and secure the lid with parafilm (≥3 layers), vortex into solution, pulse spin tube (1000 xg, 1 min, RT), adjust volume to 50 mL with ddH_2_O, cap tube and secure the lid with parafilm (≥3 layers), vortex into solution, pulse spin tube (Eppendorf 5920 R, rotor S-4x1000, 1000 xg, 1 min, RT), filter-sterilize (0.2 µm PES syringe filter, ≥25 mm diameter) into microcentrifuge tubes as 1.0 mL aliquots, and store at 4°C.

***\*Though references for 1x NATIVE-SB do not mention the inclusion of azide to prevent microbial growth, 0.01% (m/v) sodium azide was added to this stock so that it could be stored at 4°C*.**

### 10x NATIVE PAGE Running Buffer (NATIVE-RB a.k.a., Tris/Glycine Buffer)

250 mM Tris, 1.92 M Glycine, pH 8.3*

*∼0.06 N HCl added to adjust pH to 8.3.

#### Assembly of 1.0 L of 10x NATIVE-RB

Add 0.5 L ddH_2_O to a 2.0 L flask; add a 2 in. stir bar; place on a stir plate; set stir rate to a slight vortex (no frothing) and temperature to 70°C; add 144.13 g glycine (75.07 g/mol); add 30.28 g Tris-base (121.14 g/mol); add 6.0 mL 10 N HCl; adjust volume to ∼0.9 L with ddH_2_O; stir until all reagents are in solution; turn off stir plate heat (flask should not feel warm); check pH and add HCl/NaOH to adjust pH to 8.3; remove stir bar; adjust volume to 1.0 L with a 1.0 L graduated cylinder; pour solution back into 2.0 L flask with 2 in. stir bar on the stir plate; set stir rate to a slight vortex (no frothing) without heat; check pH and adjust to 8.3 (if necessary); and filter-sterilize (0.2 µm PES, 500 mL bottle-top filter) into a clean, dust-free, 1.0 L borosilicate bottle; and store at RT.

## Gel Stain Solutions

### o-Dianisidine Stain (ODS = Heme Stain) Solutions

#### 0.5 M Citrate (pH 4.4)

##### Assembly of 50 mL of 0.5 M Citrate (pH 4.4)

Mass 7.35 g sodium citrate (MW 294.10 g/mol) into a 50 mL conical tube, add ddH_2_O to reach 40 mL volume, vortex into solution, pulse-spin (Eppendorf 5430 R, rotor F35-6-30, 1000 xg, 1 min, RT), add 6.0 N HCl drop-wise (∼180 drops from a 2 mL borosilicate Pasteur pipet, ∼9 ml) to reach pH 4.4 (e.g., add small HCl volume, vortex, pulse-spin, assess pH, repeat process until pH 4.4), adjust volume to 50 ml with ddH_2_O, vortex 10 sec, pulse-spin, store at room temperature. Does not need to be filter-sterilized.

#### 10x *o*-Dianisidine Dihydrochloride (60 mM)

##### Assembly of 10 mL of 10x o-Dianisidine Dihydrochloride (60 mM)

Mass 0.19 g o-dianisidine-2HCl (MW 317.21 g/mol, stored at 4°C) to a 15 mL conical tube, add ddH_2_O to 10 ml, vortex into solution, pulse-spin (Eppendorf 5430 R, rotor F35-6-30, 1000 xg, 1 min, RT), store at 4°C covered in foil. Does not need to be filter-sterilized. Stable for up to 6 months (data not shown).

### Protein Binding Assay Solutions

<u>NP buffer (NPB)</u> – Heme-agarose (HA) Pulldown Assay Solution 10 mM Na phosphate, 0.5 M NaCl, pH 7.5

#### Assembly of 100 mL of NPB

Add 50.0 mL ddH_2_O to a 100.0 mL borosilicate beaker; add a 1 in. stir bar; place on a stir plate (slight vortex without frothing, RT); add 188 µL 1.0 M sodium phosphate monobasic; add 812 µL 1.0 M sodium phosphate dibasic; add 10.0 mL 5.0 M NaCl; adjust to 95.0 mL with ddH_2_O; adjust pH from 7.3 to 7.5 with 1.0 N NaOH; remove stir bar; adjust volume to 100 mL; filter-sterilize using a 0.2 µm PES syringe filter (≥25 mm diameter); and store at RT.

## Porphyrin Stock Solutions

### 0.226 M Hematin in 1.4 N NaOH*

#### Assembly of ∼1.018 mL of 0.226 M Hematin in 1.4 N NaOH

Mass 0.15 g hemin chloride (651.94 g/mol) into a 50 mL conical tube, add 0.418 mL 1.4 N NaOH, vortex the tube at maximum setting for 1 min, pulse spin (Eppendorf 5920 R, rotor S-4x1000, 1000 xg, 1 min, RT), add 0.2 mL 1.4 N NaOH, vortex the tube at maximum setting for 1 min, pulse spin (Eppendorf 5920 R, rotor S-4x1000, 1000 xg, 1 min, RT), inspect tube for solubilization of hematin (e.g., particulates still visible in solution), add 0.2 mL 1.4 N NaOH, vortex the tube at maximum setting for 1 min, pulse spin (Eppendorf 5920 R, rotor S-4x1000, 1000 xg, 1 min, RT), inspect tube for solubilization of hemin (e.g., solution filled with bubbles), add 0.2 mL 1.4 N NaOH, vortex the tube at maximum setting for 1 min, pulse spin (Eppendorf 5920 R, rotor S-4x1000, 1000 xg, 1 min, RT), inspect tube for solubilization of hemin (e.g., solution filled with bubbles), first hard spin of hematin solution (Eppendorf 5920 R, rotor S-4x1000, 3428 xg, 10 min, RT), inspect tube for solubilization of hemin (e.g., some bubbles popped), transfer supernatant to two new 2.0 mL microcentrifuge tubes, second hard spin of hematin solution (Eppendorf 5425R, rotor FA-24x2, 21300 xg, 10 min, RT), inspect tube for solubilization of hemin (e.g., most bubbles gone and no sign of pellet/precipitate), transfer supernatant to a syringe, sterile filter (0.22 µm PES, ≥25 mm diameter) the solution (filtered easily), aliquot solution into amber (light protection) 1.5 mL microcentrifuge tubes, and store tubes at -80°C.

**\*Approximate concentrations. Based on hemin chloride density (7.87 g/cm^3^ at 25°C), the volume added from hemin chloride was likely ≤0.019 mL, which would make the final volume ≤1.037 mL. As such, the final concentrations may have been decreased to: [hematin]^final^ = 0.222 M, [NaOH]^final^ = 1.37 M.**

#### Assessment of effect of 0.226 M Hematin in 1.4 N NaOH stock on HSB samples

Binding reactions were performed in HSB, and so the effect of the addition of this hematin stock on an HSB sample was assessed. To assess the effect of the 0.226 M Hematin in 1.4 N NaOH on HSB samples, we used pH strips to assess the pH of dilutions of this stock in HSB +/- Tris (pH 7.0) at a 1:1 molar ratio to the NaOH concentration. The 0.226 M hematin in 1.4 N NaOH stock was diluted 25-fold into HSB to create **<u>stock A</u>**: 1.0 mL stock of 9.0 mM hematin, 56.0 mN NaOH, 0.96x HSB. **<u>Stock A</u>** was then split into two 1.5 mL microcentrifuge tubes. A 1.0 M Tris (pH 7.0) stock was then diluted in HSB (e.g., 55.6 µL 1.0 M Tris (pH 7.0), 944.4 µL HSB) to create **<u>stock</u> <u>B:</u>**1.0 mL of 55.6 mM Tris (pH 7.0), 0.94x HSB. Then, 500 µL of **<u>stock B</u>**was added to one of the **<u>stock A</u>** tubes to create a 1:1 (v/v) mixed **<u>stock AB:</u>** 4.5 mM hematin, 28.0 mN NaOH, 27.8 mM Tris (pH 7.0), 0.95x HSB. The remaining **<u>stock A</u>** tube and the **<u>stock AB</u>** were then used to create a 10-fold dilution series in HSB (e.g., 500 µL of 1/10, 1/100, and 1/1000 dilutions). Finally, pH strips were used to evaluate these eight samples (see **SI Table 10**). Hematin alone samples from 9.04-904 µM could be assessed and were neutral pH. The addition of Tris had no discernable effect on the final pH of those samples that could be assessed using pH strips.

#### Assembly of Porphyrin in DMSO Stocks

##### Initial 4.5 mM Hemin in DMSO Stock

Mass 24.6 mg hemin chloride (651.9 g/mol) into a 15 mL conical tube, add 8.4 mL DMSO, vortex into solution, inspect tube for solubilization (no particulates visible), pellet insoluble material (Eppendorf 5430 R, rotor F35-6-30, 6110 xg, 5 min, 4°C), inspect tube for precipitate (minor precipitate visible at bottom of tube), wrap in aluminum foil to protect from light degradation, store at 4°C (solidifies DMSO), thaw DMSO stock at RT immediately prior to use, and pulse-spin (Eppendorf 5430 R, rotor F35-6-30, 1000 xg, 1 min, RT).

##### Subsequent 10.0 mM Hemin in DMSO Stock*

Mass 40.00 mg hemin chloride (Sigma Cat No. D45124, MW 651.94 g/mol) into a 15 mL conical tube, add 6.136 mL DMSO, vortex for 1 minute, pulse-spin (Eppendorf 5810 R, rotor FA-45-6-30,1000 xg, 1 min, RT), transfer hemin solution to ten new 2.0 mL microcentrifuge tubes, hard spin hemin solution (Eppendorf 5430 R, rotor FA-45-30-11, 20817 × g, 10 min, RT), inspect tube for solubilization of hemin**, syringe filter (0.2 µm PTFE, 25 mm diameter) the solution into a new 15 mL conical tube, aliquot solution (200 µL/tube) into 1.5 mL tubes, store at -80°C, thaw DMSO stock at RT immediately prior to use, and pulse-spin (Eppendorf 5430 R, rotor F35-6-30, 1000 xg, 1 min, RT).

**\*Hemin DMSO solubility of up to 20 mg/mL (30.7 mM) has previously been reported by commercial sources of hemin (e.g.,** https://www.selleckchem.com/products/hemin.html).

**\*\*No obvious pellet was observed at this step.**

##### Optimized Assembly of 5.0 mM Hemin in DMSO Stock

Mass 32.60 mg hemin chloride (651.94 g/mol) into a 15 mL conical tube; add 10.0 mL DMSO; vortex the tube at maximum setting for 1 min; pulse spin (Eppendorf 5920 R, rotor S-4x1000, 1000 xg, 1 min, RT); inspect tube for solubilization of hemin; if particulates are still visible in solution, repeat vortex-pulse-inspect until all hemin is solubilized; first hard spin of hemin solution (Eppendorf 5920 R, rotor S-4x1000, 3428 xg, 10 min, RT); inspect tube presence of any precipitate; transfer supernatant to ten new 2.0 mL microcentrifuge tubes; second hard spin of hemin solution (Eppendorf 5425R, rotor FA-24x2, 21300 xg, 10 min, RT); inspect tubes for presence of any precipitate; transfer supernatant to a 10 mL syringe; syringe filter (0.2 µm PTFE, 25 mm diameter) the solution into a new 15 mL conical tube; aliquot solution into amber (light protection), 1.5 mL microcentrifuge tubes; store tubes at - 80°C; thaw DMSO stock immediately prior to use; and pulse-spin (Eppendorf 5425R, rotor FA-24x2, 1000 xg, 1 min, RT).

Protoporphyrin IX (PPIX) and Coproporphyrin III (CPIII) in DMSO Stocks

Followed the same general protocol as outlined for “Initial Hemin in DMSO Stock” except that DMSO volume and porphyrin mass were modified to prepare 5.0 mM stocks. Molecular weight (MW) of metal-free porphyrins: PPIX 562.66 g/mol, CPIII dihydrochloride 727.63 g/mol.

## COMPUTATIONAL ANALYSIS

### RoxP Ortholog Selection and Sequence Analysis

RoxP orthologs were identified using a BlastP [2] search. The search query was chosen to be the post-signal peptide cleavage sequence of the reference RoxP ortholog (Sequence ID: WP_002515361.1) found in strain KPA171202, which is referred to as RoxP_1 in this manuscript. In 2018, we used SignalP - 4.1 [3] to identify the likely RoxP_1 signal sequence (residues 1-22, MFV..PGA). Of note, a later version of this program (SignalP – 6.0) that our group used in 2023 predicted a slightly different RoxP_1 signal sequence (residues 1-18, MFV..ALG), which suggests there may be four additional N-terminal residues (IPGA) after signal peptide cleavage. Then, the SignalP - 4.1 predicted RoxP_1 signal sequence (MFV..PGA) was removed from the nascent RoxP_1 amino acid sequence (MFV..ALN) to create a “post-signal peptide cleavage sequence search query,” and this search query (ATP…ALN) was used in a 2019 BlastP [2] search of the “Non-redundant protein sequences (nr)” database.

Ten orthologous sequences (RoxP_1-10) were initially identified via this BlastP search, and a repeat of this search in 2022 identified 11 additional orthologs for 21 total RoxP orthologs (**SI File 1**; **SI Table 1**,**3**). **SI File 1** is a fasta file listing these sequences with the ortholog name, RefSeq accession number, and expression construct in this paper (if available) listed in the header (e.g., RoxP_1, WP_002515361.1, RoxP_pET22b). MUSCLE [4] and Ugene [5] were used to construct a multiple sequence alignment (MSA) of these 21 RoxP ortholog sequences where the residue background was colored based on its percentage identity to RoxP_1 (**SI Fig 1**). Each RoxP ortholog’s identity to RoxP_1 is listed to two decimal places in **SI Table 3**, and it is listed after rounding to the nearest whole integer in all other parts of this manuscript. Solvent accessible surface area (SASA) values for each residues of RoxP solution structure (PDB: 7BCJ) was then calculated using PolyView [6] and confirmed using GetArea [7]. GetArea SASA is indicated below each residue in **SI Fig 1**.

The RefSeq accession number of each ortholog (**SI File 1**) was then used to fetch its identical protein group (IPG) identifier (https://www.ncbi.nlm.nih.gov/ipg/). A table of selected protein sequence accession numbers and their assigned names for our experiments are given in **SI Table 1**. All available protein sequences associated with IPG identifiers were selected for further analysis, unique entries were identified (e.g., duplicate entries were consolidated into a single entry), and a RoxP sequence space of 946 available RoxP sequences was defined. From this list of RoxP entries, all *Cutibacterium* species (spp.) sequences were identified by selecting entries that had “Cutibacterium” as part of the organism designation (total 905). *Cutibacterium acnes* sequences were selected from the *Cutibacterium* spp. group by selecting for entries that had “*Cutibacterium acnes”* as the organism designation (total 862). *Cutibacterium acnes* subspecies (subsp.) *acnes*, *defendens*, and *elongatum* sequences were selected from the C*utibacterium acnes* group wherever subspecies information was available in the organism designation. No RoxP sequence was identified for *C. acnes* subps. *elongatum*. For RoxP_8, IPG listed two *C. namnetense* strains (OGSA_21, NTS 31307302) and one unnamed strain deposited as “*C. acnes,*” but the latter strain’s submitted GenBank assembly (GCA_005768695.1) failed it’s taxonomy check. This strain’s best match type-strain was *C. namnetense* (GCA_001642025.1, average nucleotide identity [ANI] 99.74%, assembly coverage 79.33%, type assembly coverage 90.96%). Comparisons of RoxP ortholog carriage on a species level was made using both (**Fig 1b**) protein accession numbers (e.g., RefSeq, GenBank) listed by IPG and (**Fig 1c**) IPG entries alone, and the latter strategy identified the true RoxP ortholog spectrum carried by *C. acnes* subsp. *acnes* and *defendens*.

#### Computational Analysis of RoxP Orthologs

Signal peptide cleavage sites were initially predicted using SignalP - 4.1 [3] and later re-evaluated using an updated version of this program, SignalP – 6.0 [8]. All signal peptide cleavage sites described are from SignalP - 4.1 unless otherwise noted. Each protein’s amino acid sequences was used in ExPASy ProtParam [9] to calculate the protein’s MW and theoretical isoelectric point (pI). Unless otherwise noted, pI values in this study were calculated from a protein’s sequence using ProtParam. Solvent accessible surface area (SASA) of each amino acid side chain in the apo_RoxP_1 NMR structure (PDB 7bcj) [10] was initially assessed using PolyView [6] and then confirmed using GetArea [7]. SASA analysis and RoxP ortholog residue variation is summarized in **SI File 2**, which includes a definitions of conservative/non-conservative amino acid substitutions. RoxP conservation was further assessed using ConSurf [11]. RoxP conservation analysis was used to generate images in PyMOL [12] showing RoxP ortholog variation colored by SASA as defined by GetArea (**Fig 2**). Electrostatic surfaces were generated in PyMOL using the APBS plugin [13].

#### Generation of Three-dimensional Models of RoxP Orthologs

Phyre2 one-to-one threading [12] using the nuclear magnetic resonance (NMR) model (state 1) of apo_RoxP_1 (PDB 7bcj) [10] was used to generate three-dimensional models of RoxP orthologs. RoxP ortholog protein sequences used to generate Phyre2 models were based on recombinant RoxP constructs (**SI File 3**) after removal of the PelB signal peptide and N-terminal 8His-TEV tag (e.g., RoxP_5 GAT…AIN). Phyre2 did not include the TEV site glycine or residue 1 of the four orthologs. Using the RoxP_1 NMR model (state 1) residue 1 (A1), an alanine was added in PyMOL to RoxP_5, 6, 8, and 11 at position 1, though the latter two orthologs have P and V at that position, respectively. Modeling an alanine at that position was performed because (1) all models had an extremely small Root Mean Square Deviation (RMSD) after performing cealign [14] in PyMOL against RoxP_1 (0.000001 Å over 136 residues), (2) individual *in silico* addition of position 1 to each ortholog followed by localized molecular dynamic simulation could not be scripted in PyMOL, (3) this residue’s molecular surface is distant from all surfaces being interpreted in this paper, and (4) minimal change in the molecular surface would be expected at this position for A/V/P due to its presence as random coil in PDB 7bcj with no interaction with nearby secondary/tertiary structural elements. Phyre2 also flipped the W66 side chain dihedral angle χ1, which results in further exposure of this residue’s side chain and the appearance of a new cavity on the left side of the four ortholog models (**Fig 5b**-**e**,**g**-**j**; **SI Fig 7b**-**c**,**e-f).** It is not yet clear if the NMR model (7bcj) or the Phyre2 models in this work have correctly modeled the W66 dihedral angle χ1.

### Design of Recombinant Expression Constructs for RoxP Orthologs

Ten RoxP orthologs (RoxP_1-10) were initially chosen for recombinant expression based on the initial description of RoxP_1 in the literature [15] and a 2019 BlastP query of that ortholog’s post-signal peptide cleavage sequence (ATP..ALN) against the non-redundant protein sequences (nr) database. In 2022, that BlastP search was run again, which identified eleven additional RoxP orthologs. To expand the sampled RoxP amino acid sequence space, RoxP_11 (71.22% identity to RoxP_1, identified in *Sus domesticus* gastrointestinal tract [16]) was added to the RoxP orthologs chosen for recombinant expression.

For each selected RoxP ortholog, the RoxP coding region was identified (e.g., RoxP_1 from KPA171202; NCBI Reference Sequence: WP_002515361.1; 161 amino acids, MFV..ALN), and SignalP - 4.1 [3] was used to identify the ortholog’s endogenous signal peptide sequence. The post-signal peptide cleavage sequence of the ortholog (e.g., RoxP_1 ATP..ALN) was then codon-optimized for expression in *Escherichia coli* using commercially-available website software (e.g., IDT, GENEWIZ, GenScript). Finally, the nucleotide sequence of a new N-terminal leader peptide sequence ^a^ was added to the codon-optimized RoxP nucleotide sequence.

This leader sequence included an *E. coli* periplasmic secretion signal (PelB: MKY-AMA), 8His-tag (MDH_8_), flexible linker (GGS), and TEV protease site (ENLYFQG). The nucleotide sequence for this amino acid leader sequence utilized an established PelB nucleotide sequence from pET22b (PelB: MKY-PAM, ATG..TGG) fused to an *E. coli* codon-optimized nucleotide sequence (CCA..GGT) that included two nucleotides (CC) to code for the last alanine of the PelB leader and the remaining N-terminal tag sequence (MDH..FQG). The PelB leader peptide was included to direct RoxP to the periplasm for proper disulfide bond formation. The same nucleotide sequence for this amino acid leader sequence was used for all RoxP constructs. See **SI File 3** for the protein sequence of all recombinant RoxP constructs.

Recombinant RoxP expression nucleotide sequences were submitted to GenScript, where they were synthesized, cloned as an NcoI-XhoI fragment into the pET22b expression vector, and then verified by DNA sequencing. GenScript then sent purified plasmid DNA and all data related to synthesis/cloning to the McCoy Lab. All RoxP ortholog expression constructs were then transformed into the *E. coli* protein expression strain BL21(DE3)RIL and stored as glycerol stocks. **^a^ N-terminal leader peptide protein sequence: MKYLLPTAAAGLLLLAAQPAMAMDHHHHHHHHGGSENLYFQG**

## RECOMBINANT PROTEIN PRODUCTION

### Large-scale Expression and Purification of Recombinant RoxP Orthologs

Large-scale preparation of all recombinant RoxP proteins involved autoinduction, lysate preparation, IMAC purification, and tag removal. The protocol for producing recombinant RoxP_1-tagless is described below. Unless stated otherwise, the following statements apply to all experiments described in this section.

1. All solutions in this section are 1x unless noted otherwise.
2. All solutions were pre-chilled to 4°C prior to use and kept at 4°C during the experiment.
3. All sample/solution assembly steps were followed by mixing and a pulse spin step.

a. Microcentrifuge tubes: flick-to-mix five times, then 3 sec in a miniature microcentrifuge.
b. 15/50 mL conical tubes: invert five times, then centrifuge for 10 sec at 1000 xg.
4. A mix-pulse step was performed after the addition of each reagent to a sample/solution.
5. Whenever possible, all samples containing protein (e.g., cell pellet, lysate, chromatography washes/elutions, filtrates, concentrates, reactions) were kept at 4°C.
6. All centrifugation steps performed at 4°C were preceded by cooling the centrifuge and rotor to 4°C (e.g., Eppendorf ***fast temp*** function set to 4°C).
7. Filtration of protein-containing samples used hydrophilic, low protein binding filters (PES; PVDF: polyvinylidene difluoride; CA: cellulose acetate) with diameters matched to sample volumes: 4 mm, ≤1 mL; 13 mm, 1-10 mL; 25/33 mm, 10-60 mL; >60 mL, bottle top filter.
8. Filtration of protein-containing samples used specific pore sizes depending on the sample: <u>Non-HPLC Lysate Work</u>

a. Lysate from ≤500 mL culture was filtered using a ≤0.45 µm filter.
b. Lysate from >500 mL culture was filtered using a 5.0 µm filter in series with a downstream ≤0.45 µm filter. <u>HPLC Lysate Work</u>
c. Lysate from ≤500 mL culture was filtered using a ≤0.22 µm filter.
d. Lysate from >500 mL culture was filtered using a 5.0 µm filter in series with a downstream ≤0.22 µm filter. All Other Protein Samples* **\*Examples include a sample for a concentrator, a non-lysate sample for an HPLC, and the final filtration step for a protein sample prior to its use in an experiment.**
e. ≤0.22 µm filter
9. New, prechilled (4°C), plastic (polypropylene/polystyrene) bottles/tubes/concentrators were used to collect all protein-containing filtrates.
10. Ni-NTA settle steps were repeated until the solution above the column bed was clear.
11. All lysate/buffer additions to a gravity flow column were performed gently (e.g., slowly pour or pipet down the side of the funnel/column) to minimize disturbing the column bed.
12. All centrifugation ***hard-spin*** steps to pellet out protein precipitates/aggregates were followed by supernatant (soluble protein) transfer to a new, pre-chilled (4°C), 1.5 mL Low Protein Binding Microcentrifuge (LPBM) tube.

#### Autoinduction

**<u>Day 1</u>** – Streak RoxP_pET22b:BL21(DE3)RIL glycerol stock on an LB plate supplemented with 100 µg/mL Carbenicillin (Carb) and 25 µg/mL Chloramphenicol (Chlor) ^a^. **<u>Day</u> <u>2</u>** – ***(1)*** Pick a single colony along the streak to inoculate a 5.0 mL LB liquid culture supplemented with 100 µg/mL Carb and 25 µg/mL Chlor ^a^, ***(2)*** grow the LB-Carb+Chlor ^a^ culture overnight (≥16 hr) at 37°C (shaking, 200-220 rpm) in a 14 mL round bottom Falcon tube. **<u>Day 3</u>** – ***(1)*** Add 0.5 mL of the overnight culture to a 1.0 L flask containing 0.5 L of ZYM-5052 supplemented with 2 mM MgSO_4_, 0.5x Trace Metals, 0.1 mM FeCl_3_, 100 µg/mL Carb, and 25 µg/mL Chlor ^a^ [1]; ***(2)*** grow the 0.5 L ZYM-5052-Carb+Chlor culture at 37°C (shaking, 200-220 rpm) to optical density (OD) at 600 nm (OD_600_) ≥1.0 ^b^ but ≤3.0 ^c^; ***(3)*** drop the growth temperature to 18°C; and ***(4)*** grow for 5 days at 18°C to reach an OD_600_ maximum of 11-12. **<u>Day 8</u>** – Place 1.0 L flask in an ice bath for ≥20 min to chill culture to 4°C.

**^a^ Chlor can be omitted, as RoxP expression was not found to require extra copies of *argU/ileY/leuW* tRNA genes. Those genes are encoded on a pACYC-based plasmid (3.5 kB, ColE1-compatible, Chlor resistance [ChlorR]) included in BL21-CodonPlus(DE3)-RIL cells to help support expression of heterologous proteins in *E. coli*.**

**^b^ Typically takes ∼7 hr to reach OD^600^ ≥1.0.**

**^c^ Chosen to avoid autoinducing at 37°C, which causes RoxP_1 to be insoluble (*data not shown*). In general, autoinduction occurs in ZYM-5052 medium at OD^600^ >3.0.**

#### IMAC Protocols*

**\*Initial work used (1) gravity-flow IMAC, which was later adapted to an (2) HPLC IMAC protocol.**

## 1. Gravity-Flow IMAC Protocol

### Used to prepare apo-RoxP_1/5/6/8-tagless

#### Lysate Preparation

**<u>Day 8</u>** – ***(1)*** After chilling culture to 4°C, spin down *E. coli* cells in 500 mL polypropylene centrifuge bottle (Cat # 361691) using a Beckman Coulter Avanti J-E centrifuge (rotor JLA-10.500, 10000 xg, 10 min, 4°C); ***(2)*** mass the cell pellet (typically ∼18 g per 500 mL autoinduction culture) and then store it in its centrifuge bottle on ice; ***(3)*** pre-chill 45 mL of NiNTA-LB in a 50 mL conical (Corning# 352070) to 4°C; ***(4)*** prepare Complete NiNTA-LB by adding each of the following reagents to the pre-chilled NiNTA-LB: 225 µL of 200 mM PMSF (in 100% EtOH), 112.5 µL of 400x Divalent Cation Stock (DCS: 400 mM MgCl_2_, 400 mM CaCl_2_), 45 mg of lysozyme, and 0.45 mg of DNase; ***(5)*** add 10-15 mL Complete NiNTA-LB per g cell pellet ^a^ to the centrifuge bottle; ***(6)*** add a 2-in stir bar to the centrifuge bottle, cap the bottle, and shake bottle to dislodge the cell pellet from the centrifuge bottle side wall; ***(7)*** using a pre-chilled stir plate in a cold room (4°C), stir (without frothing) the cell pellet for ≥30 min to resuspend the cells in Complete NiNTA-LB; ***(8)*** remove the stir bar and pack the centrifuge bottle in ice; ***(9)*** sonicate cell suspension on ice using a Fisherbrand™ Model 505 Sonic Dismembrator (0.5 in. probe; 75% amplitude; 3 sec ON, 30 sec OFF, 3 min total sonication time, 33 min total time); ***(10)*** place 2-in stir bar back in the centrifuge bottle and stir on a pre-chilled stir plate in a cold room (4°C) for 15 min; ***(11)*** repeat steps (8-10) twice for three total rounds of sonication (9 min total sonication time) and stirring (45 min total post-sonication stirring time) ^b^; ***(12)*** remove the stir bar, transfer the lysate to a 500 mL polycarbonate centrifuge bottle (Cat # 361690), and spin out insoluble material using a Beckman Coulter Avanti J-E centrifuge (rotor JLA-10.500, 18600 xg, 60 min, 4°C); and ***(13)*** filter lysate.

**^a^ A lower ratio of 1:1 buffer:cells (mL:g) may be used, but this more viscous lysate may require extended centrifugation and/or additional filtration in later steps to remove insoluble/aggregated material.**

**^b^ The sample should change color from light brown to muddy, red-brown by the end of this step, which reflects release of intracellular materials.**

#### Chromatography

**<u>Day 8</u>** – ***(1)*** Assemble the gravity flow column hardware (Econo-Column Funnel → Econo-Column [2.5 × 10 cm] → two-way stopcock) and place it in a ring stand in a cold room ^a^; ***(2)*** rinse assembled column hardware extensively with diH_2_O; ***(3)*** add 10.0 mL of 50% Ni-NTA agarose slurry (QIAGEN #30210 or GOLDBIO #H-350-25) to the column, allow Ni-NTA to settle for ≥20 min, open stopcock until 30% EtOH storage solution meniscus is 1-2 mm above the Ni-NTA; ***(4)*** add 10 column volumes (CV) of NiNTA-LB (50 mL), allow Ni-NTA to settle for ≥10 min, and open stopcock until meniscus is 1-2 mm above the Ni-NTA; ***(5)*** apply filtered lysate to column, allow Ni-NTA to settle for ≥10 min, and open stopcock until meniscus is 1-2 mm above the Ni-NTA; ***(6)*** repeat step (5) ≥2 times to maximize RoxP yield ^b^; ***(7)*** add 5 CV of NiNTA-LB (25 mL), allow Ni-NTA to settle for ≥10 min, and open stopcock until meniscus is 1-2 mm above the Ni-NTA; ***(8)*** repeat step (7); ***(9)*** add 5 CV of NiNTA-WB (25 mL), allow Ni-NTA to settle for ≥10 min, and open stopcock until meniscus is 1-2 mm above the Ni-NTA; ***(10)*** repeat (9); ***(11)*** after performing four 25 mL washes, elute RoxP by adding 5 CV of NiNTA-EB (25 mL), allowing Ni-NTA to settle for ≥10 min, opening the stopcock, and collecting five 5 mL (1 CV) fractions until meniscus is 1-2 mm above the Ni-NTA; ***(12)*** evaluate SDS PAGE samples from all expression/purification steps, and ***(13)*** store protein samples at 4°C. Average gravity column flow rate for all steps was 5.8 mL/min.

**^a^ Maximum RoxP recovery requires that this purification be performed in a cold room, but it can be performed at room temperature with minimal yield loss for RoxP_1 if all buffers/samples are kept at 4°C.**

**^b^ Saturation of 1 mL of Ni-NTA agarose with RoxP_1 lysate will yield 22-35 mg of protein.**

#### Buffer-Exchange (BE)

**<u>Day 9</u>** – ***(1)*** Filter Ni-NTA elution into a centrifugal concentrator (Vivaspin 20 mL MWCO 10 kDa ^a^); ***(2)*** centrifuge (Eppendorf 5430 R, rotor F-35-6-30, 7197 xg, 4°C) to concentrate sample to ≤1 mL but >0.5 mL; ***(3)*** add HSB to concentrator to 14 mL mark for a >10-fold dilution into HSB; ***(4)*** repeat steps (2-3) ***<u>two times</u>*** for >2744-fold total dilution (i.e., BE) of the original buffer (NiNTA-EB) to reduce imidazole to ≤0.1 mM; ***(5)*** repeat step (2); ***(6)*** hard-spin sample (Eppendorf 5430 R, rotor FA-45-30-11, 20817 xg, 10 min, 4°C); ***(7)*** measure sample A_280_ (Tecan Spark, NanoQuant Plate with optical path length of 0.05 cm) to calculate the 8His-TEV-RoxP (MDH..ALN) concentration (ɛ280 14565 M^-1^ cm^-1^, MW 17228.04 g/mol) and yield ^b^; and ***(8)*** then store RoxP protein sample at 4°C.

**^a^ RoxP_1 was retained by MWCO 10 kDa, but two RoxP orthologs (e.g., RoxP_8, _11) were not retained by this cutoff and required MWCO 3 kDa to be concentrated.**

**^b^ Expected 8His_TEV_RoxP_1 yield after BE: ≥60 mg from a 500 mL autoinduction culture. Some orthologs did not express as well, like RoxP_6 (RoxP_HypProt_pET22b) that required scaling up of the autoinduction culture volume 24-fold (12.0 L) to purify sufficient protein for analysis.**

### Tag Removal Using pRK793 6His-TEV Protease

Tagless RoxP was then prepared using the following protocol. **<u>Day 9</u>** – ***(1)*** Dilute RoxP protein sample with HSB to 10 mg/mL ^a^; ***(2)*** add 6His-TEV ^b^ to sample; ***(3)*** place sample in a tube rotator at 4°C, and ***(4)*** gently rotate sample overnight (≥16 hr). **<u>Day 10</u>** – ***(1)*** Hard-spin sample (Eppendorf 5430 R, rotor FA-45-30-11, 20817 xg, 10 min, 4°C); ***(2)*** repeat “*Buffer-Exchange (BE).* **<u>Day 9</u>** – ***(7)***” above to assess soluble protein concentration; ***(3)*** add 1/9 sample volume of 10x NiNTA-LB to your sample to adjust chemical/salt concentrations to 18 mM HEPES, 435 mM NaCl, 0.009% azide, 50 m Na phosphate, 10 mM Imidazole; ***(4)*** filter sample; ***(5)*** remove His-tagged proteins from sample by repeating “*Chromatography.* **<u>Day 8</u>**– ***(1-11)***” above with two modifications: current sample substituted for lysate, save Ni-NTA flow-through (FT) containing RoxP-tagless; ***(6)*** assess uncut 8His-TEV-RoxP on SDS PAGE and consider repeating “*Buffer-Exchange(BE)*” above and “*Tag Removal.* **<u>Day 9</u>** – ***(1-6)***” above ^c^; ***(7)*** repeat “*Buffer-Exchange (BE).* **<u>Day 9</u>**” with four modifications: use Ni-NTA FT instead of elution, repeat steps (2-3) three times for >38,416-fold total dilution, add “filter sample” step after (6), calculate the RoxP-tagless (GAT..ALN) concentration using ɛ280 (13075 M^-1^ cm^-1^) and MW (14888.59 g/mol).

**^a^ The optimal RoxP concentration for N-terminal tag removal by Tobacco Etch Virus Protease (TEV) was experimentally determined to be 10 mg/mL, though 1-50 mg/mL will also work. Lower concentrations (<1 mg/mL) required extended incubation times.**

**^b^ His6-Tagged TEV Protease (6His-TEV) was prepared by transforming pRK793 plasmid into *E. coli* strain BL21(DE3)RIL, purifying as previously described** [17]**, and storing single-use, 100 µL aliquots of 11.6 mg/mL 6His-TEV at -80°C. Optimal mass ratio of 6His-TEV to target protein is ∼100:1, so each aliquot was used to cut ≤116 mg of 8His-TEV-RoxP.**

**^c^ For some preparations, a significant amount of uncut RoxP could be recut and prepared as RoxP-tagless.**

## 2. HPLC IMAC Protocol

### Used to prepare apo-RoxP_1-tagless

#### Lysate Preparation

**<u>Day 8</u>** – ***(1)*** After chilling culture to 4°C, spin down *E. coli* cells in 1000 mL polypropylene centrifuge bottle (Cat # # 3120-1000) using an Eppendorf 5920 R centrifuge (rotor S-4x1000, 3428 xg, 30 min, 4°C); ***(2)*** mass the cell pellet (typically ∼18 g per 500 mL autoinduction culture) and then store it in its centrifuge bottle on ice; ***(3)*** pre-chill 42 mL of ddH_2_O in a 50 mL conical (Corning# 352070) to 4°C; ***(4)*** prepare 50 mL of Complete IMAC lysis buffer (C-IMAC-LB) ^a^ by adding each of the following reagents to 42 mL of ddH_2_O: 5.0 ml of 10x IMAC-BB, 0.125 ml of 400x DCS, 0.1 mL of 5.0 M Imidazole (pH 7.8), 50 mg Lysozyme, 0.5 mg DNase, 0.5 ml of 100 mM (100x) AEBSF (ddH_2_O), 0.01 ml of 10 mg/ml (5000x) Aprotinin (HSB), 0.05 ml of 10 mM (1000x) Bestatin (100% methanol), 0.1 ml of 10 mM (500X) Leupeptin (ddH_2_O), 0.1 ml of 1 mM (500x) Pepstatin A (100% ethanol), 0.25 ml of 200 mM (200x) PMSF (100% ethanol), ice-cold ddH_2_O to 50 ml volume; ***(5)*** add C-IMAC-LB to the centrifuge bottle; ***(6)*** use a metal spatula and hand vortexing (swirl bottle) to dislodge the cell pellet from the centrifuge bottle sidewall; ***(7)*** split partially resuspended cell pellet between two 50 mL conical tubes (Corning# 352070), place those tubes in a tube rotator at 4°C, and gently rotate for 15 min to complete cell resuspension in C-IMAC-LB; ***(8)*** during step (7), prepare sonication sample cooling rig (SSCR) ^b^ by placing a pre-chilled (4°C), 100 mL, polypropylene, conical graduated cylinder (CGC) (Fisher# 08-570-20A) in a 1.0 L, EVA Foam, square, ice bucket (Fisher# 03-395-152), pouring 0.925 L of pre-chilled (4°C) aluminum beads (Lab Armor Beads, SHELDON# 42370-04) into the ice bucket to half bury the CGC, submerging two -20°C ice packs into the beads without contacting the CGC, and securing aluminum foil over the beads to seal out room air; ***(9)*** transfer cell suspension to the SSCR, insert a temperature probe (Inkbird# IBS-TH1) into the beads to monitor the SSCR, insert the 0.5 in. sonication probe into the cell suspension, and adjust probe position until only 2-5 mm of liquid separates the probe from CGC walls; ***(10)*** sonicate cells using a Fisherbrand™ Model 505 Sonic Dismembrator (0.5 in. probe; 75% amplitude; 1 sec ON, 20 sec OFF, 3 min total sonication time, 63 min total time); ***(11)*** move the temperature probe into the lysate for 20 sec to measure the post-sonication lysate temperature ^b^; ***(12)*** split the ∼70 mL of lysate between two high centrifugation speed, 50 mL conicals (VWR# 21008-177) and spin out insoluble material using an Eppendorf 5920 R centrifuge (rotor FA-6x50, 20000 xg, 60 min, 4°C); and ***(13)*** filter lysate.

**^a^ Sufficient volume for a ≤50 g cell pellet.**

**^b^ While the temperature of the beads was ∼4°C throughout sonication, the temperature of the lysate rose from ∼4°C to ≤13°C by the end of sonication.**

#### Chromatography

**Day 8** – ***(1)*** Place HPLC IMAC Buffers A and B on the BIO-RAD NGC Discover 10 Pro Chromatography System located in a chromatography refrigerator (4°C); ***(2)*** place lysate in the chromatography refrigerator and then flush NGC Sample Pump with lysate; ***(3)*** IMAC purify lysate over a 5 mL IMAC (HisTrap FF Crude) ^a^ using the following method: lysate application (1 mL/min, forward flow), 10 mM imidazole wash (1% Buffer B, forward flow, 10 CV), 20 mM imidazole wash (2% Buffer B, forward flow, 10 CV), 250 mM imidazole elution (25% Buffer B, reverse flow, 2.46 mL – pre-peak, 10.00 mL – peak, 10.8 mL – post-peak), absorbance data was collected every 1 µL.

**^a^ A 5 mL column was used to avoid having to pass lysate over the column multiple times (see “*Gravity-Flow IMAC Protocol. Chromatography.* <u>Day 8</u> – *(6)*”).**

#### Buffer-Exchange (BE)

**<u>Day 9</u>** – ***(1)*** Filter HPLC IMAC elution into a centrifugal concentrator (Vivaspin 20 mL MWCO 10 kDa ^a^); ***(2)*** centrifuge (Eppendorf 5920 R, rotor FA-6x50, 6000 xg ^b^, 4°C) to concentrate sample to ≤1 mL but >0.5 mL; ***(3)*** add HSB to concentrator to 14 mL mark for a >10-fold dilution into HSB; ***(4)*** repeat steps (2-3) ***<u>three times</u>*** for >38,416-fold total dilution (i.e., BE) of the original buffer (IMAC Buffer: 75% A, 25% B) to reduce imidazole to ≤6.5 µM; ***(5)*** repeat step (2); ***(6)*** hard-spin sample (Eppendorf 5425 R, rotor FA-24x2, 21300 xg, 10 min, 4°C); ***(7)*** measure sample A_280_ (DeNovix DS-11 FX+) to calculate 8His-TEV-RoxP (MDH..ALN) concentration (ɛ280 14565 M^-1^ cm^-1^, MW 17228.04 g/mol) and yield ^c^; and ***(8)*** then store RoxP protein sample at 4°C.

**^b^ GE Healthcare Vivaspin centrifugal concentrators were used for the original gravity-flow IMAC work, and GE recommended a maximum centrifugation speed (× g) of 8000 xg. When that protocol was adapted to an HPLC approach, Sartorius was then the distributor of these centrifugal concentrators, and Sartorius recommended a maximum centrifugation speed (× g) of 6000 xg**

**^c^ Expected 8His_TEV_RoxP_1 yield after BE: ≥60 mg from a 500 mL autoinduction culture.**

#### Tag Removal Using NEB TEV Protease

Tagless RoxP was then prepared using the following protocol. **<u>Day 9</u>** – ***(1)*** Prepare 250 µL of 2.0 mg/mL RoxP in HSB ^a^; ***(2)*** add 28.2 µL of 10x NEB TEV Protease Reaction Buffer (0.5 M Tris-HCl pH 7.5, 5.0 mM EDTA, 10.0 mM DTT); ***(3)*** add 4 µL (40 units) of NEB TEV Protease Enzyme; ***(4)*** place sample in a tube rotator at 4°C, and ***(5)*** gently rotate sample overnight (≥16 hr). **<u>Day 10</u>** – ***(1)*** Hard-spin sample (Eppendorf 5425 R, rotor FA-24x2, 21300 xg, 10 min, 4°C); ***(2)*** repeat “*Buffer-Exchange (BE).* **<u>Day 9</u>** – ***(7)***” above to assess soluble protein concentration in supernatant ^b^; ***(3)*** add 1/9 sample volume (31.4 µL) of 10x IMAC-BB and 0.63 µL of 5.0 M imidazole (pH 7.8) to your sample to adjust chemical/salt concentrations to 16 mM HEPES, 420 mM NaCl, 0.008% azide, 50 m Na phosphate, 10 mM imidazole, pH ∼8; ***(4)*** hard-spin sample (Eppendorf 5425 R, rotor FA-24x2, 21300 xg, 10 min, 4°C); ***(5)*** remove His-tagged proteins from sample using one His-Spin Protein Miniprep (Zymo Research) and collect His-Spin FT containing RoxP-tagless; ***(6)*** perform BE into HSB using 2.0 mL Zeba Desalting Column (Thermo Scientific); ***(7)*** hard-spin sample (Eppendorf 5425 R, rotor FA-24x2, 21300 xg, 10 min, 4°C); ***(8)*** measure sample A_280_ (DeNovix DS-11 FX+) to calculate the RoxP-tagless (GAT..ALN) concentration (ɛ280 13075 M^-1^ cm^-1^, MW 14888.59 g/mol); ***(9)*** store RoxP protein sample at 4°C; and ***(10)*** verify BE and tag removal samples by SDS PAGE.

**^a^ Reaction volume was scaled up as dictated by RoxP-tagless needs of planned experiments.**

**^b^ Only an estimation, as cleaved N-terminal tag is less soluble (i.e., may be in the post-centrifugation pellet) and a small amount of NEB TEV protease is present, though it is likely ≤5 µg per reaction.**

### Small-scale Expression and Purification of Recombinant RoxP

This protein preparation protocol was used to prepare apo-RoxP_1-5/7-8/10-11-tagless and apo-8His-TEV-RoxP_1/6. The pET22b, periplasmic expression constructs for eleven RoxP orthologs (RoxP_1-11, **SI Table 3**, **SI File 3**) were transformed into *E. coli* strain BL21(DE3)RIL, expressed as small-scale autoinduction cultures [1], and purified using IMAC spin columns (Zymo Research His-Spin Protein Minipreps) as outlined in the following protocol. **<u>Day 1</u> –** Streak *E. coli* glycerol stock onto LB-Carb(100 µg/mL) agar plate and grow at 37°C overnight (≥16 hr). **<u>Day 2</u> – *(1)*** Inoculate 5 mL LB-Carb(100 µg/mL) with a single *E. coli* colony and ***(2)*** grow that culture at 37°C overnight (≥16 hr, shaking). **<u>Day 3</u> – *(1)*** Prepare 95 mL of ZYM-5052-Carb (ZYM-5052 supplemented with 2 mM MgSO_4_, 0.5x Trace Metals, 0.1 mM FeCl_3_, and 100 µg/mL Carb) in a 1.0 L flask; ***(2)*** take 95 µL of the LB-Carb(100 µg/mL) overnight culture and dilute it 1000-fold into the 95 mL of ZYM-5052-Carb; ***(3)*** incubate the 1.0L flask at 37°C (shaking) for several hours (∼6 hr) until OD_600_ reaches ≥1 and ≤3; ***(4)*** and then transfer the culture to an 18°C shaker (200 rpm) to grow overnight (≥16 hours). **<u>Day 4</u> – *(1)*** Incubate 10.0 mL of B-PER Complete Protein Extraction (stored at 4°C) in a ≥100 mL RT aluminum bead bath for ≥20 min to raise the temperature of this reagent to RT; ***(2)*** Incubate 25.0 mL of NiNTA-WB in an ice-bath for ≥30 min to prechill it to 4°C; ***(3)*** spin down 70 mL of ZYM-5052-Carb autoinduction culture (Eppendorf 5920 R, rotor FA-6x50; 10000 x g, 10 min, 4°C); ***(4)*** mass the cell pellet ^a^; ***(5)*** prepare 25 mL of RT B-PER with protease inhibitors (BPER-PI) ^b^; ***(6)*** collect cells by centrifugation (Eppendorf 5920 R, rotor FA-6x50, 10000 xg, 10 min, 4°C); ***(7)*** freeze-thaw the cell pellet (-20°C for 30 min then RT until thawed) ^c^; ***(8)*** gently resuspend cell pellet in BPER-PI and incubate with gentle inversion at RT for 15 min; ***(9)*** centrifuge lysate (Eppendorf 5920 R, rotor FA-6x50; 16000 x g, 20 min, 4°C) and transfer supernatant to a new 50 mL conical; ***(10)*** add 10 mL of ice-cold NiNTA-WB to lysate supernatant (1:1 lysate:buffer volume; imidazole final concentration: 10 mM; non-EDTA protease inhibitor final concentration: 1x); ***(11)*** filter sample; ***(12)*** concentrate to 2.0-4.0 mL using a Vivaspin 20 mL MWCO 10 ^d^; ***(13)*** hard spin sample (Eppendorf 5425 R, rotor FA-2x24; 21300 x g, 10 min, 4°C); ***(14)*** filter sample; ***(15)*** purify 8His-TEV-RoxP using one His-Spin Protein Miniprep column (Zymo Research) according to manufacturer’s instructions; ***(16)*** use a 2.0 mL Zeba Desalting Column (Thermo Scientific) to BE the His-Spin elution into PBS; ***(17)*** measure sample A_280_ (DeNovix DS-11 FX+) to calculate 8His-TEV-RoxP concentration (MDH..ALN, ɛ280 14565 M^-1^ cm^-1^, MW 17228.04 g/mol) ^e^ and yield ^f^; ***(18)*** add 26.7 µL of 10x NEB TEV Protease Reaction Buffer; ***(19)*** add 4 µL (40 units) of NEB TEV Protease Enzyme; ***(20)*** place sample in a tube rotator at 4°C, and ***(21)*** gently rotate sample overnight (≥16 hr). **<u>Day 5</u>** – ***(1)*** Add 1/9 sample volume (30.1 µL) of 10x IMAC-BB and 0.60 µL of 5.0 M imidazole (pH 7.8) to the sample to adjust chemical/salt concentrations to 16 mM HEPES, 420 mM NaCl, 0.008% azide, 50 m Na phosphate, 10 mM imidazole, pH ∼8; ***(2)*** hard-spin sample (Eppendorf 5425 R, rotor FA-24x2, 21300 xg, 10 min, 4°C); ***(3)*** remove His-tagged proteins from sample using one His-Spin Protein Miniprep and collect His-Spin FT containing RoxP-tagless; ***(4)*** perform BE into HSB using 2.0 mL Zeba Desalting Column; ***(5)*** measure sample A_280_ (DeNovix DS-11 FX+) to calculate the RoxP-tagless concentration (GAT..ALN, ɛ280 13075 M^-1^ cm^-1^, MW 14888.59 g/mol) ^i^ and yield ^j^; ***(6)*** store RoxP protein sample at 4°C; and ***(7)*** verify BE and tag removal samples by SDS PAGE ^k^.

**^a^ Average cell pellet yield from 70 mL of autoinduction culture grown overnight at 18°C was 1.96 g, which is 78% of the cell pellet yield expected based on the larger scale (500 mL), 5-day at 18°C autoinduction culture.**

**^b^ A 10.0 mL aliquot of RT B-PER Complete Protein Extraction Reagent was supplemented with protease inhibitors aprotinin (4.0 µg/mL), leupeptin (40.0 µM), Pepstatin A (4.0 µM), PMSF (2.0 mM), and EDTA pH 8.0 (2.0 mM). See RoxP large-scale expression and purification protocols above for details on protease inhibitor stocks. Except for EDTA, all protease inhibitors in BPER-PI are at 2x concentration. EDTA in BPER-PI is at 1x concentration.**

**^c^ Thaw normally takes ∼35 min.**

**^d^ RoxP_1 and most other RoxP orthologs were retained by MWCO 10 kDa, but two RoxP orthologs (e.g., RoxP_8, _11) were not retained by this cutoff and required MWCO 3 kDa to be concentrated.**

**^e^ Values for 8His-TEV-RoxP_1.**

**^f^ 8His-TEV-RoxP_1 yield: 1.148 mg/mL in ∼240 µL, 276 µg total ^g,h^.**

**^g^ Total yield limited by amount of His-Affinity Gel per spin column (75 µL), as significant 8His-TEV-RoxP_1 was in the His-Spin FT (SI Fig 14b).**

**^h^ If one accounts for differences in culture volume (500/70 = 7.14), cell pellet mass from 5-days vs. ≥16 hr at 18°C (((((70 mL)/(500 mL))*18 g)/1.96 g) = 1.29), and IMAC volume required to capture the expected ≥60 mg from 5-days at 18°C (2.1/0.075 = 28), then scale up of this small-scale to a 500 mL autoinduction culture grown for 5 days would yield 70.9 mg, which is consistent with ≥60 mg observed in the large-scale protocols.**

**^i^ Values for RoxP_1-tagless.**

**^j^ RoxP_1-tagless yield: 0.418 mg/mL in ∼390 µL, 163 µg total ^g^.**

**^k^ RoxP_6 and RoxP_9 suffered from solubility issues during this protocol. RoxP_6 was not soluble after removal of its N-terminal tag, so work with this ortholog was performed without TEV cleavage. RoxP_9 was not soluble with or without its N-terminal tag, so it was not present on SDS PAGE at either the tagged or tagless stages of this purification.**

### Expression and Purification of Human Siderocalin for Biochemical Assays

Human siderocalin (hSCN) was prepared in a similar manner to RoxP (*E. coli* periplasmic expression followed by IMAC purification), so that it could be used as a negative control protein (NCP) in biochemical assays. This protein is also known as lipocalin 2 (LCN2), neutrophil gelatinase-associated lipocalin (NGAL), or 24p3. SCN is a nutritional immunity protein that sequesters iron (Fe^3+^) coordinated by microbially-produced enterobactin (ENT) or urinary catechols [18]. The hSCNkk_pET22b construct (a kind gift of Dr. Roland Strong, see **SI File 4**) was used in this work, because it (1) directs hSCNkk to the periplasm via a PelB signal peptide and (2) contains two calyx mutations (K125A/K134A) that macroscopically ablate visible Fe-enterobactin (ENT) binding [19]. This protein was also selected as NCP because it has a MW (nascent [MKY-HHH]: 24.5 kDa, post-PelB signal peptide cleavage [MGQ-HHH]: 22.3 kDa, post-thrombin digest to remove the C-terminal 6His-tag [MGQ-VPR]: 21.1 kDa) that is very similar the post-signal peptide cleavage MW of native RoxP (ATP..ALN, 14.8 kDa) and recombinant, apo-RoxP_1-tagless (GAT..ALN, 14.9 kDa).

Recombinant *E. coli* expression of hSCNkk was performed using the following protocol. Unless otherwise stated below, all sample assembly steps include mixing (flick-to-mix five times) and a pulse-spin (3 sec) in a tabletop, miniature (mini) microcentrifuge. **<u>Day 1</u>** – Streak a hSCNkk_pET22b:BL21(DE3) glycerol stock on an LB-Carb(50 µg/mL) plate. **<u>Day 2</u>** – Pick a single colony along the streak to inoculate a 5.0 mL LB-Carb (50 µg/mL) liquid culture to grow overnight (≥16 hr) at 37°C (shaking, 200 rpm). **<u>Day 3</u>** – (***1)*** Add 3.0 mL of the overnight culture to 1.0 L of LB-Carb in a 2.0 L flask; ***(2)*** grow that 1.0 L culture at 37°C (shaking, 200 rpm) for ∼3 hr to reach OD_600_ ∼0.6; ***(3)*** add 1.0 mL of IPTG (Isopropyl ß-D-1-thiogalactopyranoside) to the culture for [IPTG]_final_ of 1.0 mM; ***(4)*** induce for 4 hr at 37°C (shaking, 200 rpm), and then ***(5)*** store the induced culture overnight at 4°C. **<u>Day 4</u>** – ***(1)*** Spin down *E. coli* cells in 50 mL conical tubes (Eppendorf 5810 R, rotor S-4-104, 4°C, 3197 xg, 30 min); ***(2)*** pre-chill 3.5 mL of NiNTA-LB to 4°C; ***(3)*** add 17.5 µL of 100 mM PMSF (in 100% EtOH) to the lysis buffer; ***(4)*** use a 5 mL serological pipet to gently and completely resuspend (e.g., aspirate-dispense 2.5 mL repeatedly without frothing the cell suspension) the cell pellets in lysis buffer; ***(5)*** add 8.75 µL of 400x DCS; ***(6)*** add 70 µL of 50x lysozyme (50 mg/mL, lyophilized protein resuspended in ddH_2_O); ***(7)*** add 3.5 µL of 1000x DNase (10 mg/mL, lyophilized protein resuspended in ddH_2_O); ***(8)*** incubate on ice for 15 min; ***(9)*** sonicate cells packed in ice using a Fisherbrand™ Model 505 Sonic Dismembrator (0.5 in. probe; 20% amplitude; 10 sec ON, 10 sec OFF, 60 sec total sonication time); ***(10)*** split lysate between 1.5 mL microcentrifuge tubes (875 µL lysate/tube); ***(11)*** spin out insoluble material (Eppendorf 5430 R, rotor FA-45-30-1, 20817 xg, 30 min, 4°C); ***(12)*** prepare a BIO-RAD Econo-Column (1.0 × 10 cm, #7371012) with 1.0 mL Ni-NTA buffer-exchanged into lysis buffer by assembling the column (Econo-Column Funnel → Econo-Column [0.5 × 10 cm] → two-way stopcock), adding 2.0 mL of 50% Ni-NTA agarose slurry, allowing the beads to settle over 10-15 min, draining off the 30% (v/v) EtOH storage solution by gravity flow until the meniscus is 1-2 mm above the Ni-NTA column bed, adding 10 mL (10 CV) of NiNTA-LB to the column, settling the beads again for 10 min, and draining the NiNTA-LB until meniscus is 1-2 mm above the column bed; ***(13)*** filter lysate supernatant through a 0.45 µm PES syringe filter (33 mm diameter) into the prepared column; ***(14)*** wash the column (same approach as initial BE into NiNTA-LB) with 4.0 mL (4 CV) NiNTA-WB; ***(15)*** repeat the 4.0 mL NiNTA-WB step for 8 CV of wash split over two steps; ***(16)*** elute the hSCNkk from the column (same approach as initial BE into lysis buffer) using 2.0 mL NiNTA-EB; ***(17)*** repeat the 2.0 mL NiNTA-EB step for 4 CVs of elution split over two steps, and then ***(18)*** evaluate SDS PAGE samples from all expression/purification steps. **<u>Day 5</u>** – ***(1)*** BE the eluted hSCNkk protein into phosphate-buffered saline (PBS) using a Vivaspin 2 mL MWCO 3 kDa centrifugal concentrator (>10,000-fold total dilution of original buffer, same Vivaspin BE strategy as that described above for RoxP); ***(2)*** hard-spin hSCNkk (Eppendorf 5430 R, rotor FA-45-30-1, 4°C, 20817 xg, 10 min) to pellet out any precipitates and/or aggregated protein; ***(3)*** transfer the soluble hSCNkk (supernatant) to a new 1.5 mL microcentrifuge tube; ***(4)*** obtain a protein concentration ^a^ via an absorbance 280 nm (A_280_) on a Tecan Spark using the NanoQuant Plate (optical path length of 0.05 cm; ɛ280 26025 M^-1^ cm^-1^, MW 22280.37 g/mol); ***(5)*** verify all post-Ni-NTA samples on an SDS PAGE gel; ***(6)*** flash-freeze 50 µL aliquots in liquid nitrogen, and then ***(7)*** store frozen aliquots at -80°C.

Tagless hSCNkk was then prepared using the following protocol. **<u>Day 1</u>** – ***(1)*** Add 1 µL 1000 units/mL thrombin (∼0.7 µg, MW ∼36 kDa, resuspended in H_2_O per MP company protocol) to 50 µL of 2.0 mg/mL hSCNkk (100 µg, MW 22.3 kDa) for ∼230-fold molar excess of target:protease; ***(2)*** incubate at RT for 4 hr without inversion; ***(3)*** stop proteolysis by adding 0.5 µL 100 mM PMSF (100% EtOH stock); ***(4)*** BE 20 µL of 50% QIAGEN Ni-NTA agarose slurry ^b^ in HSB via three pulse-spin exchanges in a tabletop microcentrifuge (e.g., 10 µL agarose + 100 µL HSB, invert-to-mix three times, pulse spin for 3 sec at RT, aspirate away supernatant); ***(5)*** add 10 µL of HSB buffer-exchanged Ni-NTA to the thrombin reaction to remove the uncut hSCNkk and free-His-tag; ***(6)*** incubate Ni-NTA with the reaction for 1 hr at RT with gentle inversion; ***(7)*** remove the Ni-NTA via centrifugation (Eppendorf 5430 R, rotor FA-45-30-1, 4°C, 1000 xg, 10 sec); ***(8)*** carefully aspirate the supernatant containing hSCNkk-tagless and transfer to a new pre-chilled tube; ***(9)*** verify samples from all steps via SDS PAGE; and then ***(10)*** obtain a protein concentration via an absorbance 280 nm (A_280_) on a Tecan Spark using the NanoQuant Plate (optical path length of 0.05 cm; ɛ280 26025 M^-1^ cm^-1^, MW 21071.12 g/mol).

**^a^ Expected yield: ∼200 µL of 2.0 mg/mL or 0.4 mg from a 1.0 L culture.**

**^b^ Per QIAGEN, 10 µL of Ni-NTA agarose is sufficient to remove ∼0.5 mg of His-tagged protein, which is 5-fold more protein than the hSCNkk input in this reaction.**

## PROTEIN ASSESSMENT EXPERIMENTS

### SDS PAGE Experiments and Gel Imaging

Unless otherwise noted, all SDS PAGE experiments were run using the following experimental setup: BIO-RAD Any kD™ Mini-PROTEAN® TGX Precast Gel (#4569034, #4569036), 1.0 L of SDS-RB, protein ladder (BIO-RAD Precision Plus Protein Dual Color Standard, #1610374), 300V, 10-15 min ^a^, stained with Coomassie Brilliant Blue (CBB) R-250 (Imperial Stain# 24615). All gel images were obtained on either a BIO-RAD Gel Doc XR or a Thermo Scientific Bright™ CL1500 Imaging System. Images were saved as Tag Image File Format (TIFF) files at ≥800 dots per inch (dpi). TIFF gel image files were used in the assembly of all figures in this paper, and no gel lanes relevant to an experiment were cropped from a gel image presented in this paper. Any gel images with were cropped vertically (i.e., parallel to a lane) were cropped to remove blank lanes, sample lanes that were not part of the described experiment (e.g., a sample added to a gel to check that sample for another lab member’s unrelated project), or the left/right sides of gel before/after the sample wells.

**^a^ Gels were run until the BPB dye (669.96 g/mol) reached the bottom of the gel.**

### Heme-Binding Assays

#### Heme-Agarose (HA) Pulldown

During the development of this HA pulldown assay, 6His-tagged hSCNkk [MGQ-HHH, MW 22.3 kDa] was evaluated as NCP, but it was pulled down by HA in 3 separate experiments. Next, hemoglobin (10 mg/mL stock in HSB, tetramer MW 64.5 kDa, monomer MW 16.1 kDa) was evaluated as a NCP, but it was found to be pulled down due to hemin-exchange (1 experiment), which has been previously reported to occur at a rate of 60% exchange under similar conditions [20]. During the evaluation of potential control proteins, bovine serum albumin (BSA, SIGMA#A-7906) was also shown to be pulled down by HA (1 experiment).

Tagless versions of RoxP_1 and the NCP protein hSCNkk were then prepared to eliminate low-affinity His-tag [21–23]) binding to HA that we previously observed for tagged hSCNkk. This NCP (hSCNkk-tagless, [MGQ-VPR]: 21.1 kDa) has a higher predicted isoelectric point (pI 8.97) compared to apo-RoxP-tagless (pI 6.19), which theoretically could favor non-specific (charge-related) interactions with HA. Fortunately, this potential HA interaction only resulted in minor hSCNkk-tagless binding to HA, which could be eliminated by pre-incubation with hemin (**Fig 4a**, SCN, lane 5).

The final HA pulldown assay that demonstrated RoxP_1-tagless pulldown in biological triplicate was performed as outlined below. NP buffer (10 mM Na phosphate, 0.5 M NaCl, pH 7.5) used in previous HA pulldowns [24] was prepared. For each HA pulldown assay, the following protocol was followed with all steps performed at RT: (1) 20 µL HA (hemin content: 4 µmol/mL, hemin input via HA: 0.08 µmol [≥275-fold molar excess over protein input below) was aliquoted into six 1.5 mL LPBM tubes, (2) each tube of HA was washed with 100 µL of NP buffer three times (e.g., add buffer, invert tube three times, pulse-spin at RT in a miniature microcentrifuge for 3 sec, remove supernatant), and (3) six 100 µL reaction tubes were assembled as shown below.

1. No protein controls: (i) NP buffer + 1% DMSO, (ii) NP buffer + 1% DMSO + 37 µM hemin
2. Control protein samples: Same as (1) with either 4.4 µg hemoglobin (2.7 µM final monomer [heme-binding subunit] concentration in reaction, 0.00027 µmol in reaction) or 6.4 µg hSCNkk-tagless (2.9 µM final concentration in reaction, 0.00029 µmol in reaction).
3. Experimental (RoxP_1-tagless) samples: Same as (1) with 5.0 µg RoxP_1-tagless (3.4 µM final concentration in reaction, 0.00034 µmol in reaction) and 27.5 µM hemin (see below).

#### Note on the reactions assembled above

Input protein masses were chosen to provide similar input protein band staining by Coomassie on an SDS PAGE gel. Molar ratios of HA-hemin:protein were ≥235:1 (e.g., HA-hemin:hemoglobin – 296:1; HA-hemin:hSCNkk-tagless – 276:1; HA-hemin:RoxP_1-tagelss – 235:1). Since our hemin stocks were prepared in DMSO, the [DMSO] was set to 1% by either the addition of DMSO or hemin stock, so that it would be a constant in our experiment. The [hemin] was planned to be 10-fold greater than [protein] for all samples, but an experiment planning error was identified during the assembly of this manuscript that revealed variable [hemin] excesses of 14-, 13-, and 8-fold for hemoglobin, hSCNkk-tagless, and RoxP_1-tagless, respectively. While pre-incubation with excess hemin was evaluated for RoxP_1-tagless in triplicate, co-incubation of HA with RoxP_1-tagless and hemin was not performed. Therefore, heme-exchange for RoxP during HA binding cannot be excluded, which might account for a lack of observed heme competition in **Fig 4a** (RoxP lane 5).

After assembly of the reaction tubes, the reaction tube samples were (4) incubated without rotation for 30 min at RT, (5) buffer-exchanged into NP buffer using a 0.5 mL Zeba column (Thermo Scientific), and then (6) added to the previously NP buffer-exchanged HA tubes. Sample+HA tubes were then (7) inverted three times to mix, (8) incubated (without rotation) at RT for 2 hours, and (9) pulse-spun (miniature microcentrifuge, RT, 3 sec) to pellet the HA. (10) The supernatant was then aspirated away from the HA to evaluate unbound protein, and (11) the HA was washed with 1.0 mL NP buffer five times (e.g., add buffer, invert tube three times, pulse-spin in a miniature microcentrifuge for 3 sec at RT, remove supernatant). After washing the HA, (12) a final centrifugation (Eppendorf 5430 R, rotor FA-45-30-1, 8000 xg, 2 min, RT) followed by (13) aspiration of supernatant was performed to remove as much liquid as possible. Then, (14) boiled-bead samples were prepared (40 µL 1x SDS LB+BME added to HA, flick-to-mix five times, incubated at 95°C for 10 min, pulse-spun for 3 sec in miniature microcentrifuge at RT) and (15) loaded onto BIO-RAD Tris-Gly AnyKd for SDS PAGE analysis (12.5 µL sample per lane) to evaluate HA pulldown in each sample.

#### NATIVE Gel Shift Assay

In brief, RoxP-porphyrin binding was assessed using NATIVE gel electrophoresis. Porphyrin ligand titration of RoxP used molar ratios (porphyrin:RoxP) that ranged from 0.03:1 to 67:1. Each sample component (e.g., RoxP, individual porphyrins) was also alone run to evaluate their NATIVE PAGE migration. Human hemoglobin (Hgb, theoretical pI 8.13), human hemopexin (Hpx, theoretical pI 6.43), bovine serum albumin (BSA, theoretical pI 5.60), and bovine pancreatic deoxyribonuclease (DNase, theoretical pI 5.08) were used as control proteins on some NATIVE gels to evaluate their migration as a function of heme-binding and pI. All theoretical pI values were calculated with ExPASy ProtParam using a protein’s primary sequence after (1) signal peptide cleavage and (2) pro-peptide removal (if relevant). Hemopexin and BSA showed heme-binding dependent NATIVE gel-shifts by CBB and tryptophan protection by 2,2,2-trichloroethanol (TCE) staining [25,26] (data not shown).

Unless otherwise noted, proteins were incubated with and without porphyrin ligands for 30 minutes under aerobic conditions. Samples were then loaded onto a precast 12% PAGE gel lacking SDS, which then underwent non-denaturing electrophoresis at 4°C. Electrophoresis was performed with standard forward polarity (FP) for all proteins and reverse polarity (RP) for those proteins with high pI values that might prevent gel entry at pH 8.3 (see section below: <u>Electrophoresis Polarity Variation)</u>. Gels were then imaged using (1) BIO-RAD stain-free technology to detect exposed tryptophans (TCE staining)*, (2) CBB (Imperial Protein Stain) to visualize proteins, and (3) ODS (heme stain) to identify heme-containing bands.

\***Apo-RoxP W66 is partially-surface-exposed (Fig 2) and thus can be modified by TCE.**

### Standard NATIVE Gel Shift Assay Protocol

#### Standard Preparation for NATIVE PAGE Electrophoresis Protocol

Unless otherwise noted, all NATIVE gel shift assays in this work used the following preparation for NATIVE PAGE gel electrophoresis protocol conducted in a cold room set to 4°C. (1) Pre-chill 1.0 L of 1x NATIVE-RB overnight to 4°C. (2) Rinse out the gel rig, gel box, and buffer dam (if needed to run only one gel) with diH_2_O to rid them of any SDS from prior electrophoresis runs. (3) Repeat rinse (2) if any salt crystals or detergents bubbles were present prior to rinsing. (4) Rinse the sides of two pre-cast gels (12% Mini-PROTEAN TGX Stain-Free Protein Gel, BIO-RAD #4561044) with diH_2_O, remove the combs from both gels, and then diH_2_O rinse the gel wells. (5) Use a P200 gel-loading tip to straighten any gel spacers between wells that were bent by comb removal and/or diH_2_O rinsing. (6) Place all electrophoresis equipment/consumables (gel rig, gel box, power supply, two 12% pre-cast gels, buffer dam if only running one gel) in the cold room to chill for ≥30 min. (7) Assemble the gel rig with two 12% gels. (8) Pour the entire 1.0 L of NATIVE-RB into gel-rig allowing it to rinse the gel wells and then overflow into the gel box. (9) Examine the gel for leaks. (10) Prepare gel samples and start their 30 min incubation (see below).

#### NATIVE Gel Shift Assay Binding Reaction Assembly

Unless otherwise noted, each sample was assembled using the following protocol. Thaw porphyrin-DMSO stocks at RT, pulse spin (miniature microcentrifuge, max speed, 3-5 sec), and store at RT to prevent DMSO from forming a solid at 4°C. Thaw hematin stocks at RT, pulse spin (miniature microcentrifuge, max speed, 3-5 sec), and then store on ice. Assemble each sample on ice in a pre-chilled (4°C), 1.5 mL LPBM tube (#90410). Each sample is an 84 µL sample containing: 0.9x HSB ^a^, 10% (v/v) DMSO, +/-9.4 µg of protein (e.g., 7.5 µM RoxP), +/- porphyrin at a molar ratio to protein ranging from 0.03-67:1. After assembling each sample, flick-to-mix each sample three times, pulse-spin (Eppendorf 5430 R, 1000 xg, 10 sec, 4°C), and then store on ice until all samples are assembled. After assembly of all samples, incubate samples at room temperature (RT) for 30 min. This assembly plan was used to prepare a sample that could then be adjusted to 1x NATIVE-SB by adding 28 µL 4x NATIVE-SB, and then that 112 µL samples could be used to load two gels with 50 µL/lane where each lane would contain 4.2 µg of input protein. This input protein mass per lane was chosen to ensure robust gel staining using the three chosen gel staining techniques (CBB, TCE, ODS).

**^a^ Porphyrin-DMSO stocks were diluted 10-fold into HSB to create 4.5-5.0 mM porphyrin in 10% DMSO and 0.9x HSB (18 mM HEPES, 135 mM NaCl, 0.009% (m/v) Na azide, pH 7.4).**

#### Standard NATIVE PAGE Electrophoresis Protocol

(11) After sample RT incubation is started, start a pre-run of the gels (120V, 0.5 hour, 4°C). (12) After samples have finished their RT incubation, place all samples on ice, add 28 µL of 4X NATIVE-SB to each sample (final volume 112 µL), flick-to-mix each tube three times, pulse-spin all samples (Eppendorf 5430 R, 1000 xg, 10 sec, 4°C), and place samples back on ice. (13) Load 50 µL of each sample into the same well/lane of the two gels, and load all lanes without sample (i.e., blank lanes) with an equivalent volume (50 µL) of 1x NATIVE-SB. (14) Run the gels at 120V for 2 hr ^a^. (15) Disassemble the gel rig, extensively rinse both gel cassettes in diH_2_O, open each cassette, and place each gel in a weigh boat containing 100 mL of diH_2_O. (16) Carefully, use two gloved fingers to pin each gel to the bottom of its weigh boat, and then pour out the diH_2_O. (17) Add 100 mL of diH_2_O to each weigh boat and then repeat (16). (18) Repeat (17) for three total 100 mL diH_2_O rinses. (19) Take one gel to the BIO-RAD Gel Doc XR and activate (via ultraviolet [UV] light) the proprietary trihalo compound in the gel (TCE staining ^b^) using the stain-free imaging protocol below. (20) Return the TCE imaged gel to its weigh boat containing 100 mL diH_2_O, repeat (16), and then add 200 mL of diH_2_O to the weigh boat. (21) For the TCE imaged gel, follow the <u>Imperial Protein Stain Protocol</u> below. For the other gel, follow the <u>Heme Stain Protocol</u> below.

**^a^ These NATIVE PAGE run conditions result in BPB migrating to the bottom of the gel. BPB runs at the same apparent MW as unbound hemin.**

**^b^ These BIO-RAD gels contain a proprietary trihalo compound (similar to TCE) that is activated by UV light and reacts with tryptophans in the gel. As such, TCE in this manuscript refers to this technique using the BIO-RAD proprietary trihalo compound.**

##### Tri-halo Stain-Free Imaging Protocol

New Protocol è Application Settings: Gel Imaging è Protein Gels è Stain Free Gel è Gel Activation Time: Best sensitivity (5 min)è Imaging Area: Select gel type: Bio-Rad Mini-PROTEAN Gel è Image Exposure: Faint band èImage color: Gray.

### Imperial Protein Stain Protocol

Gently shake the gel in 200 mL of diH_2_O on an orbital gel shaker for 15 min. (2) Use two gloved fingers to pin the gel the bottom of the weigh boat, pour out the diH_2_O, and absorb any remaining diH_2_O with a Kimwipe. (3) Add sufficient Imperial Protein Stain to the weigh boat to barely cover the gel (∼25 mL), gently shake the gel on an orbital gel shaker for 10 min ^a^, use two gloved fingers to pin the gel the bottom of the weigh boat, and then pour out the Imperial Protein Stain. (4) Add 100 mLdiH_2_O to the weigh boat, gently shake the gel on an orbital gel shaker for 1 min, and repeat (2). (5) Repeat (4) once for three total brief (1 min) diH_2_O washes. (6) Add 100 mL diH_2_O to the weigh boat, note the diH_2_O fluid depth, ball up three Kimwipes and place them in a corner of the weigh boat, repeat Kimwipe “ball” assembly and placement twice, add sufficient diH_2_O to reach the previously noted diH_2_O depth, and then destain the gel in diH_2_O overnight. (7) Go to a gel imager and take an image. (8) Store the gel in diH_2_O.

***^a^ Imperial staining for NATIVE Gel Shift Assays was performed for 10 min, but staining for some other gels in this work (e.g., SDS PAGE) was extended (e.g., 2 hours, overnight) to increase stain sensitivity, as suggested in the company protocol*.**

### Heme Stain Protocol

Adapted from prior publications [27,15]. All rinse/wash steps were performed in a sturdy, polypropylene container (e.g., P200 micropipette tip box or lid) with gentle shaking (i.e., no splashing) on an orbital gel shaker. (1) Rinse the gel in diH_2_O for 15 min, use two gloved fingers to pin the gel the bottom of the weigh boat, pour out the diH_2_O, and absorb any remaining diH_2_O with a Kimwipe. (2) Wash the gel with 30 ml of 10% trichloroacetic acid (TCA) for 10 min. (3) Pour 10% TCA into a TCA/H_2_O waste bottle in a chemical hood. (4) Wash the gel with 30 ml of diH_2_O for 10 min, dispose of diH_2_O water wash in the TCA/H_2_O waste bottle. (5) Repeat step (4) for two total 30 mL diH_2_O washes. (6) While performing steps (4-5), prepare 30 ml of heme stain (50 mM sodium citrate [pH 4.4], 6.0 mM *o*-dianisidine-2HCl, 180 mM* [0.55 w/w %] H_2_O_2_) by mixing together 23.4 mL diH_2_O, 3.0 mL 0.5 M sodium citrate (pH 4.4), 3.0 ml 10x *o*-dianisidine-2HCl (0.06 M), and 0.6 ml 30% (8.8 M) H_2_O_2_. ***Particulate matter will be visible in the stain after vortexing the components together.*** (7) After performing step (5), use a Kimwipe to absorb any remaining diH_2_O from the side of the gel, pour 30 ml of heme stain on gel, and shake for 30 min. ***Bands should be visible within 5-10 min^**,***^.*** (8) Dispose of heme stain in the heme stain waste bottle. (9) Wash the gel with 30 ml of 5% acetic acid for 10 min and then dispose of wash in an acetic acid wash waste bottle in a chemical hood. (10) Wash the gel with 30 ml of diH_2_O for 10 min and then dispose of wash in acetic acid wash waste bottle. (11) Repeat step (10) twice for three total 30 mL diH_2_O washes. ***Multiple washes are required to remove particulate matter from the heme stain.*** (12) Go to a gel imager and take an image. (13) Store the gel in diH_2_O.

**\*Allhorn heme stain protocol** [15] **listed 18 mM H^2^O^2^, but their stain solution assembly would have resulted in 180 mM H^2^O^2^. The other staining protocol used to prepare the above protocol** [27] **used 419 mM H^2^O^2^. As such, we believe that the 18 mM H^2^O^2^ was a typo in Allhorn et al. (2016).**

**\*\*TCE Porphyrin Staining:**

- **Hemin appears as white bands in TCE staining of a NATIVE gel, which we hypothesize is due to heme-generated free radicals reacting with TCE in a manner similar to how the UV-activated tryptophan indole ring (six-membered benzene ring fused to a five-membered pyrrole ring) reacts with TCE** [26].
- **Since UV-activated tryptophan indole ring reacts with TCE reacts and metal-free porphyrins PPIX and CPIII react with TCE, we hypothesize that TCE is modifying the inner part of their porphyrin rings i.e., their five-membered pyrrole rings.**
- **CPIII has two additional carboxylates compared to hemin, which we hypothesize leads it to migrate more quickly through the gel than hemin. Hemin also has a more positive charge due to its iron atom, which would be expected to slow migration through the gel. These attributes of CPIII would also be expected to have it have a less intense reaction with CBB, which we did observe. The latter situation is hypothesized to be due to CBB in acidic conditions binding to basic and hydrophobic residues of proteins (or other macromolecules) leading to a change in color from a dull reddish-brown to intense blue.**
- **The TCE staining profile of PPIX is more complex likely due to varying TCE sensitivity of various protonated states of PPIX separated via the glycine-chloride gradient established during electrophoresis.**

**\*\*\*W66 protection from TCE modification could be assessed for heme, but it could not be assessed for PPIX due to the TCE(+) bands in PPIX-only control lanes that migrate at the same apparent MW as holo-RoxP.**

### Electrophoresis Polarity Variation

The pI of apo-RoxP_1-tagless (pI 6.19) allowed it to enter a 12% NATIVE gel run using the standard electrophoresis conditions described above (e.g., pH 8.3 NATIVE-RB, FP). Heme-binding by four RoxP orthologs (1, 5, 6, 8; pI range 6.19-9.56) also allowed their holo-forms to enter a 12% NATIVE gel run using these standard conditions.

The pI of three of these RoxP orthologs (5, 6, 8) are higher than NATIVE-RB (pH 8.3), which impeded the entry of their apo-forms into a NATIVE gel run using standard conditions. To address this issue, high pI RoxP orthologs underwent NATIVE gel electrophoresis using both FP and RP as described below.

***Apo-RoxP_5-tagless*** (pI 9.56) did not enter the NATIVE gel under standard FP conditions (120 V, 2 hr, 4°C). RP electrophoresis (250 V, 8.5 hr*, 4°C) led apo-RoxP_5-tagless to traverse nearly the entire NATIVE gel (band at the bottom of the gel, data not shown), so a shorter run time (2.6 hr) was trialed which clearly showed (i) apo-RoxP_5-tagless migration into the gel and (ii) disappearance of this band with titration of hemin, which supported heme-binding by this ortholog (data not shown).

**\*E1 error (no load detected) noted at 15 hr, but evidence of gel running until 8.5 hr was available. Gel bands were able to diffuse due to lack of current for ∼6.5 hr.**

***Apo-RoxP_6-tagless*** (pI 9.56) did not enter the NATIVE gel under standard FP conditions. RP electrophoresis (250 V, 8.5 hr*, 4°C) did not indicate that apo-RoxP_6-tagless had entered the NATIVE gel (data not shown), though band diffusion prior to staining and/or excessive destaining may have impeded identification of that band. RP electrophoresis (250 V, 2.6 hr, 4°C) of apo-RoxP_6-tagless did allow it to enter the NATIVE gel (top of the gel, data not shown) and disappearance of this band with titration of hemin supported heme-binding by this ortholog (data not shown).

**\*E1 error (no load detected) noted at 15 hr, but evidence of gel running until 8.5 hr was available. Gel bands were able to diffuse due to lack of current for ∼6.5 hr.**

***Apo-RoxP_8-tagless*** (pI 8.98) barely enters the gel (e.g., slight staining of sides and bottom of wells) under standard FP conditions (**Fig 4g**; Gel#2; lanes 2,4). RP electrophoresis showed a run time-dependent increase (120V; 1 vs. 4 hr, 4°C) in apo-RoxP_8-tagless gel migration, but further run time extension to 16 hr did not increase apo-RoxP_8-tagless migration into the gel (data not shown).

**\*E1 error (no load detected) noted at 22.5 hr, but evidence of gel running until 20 hr was available. Gel bands were able to diffuse due to lack of current for ∼2.5 hr. It was not clear why this gel appeared to run longer (20 vs. 8.5 hr) before having an E1 error under the same conditions as described above (RP, 250V).**

#### Soret peak analysis

Absorbance scans (300-800 or 170-810 nm) were performed on tagless proteins (1) alone and (2) in the presence of heme. Reported molar ratios were rounded to the closest whole number. After blank (solution alone) subtraction, individual components (e.g., protein alone, heme alone) were subtracted from the binding reaction (protein + heme) to identify the absorbance spectrum unique to protein-heme binding, which allowed identification of Soret peaks (370-420 nm) and the smaller Q bands (500-700 nm). Two established heme-binding proteins, hemopexin (Hmp) and bovine serum albumin (BSA) were included as controls. Several experimental conditions were evaluated in this Soret peak analysis work. Those experiments are listed below (Author[s] – Experiment Date) with experimental differences noted and references to the corresponding manuscript figures listed.

##### WHM – 12/27/2018

The reaction buffer for all samples was 0.44% (v/v) DMSO in HSB. Samples were initially set up as 100 µL volumes containing twice the planned, final protein concentration. Control proteins Hmp (0.5 mg/mL, 57 kDa, HSB + PBS) and BSA (10 mg/mL, 67 kDa, HSB) were run as positive controls. Hmp in this experiment was SIGMA #H9291-500UG (Hemopexin from human plasma), which had a product insert that listed an Hmp MW of 57 kDa. Hmp from human serum has a MW of 63 kDa (unmodified polypeptide: 49.3 kDa, 0.6 kDa for the galactosamine oligosaccharide, 12.5 kDa the five glucosamine oligosaccharides) [28]. The Hmp MW of 63 kDa was used in our experiment planning and analysis.

This Hmp stock ships as lyophilized protein that contributes 1x PBS (11.9 mM phosphate, 137 mM NaCl, 2.7 mM KCl, pH 7.4) when resuspended at 0.5 mg/mL. This stock was resuspended in HSB. As such, the Hmp samples contained 0.44% (v/v) DMSO, HSB, and 0.42x PBS (5.0 mM phosphate, 57.8 mM NaCl, 1.1 mM KCl, pH 7.4).

Final DMSO concentration (0.44%) and protein:hemin molar ratios were dictated by the hemin (4.5 mM) and protein stock solutions available at that time. The protein:hemin molar ratios were: 10:1 RoxP:hemin, 8:1 BSA:hemin, 1:3 Hmp:hemin. RoxP in this experiment was RoxP_1-tagless. Samples were assembled without hemin in pre-chilled, 1.5 mL, LPBM tubes; flicked-to-mix three times; pulse spun (Eppendorf 5430 R, Rotor FA-45-30-11, 1000 xg, 4°C, 10 sec); and then transferred to a Corning 96-well, not treated, clear, flat bottom microplate.

Immediately prior to starting the absorbance scans, 100 µL of 20 µM hemin (0.44% (v/v) DMSO in HSB) was added to the heme-containing samples, while all other samples received 100 µL of 0.44% (v/v) DMSO in HSB. All heme-containing samples had a final concentration of 10 µM hemin. All samples were then mixed using a P200 multi-channel micropipettor (aspirate-dispense 100 µL three times), placed in the TECAN Spark microplate reader, and then analyzed on a TECAN Spark microplate reader using the protocol below. Data from this experiment is presented in: **Fig 4b**; **SI Fig 2a,c**.

#### Soret Peak Absorbance Scan Protocol #1

Using a TECAN Spark microplate reader, absorbance scans (300-800 nm, bandwidth 3.5, step size 2) were performed every 1 min for 5 min, every 5 min for 25 min, every 10 min for 30 min, and then every 30 min for 120 min. The total experiment time was ∼3.3 hr. Though the TECAN Spark was set to RT (i.e., no plate heating), the recorded temperature averaged 27.37±0.13°C, which was ∼5°C above RT. All wells were blank subtracted (0.44% [v/v] DMSO, HSB) and then the hemin alone well was subtracted. Protein alone wells were not subtracted in this initial experiment from their respective RoxP+hemin binding reactions. Of note, none of the examined protein alone samples, HSB, or PBS have significant absorbance in the examined absorbance range of 300-800 nm.

##### WHM – 07/08/2019

Same as “WHM – 12/27/2018” except (1) only examined RoxP_1-tagless, (2) reaction buffer was 1.0% (v/v) DMSO in PBS, (3) final hemin concentration was 50 µM, (4) final apo-RoxP_1-tagless concentration was 10 µM, and (5) RoxP:hemin molar ratio was 1:5. Data from this experiment is summarized in **SI Table 5**.

##### SBS – 07/08/2019

Same as “WHM – 12/27/2018” except: (1) reaction buffer was 10.0% (v/v) DMSO in 0.9x PBS^*,**^; (2) final hemin concentration was 12.5 µM; (3) protein:hemin molar ratio of 1:1 was examined due to the assessment of a novel fluorescent-heme ligand in the same experiment (not reported in this paper) which was only available as a 100 µM stock in DMSO; (4) RoxP/BSA stock solutions were in HSB and thus resulted in slight changes to the final sample buffers (RoxP samples: 0.06x HSB, 0.94x PBS; BSA samples: 0.07x HSB, 0.93x PBS); (5) samples were assembled in the 96-well microplate without protein and then mixed (P200 aspirate-then-dispense 100 µL five times); (6) immediately prior to starting the experiment, protein was added to the relevant wells and mixed (P200 aspirate-then-dispense 100 µL five times); and (7) the Tecan protocol below was used. Data from this experiment is presented in: **SI Fig 2b,d,e**.

**\*Base solution was changed from HSB to PBS to match Hmp stock solution after reconstitution (Cat # MBS143111, 1.22 mg/mL in PBS, 17.4 µM based on MW of 70 kDa listed on the product insert).**

**\*\*Blank in this experiment was accidentally assembled as only 1x PBS, rather than the planned assembly of 20 µL DMSO + 180 µL PBS for 10.0% (v/v) DMSO in 0.9x PBS. All protein:hemin samples underwent subtraction of hemin alone, thus removing absorbance from solution components (e.g., PBS, HSB, DMSO) and hemin alone. As such, this mistake in blank assembly did not impact our final Soret analysis. Also, a comparison of the (SI Fig 2b,d,e) Hmp/BSA/RoxP difference spectra to the (Fig 4b) blank-subtracted RoxP alone spectra near 300 nm suggests that the protein alone raw spectra in SI Fig 2b,d,e is due to DMSO.**

#### Soret Peak Absorbance Scan Protocol #2

Using a TECAN Spark microplate reader, absorbance scans (300-800 nm, bandwidth 3.5, step size 2) were performed every 1 min for 120 min (2 hr). Though the TECAN Spark was set to RT (i.e., no heating), the recorded temperature during the experiment averaged 27.71±0.02°C, which as ∼5°C above RT. All wells were blank subtracted (0.5% [v/v] DMSO, PBS). Then, each protein:hemin binding reaction had hemin alone subtracted. Protein-only samples containing 10% DMSO were supposed to be blank-subtracted (10.0% (v/v) DMSO in 0.9x PBS), but that blank sample was not available. As such, protein-only sample spectra have absorbance descending from a maximum at 300 nm to ∼425 nm, which is favored to be DMSO rather than protein (see above).

##### MRV – 07/18/2019

Same as “SBS – 07/08/2019” except: (1) planned reaction buffer was 0.25% (v/v) DMSO in HSB; (2) blank contained 0.25% (v/v) DMSO in HSB (0.5 µL DMSO + 199.5 µL HSB); (3) only RoxP_8-tagless ortholog was examined; (4) protein stock was in HSB so that final binding reaction solution had the same DMSO and HSB concentrations; (5) the experiment was performed in biological duplicate; (6) the RoxP:hemin molar ratio was 1:1; (7) protein:hemin samples were assembled initially without hemin; (8) samples were prepared in separate 1.5 mL LPBM tubes, flicked to mix three times, pulse spun (miniature microcentrifuge, 3-5 sec, RT), transferred to the 96-well microplate, hemin was added to the protein:hemin samples, all wells were mixed by pipetting (aspirate-then-dispense 100 µL five times), the 96-well plate was placed in the TECAN Spark microplate reader, and then the Tecan protocol below was used. Data from this experiment is presented in: **SI Fig 2f**.

#### Soret Peak Absorbance Scan Protocol #3

Using a TECAN Spark microplate reader, absorbance scans (300-800 nm, bandwidth 3.5, step size 2) were performed every 1 min for 120 min (2 hr). Though the TECAN Spark was set to RT (i.e., no heating), the recorded temperature during the experiment averaged 27.91±0.02°C, which was ∼5°C above RT. All wells were blank subtracted (0.25% [v/v] DMSO, PBS). Data processing for RoxP:hemin samples included (1) subtraction of RoxP(blank-subtracted) from the RoxP:hemin(blank-subtracted) and then (2) subtraction of hemin(blank-subtracted) from the RoxP:hemin(blank-subtracted, RoxP(blank-subtracted)). This data processing strategy was designed to remove all confounding absorbances (DMSO, HSB components, unbound hemin, apo-protein absorbance*) leaving only heme-binding-specific Soret peaks and Q-bands.

**\*There is essentially no apo-RoxP protein absorbance beyond 300 nm. (Fig 4b, SI Fig 2 f-h).**

##### GWP – 12/20/2021

Adaptation of microplate Soret protocols above to cuvette assay using DeNovix DS-11 FX+ Spectrophotometer/Fluorometer. Samples were 1.0 mL 0.25% (v/v) DMSO in HSB, and they contained a range of hemin concentrations (0.0, 2.8, 5.6, 11.3 µM) and apo-RoxP_1-tagless concentrations (0.0, 22.9, 38.2, 76.5 µM). Stock solutions were: 4.5 mM hemin in DMSO, 35.9 mg/mL apo-RoxP_1-tagless in HSB. The molar ratios of RoxP:hemin examined were: 7:1, 8:1, 14:1. Each sample was assembled at RT in 1.5 mL LPBM tubes; inverted five times to mix; pulse spun (miniature microcentrifuge, 3-5 sec, RT); transferred to a disposable cuvette (#14-377-016); and incubated at RT incubation for 30 min. Then, a DeNovix DS-11 FX+ was blanked against 0.25% (v/v) DMSO in HSB, and an absorbance scan (170-810 nm) was performed on each sample. The cuvette’s poly(methyl methacrylate) (PMMA) plastic did not allow for measurement below 220 nm, and the light path (Z height 8.5 mm) did not require the minimum cuvette volume of 1.5 mL. In addition to the scan, absorbance at specific wavelengths (280, 415, 518, 546, 646 nm) were measured. After this measurement, the 30 min RT incubation and subsequent absorbance measurements were repeated three times for scans at 60-, 90-, and 120-min total reaction time. Data processing included (1) subtraction of RoxP(blank-subtracted) from the protein:hemin(blank-subtracted) and then (2) subtraction of hemin(blank-subtracted) from the protein:hemin(blank-subtracted, RoxP(blank-subtracted)). This data processing strategy was designed to remove all confounding absorbances (DMSO, HSB components, unbound hemin) leaving only heme-binding-specific Soret peaks and Q-bands. Data from this experiment is presented in: **SI Fig 2g**.

##### GWP – 01/11/2022

Same as “GWP – 12/20/2021” except: (1) Hematin was used instead of hemin; (2) a working hematin solution was prepared by diluting 0.226 M hematin in 1.4 N NaOH to 9.04 mM hematin using HSB for a final stock of 9.04 mM hematin, 56 mM NaOH, 0.96x HSB; (3) a single hematin concentration was examined (5.625 µM); and (4) and absorbance measurements were taken at time 0 min prior to RT incubation. There was no DMSO in this experiment. Data from this experiment is presented in: **SI Fig 2h**.

### Size-exclusion chromatography

Several chromatography systems and size-exclusion chromatography (SEC) columns were used in this study. All columns were calibrated with ≥3 MW standards to generate MW calibration curves to estimate protein MWs (see **SI Fig 5a**,**b** for example calibration curves). Ovalbumin (SIGMA#A-5253) required SEC purification prior to use as a MW standard. The running buffer for all SEC runs was 1x HSB (20 mM HEPES pH 7.3, 150 mM NaCl, 0.01% Na azide) unless otherwise indicated. All SEC runs were run at 4°C and applied as a 0.5 mL sample to an ∼24 mL column. Those experiments are listed below (Author[s] – Experiment Date) with experimental differences noted and references to the corresponding manuscript figures listed.

#### Experiment Details on the SEC Evaluation of RoxP-ligand Binding

##### WHM – 12/18/2018

This experiment was performed as follows to examine how a 10-fold molar excess of hemin affects RoxP_1 SEC elution: (1) Assemble two 900 µL 0.9xHSB, 10% (v/v) DMSO SEC samples containing 0.67 mg/mL (45 µM) apo-RoxP_1-tagless +/- 450 µM hemin; (2) mix gently (flick with a finger three times); (3) pulse-spin (Eppendorf 5430 R, rotor FA-45-30-11, 1000 xg, 10 sec, 4°C); (4) incubate for 30 min on ice (4°C); (5) hard spin (Eppendorf 5430 R, rotor FA-45-30-1, 20000 xg, 10 min, 4°C) to pellet insoluble material (e.g., aggregated protein, unbound hemin); (6) transfer to a new, pre-chilled LPBM tube; (7) if insoluble material was observed in (5) (e.g., +hemin sample), then repeat (5-7) until no aggregate is visualized; (8) syringe filter (0.2 µm, 4 mm diameter, PVDF, # SLGV004SL) sample into a new, pre-chilled LPBM tube; (9) load 0.5 mL loop on the HPLC (AKTA Purifier 100); and (10) run the sample over the S75 Increase column. Absorbance data was collected every 1 µL with column run at 0.5 mL/min. Data was plotted in **Fig 4e** as 50 µL per datapoint to clearly show individual data points. Note, HPLC fraction collection was started at 4.72 and 8.25 mL for apo- and holo-RoxP runs, respectively.

Initial NATIVE PAGE (45 µL SEC fraction + 15 µL 4x NATIVE-SB, 50 µL loaded per lane) clearly showed holo-RoxP gel-shift (**SI Fig 5c**), but the gel-shift bands were faint due to the main holo-RoxP peak (∼5.3 mL wide) being broader due to ≥2 overlapping peaks, as compared to the monodispersed, main apo-RoxP peak (∼2.7 ml). The suboptimal holo-RoxP peak separation at 71 kDa is due in part to this this column’s ideal fractionation range of 3000 to 70000 daltons. The apo-RoxP peak fractions also appeared overloaded, as there was an apo-RoxP doublet (**SI Fig 5c**, lane 3,4).

A 500 µL aliquot of holo-RoxP peak fractions 2-6 were then concentrated 5-fold using a centrifugal concentrator (Vivaspin 500 µL, MWCO 3, PES). New NATIVE and SDS PAGE samples were assembled containing 5-fold of the initial fraction material for the holo-RoxP peak fractions, as compared to the apo-RoxP fractions. SDS PAGE gel and two NATIVE PAGE gels (#1: TCE then CBB; #2: ODS) were then run on these samples. The SDS PAGE CBB band intensity of the best matched SDS PAGE samples were compared (**Fig 4e**) and suggest that the holo-RoxP A_280_ HPLC trace is predominantly (∼94%) due to hemin bound to RoxP_1. Slight offsets of these fractions from the observed A_280_ peaks were due to a larger fraction size (1.0 mL) compared to the column volume (V_c_) of ∼24 mL and sample dilution post-column (e.g., PEEK tubing, A_280_ detector) prior to reaching the fraction collector.

##### SBS – 04/29/2019

This experiment was performed as follows: (1) Assemble three 900 µL, 0.9xHSB, 10% (v/v) DMSO SEC samples containing 0.67 mg/mL (45 µM) apo-RoxP_1-tagless alone, apo-RoxP + 500 µM hemin (HD, hemin in DMSO); and apo-RoxP + 500 µM protoporphyrin IX (PPIX); (2) mix gently (invert five times); (3) pulse-spin (Eppendorf 5430 R, rotor FA-45-30-11, 1000 xg, 10 sec, 4°C); (4) incubate for 30 min on ice (4°C); (5) hard spin (Eppendorf 5430 R, rotor FA-45-30-1, 20000 xg, 10 min, 4°C) to pellet insoluble material (e.g., aggregated protein, unbound hemin)*; (6) transfer to a new, pre-chilled LPBM tube; (7) syringe filter (0.2 µm, 4 mm diameter, PVDF, # SLGV004SL) sample into a new, pre-chilled LPBM tube; (8) load 0.5 mL loop on the HPLC (AKTA Purifier 100); and (10) run each sample over the analytical S200 column (AS200, Superdex^TM^ 200 10/300 GL). The column was run at 0.5 mL/min, and absorbance data was collected every 1 µL. Samples were run in the following order: (1) RoxP:PPIX, (2) RoxP:HD, (3) RoxP:DMSO. A small front shoulder was observed for RoxP:DMSO (data not shown), which was likely holo-RoxP formed from trace hemin remaining on the column from the previous RoxP:HD run. See “GWP/ALM – 11/19/2023-11/29/2023” for more information on trace hemin absorbed to SEC columns and how to remove it.

**\*No pellet was observed for the sample without HD/PPIX (i.e., DMSO-only control), while small and large pellets were observed for the HD and PPIX samples, respectively.**

##### GWP/ALM – 11/19/2023-11/29/2023

The following experiment was performed to examine how a 10-fold molar excess of hemin affects the SEC elution of four RoxP orthologs (RoxP_4, 5, 7, 11). RoxP_1 was included as a control in this experiment. This experiment was performed for each ortholog as follows: (1) Assemble two 1000 µL SEC samples in HSB (**RoxP alone**: 2 µM apo-RoxP-tagless; **RoxP:hemin**: 2 µM apo-RoxP-tagless, 20 µM hemin, 0.4% DMSO*); (2) mix gently (flick with a finger five times); (3) pulse-spin (Eppendorf 5425 R, rotor FA-24X2, 1000 xg, 10 sec, 4°C); (4) incubate for ≥ 16 hr at 4°C with rotation (10 rpm); (5) hard spin (Eppendorf 5425 R, rotor FA-24X2, 20000 xg, 10 min, 4°C) to pellet insoluble material (e.g., aggregated protein, unbound hemin); (6) transfer supernatant to a new, pre-chilled LPBM tube; (7) syringe filter (0.2 µm, 4 mm diameter, PVDF) sample into a new, pre-chilled LPBM tube; (8) load 0.5 mL HPLC loop on the Bio-Rad Discover 10 Pro; and (9) run the sample at 0.5 mL/min over the Bio-Rad ENrich™ SEC 70 column with a total SEC run volume of 1.3 column volumes (CV).

Absorbance data was collected at 215, 280, 374, and 500 nm. Absorbance at 215 nm was used to monitor small molecules (e.g., HEPES, DMSO), protein, and hemin. Absorbance at 280 nm was used to monitor protein/hemin. Absorbance at 374/500 nm was used to monitor hemin, Soret peak, and Q bands. HSB samples containing only 20 µM hemin (+0.4% DMSO*) were also run to evaluate the elution profile of hemin alone, which resulted in no significant absorbance in any measured wavelength (e.g., 280 nm: ± 1 mAU, 374 nm: ± 3 mAU). Lack of hemin SEC absorbance was due to the insolubility of hemin in aqueous solution, which likely explains its adherence to SEC columns as trace hemin**. All samples were run at least in duplicate. The order of SEC runs in this experiment was: (1) removal of trace hemin from the SEC column**, (2) run all apo-RoxP samples, (3) repeat trace hemin removal, (4) run all holo-RoxP samples, (5) run all hemin-alone samples, and (6) repeat trace hemin removal. Data is presented in **Fig 4h**.

* **DMSO was contributed by the stock of 5.0 mM hemin in 100% DMSO. The original experimental plan included “RoxP alone” samples that included 0.4% DMSO, but an assembly error led to the omission of DMSO in the final “RoxP alone” samples. Prior SEC of an apo_RoxP_1-tagless with 10% DMSO (SBS – 04/29/201) did not significantly change RoxP’s elution profile (data not shown).**

**\*\* <u>Removal of Trace Hemin from SEC 70.</u> During the development of a RoxP:ligand SEC binding assay, trace hemin was found to remain on SEC columns after the application of a hemin-containing sample, like RoxP:hemin or hemin alone. Subsequent runs with heme-binding proteins would then demonstrate very small. heme-specific peak(s). Due to this observation, the following protocol was developed to remove trace hemin from a SEC 70 column: (1) Assemble one 1000 µL SEC sample of 20 mg/mL (300 µM) BSA in HSB; (2) invert to mix five times; (3) hard spin (Eppendorf 5425 R, rotor FA-24X2, 20000 xg, 10 min, 4°C) to pellet insoluble material (e.g., aggregated protein); (4) transfer supernatant to a new, pre-chilled LPBM tube; (5) syringe filter (0.2 µm, 4 mm diameter, PVDF) sample into a new, pre-chilled LPBM tube; (6) load 0.5 mL HPLC loop on the Bio-Rad Discover 10 Pro; and (7) run the sample at 0.5 mL/min over the Bio-Rad ENrich™ SEC 70 column (total SEC run: 1.3 CV); (8) run 3 more CV of HSB over the Bio-Rad ENrich™ SEC 70 column at 0.5 mL/min; and then (9) flush the 0.5 mL HPLC loop on the Bio-Rad Discover 10 Pro with 20 mL of HSB at 2.5 mL/min.**

### Analytical ultracentrifugation (AUC)

This experiment was performed on RoxP_1-tagless as follows: (1) Assemble one SEC sample with a 30:1 molar ratio of hematin:RoxP_1 (1000 µL 0.9x HSB, 55.6 mM Tris pH 7.0, 61.9 mM NaOH, 10 mM hematin, 0.34 mM apo-RoxP_1-tagless); (2) mix gently (invert to mix five times); (3) pulse-spin (Eppendorf 5425 R, rotor FA-24X2, 1000 xg, 10 sec, 4°C); (4) incubate for 2.0 hr at RT; (5) hard spin (Eppendorf 5425 R, rotor FA-24X2, 20000 xg, 10 min, 4°C) to pellet insoluble material (e.g., aggregated protein, unbound hemin); (6) transfer to a new, pre-chilled (4°C), LPBM tube; (7) syringe filter (0.2 µm, 4 mm diameter, PVDF) sample into a new, pre-chilled (4°C), LPBM tube; (8) load 0.5 mL HPLC loop on the Bio-Rad Discover 10 Pro; and (9) run the sample at 0.5 mL/min over the Bio-Rad ENrich™ SEC 70 column (total SEC run: 1.3 CV). Absorbance data was collected at 280, 300, 374, and 500 nm. See “GWP/ALM – 11/19/2023-11/29/2023” for an explanation of the selection of the 280, 374, and 500 nm wavelengths. Absorbance data at 300 nm was collected as an alternative wavelength for monitoring heme.

To identify holo-RoxP fractions that could be used to prepare AUC samples at defined absorbances (1.0, 0.5, 0.3) for a specified wavelength, A_374_ was assessed for holo-RoxP SEC fractions using a DeNovix DS-11 FX+ spectrophotometer. Apo-RoxP_1-tagless protein stock dilutions were also prepared and assessed at A_280_ to prepare apo-RoxP AUC samples. Protein concentrations for apo-RoxP_1-tagless (with disulfide present) samples were calculated using the following A_280_ concentration constants: Abs 0.1% [=1 g/L] = 0.878, ε_280_ = 13075 M^-1^ cm^-1^.The apo-RoxP_1 protein stock used in this experiment was IMAC-purified but not SEC-purified, which may explain a small amount of aggregate (<2%) seen by AUC (data not shown, see further details below).

SDS PAGE was then used to assess the holo-RoxP SEC fractions and determine the approximate RoxP concentration in these fractions. Absorbance at 280 nm of holo-RoxP could not be used to calculate the holo-RoxP SEC fraction protein concentration due to both protein and hemin absorbing at 280 nm. Control apo-RoxP_1 gel samples were prepared from the apo-RoxP_1-tagless protein stock and included on this gel. All apo-/holo-RoxP gel samples were >95% pure by SDS PAGE.

CBB staining of RoxP in the apo-RoxP lanes was compared to the holo-RoxP fraction lanes to determine the approximate RoxP protein concentration in the holo-RoxP fractions. Control samples loaded a defined protein mass per lane (835, 209, 52, 13 ng), which allowed for an approximation of RoxP protein concentration in holo-RoxP fractions (e.g., P7/6: 2.5 mg/mL [168.0 µM], P7/8: 0.4 mg/mL [28.0 µM],). Fraction P7/8 was used for the holo-RoxP A_374_ 1.0 sample, while fraction P7/6 was used for the holo-RoxP A_374_ 0.5 and 0.3 samples. The approximate MW of these fractions from SEC MW calibration were: P7/6: 43.3 kDa, P7/8: 27.0 kDa. The increased protein concentration and SEC-estimated MWs in AUC samples from P7/6 support the higher order oligomerization (pentamer) observed in **Fig 4f** for the A_374_ 0.5 and 0.3 samples.

AUC samples were then assembled from the apo-RoxP protein stock and holo-RoxP peak fractions as follows: (1) Assemble three 1.2 mL apo-RoxP_1-tagless samples in HSB on ice with a final A_280_ concentrations of 1.0 (1.14 mg/mL, 76.48 µM), 0.5 (0.57 mg/mL, 38.24 µM), and 0.3 (0.34 mg/mL, 22.94 µM); (2) assemble three 1.2 mL holo-RoxP_1-tagless:hematin samples in HSB on ice with a final A_374_ of 1.0 (0.19 mg/mL, 13.11 µM), 0.50 (0.21 mg/mL, 14.42 µM), and 0.30 (0.13 mg/mL, 8.69 µM); (3) mix all samples gently (invert to mix five times); (4) hard spin (Eppendorf 5425 R, rotor FA-24X2, 20000 xg, 10 min, 4°C) to pellet insoluble material (e.g., aggregated protein, unbound hemin); (5) transfer supernatant to a new, pre-chilled LPBM tube; (6) syringe filter (0.2 µm, 4 mm diameter, PVDF,) sample into a new, pre-chilled LPBM tube; (9) ship samples (6 proteins samples, 7.0 mL of HSB sample buffer) overnight at 4°C* to the University of Connecticut (UCONN), Center for Open Research Resources & Equipment (COR²E); and (10) have the UCONN COR^2^E Biophysics CORE Director (Heidi Erlandsen, Ph.D.) perform AUC sedimentation velocity experiments as outlined below.

AUC was performed on a Beckman-Coulter Optima AUC analytical ultracentrifuge with double sector cells (pathlength 1.2 cm) equipped with sapphire windows for absorbance scans at 280 nm and 374 nm. Prior to ultracentrifugation, the rotor was equilibrated under vacuum at 20°C for ∼1 hour. Then, the rotor was accelerated to 45,000 RPM, and sedimentation velocity analysis was conducted in one run (20°C, 45,000 RPM, scans acquired every 20 seconds, ∼12-hour total run time). AUC data was analyzed using the: program - Sedfit (version 16.36); method - direct boundary modeling program for individual data sets using model-based numerical solutions to the Lamm equation [29]; model - continuous sedimentation coefficient distribution, c(s), with resolution of 0.05 S and maximum entropy regularization confidence limits (55% holo-RoxP, 68% apo-RoxP); and plot normalization - c(s) distributions were normalized to the maximum c(s) peak height. Buffer density and viscosity were calculated using Sednterp [30,31] to be 1.006 g/mL and 0.0103091 poise (centimeter-gram-second [cgs] unit of viscosity) at 20°C, respectively.

Apo-RoxP-tagless used in this experiment was not HPLC purified prior to the preparation of the AUC samples. Multiple SEC runs of this apo-RoxP protein preparation previously demonstrated a single, mono-dispersed peak at the MW of monomeric apo-RoxP. While the holo-RoxP AUC sample underwent SEC, the apo-RoxP AUC sample did not undergo SEC, which likely explains the AUC observation of a minor percentage of aggregates (<2%).

* **Samples were packaged immediately prior to shipment as follows: (1) two frozen (-20°C), 12 oz. ULINE Cold Packs were placed in the bottom of a small Styrofoam cooler; (2) two 18-inch pieces of 1-inch lab tape were used to make an “X” with their adherent sides facing up; (3) a pre-chilled (overnight at 4°C), 12 oz. ULINE Cold Pack was placed on top of the center of tape strip “X”; (4) AUC samples were placed in a Ziploc bag (1 Quart), the air was evacuated from the bag, the bag was sealed and then folded to minimize its area, and a piece of 1-inch lab tape bag was wrapped around the bag; (5) the bag was placed in the center of the pre-chilled cold pack; (6) a second pre-chilled (overnight at 4°C), 12 oz. ULINE Cold Pack was placed on top of sample bag; (7) the two free pieces of tape were then used to secure the pre-chilled cold packs to one another to prevent sample movement during shipping; (8) a third frozen (-20°C), 12 oz. ULINE Cold Packs was placed in the lid of the small Styrofoam cooler; (9) the lid of the cooler was affixed to the bottom of the cooler with packing tape; and (10) then samples were taken directly to FedEx (≤15 minute transit time) to be shipped overnight to UCONN. The five cold packs (exterior x3 – frozen, internal x2 – pre-chilled) sandwiched the samples to maintain them at 4°C**, while effectively filling the entire interior of the Styrofoam cooler to avoid sample movement during shipping.**

**\*\* Prior to the UCONN shipment, this packing strategy was evaluated using a SensorPush temperature sensor as a mock sample in a Ziploc bag, which showed that this strategy maintained 4°C for >24 hours when the cooler was stored at RT.**

#### Differential Scanning Fluorimetry (DSF)

Assessment of protein apparent melting temperature (T_ma_) using SYPRO™ Orange Protein Gel Stain (excitation/emission 470/570 nm) and two real-time PCR detection system (BIO-RAD CFX96 Real-Time System, QuantStudio™ 5 Real-Time PCR [384-Well Block]) was performed as previously described [32] with the following modifications. At least 2 replicates were performed for each DSF experiment. All RoxP proteins used in DSF experiments were tagless and in their apo-form. Those experiments are listed below (Author[s] – Experiment Date) with experimental differences noted and references to the corresponding manuscript figures listed.

##### WHM/SBS/JZT – 2019

To identify the optimal signal-to-noise ratio for RoxP DSF samples, DSF relative fluorescence unit (RFU) signal for RoxP_1 was initially monitored in all five CFX96 channels (channel#, example fluorophore, excitation range, emission range; 1, FAM, 450-490 nm, 510-530 nm; 2, HEX, 515-535 nm, 560-580 nm; 3, Texas Red, 560-590 nm, 610-650 nm; 4, Cy5, 620-650 nm, 675-690 nm; 5, Quasar 705, 672-684 nm, 705-730 nm). Based on the RFU signal, all subsequent experiments monitored RFU in the HEX channel. The ratio of RoxP protein concentration (µM) to SYPRO™ Orange stock concentration (5000x initial stock) was also varied (5-96 µM, 5-100x) to identify optimized ratios for each RoxP ortholog (RoxP_1: 72 µM:30x; RoxP_5/6: 96 µM:25x). Initial RoxP_1 thermal shift assays were run with 60 µM RoxP_1 and 5x SYPRO™ Orange Protein Gel Stain, while later runs were with 72 µM RoxP_1 and 30x SYPRO™ Orange Protein Gel Stain. Samples were assembled as either 50 µL (RoxP_1) or 25 µL (RoxP_5,6) samples in 1.5 mL LPBM tubes immediately prior to the experiment, mixed gently (flicked with a finger five times and then pulse spun). Samples were then pipetted into BIO-RAD Hard-Shell 96-Well PCR Plates (low profile, thin wall, skirted, white/clear #HSP9601), plates were sealed with BIO-RAD Microseal ’B’ PCR Plate Sealing Film (adhesive, optical #MSB1001), plates were vortexed lightly, pulse spun (800 xg, 25°C, 2 min), and then incubated at room temperature (23.6°C) for 30 min prior to performing the thermal shift assay in a BIO-RAD CFX96 Real-Time System. All RoxP proteins examined were tagless. The melting protocol used was a 5 min hold at 10°C, 10°C to 98°C temperature ramp (RoxP_1: 0.5°C/cycle, RoxP_5,6: 1.0°C/cycle) with 1 min hold at each temperature, followed by return to 25°C to examine if denaturation was reversible (∼100 min total experiment time).

##### WH/GWP – 2024

Based on our initial DSF work in 2019, DSF was then used to determine the effect of pH on RoxP_1, 5, and 11 thermal stabilities. The 2019 DSF strategy above was adapted as outlined below to be run on a QuantStudio™ 5 Real-Time PCR with 384-Well Block. Initially, the DSF T_ma_ of each RoxP ortholog was assessed at pH 7.4 using HSB. Then, the effect of pH on T_ma_ was examined using a sample buffer containing 150 mM NaCl and a Citrate: BIS-TRIS propane (CBTP) buffer system (pH 2.30-9.81, **SI Table 11**). A 40 µL sample comprising 70 µM protein and 25x SYPRO Orange was prepared in each sample buffer. The protein, dye, and buffer were combined via gentle pipetting into a 1.5 mL LPBM tube on ice, and then 10 µL of each sample was dispensed into three individual wells (technical triplicates) of a 384-well qPCR plate (Thermo Scientific# AB1384). Following sample preparation, the plate was sealed with an adhesive sealing sheet (Thermo Scientific# AB0558), centrifuged (Eppendorf 5920 R, 3000 xg, 4°C, 3 minutes) to eliminate air bubbles, and placed in the QuantStudio™ 5 Real-Time PCR machine. A 30-minute room temperature incubation was not performed prior to running the real-time PCR experiment. Fluorescence intensity was continuously monitored using emission #3 (excitation 539-561 nm, emission 576-596 nm; similar to Hex). The temperature was initially stabilized at 10°C for 2 minutes and subsequently ramped from 10°C to 99°C at a rate of 0.05°C per second run over 168 cycles (∼32 min per experiment). DSF data analysis was conducted using DSFworld (https://gestwickilab.shinyapps.io/dsfworld/) [33] to assess each protein’s fitting curves and determine its apparent melting temperature (T_ma_). DSFworld model #2 was used for RoxP_1 and RoxP_5, but RoxP_11 required the use of DSFworld model #4 to fit its two transitions. While color intensity of SYPRO Orange solutions have been reported to decrease below pH ∼5 [34], SYPRO Orange fluorescence increased when RoxP underwent thermally-induced unfolding. Each DSF sample was assessed in technical triplicate. Four biological replicates (performed in technical triplicate) of apo-RoxP_1-tagless in HSB were assessed by DSF, while apo-RoxP_5/11-tagless were assessed in HSB once in technical triplicate. The effect of pH on each ortholog was assessed in technical triplicate one time (single biological replicate). The findings of these experiments shown in **Fig 3d-f** and **SI Fig 6** reproduced what had been previously observed in “<u>WHM/SBS/JZT – 2019</u>” experiments for HSB T_ma_ values for RoxP_1/5, as well as a pilot pH-dependence of apo-RoxP_1-tagless T_ma_ performed in a similar buffer system.

## ANTIBODY EXPERIMENTS

### Antibody generation and modification

RoxP_1-tagless prepared using the “<u>Autoinduction</u> → *Gravity-Flow IMAC Protocol”* was SEC-purified on a GE Healthcare Superdex 75 Increase 10/300 GL and then submitted to GenScript to generate murine anti-RoxP monoclonal antibodies. Supernatants from twenty anti-RoxP hybridomas were prepared by GenScript and then screened by the McCoy Lab to identify non-competitive antibodies. Hybridoma screening was performed using biolayer-interferometry (BLI) on an Octet using Anti-Mouse Fc Capture (AMC) biosensors (data not shown). Four antibodies (2H2F3, 3E10E3, 4D8H1, 5F3D11) appeared to be non-competitive by BLI, and they were selected for further work at GenScript that included (1) frozen preparation of hybridoma stocks, (2) purification of milligram (mg) quantities (2 mg, 15 mg) of each antibody from mouse ascites using protein A, and variable domain sequencing of each hybridoma (**SI File 5**). Hybridoma cell lines, purified antibodies, sequencing information, and all other pertinent related data were then sent to the McCoy Lab. Antibodies were biotinylated using an EZ-Link NHS-PEG4-Biotinylation Kit (Thermo Scientific). Anti-RoxP antibody isotypes: IgG2b,κ (2H2F3, 3E10E3, 5F3D11); IgG2b,λ (4D8H1)(

### Enzyme-linked Immunosorbent Assay (ELISA) Experiments

Unless stated otherwise, the following statements apply to all experiments in this section.

1. All solutions are 1x.
2. All experiments used high-binding, clear, half-area plates (Greiner, Cat # 675061).
3. ELISA Coating Buffer (ELISA-CB): 50 mM Carbonate (pH 9.6).
4. ELISA Wash Buffer (ELISA-WB): 1x PBS + 0.1% (v/v) Tween-20, a.k.a. PBST.
5. ELISA Blocking Buffer (ELISA-BB): 1% (w/v) BSA in PBST
6. Blocking was performed by applying 50 µL/well of ELISA-BB and incubating at RT for 1 hr.
7. Other than antigen coating stocks, all other protein dilutions were prepared in ELISA-BB.
8. All plate incubations were performed without shaking.
9. All washes were 100 µL/well ELISA-WB, a.k.a. PBST.
10. All experiments were run using technical triplicates and performed ≥2 times.
11. When plotted, all error bars are the standard deviation (SD) of technical triplicates.
12. OD at 450 nm (OD_450_) was measured for all plates*.
13. OD at 570 nm (OD_570_) was measured for some plates and used to calculate the delta (OD_450_-OD_570_)*.
14. Blue color development was only monitored visually initially, but then later it was monitored both visually and at 650 nm (A_650_).
15. For some plates, data was normalized to a positive control (e.g., 100% for apo-RoxP_1-tagless) and a negative control (e.g., 0% for a “blank” well coated with ELISA-CB that did not contain any protein).

\***While both wavelengths were routinely measured, the OD^570^ was very low for all wells and displayed very little range from well-to-well (e.g., 0.0468 ± 0.0047 across an entire 96-well plate), which resulted in the subtraction to calculate a delta (OD^450^-OD^570^) being essentially a constant subtracted from all values that was equivalent to ∼20% of the buffer-alone negative control. As such, some experiments only monitored OD^450^ and did not calculate a delta (e.g., Fig 6c).**

#### Indirect ELISA (iELISA)

Development of an anti-RoxP iELISA used RoxP_1-tagless that was prepared using the “<u>Autoinduction</u> → *Gravity-Flow IMAC Protocol”* and then diluted from >30 mg/mL (HSB) to 1000 ng/mL in ELISA-CB. ELISA plates were then coated with 50 µL/well of RoxP_1-tagless (**Fig 6a** basic iELISA:1000 ng/mL; adaptations of iELISA [**Fig 6b**,**f**; **SI Fig 15**]: 500-10000 ng/mL), and the plates were incubated at 4°C overnight. After three ELISA-WB (PBST) washes, the plates were blocked with 50 µL/well ELISA-BB for 1 hr at RT. After removing the blocking solution, wells were washed four times with PBST, and then 50 µL/well of primary antibodies were added to their respective wells and incubated at RT for 1 hour. For **Fig 6a** iELISA experiments, the following antibody titrations were evaluated: **2H2F3** (2.3-5000 ng/mL), **3E10E3** (4.6-10000 ng/mL), **4D8H1** (1.1-2500 ng/mL), and **5F3D11** (1.1-2500 ng/mL). Plates were washed three times with PBST, and then a 1:5000 dilution of secondary antibody biotin-Goat anti-mIgG (BioLegend; Cat # 405303) was added to all wells and for 1 hr at RT. After washing three times with PBST, 50 µL/well of 1:20000 diluted **(Fig 6a, SI Fig 15a)** Streptavidin-HRP (SA-HRP; GenScript, Cat #M00091) or (**Fig 6f**, **SI Fig 15b**) Streptavidin Poly-HRP (SA-Poly-HRP; Thermo Scientific, Cat # 21140) was dispensed into each well and incubated at RT for 1 hour. After washing three times with PBST, color was developed by adding 50 µL/well TMB substrate (BioLegend; Cat # 421101), blue color development was monitored visually and by A_650_, and then the reaction was stopped by adding 25 µL/well TMB stop solution (BioLegend; Cat # 423001). Finally, the OD of each well was measured on a plate reader. Using GraphPad Prism (version 8.4.3 for Mac, GraphPad Software, Boston, Massachusetts USA, www.graphpad.com), nonlinear regression was used to calculate EC50 and EC90 values for each antibody using a plot of log[mAb] vs. response (OD_450_). EC90 95% confidence interval (CI) is shown in **Fig 6a**, and the ECF values from EC90 fits was used in iELISA ortholog specific experiments (**Fig 6f**, **SI Fig 15**). The iELISA shown in **Fig 6a** was performed in technical triplicate and biological duplicate. For evaluations of the effect of biotinylation on anti-RoxP antibody antigen recognition, the iELISA shown in **Fig 9** evaluating all four biotinylated anti-RoxP antibodies was performed in technical duplicate, a subsequent experiment titrating RoxP coating concentration against set antibody concentrations (**2H2F3**: 500ng/ml, **4D8H1**: 250ng/ml; **5F3D11**: 250ng/ml) was performed in technical duplicate, and a comparison of two b-2H2F3 lots was performed in technical triplicate.

#### Use of iELISA to Assess Anti-RoxP Antibody Specificity for RoxP Orthologs

These experiments were performed as described above for “*<u>Indirect ELISA (iELISA)</u>*” with the following modifications. For RoxP ortholog iELISA experiments (**Fig 6f**, **SI Fig 15**), recombinant RoxP_1-11 was prepared using the “**Small-scale Expression and Purification of Recombinant RoxP**” as 0.2 to >1.0 mg/mL (HSB) stocks and then diluted to 1-10 µg/mL in ELISA-CB. RoxP ortholog preparation concentrations were ascertained by A_280_ using ExPASy ProtParam ε280 for the post-TEV removal of the N-terminal tag, and then SDS PAGE was used to confirm that iELISA coating samples prepared using these concentrations contained equivalent amounts of protein. RoxP ortholog iELISA was performed in technical triplicate for each coating antigen concentration and HRP detection reagent (SA-HRP, SA-Poly-HRP). Three separate experiments were performed (**Fig 6f** – 10.0 µg/mL coating concentration, SA-Poly-HRP; **Fig 15a** – 1.0 µg/mL coating concentration, SA-HRP; **Fig 15b** – 1.0 µg/mL coating concentration, SA-Poly-HRP). RoxP_9 was not soluble, and so it was not able to be assessed in this experiment. RoxP_6 became insoluble after removal of its N-terminal His-tag, so RoxP_6+Tag was compared to RoxP_1+Tag. All other RoxP orthologs were tagless (RoxP_2-5,7,8,10,11-tagless) and compared to apo-RoxP_1-tagless. Apo-RoxP_1-tagless prepared using the (1) “<u>Autoinduction</u> → *Gravity-Flow IMAC Protocol”* and (2) “**Small-scale Expression and Purification of Recombinant RoxP**” protocol were compared to one another (**SI Fig 15a**), and they displayed equivalent binding for all antibodies. For RoxP ortholog iELISA experiments (**Fig 6f**, **SI Fig 15**), the concentrations of the primary antibodies were the ECF90 values calculated by iELISA (see **Fig 6a**): 2H2F3 (330 ng/mL), 3E10E3 (2900 ng/mL), 4D8H1 (155 ng/mL), and 5F3D11 (2100 ng/mL).

#### Competition iELISA (ciELISA)

For ciELISA experiments, the iELISA protocol was used with the following modifications: apo-RoxP_1-tagless was coated on plates using two RoxP concentrations in ELISA-CB: 500 ng/mL (b-4D8H1 plate), 1000 ng/mL (b-2H2F3 and b-5F3D11 plates); serially diluted unlabeled (27-20000 ng/mL) antibodies and biotinylated monoclonal antibodies (b-mAbs) were added separately at the primary antibody step, no secondary antibody was used because detection was based on biotinylated anti-RoxP antibody. Biotinylated anti-RoxP antibody concentrations were b-2H2F3 (500 ng/mL), b-4D8H1 (250 ng/mL), and b-5F3D11 (250 ng/mL). The level of biotinylation per antibody was b-2H2F3 (lot #1: 11 biotins/IgG, #2: 10 biotins/IgG,), 3E10E3 (11 biotins/IgG), 4D8H1 (18 biotins/IgG), 5F3D11 (23 biotins/IgG). This experiment was performed in technical triplicate using three different maximum competitor mAb concentrations (2000, 5000, and 20000 ng/mL) diluted either 2- or 3-fold to evaluate 7 different competitor concentrations (e.g., 20000-27 ng/mL), and all three experiments displayed the same competition pattern shown in **Fig 6a** and **SI Fig 10**.

#### Sandwich ELISA (sELISA)

ELISA plates were coated with 50 µL/well capture antibody 4D8H1 (2.0 µg/mL in ELISA-CB) and incubated at 4°C overnight. After three washes with PBST, the plates were blocked with 50 µL/well blocking solution for 1 hr at RT. After removing the blocking solution, wells were washed three times with PBST. Then, 50 µL/well of antigen (e.g., serially diluted RoxP ^a^ from 0.5-400 ng/mL ^b^) was added and incubated for 1 hr at RT. Plates were washed three times with PBST. After that, plates were incubated with 50µL/well detection antibody b-2H2F3 (lot #1: 5 µg/mL, lot #2: 250.0 ng/mL) ^c^ for 1 hr at RT. After washing three times with PBST, 50 µL/well of 1:5000 diluted Streptavidin-HRP (SA-HRP; GenScript, Cat #M00091) or Streptavidin Poly-HRP (SA-Poly-HRP; Thermo Scientific, Cat # 21140) was dispensed into each well and incubated at RT for 1 hour. After 3 washes with PBST, color was developed by adding 50 µL TMB reagents (BioLegend; Cat # 421101) for 30 min, and then the reaction was stopped by adding 25 µL TMB stop solution (BioLegend; Cat # 423001). Finally, OD_450_ and OD_570_ were measured for each well in the plate, and the delta (OD_450_-OD_570_) was calculated. Experiments were performed in technical triplicate and biological replicates (N) are indicated. The SA-HRP version of this assay demonstrated a linear RoxP standard curve from 12.5-400 ng/mL (N=7), 12.5-500 ng/mL (N=2), 12.5-600 ng/mL (N=3), and 6.25-600 ng/mL (N=1). The SA-Poly-HRP version of this assay demonstrated a linear RoxP standard curve from 0.5-15 ng/mL (N=1), and this experiment suggested that lower concentrations would have been detectable (see below).

Then, the Limit of Detection (LOD), defined as ≥3-fold the standard deviation (SD) of the blank above the blank value, and Lower Limit of Quantitation (LLOQ), defined as ≥10-fold the SD of the blank above the blank value, was determined for a set of apo-RoxP_1-tagless ^a^ samples creating a standard curve (e.g., a three-fold serial dilution of RoxP from 0.5-400 ng/mL in ELISA-BB). For SA-HRP experiments (fifteen biological replicates performed in technical triplicate), LOD ∼10 ng/mL and LLOQ ∼40 ng/mL were routinely observed, and the average values of 7 biological replicates performed specifically on the dilution series of 12.5-400 ng/mL RoxP were LOD 9 ± 5 ng/mL and LLOQ 40 ± 10 ng/mL. For SA-Poly-HRP experiments (technical triplicate), both LOD and LLOQ were <0.5 ng/mL, as the value for the lowest RoxP concentration (0.5 ng/mL) was well above the values for LOD and LLOQ.

**^a^ RoxP_1-tagless in sELISA experiments was prepared using the “Autoinduction** è ***Gravity-Flow IMAC Protocol”* and then diluted from >30 mg/mL (HSB) into PBST/biofluid.**

**^b^ This range was adjusted based on the goals of an experiment. For example, the sar-sELISA experiments did not require lower concentrations when preparing a sELISA standard curve, as the spike in concentration was 100 ng/mL.**

**^c^ Antibody lot assessments: The optimal b-2H2F3 concentration sELISA was assessed for each lot, and a slightly higher degree of biotinylation of lot #1 b-2H2F3 is believed to account for the higher concentration required for that lot.**

#### Spike-and-Recovery Sandwich ELISA (sar-sELISA)

The sELISA strategy above was followed with the following modifications. For the RoxP spike-and-recovery sELISA experiment (sar-sELISA), each complex fluid (matrix) was diluted two-fold into ELISA-BB to assess a dilution series from undiluted (neat) to either 1/128 (all samples) or 1/256 (synovial fluid only). Evaluated complex fluids were two types of *C.* acnes culture medium (RCM, BHI), human serum, human cerebrospinal fluid (CSF), and bovine synovial fluid (bSF), and human synovial fluid (hSF). All complex fluids (matrices) were evaluated after being thawed once from an aliquot that had been previously flash-frozen in liquid nitrogen and then stored at - 20°C. In addition to this assessment, bSF was evaluated a second time alongside a freshly-thawed aliquot of hSF, and that latter experiment displayed increased bSF matrix effect which may have been due to the -20°C freeze-thaw between experiments (compare **Fig 6d** to **SI Fig 11d**). After the creation of each matrix dilution series, a 2.0 µg/mL stock of apo-RoxP_1-tagless in ELISA-BB was used to spike in 100 ng/mL RoxP into each diluted complex fluid sample. These samples were added to the plate during the antigen addition step. A RoxP sELISA standard curve (12.5-400 ng/mL, 2-fold dilution series) in ELISA-BB (performed in technical triplicate per plate) and RoxP at 100 ng/mL (technical duplicate for each sample) were also included to determine which dilutions replicated the signal of 100 ng/mL RoxP. For most assays, the mock 100 ng/mL RoxP matched that predicted by the standard curve (105.50±16.69 ng/mL), so sELISA values interpolated as ng/mL were equivalent to percentage (%) recovery. For one assay (**Fig 6d**), the mock 100 ng/mL RoxP wells were calculated to be at a higher concentration (120.45±3.97 ng/mL), so those sar-sELISA values were normalized to the average empirical spiked concentration.

Prior to interpolation of complex sample dilutions, the dilution-specific blank without spiked RoxP was subtracted to remove any possible contribution of RoxP in the samples at baseline. Of note, all complex fluids had dilution-specific blank values equivalent to PBST alone except for pooled human serum, which appeared to have 22.25 ng/mL RoxP. To determine if pooled human serum in fact has RoxP, an independent assessment would need to be performed, which our lab is currently evaluating. Percentage recovery versus dilution factor (Dil.) and percentage matrix was then plotted with 100% ± 20% highlighted to identify the minimum dilution necessary for a matrix to minimize complex sample (e.g., biofluid, culture medium) matrix effects. Dilutions for each complex fluid for subsequent assays were: 4-fold dilution (human serum, bSF, BHI, RCM), 8-fold dilution (hCSF, hSF).

#### Linearity and Detection Limits of sELISA in Complex Samples

The sELISA strategy above was followed with the following modifications. All sELISA in complex samples work was performed with SA-HRP, which reduced sensitivity as described above (see comparison of SA-HRP to SA-Poly-HRP). Also, the SA-HRP used in this experiment had been stored at 4°C for an extended period of time, so a 1:2500 dilution of SA-HRP was used for these experiments. Complex samples were assessed in technical triplicate for each biological replicate (N=5: hSF, hCSF, bSF; N=4: human serum, RCM, BHI). The optimal matrix dilution identified by sar-sELISA was used in sELISA of complex samples. For example, a sample of 3200 ng/mL apo-RoxP_1-tagless ^a^ in hSF would be diluted 8-fold into ELISA-BB to make an initial sample of 400 ng/mL apo-RoxP_1-tagless in 12.5% (v/v) hSF and 87.5% (v/v) ELISA-BB. Then, that sample would undergo a 2-fold dilution series into ELISA-BB to create samples with concentrations of 200 ng/mL apo-RoxP_1-tagless in 6.25% (v/v) hSF and 93.75% (v/v) ELISA-BB, 100 ng/mL apo-RoxP_1-tagless in 3.125% (v/v) hSF and 96.875% (v/v) ELISA-BB), and so on.

Then, an apo-RoxP_1-tagless dilution series in ELISA-BB (e.g., 12.5-400 ng/mL ^b^) would be used to determine each sELISA plate’s LOD (defined as ≥3-fold the SD of the blank above the blank value) and LLOQ (defined as ≥10-fold the SD of the blank above the blank value). The RoxP standard curve was also used to calculate the empirical concentration of RoxP in all complex samples. Theoretical recovery (%) was then calculated by dividing the empirical concentration by the concentration that would be expected from 400 ng/mL RoxP (in appropriately diluted complex sample) that had undergone a 2-fold dilution series. Then, empirical recovery (%) was calculated by applying the relevant dilution factor (2- to 64-fold) to the highest empirical concentration and dividing those values by the concentration expected using the experimental design (e.g., 200-6.25 ng/mL). The comparison of theoretical to empirical recovery was used to address an ∼20% deficit in expected RoxP concentration (**SI Fig12g**,**h**,**j**), which tracked with the plate suggested that standard curve’s initial concentration was ∼80% of expected.

Percentage recovery was used to define the limit of dilution for this assay. This assessment showed that the LOD/LLOQ in ELISA-BB was essentially the same as the limit of dilution observed in complex samples. While this work was performed with SA-HRP, the expectation is that the SA-Poly-HRP would enhance the signal of sELISA in complex samples equivalently (e.g., >30-fold increase in LOD/LLOQ and limit of dilution).

**^a^ RoxP_1-tagless in sELISA experiments was prepared using the “Autoinduction** è ***Gravity-Flow IMAC Protocol”* and then diluted from >30 mg/mL (HSB) into PBST/biofluid.**

**^b^ This range was adjusted based on the goals of an experiment.**

### Biofluids

Human serum (MP Biomedicals: Cat # 092931149, Lot # S8657), human cerebrospinal fluid (Lee Biosolutions: Cat # 991-19-P, Lot # 09H4473), human synovial fluid (Innovative Research: Cat # IRHUSYNS1ML, Lot # HMN930539; Medix Biochemica: Cat # 991-42-P-1, Lot # 01J5473-14), and bovine synovial fluid (Innovative Research: Cat # IGBOSYNZ25ML, Lot # 41360) were purchased from commercial vendors. Upon receipt, biofluids were thawed (if necessary), 1.0 mL aliquots were prepared, all aliquots were flash-frozen in liquid nitrogen, initially stored at -80°C, and then stored prior to use in experiments at -20°C. Aliquots used in experiments for this work were thawed immediately prior to an experiment, and no aliquot was thawed more than twice for these studies. Aliquots that were refrozen between experiments (e.g., hSF and bSF for sar-sELISA) were frozen at -20°C.

## Funding sources

McCoy: NIH, United States grants: KL2TR002346, UL1TR002345, 5K08AR076464 Aleem: Private Donation

## Conflicts of Interest

Declaration of interest: None Conflicts of interest: None

## IRB approval status

Not applicable

## Consent

Not applicable

## Classification(s)

Original research

## Table of Manuscript Abbreviations

Abbreviation: Full Name
8His: 8 histidine tag (for protein purification)
αROS: anti-reactive oxygen species
A280,450,570,650: absorbance at wavelengths 280/450/570/650 nm
AU: absorbance unit
AUC: analytical ultracentrifugation
BHI: Brain Heart Infusion
BPB: bromophenol blue
BSA: bovine serum albumin
bSF: bovine synovial fluid
CAMP: Christie–Atkins–Munch-Peterson
CBB: Coomassie brilliant blue
CDR: complementarity-determining region
coNS: coagulase-negative staphylococci
Cons.: conservative
CPIII: coproporphyrin III
CSF: cerebrospinal fluid
Cterm: carboxyl(C)-terminus
Dil: dilution
DSF: differential scanning fluorimetry
ELISA: enzyme-linked immunosorbent assay
Fc: fragment crystallizable region
FFA: free fatty acids
FP: forward polarity
FT: flow-through
Fv: fragment variable region
HA: hemin-agarose
hc: heavy-chain
hCSF: human CSF
hCSF: human cerebrospinal fluid
HD: hemin dissolved in DMSO
Hmp: hemopexin
HN: hemin dissolved in 1.4 N NaOH
HRP: horseradish peroxidase
HSB: HEPES sizing buffer
hSerum: human serum
hSF: human synovial fluid
iELISA: indirect ELISA
IHC: immunohistochemistry
IMAC: immobilized-metal affinity chromatography
IMD: Indwelling medical device
Infxn: infection
IPG: Identical Protein Group
kDa: kilodalton
lc: light-chain
LLOQ: lower limit of quantitation
LOD: limit of detection
mAb: monoclonal antibody
mAU: milli-absorbance unit
mAU: milli-absorbance unit
MSA: multiple sequence alignment
MW: molecular weight
N: no
N=: number of replicates
NCBI: National Center for Biotechnology Information
NCP: negative control protein
nm: nanometer
NMR: nuclear magnetic resonance
Nterm: amino(N)-terminus
OD600: optical density at 600 nm
ODS: o-Dianisidine (3,3′-dimethoxybenzidine)
ORG: open-reading frame
pAB: polyclonal antibody
PAGE: polyacrylamide gel electrophoresis
PBS: phosphate-buffered saline
pI: isoelectric point
PJI: periprosthetic joint infection
POC: point-of-care
PPIX: protoporphyrin IX
PSU: pilosebaceous unit
RB: running buffer
RCM: reinforced Clostridial medium
RefSeq: Reference Sequence
RP: reverse polarity
rpm: revolutions per minute
RT: room temperature
SA: streptavidin
SA-HRP: streptavidin-horseradish peroxidase
SA-Poly-HRP: SA conjugated with polymers of HRP
sar-sELISA: spike-and-recovery sELISA
SASA: solvent-accessible surface area
SCN: siderocalin
SD: standard deviation
SDS: sodium dodecyl sulfate
SEC: size-exclusion chromatography
sELISA: sandwich ELISA
SI: supplementary information
SLST: single locus sequence typing
TCE: 2,2,2-trichloroethanol
TEV: Tobacco Etch Virus
Tma: apparent melting temperature
Trp: tryptophan
Tyr: tyrosine
W: tryptophan
Y: yes

## Supplementary Information (SI) Figure Legends

### Bibliography – SI Figure Legends

**SI Figure 1.**
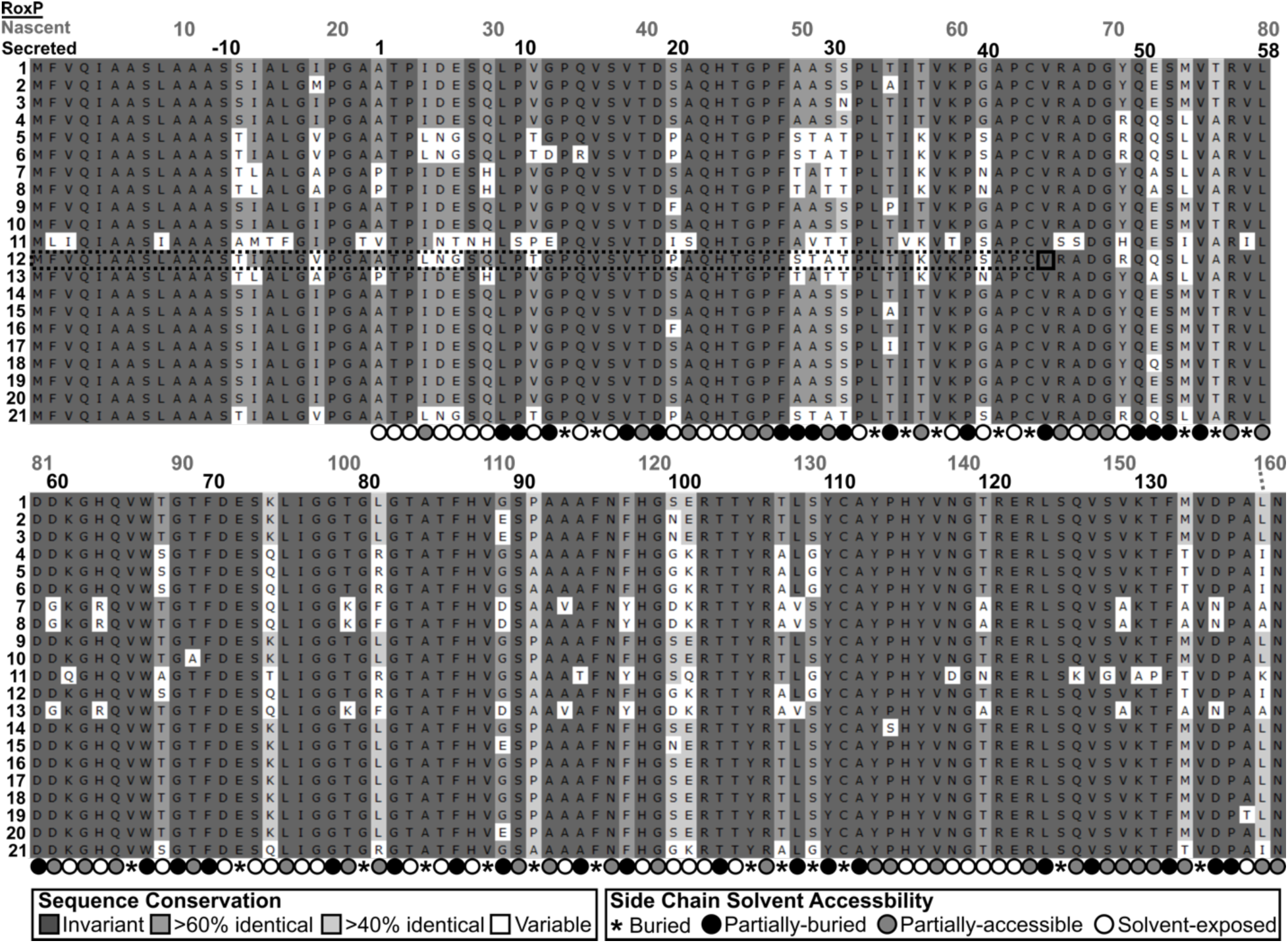
Multiple Sequence Alignment of All RoxP Orthologs. Multiple sequence alignment (MSA) of the 21 RoxP orthologs is shown. Residue numbering of the *<u>nascent</u>* polypeptide chain and predicted, mature, *<u>secreted</u>* RoxP is shown. RoxP signal peptide cleavage was predicted by SignalP-4.1^1^. All residue positions noted in the text of this paper refer to mature, *<u>secreted</u>* RoxP (i.e., post-signal peptide cleavage) numbering. Residue backgrounds are colored according to sequence identity and variable positions are highlighted in white. The surface-exposure of each residue’s side chain in the apo-RoxP NMR structure (PDB 7bcj)^2^ as calculated by GetArea^3^ is shown below the alignment as a percentage exposure compared to the same residue in a tripeptide (Gly-X-Gly) random coil. Variant positions within predicted, mature, *<u>secreted</u>* RoxP include: 38 total non-conservative variant positions (27%), 31 variant positions with exclusively non-conservative variants (22%), 27 total conservative/semi-conservative variant positions (19%), 20 variant positions with exclusively conservative/semi-conservative variants (14%), and 7 variant positions (23, 33, 53, 56, 103, 142, 160) with both non-conservative and conservative/semi-conservative variants (5%). Solvent-accessibility categories: Buried (≤1%), Partially-buried (>1% to <30%), Partially-accessible (30-60%), Solvent-exposed (>60%). The first digit of a residue position was aligned to the corresponding MSA column e.g., 1 of **<u>1</u>**0 is centered on the corresponding MSA column for position 10. RoxP_12 was initially identified as a truncated (96 instead of 161 amino acid) RoxP ortholog in *C. modestum* strain P08 (AFAM01000017.1: 304958-305248 *Cutibacterium modestum* P08 contig00017), but a later reassessment of that genome’s contig (AFAM01000017.1:304763-305248) revealed that the complete open-reading frame (ORF) for RoxP_5 is likely present. The annotation mistake was due to the depositors of this genome^4^ accidentally using the GTG at AFAM01000017.1: 304958-304960 as a methionine start codon, when it would be a valine within the full-length sequence. The full-length, 161 amino acid sequence for both RoxP_5 and RoxP_12 are included in this alignment, and the RoxP_12 sequence is labeled with the *in silico,* “truncated” N-terminal sequence (**black**-dashed box) and (likely) mistakenly used GTG start codon (**black** box). RoxP_7 and RoxP_13 are also identical in the 161 amino acids shown in this alignment, but RoxP_13 has an additional seven N-terminal amino acids (MTYWGIT) making it a distinct RoxP ortholog. With or without the inclusion of RoxP_13 (MTYWGIT) leads to the same signal peptide cleavage prediction (ALG-APG) using SignalP-6.0^5^, but without those seven amino acids, SignalP-4.1^6^ predicts the loss of three N-terminal amino acids with a cleavage site of APG-APT.

**SI Figure 2.**
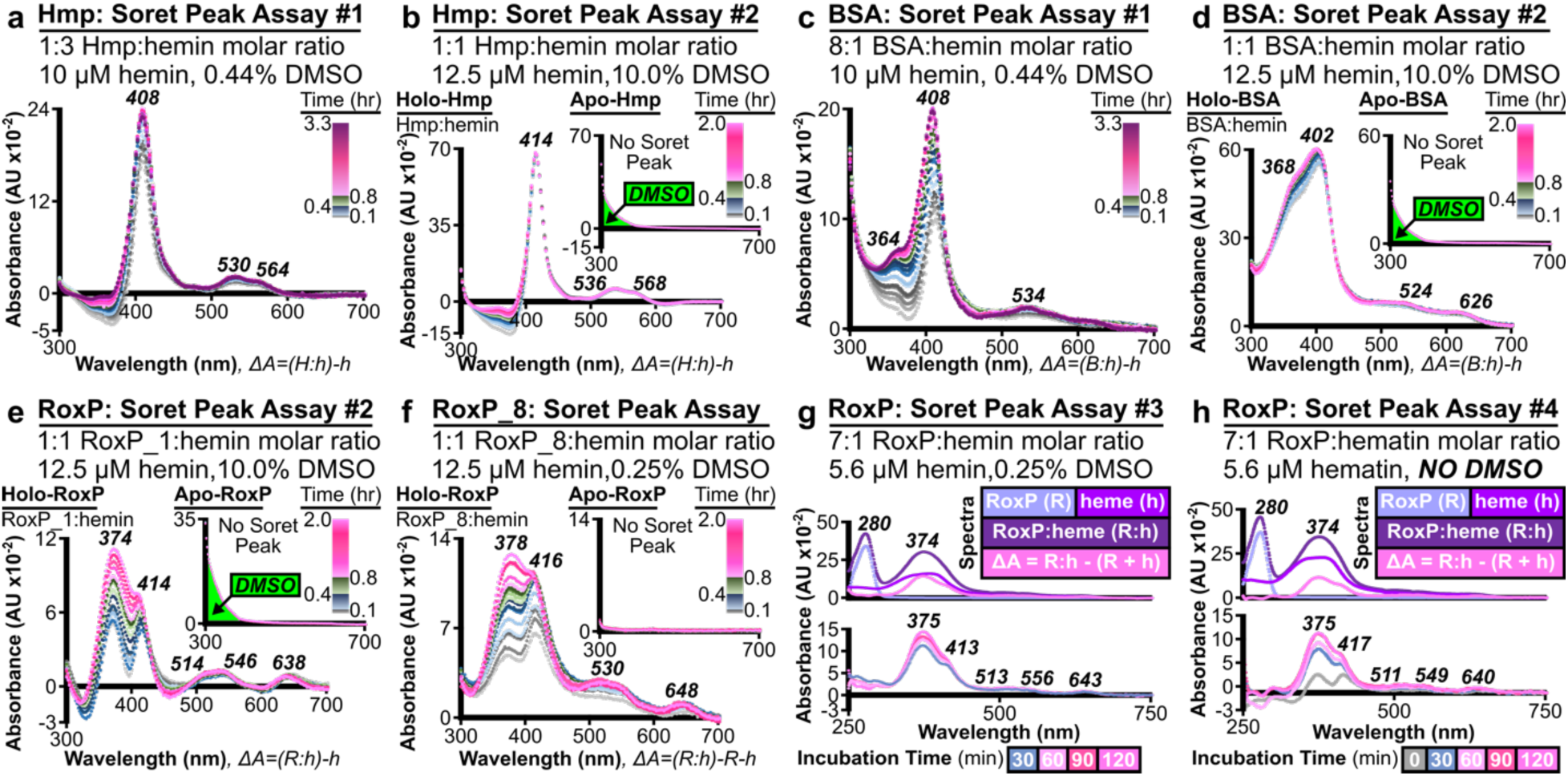
Soret Peak Analysis Controls and Extension to RoxP_8. The effect of DMSO concentration, protein:hemin molar ratio, hemin vs. hematin as a ligand, and RoxP ortholog were assessed. (**a**-**d**) Two heme-binding control proteins (high-affinity: hemopexin [Hmp]; low-affinity: bovine serum albumin [BSA]) were assessed at (**a**,**c**) low DMSO (0.44%) and (**b**,**d**) high DMSO (10%) concentrations. These heme-binding control proteins were also assessed at protein:hemin ratios with (**a**) excess hemin, (**c**) excess protein, and (**b**,**d**) equimolar protein/hemin. (**e**) RoxP_1:hemin Soret peak analysis was also performed using the high DMSO (10%) concentration and equimolar protein:hemin. (**e**,**f**) Time-dependent increase in the two RoxP Soret peaks (370-378, 412-416 nm) were observed for orthologs (Fig 4b, **SI Fig 4e**) RoxP_1 and (**SI Fig 4f**) RoxP_8. (**g**,**h**) The Soret peak microplate assay (Fig 4b, **SI Fig 4a-f**) was adapted to cuvettes, and this protocol was used to demonstrate equivalent RoxP_1 Soret peaks for (**g**) hemin (375, 413 nm) and (**h**) hematin (375, 417 nm), which both reached equilibrium by 2 hours (see time course legend in the bottom-right of each panel). In the top panel, the major absorbance peak for protein (280 nm) and heme (374 nm) are labeled in ***bold italics***. (**a**-**h**) Soret peaks and Q-bands are shown in ***bold italics***. (**a**-**f**) The lowest absorbance spectrum is the time 0 scan. Incubation time prior to each scan is shown in the legend (top-right) as a color gradient. Color gradient heights were scaled to the time span examined. (**a**,**c**) From bottom-to-top, the color gradient and number of included traces for each incubation time span are grey-scale (0-240 s, 5 traces), blue (300-1600 s, 5 traces), green (1560-2760 s, 3 traces), and magenta (2820-11819 s, 6 traces). (**b**,**d**,**e**,**f**) From bottom-to-top, the color gradient and number of included traces for each incubation time span are grey-scale (0-240 s, 5 traces), blue (300-1500, 5 traces), green (1800-2700, 4 traces), and magenta (3000—7140 s, 5 traces). (**b**,**d**,**e**) Inset protein-only plots without buffer subtraction show the absorbance (**bottom-left corner**) from DMSO in the samples, which is not seen in the (Fig 4b) blank-subtracted RoxP alone spectrum. All other spectra are blank-subtracted, difference spectra. See x-axis label for data processing e.g., ΔA=(R:h)-h is the difference spectrum (ΔA) from the RoxP:hemin (R:h) spectrum minus the hemin (h) alone spectrum. (**e**) Inset y-axis was increased compared to difference spectrum to capture the DMSO absorbance at 300 nm. (**g**,**h**) Though all plots were generated in Excel with the same axes/formatting for each plot type (all spectra vs. time course of difference spectra) and exported as vector files (.svg), the y-axis height of (**g**) was compressed compared to (**h**) upon importing into Inkscape.

**SI Figure 3.**
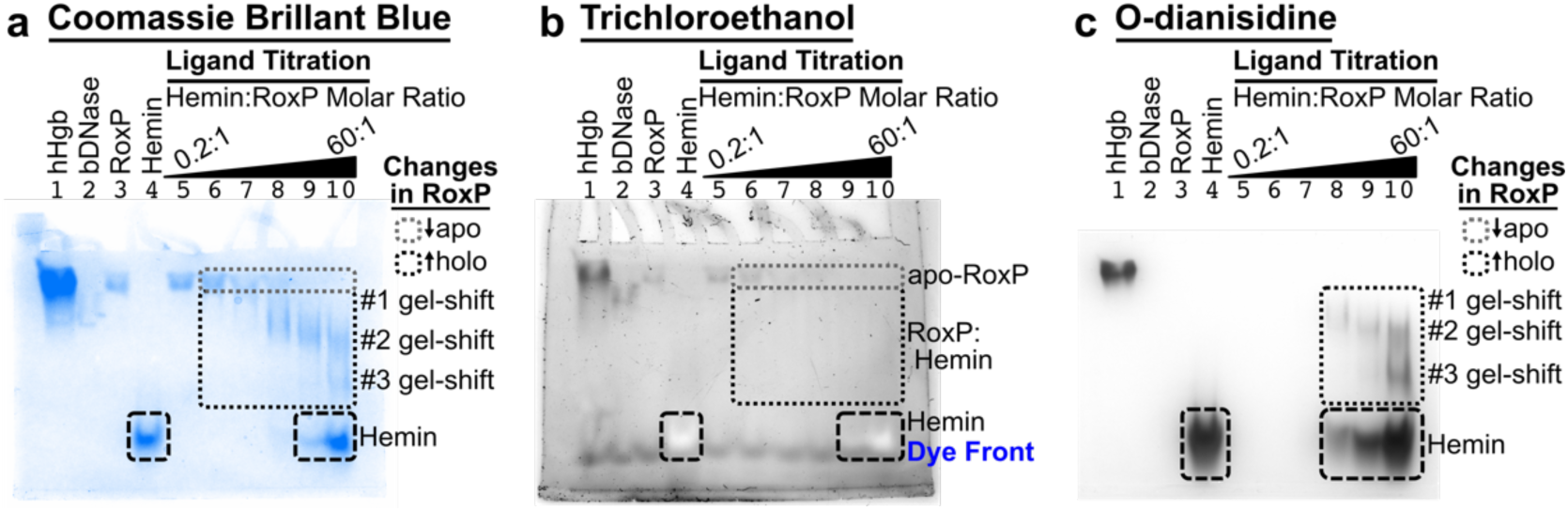
Full Images of RoxP:Hemin NATIVE Gels. This figure presents the full gel images of NATIVE gels shown in Fig 4c, so that control lanes 1-4 may be assessed by the reader. RoxP:hemin binding reactions were incubated (aerobic conditions, 0.5 hr RT), loaded into two 12% NATIVE gels run in the same gel box (pH 8.3 running buffer [RB]), and then imaged using three staining techniques: (**a**) Coomassie brilliant blue (CBB) that stains proteins, (**b**) 2,2,2-trichloroethanol (TCE) that is activated by UV light and reacts with tryptophans (exposed tryptophan stain), and (**c**) O-dianisidine (ODS) stain that reacts with heme (heme stain). Gel #1 was (**b**) UV-imaged (TCE/W stain) and then (**a**) CBB stained, while gel #2 only underwent (**c**) ODS/heme-staining. (**a**,**c**) CBB and ODS demonstrated three heme-dependent gel-shifts for holo-RoxP_1. (**a**) CBB staining of RoxP:hemin binding reactions demonstrated titratable, hemin-dependent (i) decrease in apo-RoxP_1 (**grey**-dashed box) and an (ii) increase in holo-RoxP_1 (**black**-dashed box) with three distinct gel-shift bands. CBB staining also showed the migration location of hemin alone (**black**-long-dashed box). (**b**) TCE staining of RoxP:hemin binding reactions demonstrated titratable, hemin-dependent (i) decrease in apo-RoxP_1 (**grey**-dashed box) and (ii) a barely discernable increase in holo-RoxP_1 (**black**-dashed box), though no specific gel-shift bands could be identified. Since TCE staining of holo-RoxP was much less than would have been expected based on (**a**) CBB staining (compare lane 5/6 apo-RoxP intensity with lane 10 holo-RoxP intensity), it suggested that heme-binding protects RoxP_1 W66 from TCE modification. TCE staining also showed the migration location of (i) hemin alone (**black**-long-dashed box, white gel band) and (ii) the bromophenol blue (BPB) dye front (labeled in **blue**). (**c**) ODS staining of RoxP:hemin binding reactions demonstrated titratable, hemin-dependent increase in heme-bound, holo-RoxP_1 (**black**-dashed box) with three distinct gel-shift bands. ODS staining clearly showed the migration location of hemin alone (**black**-long-dashed box). ODS staining specificity for heme-containing proteins was also demonstrated using a positive control (hHgb: human hemoglobin) and a negative control (bDNase: bovine pancreatic deoxyribonuclease). In the original experimental plan, all protein lanes were to be loaded with 4.2 µg of protein, but an error led to lane 1 containing 38 µg of hHgb. All other protein lanes (2-3 and 6-10) contained the planned 4.2 µg of their respective proteins.

**SI Figure 4.**
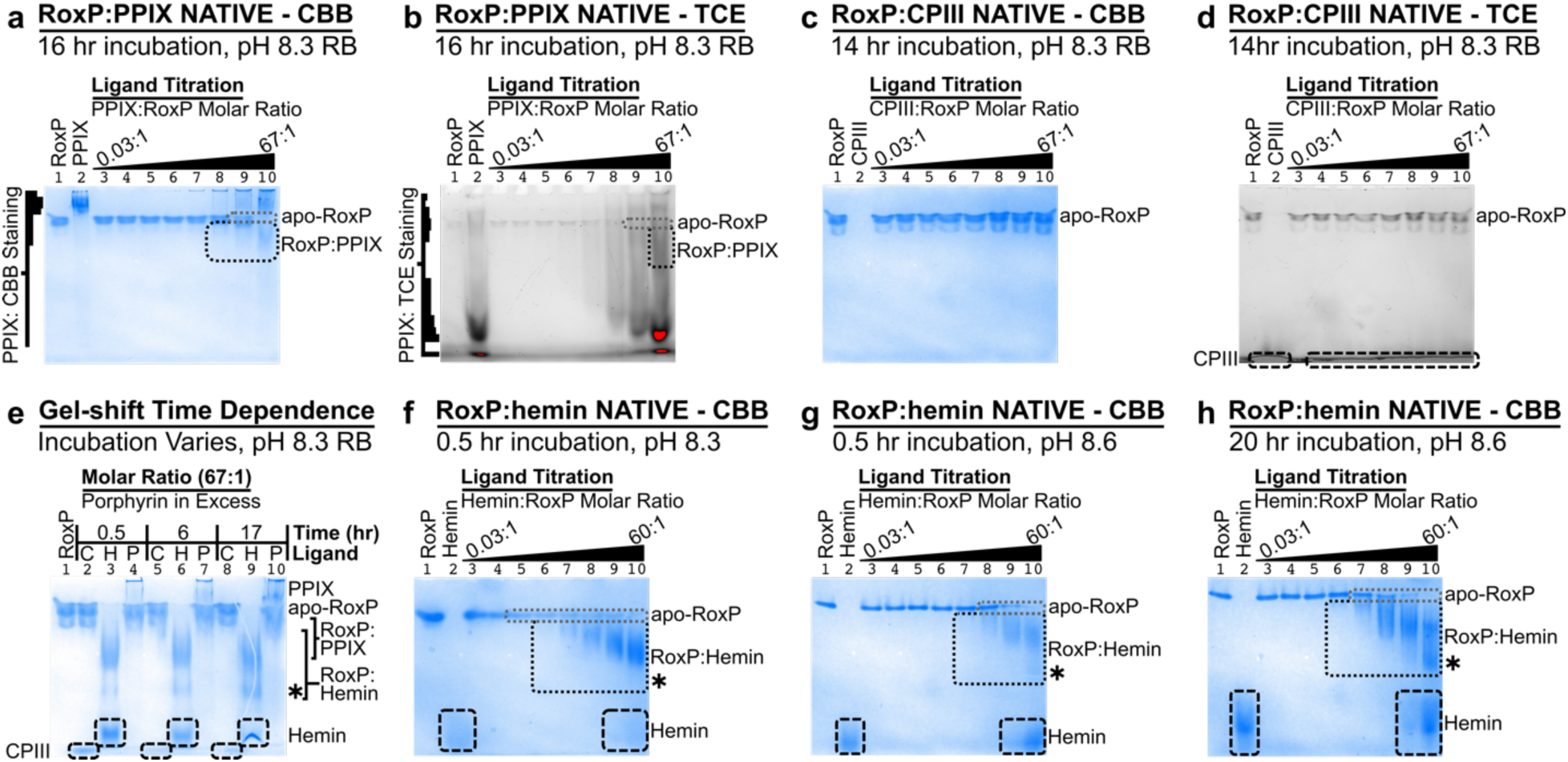
NATIVE Gel Assessment of RoxP:Porphyrin Binding. Effect of varying electrophoresis conditions (time, pH) on RoxP:porphyrin NATIVE gel migration was evaluated using CBB and TCE staining. NATIVE RB pH was varied from standard pH of 8.3 to 8.6. Individual experiments are listed by the date (YYYYMMDD) that they were performed. RoxP in this figure refers to RoxP_1-tagless. (**a**,**b**) 20190516 RoxP:PPIX binding experiment performed under aerobic conditions at pH 8.3 after 16 hr RT incubation that demonstrates titratable, PPIX-dependent (i) decrease in apo-RoxP_1 (**grey**-dashed box) and an (ii) increase in holo_RoxP_1 (**black**-dashed box). The migration location of PPIX alone (lane 2) as demonstrated by (**a**) CBB and (**b**) TCE is shown, which demonstrated that the predominant stainable PPIX species is (**a**) slower or (**b**) faster migrating depending on the gel staining technique. Qualitative staining intensity is also indicated for lane 2 PPIX to the left of each gel image to help distinguish it from the changes in apo-/holo-RoxP species indicated to the right of the gel. (**b**) Saturated pixels are shown in **red**. (**c**,**d**) 20190612 RoxP:CPIII binding experiment performed under aerobic conditions at pH 8.3 after 14 hr RT incubation did not produce a NATIVE gel-shift for RoxP:CPIII. The migration location of CPIII alone is (**c**) not visible by CBB due to destaining but (**d**) is visible by TCE (**black**-long-dashed box), which was performed immediately after running the NATIVE gel. (**e**) 20190612 RoxP:porphyrin binding experiment performed under aerobic conditions at pH 8.3 with a 67-fold molar excess of porphyrin as a function of incubation time at RT demonstrates (i) an increase in RoxP:hemin migration with extended incubation time, (ii) increase in the faster migrating RoxP:hemin(H) band (*) with extended incubation time, (iii) saturated RoxP:PPIX(P) migration across all time points, and (iv) no gel-shift (i.e., evidence of binding) for CPIII(C) to apo-RoxP_1-tagless. The migration location of CPIII as demonstrated by CBB is also shown (**black**-long-dashed box), though its location was rarely captured by CBB staining in gel images due to rapid destaining of this band prior to gel image capture. The migration location of hemin alone as demonstrated by CBB is also shown (**black**-long-dashed box). (**f**) 20190614 RoxP:hemin binding experiment performed under aerobic conditions at pH 8.3 after 0.5 hr RT incubation demonstrates titratable, hemin-dependent (i) decrease in apo-RoxP_1 (**grey**-dashed box) and an (ii) increase in holo_RoxP_1 (**black**-dashed box). The migration location of hemin alone as demonstrated by CBB is also shown (**black**-long-dashed box). (**g**) 20190603 RoxP:hemin binding experiment performed under aerobic conditions at pH 8.6 after a 0.5 hr RT incubation demonstrates titratable, hemin-dependent (i) decrease in apo-RoxP_1 (**grey**-dashed box) and an (ii) increase in holo_RoxP_1 (**black**-dashed box). The migration location of hemin alone as demonstrated by CBB is also shown (**black**-long-dashed box). (**f**,**g**) Comparison of these identical hemin-titrations (see lanes 5-10) suggests higher affinity binding at pH 8.3 compared to pH 8.6 due to the (i) increased holo-RoxP migration (**f** – lane 6 vs. **g** – lane 7) and (ii) decreased in apo-RoxP (**f** – lane 5 vs. **g** – lane 8). (**h**) 20190603 RoxP:hemin binding experiment performed under aerobic conditions at pH 8.6 after 20 hr RT incubation demonstrates titratable, hemin-dependent (i) decrease in apo-RoxP_1 (**grey**-dashed box), (ii) increase in holo_RoxP_1 (**black**-dashed box) that appears at lower [hemin] (lane 6) compared to (**g** – lane 7), and (iii) increase in the faster migrating holo-RoxP:hemin band (* in **e**-**h**) with extended incubation time compare to (**g**). The migration location of hemin alone as demonstrated by CBB is also shown (**black**-long-dashed box).

**SI Figure 5.**
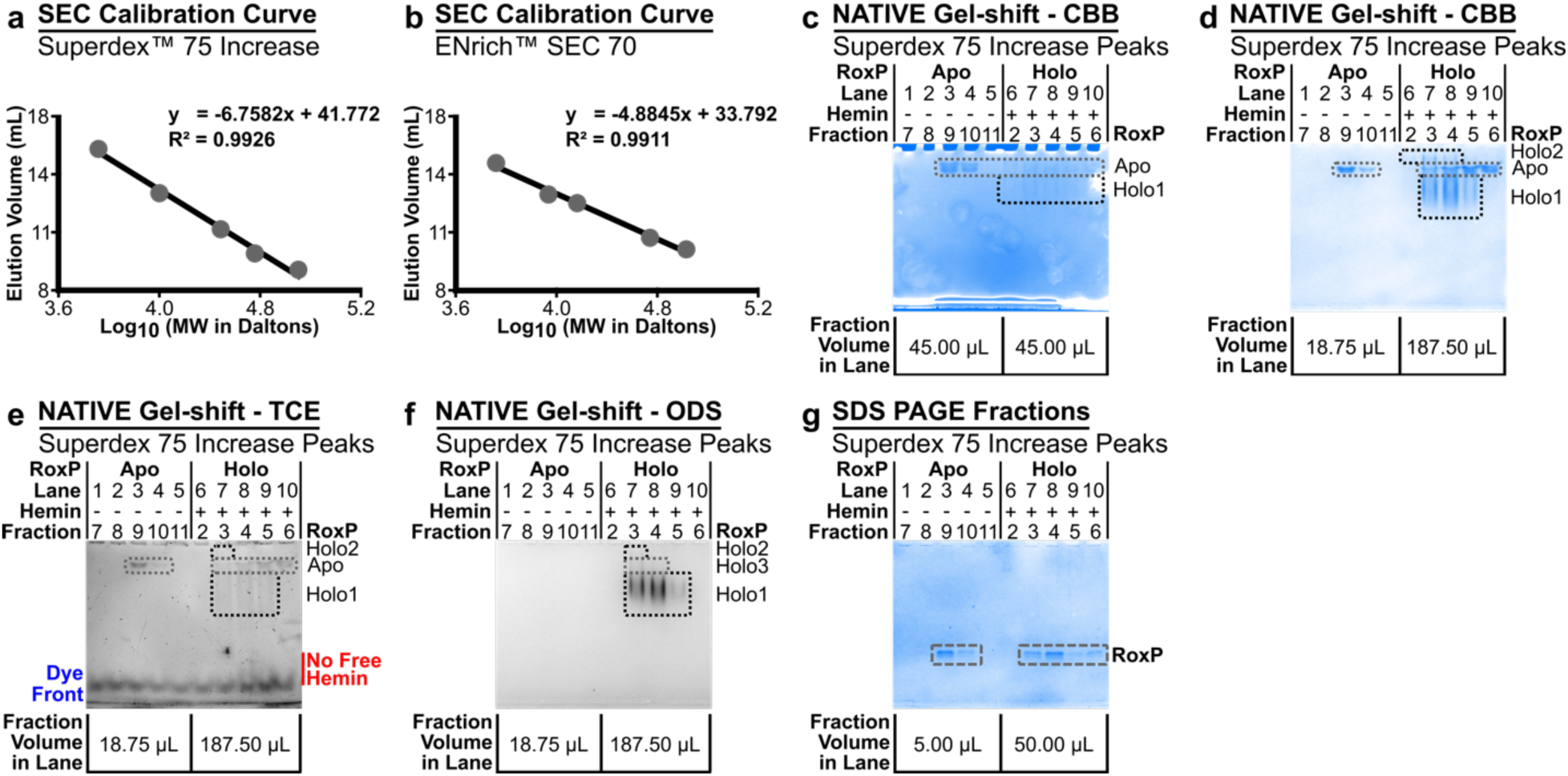
SEC Calibration Curves and Complete Gels of SEC Peaks. (**a**,**b**) SEC calibration curves are shown as elution volume (V_e_) vs. log MW to aid in reader interpretation and use of the linear fits presented in each panel. SEC MW calibration was calculated using the equation K_av_=(V_e_-V_o_)/(V_c_-V_o_) with values V_e_ (elution volume for each MW standard), V_o_ (void volume, blue dextran), V_c_ (column volume). (**a**) Superdex^TM^ 75 Increase calibrated using MW standards from GE Healthcare LMW (low molecular weight) kit: ribonuclease A, carbonic anhydrase, ovalbumin, conalbumin. Blue Dextran was not available during this column’s calibration, but it’s absence does not impact the accuracy of MW estimation via a linear fit of V_e_ vs. MW, as V_o_ and V_c_ are constants in the equation K_av_=(V_e_-V_o_)/(V_c_-V_o_). This calibration was used in Fig 4e. (**b**) ENrich^TM^ SEC 70 calibrated using MW standards from GOLDBIO (aprotinin) and Sigma (aprotinin, cytochrome C, myoglobin, ovalbumin, BSA, Blue Dextran). This calibration was used to prepare AUC samples (Fig 4f) and to evaluate RoxP orthologs (Fig 4h). **(c**-**g)** This figure shows the (**d**,**g**) full gel images of NATIVE and SDS PAGE gel lanes shown in Fig 4e, as well as (**c**,**e**,**f**) other NATIVE gels run on SEC fractions of RoxP_1 (+/-) hemin run over a Superdex^TM^ 75 Increase column. The ranges of the 280 nm absorbance (A_280_) traces examined by these gels are shown in Fig 4e as black-outlined bars with the same color as the A_280_ trace (**pale-blue: apo-RoxP**; **magenta: holo-RoxP**). (**c**) SEC fractions were initially evaluated on a 12% NATIVE gel stained with CBB, which demonstrated decreased apo-RoxP_1 (**grey**-dashed box) in the (+) hemin samples that coincided with the appearance of holo-RoxP_1 (**black**-dashed box). Due to the appearance of several potential holo-RoxP NATIVE species not observed in the NATIVE gel shift assays (**d**-**f**: Holo2; **f**: Holo3), holo-RoxP in (**c**) is labeled as “Holo1” in this figure, and (**c**-**f**) use the same **grey**-/**black**-dashed boxes. (**d**-**f**) To eliminate the apo-RoxP doublet caused by protein overloading (**c**) lanes 3-4 and better evaluate holo-RoxP in (**c**) lanes 6-10, the protein sample load (fraction volume loaded per lane) was decreased and increased, respectively (compare bottom of panels **c**-**f**, see **SI Methods**). (**d**) CBB staining of SEC fractions identified both holo-RoxP as seen in NATIVE gel shift assays (Holo1), as well as a slower migrating band (lanes 6-8, Holo2) above the apo-RoxP bands. (**e**) TCE staining shows decreased apo-RoxP staining in lanes 7-8 that correspond to larger MWs by SEC, but it also demonstrated TCE(+) bands in the regions below (Holo1) and above (Holo2) the apo-RoxP bands. (**f**) ODS stain that reacts with heme (heme stain) clearly marks the holo-RoxP bands in lanes 7-9 (Holo1), but it also partially stained apo-RoxP bands (lanes 7-8) and the slower migrating band (Holo2) in lane 7. (**d**-**f**) The slower migrating bands unique to the (+)hemin sample (Holo2) correspond to fractions with MWs of 60-36 kDa (lane 6 and 8, respectively), and we hypothesize that they represent the holo-RoxP pentamer/trimer/dimer equilibrium observed by AUC (Fig 4f). (**d**-**f**) The unexpected TCE/ODS staining of (**e**) holo-(Holo1/2) and (**f**) apo-RoxP (Holo3), respectively, is hypothesized to be due to partial dissociation of holo-RoxP during SEC/PAGE. Dissociation of holo-RoxP would be expected to be more likely to occur after SEC removal of the 10-fold molar excess of hemin. Thus, (**c**-**f**) lanes 7-8 are most comparable in the NATIVE gel shift assay experiments to the molar ratio of hemin:RoxP in **SI Fig 4f** (lane 6, 0.74:1 ratio). Further, (**f**) holo-RoxP dissociation from an oligomeric state might lead to monomeric holo-RoxP (Holo3) that migrates at the same relative position as apo-RoxP. (**e**) While TCE can inversely stain hemin (white band: see **SI Fig 3b** lanes 4, 9, 10), no free hemin was observed in these SEC fractions (see expected migration location of hemin labeled in **red**). TCE did stain the BPB dye front (labeled in **blue**). (**g**) SDS PAGE was used to examine the apo- and holo-RoxP SEC fractions to compare the amount of total RoxP present in each fraction (**grey**-long-dashed box). Though lanes 3 and 8 appear to have a similar amount of CBB-stained RoxP, there is 10-fold more fraction volume in lane 8 compared to lane 3. The lane 8 peak height is also ∼2-fold larger than the lane 3 peak, which leads to the approximation that 96% of the holo-RoxP A_280_ peak height is due to hemin absorbance, rather than RoxP protein absorbance.

**SI Figure 6.**
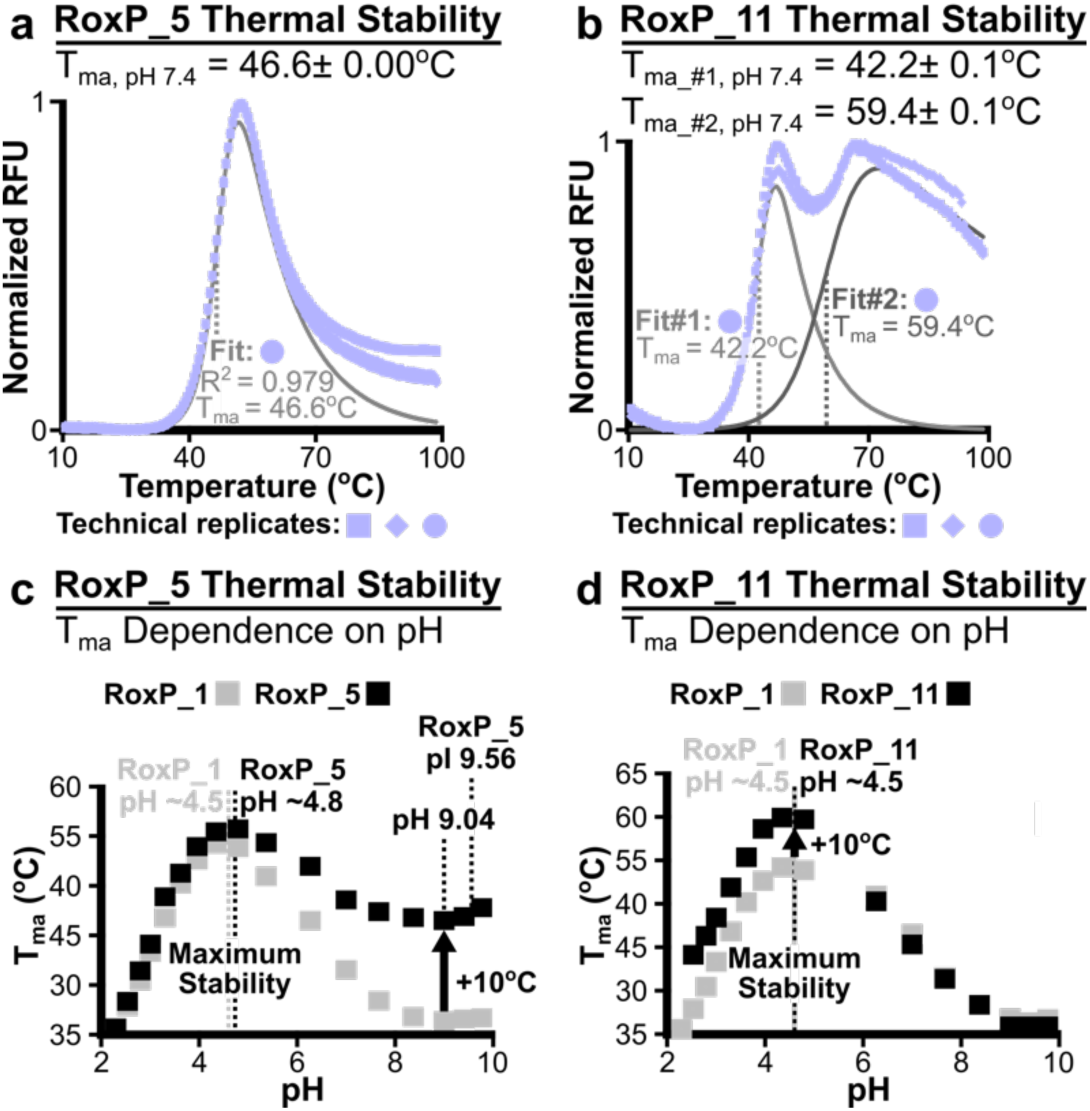
Differential Scanning Fluorimetry (DSF) pH Dependence of RoxP Orthologs. The effect of pH on the RoxP_1 (100% identity, pI 6.19) melting temperature apparent (T_ma_) shown in Fig 3e**,f** led to the evaluation of how pH affected T_ma_ for two RoxP orthologs: (**a**,**c**) RoxP_5 (82%, pI 9.56), (**b**,**d**) RoxP_11 (71%, pI 6.38). All proteins were apo-RoxP-tagless protein preparations. At pH 7.4, (**a**) RoxP_5 was found to have significantly higher thermal stability (T_ma_ 46.6±0.0°C, DSFworld model #2) than RoxP_1 (T_ma_ 39.1±0.2°C, DSFworld model #2), while (**b**) RoxP_11 had higher thermal stability and appeared to undergo two transitions (DSFworld model #4, R^2^ 0.995) unfolding with T_ma_ values of 42.2±0.1°C and 59.4±0.1°C. Use of DSFworld model #2 to fit the RoxP_11 data resulted in a T_ma_ of ∼41°C with R^2^ <0.7 due to the fit not accounting for the 2^nd^ transition. (**c**,**d**) The effect of pH on T_ma_ for (**c**) RoxP_5 (N=3) and (**d**) RoxP_11 (N=3) was then assessed and compared to RoxP_1 (N=4), which showed that both orthologs appear most stable at low pH. Symbols represent average of technical replicates at each pH. (**c**) Unlike RoxP_1/11, RoxP_5 did not exhibit as substantial a decrease in T_ma_ beyond pH 5, which may be due to higher overall stability of this ortholog and/or its higher pI. (**d**) RoxP_11 also exhibited a low pH T_ma_ that was ∼10°C higher than RoxP_1’s T_ma_ at low pH, which may have been required during the adaptation of *C. porci* to the pig (*Sus domesticus*) gastrointestinal tract^7^, as pig’s body temperature (39.67±0.09°C^8^) is 2-3°C higher than humans.

**SI Figure 7.**
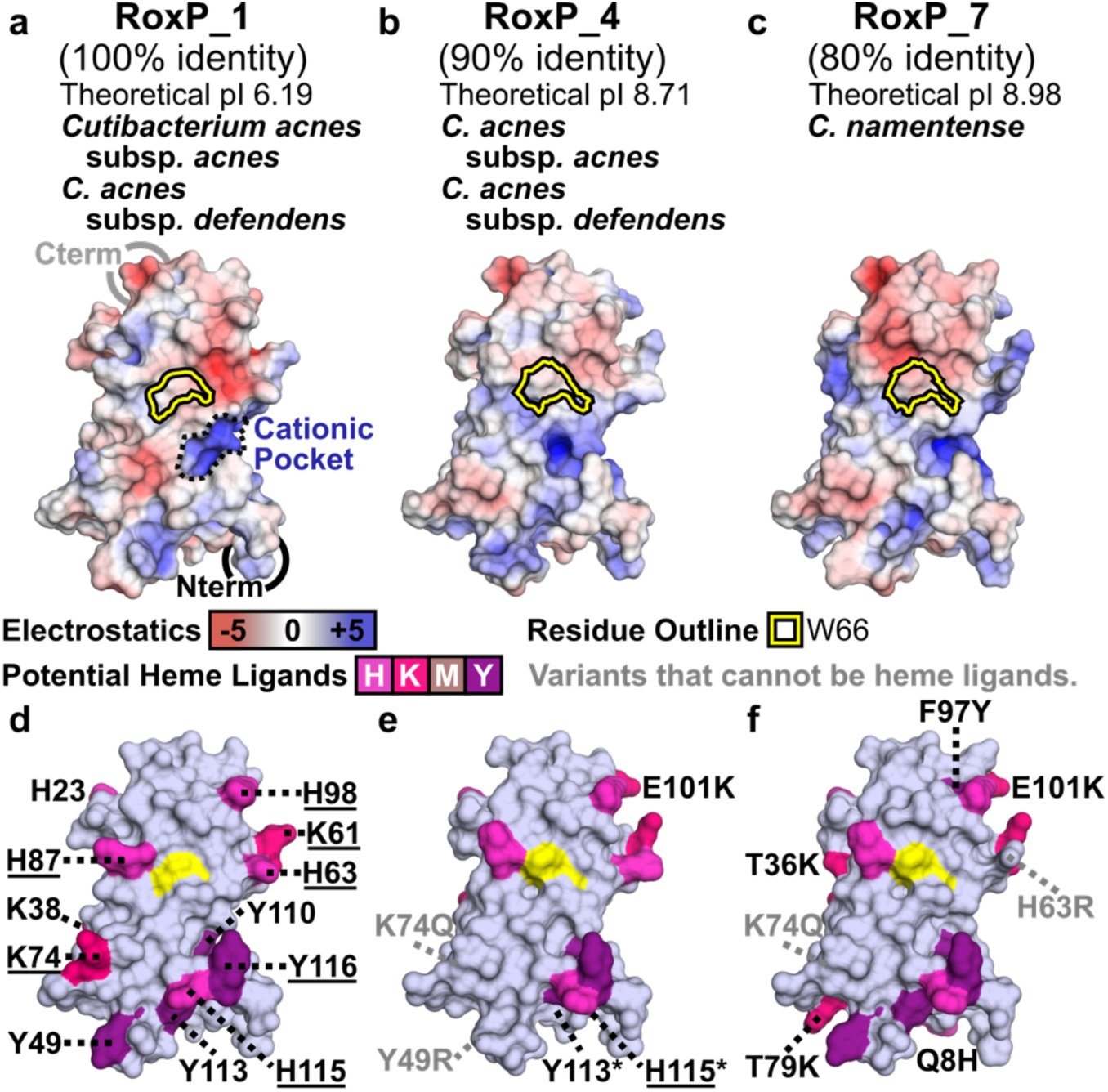
Molecular Modeling to Characterize RoxP_4/7 Heme-Binding Site. For each RoxP ortholog shown experimentally to bind heme (**SI Table 5**), sequence identity relative to RoxP_1, theoretical isoelectric point (pI), and *Cutibacterium* species/subspecies encoding the ortholog are shown. (**a**-**c**) Electrostatic surfaces and (**d**-**f**) potential heme ligands are shown for (**a**,**d**) the NMR structure of RoxP_1 (PDB 7bcj) and (**b**,**c**; **e**,**f**) three-dimensional models of two RoxP orthologs (RoxP_4, 7) shown to bind heme. The partially-buried tryptophan surface is (**a**-**c**) outlined or (**d**-**f**) colored yellow (see Fig 2) on each molecular surface, and (**a**) the **cationic pocket** is outlined by a **black**, dashed line. (**d**-**f**) Potential RoxP heme axial ligands are labeled in **black,** <u>underlined</u> if in close proximity to W66, and colored according to Fig 2e. (**e,f**) Ortholog positions lacking a potential heme axial ligand found in RoxP_1 are labeled in **grey**. (**e**) Two potential, invariant heme ligands (Y113, H115) have side chains that flipped during modeling, which makes it harder to identify them in comparison to panel (**d**). These residues are labeled with asterisks in Fig 5g, and equivalent side chain flips were observed for four RoxP orthologs (RoxP_4,5,6,11). See Fig 5 for the analysis of four additional heme-binding orthologs (RoxP_5,6,8,11).

**SI Figure 8.**
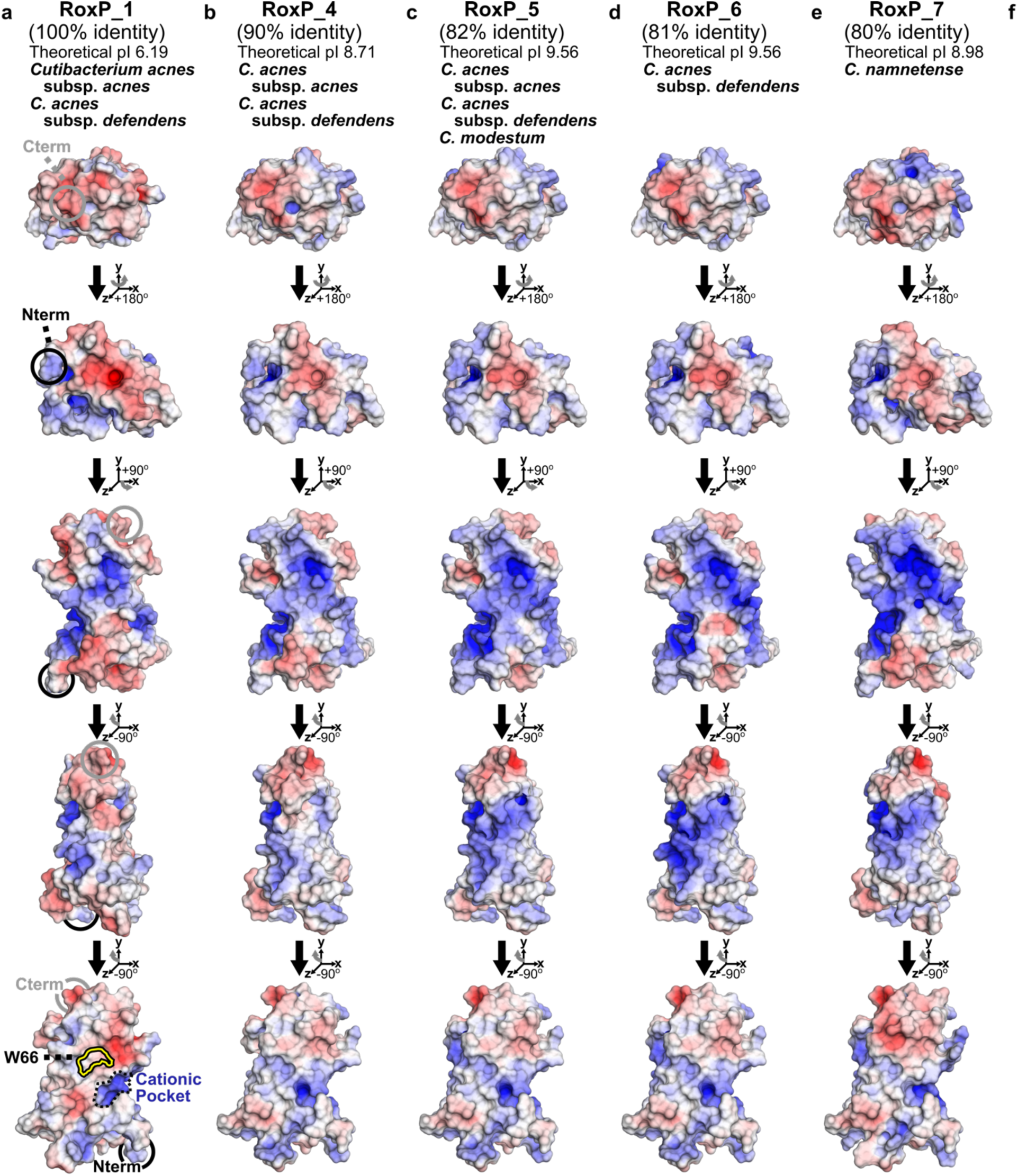
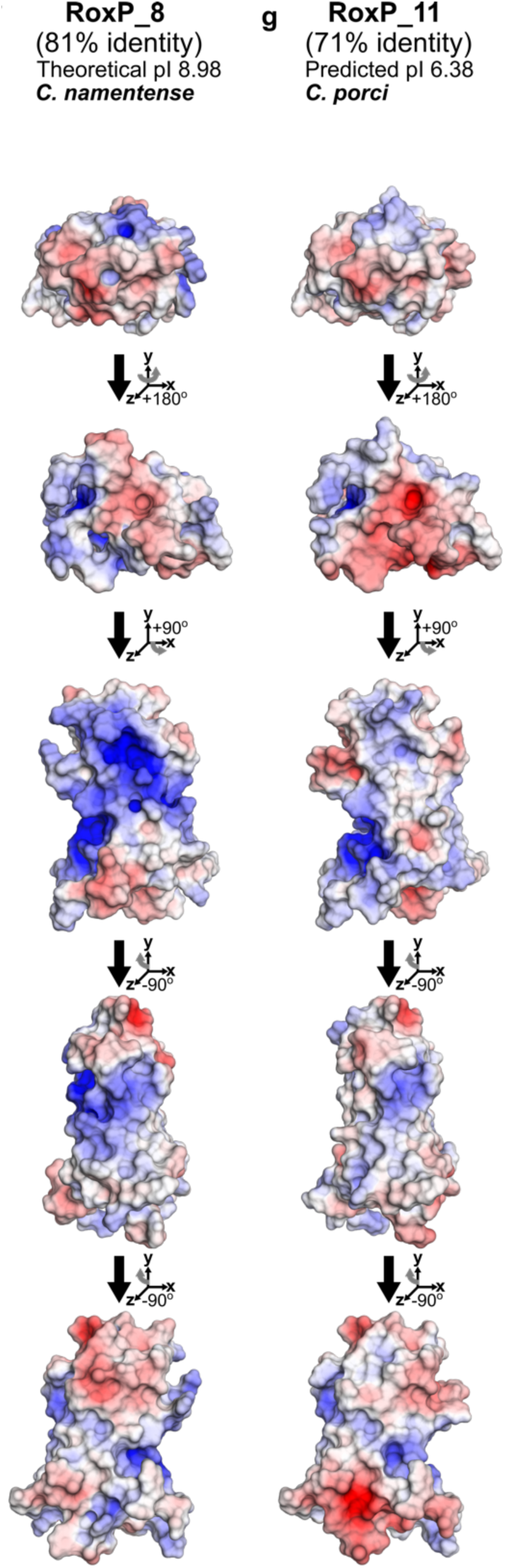
Electrostatic Surfaces of RoxP Orthologs Shown to Bind Heme. For each RoxP ortholog shown experimentally to bind heme (RoxP_1,4,5,6,7,8,11) (**SI Table 5**), sequence identity relative to RoxP_1, theoretical pI, and *Cutibacterium* species/subspecies encoding the ortholog are shown. (**a**-**g**) Electrostatic surfaces are shown for (**a**) the NMR structure of RoxP_1 (PDB 7bcj) and (**b**-**g**) three-dimensional models of six RoxP orthologs shown to bind heme. (**a**-**g**) The proposed heme-binding site (Fig 5) and (**a**) **cationic pocket** (outlined by a **black**, dashed line) are shown in the bottom row, while the opposite molecular surface is shown in the middle row. RoxP ortholog protein sequences used in this analysis were based on recombinant RoxP constructs after removal of the PelB signal peptide and N-terminal 8His-TEV tag (e.g., RoxP_5 GATP…PAIN).

**SI Figure 9.**
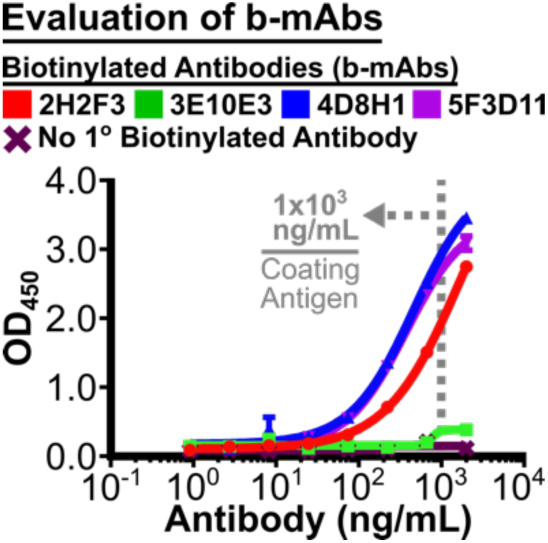
Biotinylation of 3E10E3 Leads to Loss of RoxP Binding. Indirect ELISA (iELISA) curves showing detection of recombinant apo-RoxP-tagless (coating concentration 1000 ng/mL) by four biotinylated anti-RoxP monoclonal antibodies (b-mAbs): b-2H2F3, b-3E10E3, b-4D8H1, b-5F3D11. Antibody concentrations: 1-2000 ng/mL. 2^nd^ layer: SA-HRP. Negative control: No 1° anti-RoxP b-mAb.

**SI Figure 10.**
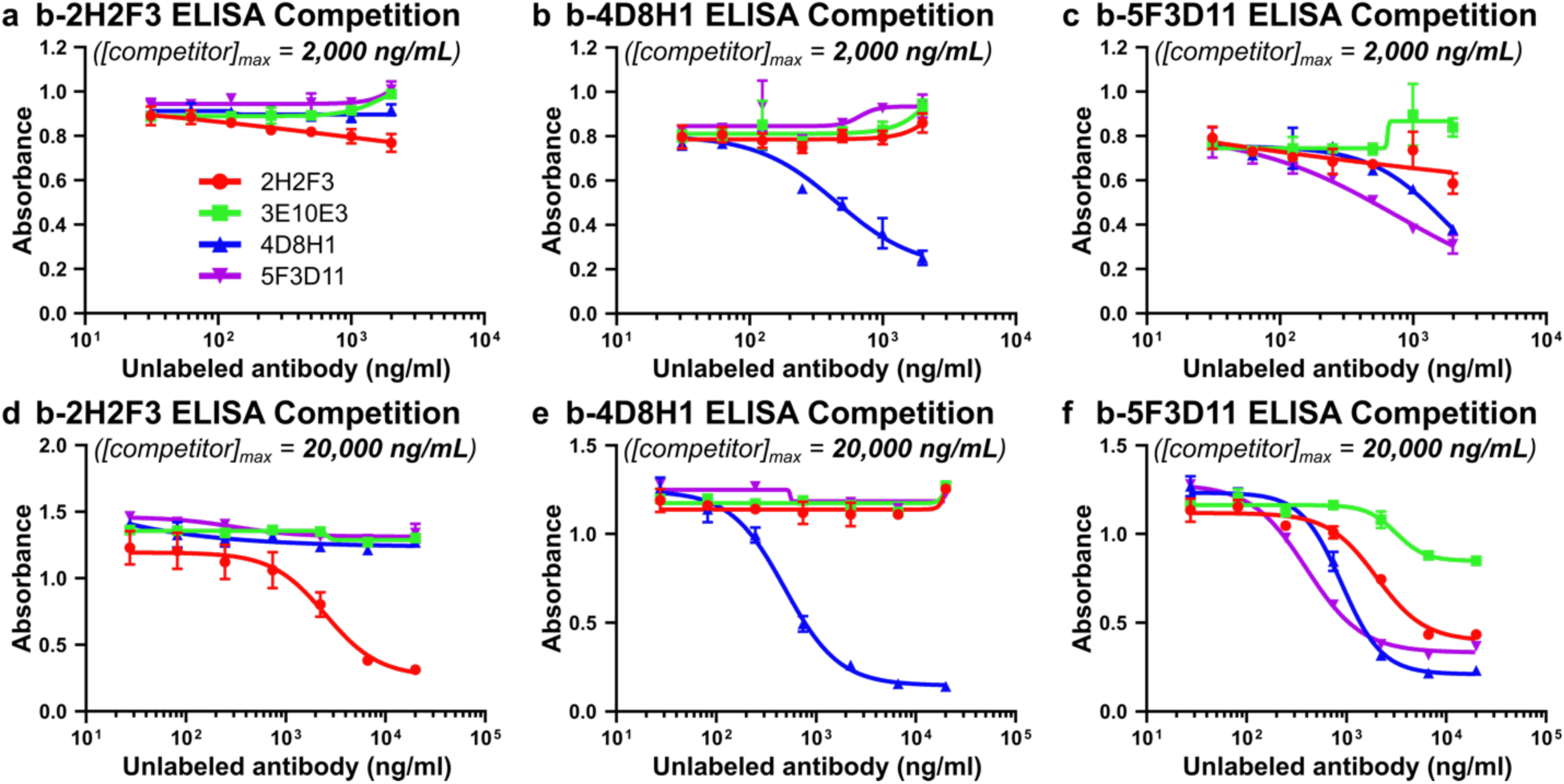
Competition ELISAs to Identify Epitope Overlaps. RoxP detection by iELISA using biotinylated monoclonal antibody (b-mAb) incubated in the presence of unlabeled competitor antibodies (2H2F3, 3E10E3, 4D8H1, 5F3D11). Primary detection reagent: (**a**,**d**) b-2H2F3, (**b**,**e**) b-4D8H1, (**c**,**f**) b-5F3D11. Titration of competitor antibodies were (**a**-**c**) 31-2000 ng/mL and (**d**-**f**) 27-20,000 ng/mL. (**d**,**f**) These panels are also shown in Fig 6b, but they were included in **SI Fig 10** for comparison to **SI Fig 10a** and **10c**, respectively. Coating concentrations: (**a**,**c**,**d**,**f**: 1000 ng/mL; **b**,**e**: 500 ng/mL).

**SI Figure 11.**
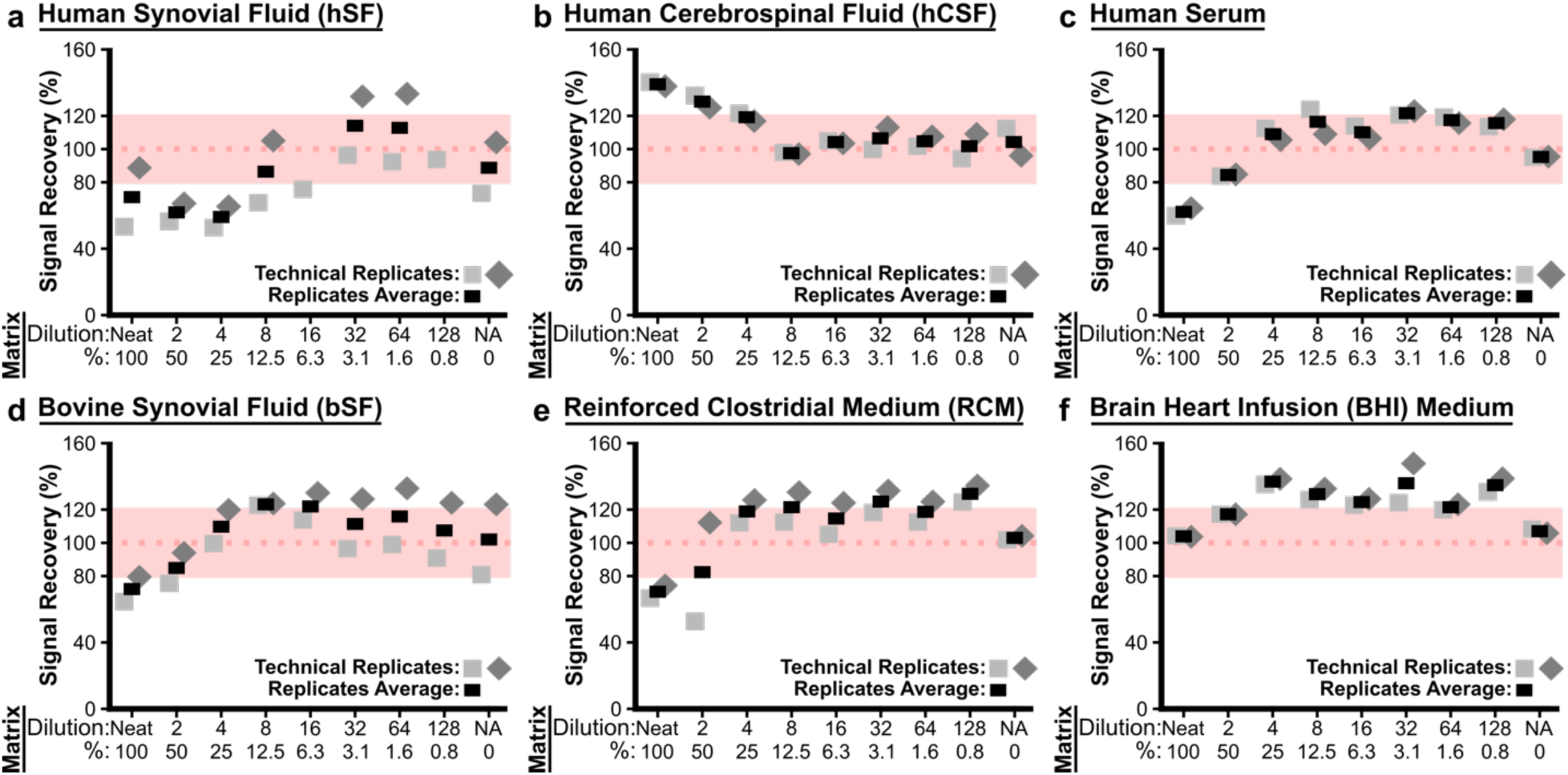
Spike-and-Recovery Sandwich ELISA (sar-sELISA) in Complex Samples. Recombinant RoxP was spiked into serially diluted **(a)** human synovial fluid (hSF), **(b)** human cerebrospinal fluid (hCSF), **(c)** human serum, **(d)** bovine synovial fluid (bSF), **(e)** reinforced clostridial medium (RCM), and **(f)** brain-heart infusion (BHI). Samples were analyzed by sandwich ELISA (sELISA) in duplicates. Each plot shows the empirically calculated percent recovery (%) based on a RoxP standard curve (12.5-400 ng/mL in ELISA-BB). These percent recovery (%) values are also equivalent to (ng/mL), as the spike concentration in this experiment was 100 ng/mL. The horizontal red line indicates 100% recovery. (**a**) Two values (32- and 128-fold) were not included in this analysis, as their values (395% and 302% signal recovery, respectively) were clear outliers likely due to an experiment artifact (e.g., loose tip(s) during ELISA workflow).

**SI Figure 12.**
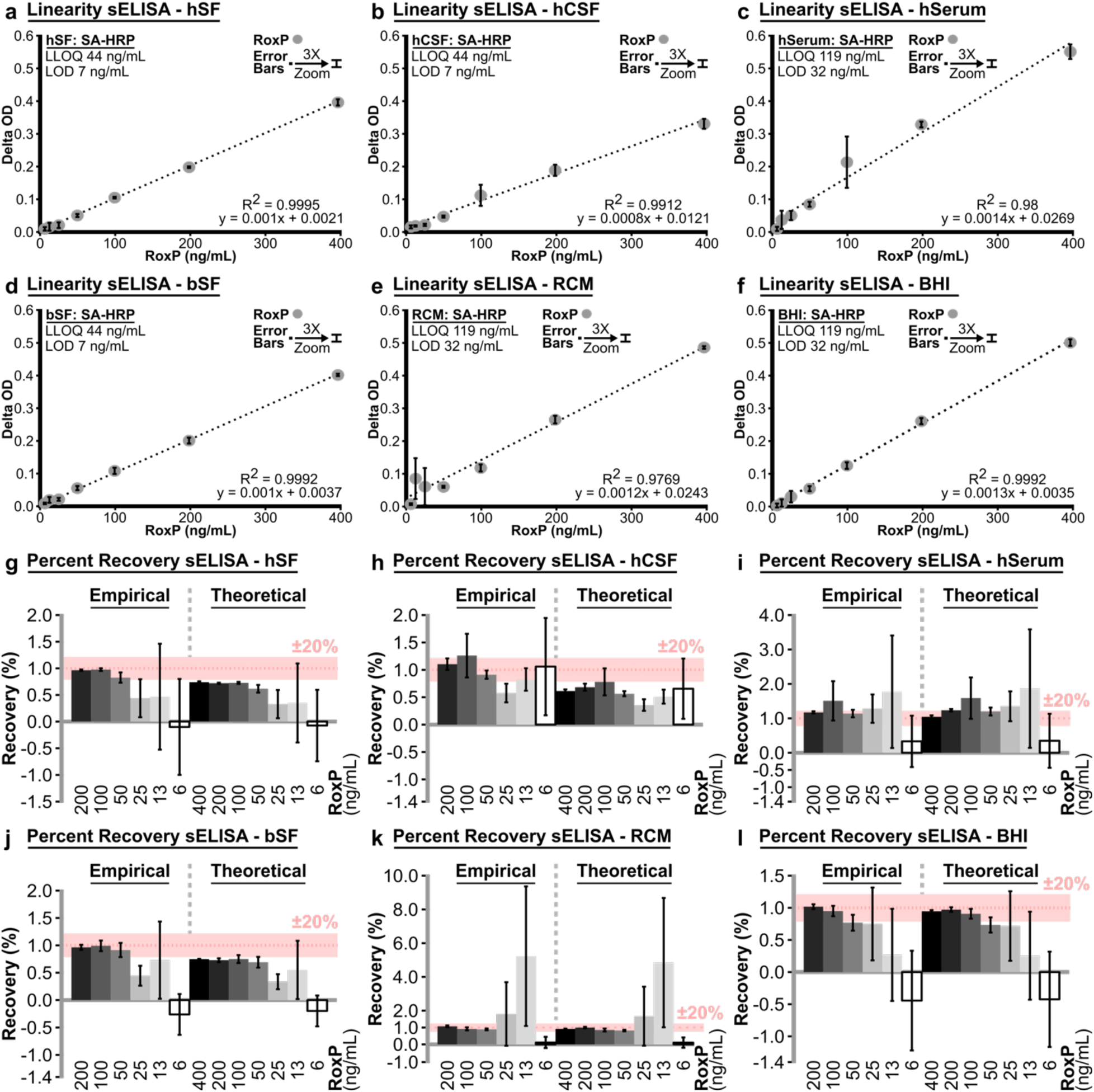
Assessing Sandwich ELISA (sELISA) Linearity/Recovery in Complex Samples. The (**a**-**f**) linearity and (**g**-**l**) percent recovery of the anti-RoxP sELISA was assessed in three human biofluids: (**a**) hSF, (**b**) hCSF, (**c**) human serum (hSerum); an animal biofluid that can be used in future studies: (**d**) bovine synovial fluid (bSF); and two types of *C. acnes* growth medium: (**e**) RCM, (**f**) BHI. (**a**-**f**) All complex samples displayed a linear relationship between concentration and delta OD under the evaluated experimental conditions with low nanogram LOD/LLOQ (LOD 7-32 ng/mL, LLOQ 44-119). LOD and LLOQ were evaluated per ELISA plate using 0 mg/mL RoxP (9 wells/plate) in ELISA-BB (plate #1: **a**,**b**,**d**; plate #2: **c**,**e**,**f**). (**g**-**l**) A comparison of empirical to theoretical recovery suggests that the standard curve RoxP stock used for plate #1 (**g**,**h**,**j**) had an initial concentration ∼20% lower (e.g., 320 instead of 400 ng/mL RoxP) than expected when compared to the stock used for plate #2 (**I**,**k**,**l**). (**a**,**g**) The data in this figure was collected on 20230512, while the data in Fig 6e is a separate hSF sELISA experiment that was performed on 20230920. (**c**,**e**,**i**,**k**) The large error bars of several wells in these panels (e.g., panel **k:**12.5 and 25 ng/mL RoxP) were likely due to a pipetting error from one or more loose micropipettor tips. (**a**-**l**) Error bars represent standard deviation (SD) of technical triplicates. Labeled concentrations were rounded to the first whole digit for the purposes of figure assembly.

**SI Figure 13.**
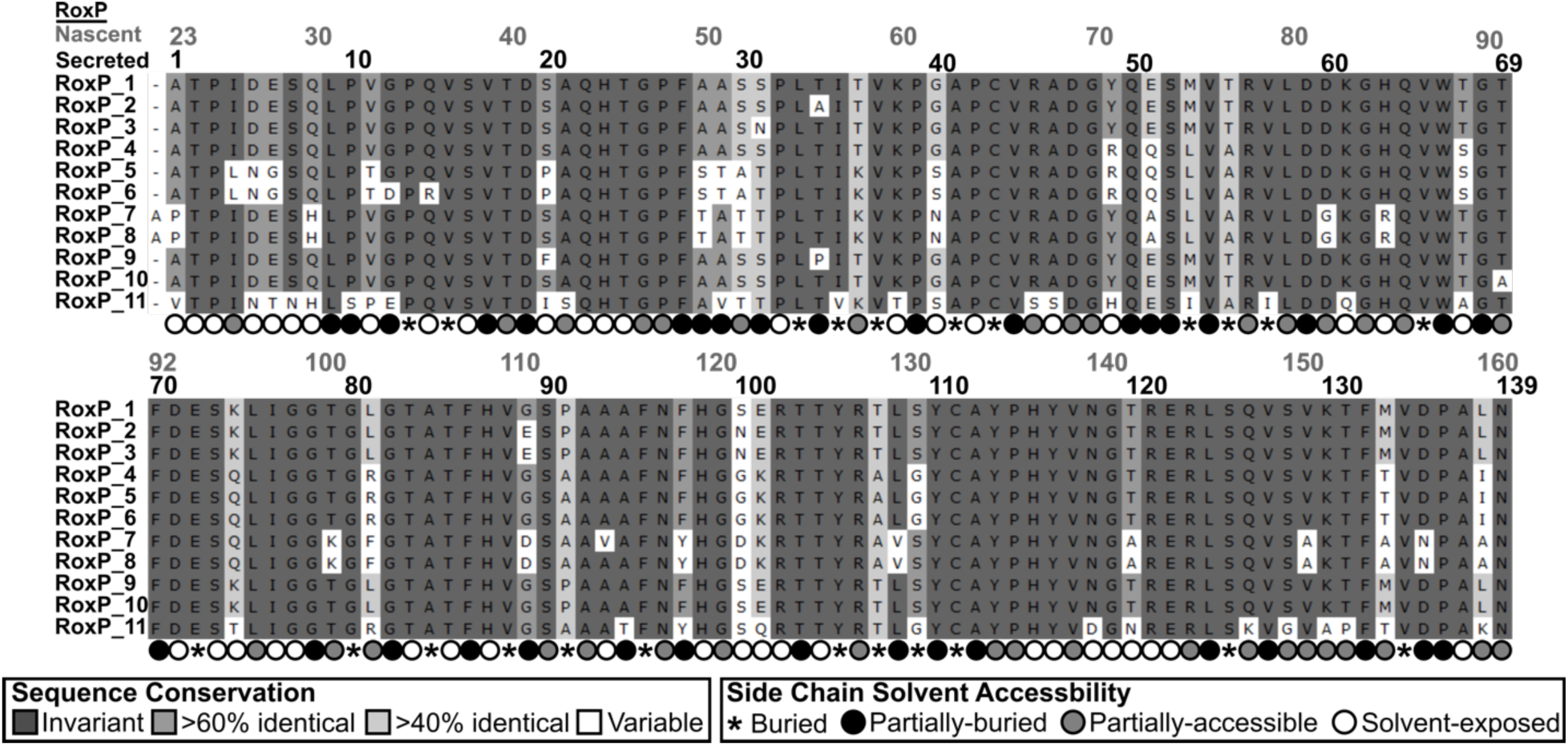
Multiple Sequence Alignment of Recombinantly Expressed RoxP. MSA of 11 RoxP orthologs expressed recombinantly in *Escherichia coli* is shown. Residue numbering of the *<u>nascent</u>* polypeptide chain and predicted, mature, *<u>secreted</u>*RoxP is shown. RoxP signal peptide cleavage was predicted by SignalP-4.1^1^. All residue positions noted in the text of this paper refer to mature, *<u>secreted</u>* RoxP (i.e., post-signal peptide cleavage) numbering. Residue backgrounds are colored according to sequence identity and variable positions are highlighted in white. The surface-exposure of each residue’s side chain in the apo-RoxP NMR structure (PDB 7bcj)^2^ as calculated by GetArea^3^ is shown below the alignment as a percentage exposure compared to the same residue in a tripeptide (Gly-X-Gly) random coil. The first digit of a residue position was aligned to the corresponding MSA column e.g., 1 of **<u>1</u>**0 is centered on the corresponding MSA column for position 10.

**SI Figure 14.**
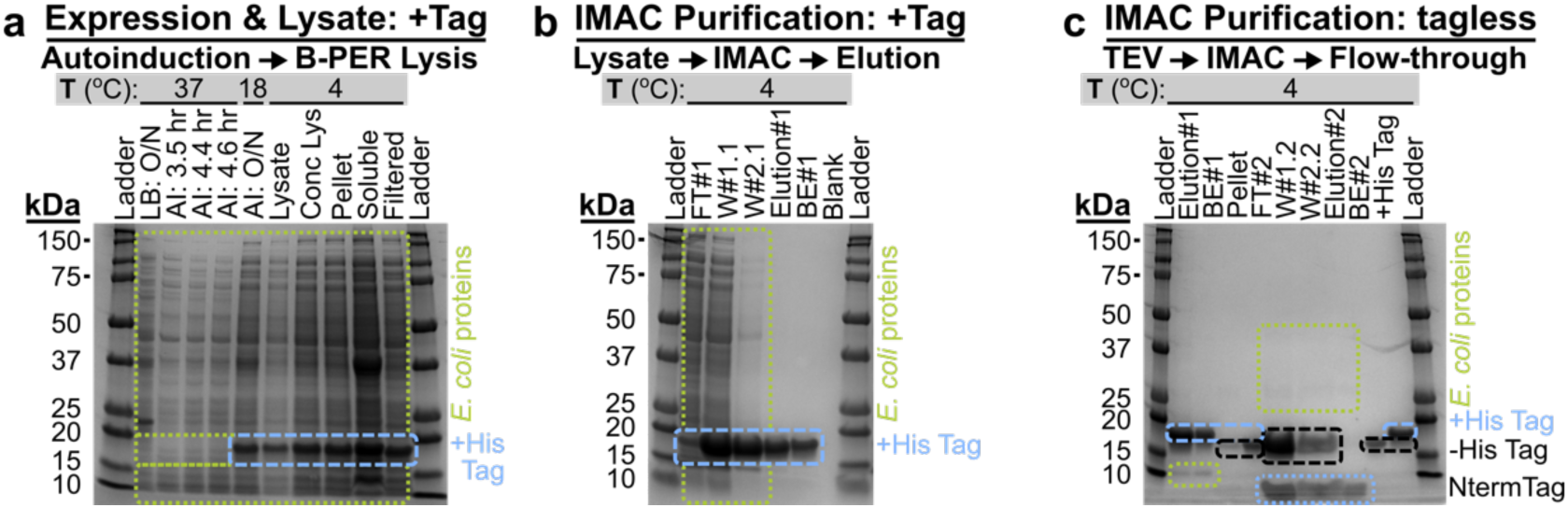
Small-scale Recombinant RoxP Expression and Purification. Representative gels from a small-scale expression and purification of recombinant RoxP are shown for one preparation of apo-RoxP_1-tagless. All other orthologs purified using the small-scale expression and purification of recombinant RoxP strategy looked similar. Samples presented in each panel were collected as a part of the small-scale expression and purification of recombinant RoxP protocol, and they are presented (from left-to-right) as they were collected in time during the protocol. Temperature (T, °C) for each protocol step is shown above the relevant lane. (**a**) Expression of 8His-TEV-RoxP_1 (+His Tag: 17.2 kDa) in *E. coli* using autoinduction followed by lysate preparation B-PER (non-ionic detergent-based cell lysis reagent) and post-lysate processing (e.g., centrifugation, filtration). (**b**) Immobilized metal affinity purification (IMAC) using one His-Spin column of 8His-TEV-RoxP_1 followed by buffer-exchange (BE) into HEPES Sizing Buffer (HSB). (**c**) TEV (Tobacco Etch Virus protease) proteolysis of 8His-TEV-RoxP_1 in HSB followed by His-Spin IMAC capture of tagged (+Tag) proteins (e.g., 8His-TEV-RoxP_1 [17.2 kDa], 8His-TEV [28 kDa], tag alone [2.3 kDa], *E. coli* proteins that bind IMAC columns). The final sample of apo-RoxP_1-tagless (-His Tag: 14.9 kDa) in HSB used in experiments is shown in BE#2. Two samples (Elution#1, BE#1) from panel (**b**) are shown in (**c**) aid in identification of N-terminal tag removal by TEV (see 2.3 kDa drop down in RoxP from 17.2 to 14.9 kDa). ***Abbreviations***: AI (autoinduction culture), Conc (Concentrate from Vivaspin centrifugal concentrator), FT (flow-through from His-Spin column), +His Tag (8His-TEV_RoxP_1), hr (hour), Lys (lysate), Nterm (N-terminal), O/N (overnight culture), +/-Tag (indicates the presence or absence of N-terminal tag on RoxP), W#1 (10 mM Imidazole IMAC wash), W#2 (20 mM Imidazole IMAC wash).

**SI Figure 15.**
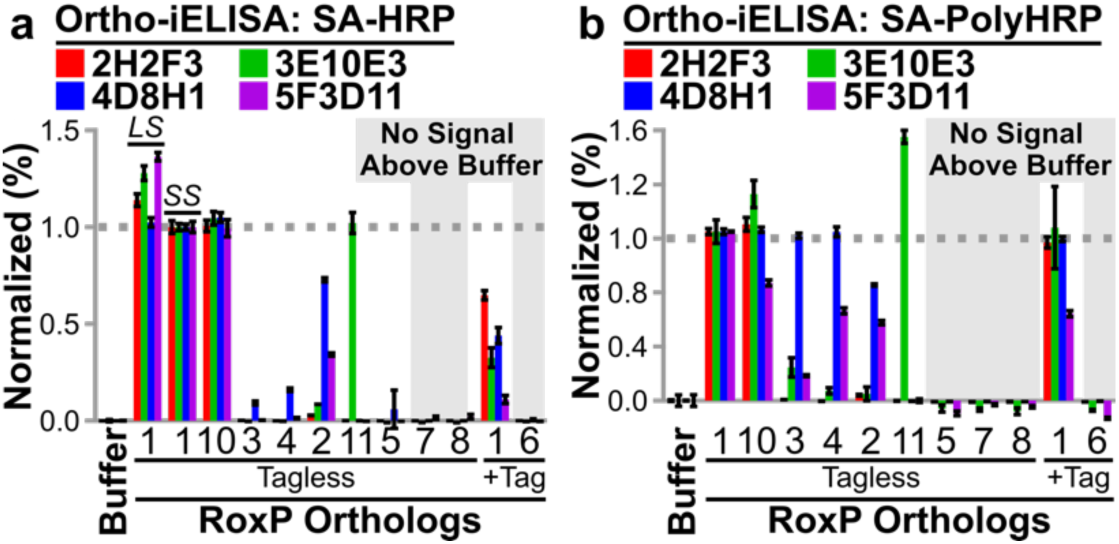
Determining Anti-RoxP Antibody Specificity for RoxP Orthologs. To determine if a panel of RoxP orthologs could be bound by anti-RoxP antibodies (2H2F3, 3E10E3, 4D8H1, 5F3D11), ten recombinant RoxP orthologs were evaluated by iELISA. (**a**,**b**) The coating concentration for both experiments was 1.0 µg/mL RoxP, but the terminal detection reagent varied between experiments; (**a**) SA-HRP, (**b**) SA-Poly-HRP). (**a**,**b**) Recombinant RoxP_2-5,7,8,10,11 were prepared as tagless proteins and compared to apo-RoxP_1-tagless. RoxP_6 was not soluble after tag removal, so RoxP_6+Tag was compared to RoxP_1+Tag. ELISA signal was normalized to SS (Small-Scale RoxP Protein Preparation – His-Spin Protocol) apo-RoxP_1-tagless (100%) and buffer-only controls (0%). (**a**) In addition to ortholog comparisons, a comparison between RoxP preparation protocols was performed that demonstrated equivalent antibody binding for SS and LS (Large-Scale RoxP Protein Preparation – Gravity-Flow Protocol) apo-RoxP_1-tagless, as the slight increase in LS signal for 3E10E3 and 5F3D11 is well within potential experimental error identified for this experiment (e.g., inter-spectrophotometer absorbance variability, assessments of high- vs. low-concentration protein stocks). All SS stocks were assessed on a DeNovix DS-11 FX+ spectrophotometer at lower protein concentrations (e.g., 0.1-2.0 mg/mL), while the LS stock was assessed on a NanoDrop® ND-1000 spectrophotometer at a high protein concentration (35.9 mg/mL) and two dilutions (1/10, 1/100).

**SI Figure 16.**
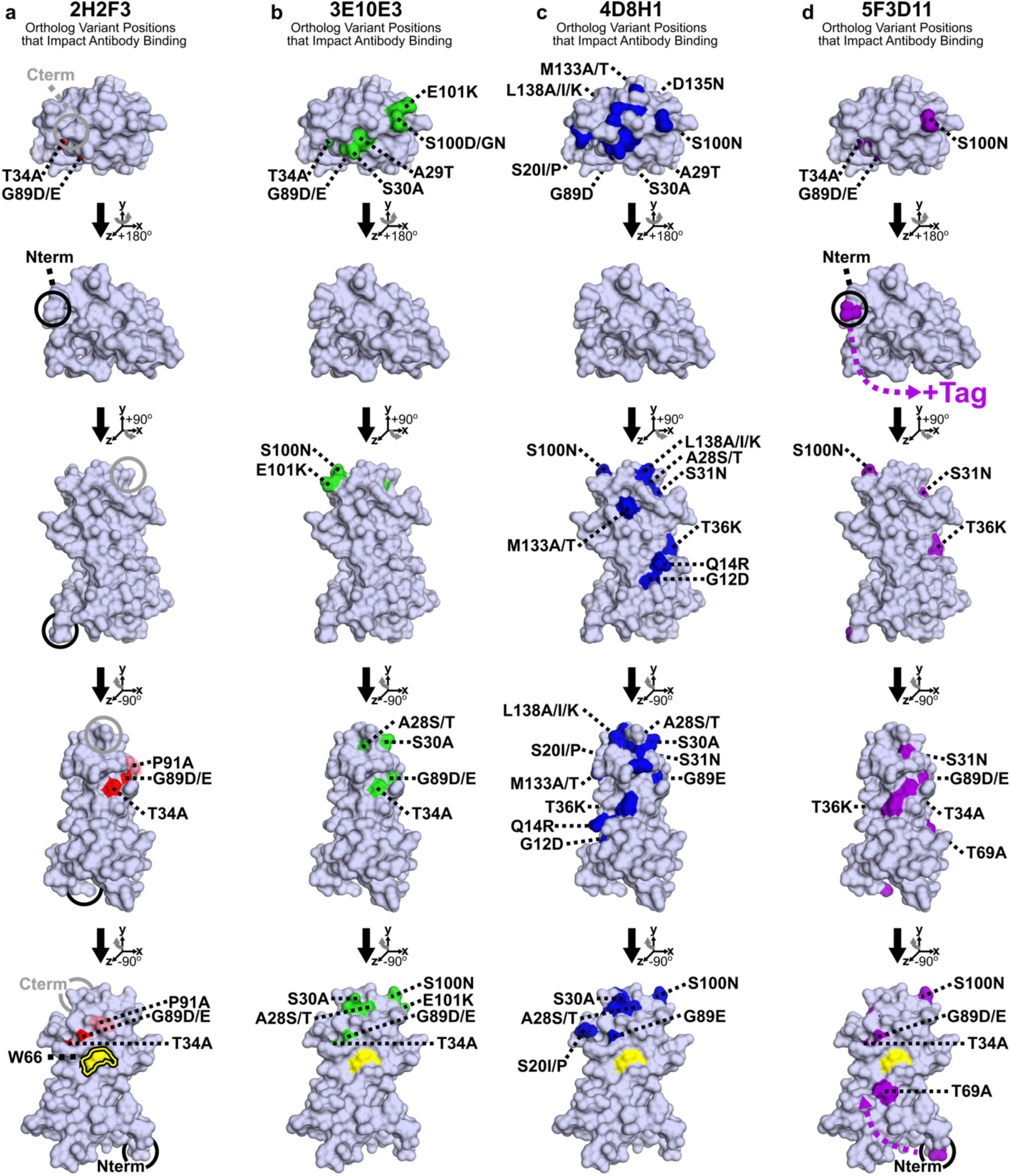
RoxP Ortholog Variant Positions that Impact Antibody Binding. Based on **SI Table 7**, RoxP ortholog variant positions that impact antibody binding were mapped onto the apo-RoxP_1 NMR model (PDB 7bcj) for (**a**) 2H2F3, (**b**) 3E10E3, (**c**) 4D8H1, and (**d**) 5F3D11. (**a**-**d**) Variant residues are colored according to antibody (see Fig 6g,**h**). RoxP variants discussed in the text are labeled. (**a**-**d**) Tryptophan 66 (W66) is (**a**) labeled and outlined (see Fig 5), and (**a**-**d**) it is also colored yellow. (**a**) RoxP P91 is not surface exposed, so it’s location and effect on 2H2F3 binding is indicated by a red oval with 30% transparency in the bottom two models (2^nd^ from bottom model: right side; bottom model: center of model). (**d**) The proposed extension of the N-terminal tag causing partial inhibition of 5F3D11 is shown as “+Tag” (from-the-top: 2^nd^ and 5^th^ RoxP models). See Fig 2 for additional structure figure details.

- Nielsen, H. *Methods Mol. Biol. Clifton NJ* 1611, 59–73 (2017)
- Stødkilde, K. *et al. Front. Cell. Infect. Microbiol.* 12, 803004 (2022)
- Fraczkiewicz, R. *et al. J. Comput. Chem.* 19, 319–333 (1998)
- Butler-Wu, S. M. *et al. J. Bacteriol.* 193, 3678–3678 (2011)
- Teufel, F. *et al. Nat. Biotechnol.* 40, 1023–1025 (2022)
- Petersen, T. N. *et al. Nat. Methods* 8, 785–786 (2011)
- Wylensek, D. *et al. Nat. Commun.* 11, 6389 (2020)
- McCance, R. A. *et al. Br. J. Nutr.* 14, 509–518 (1960)

**SI Table 1.**
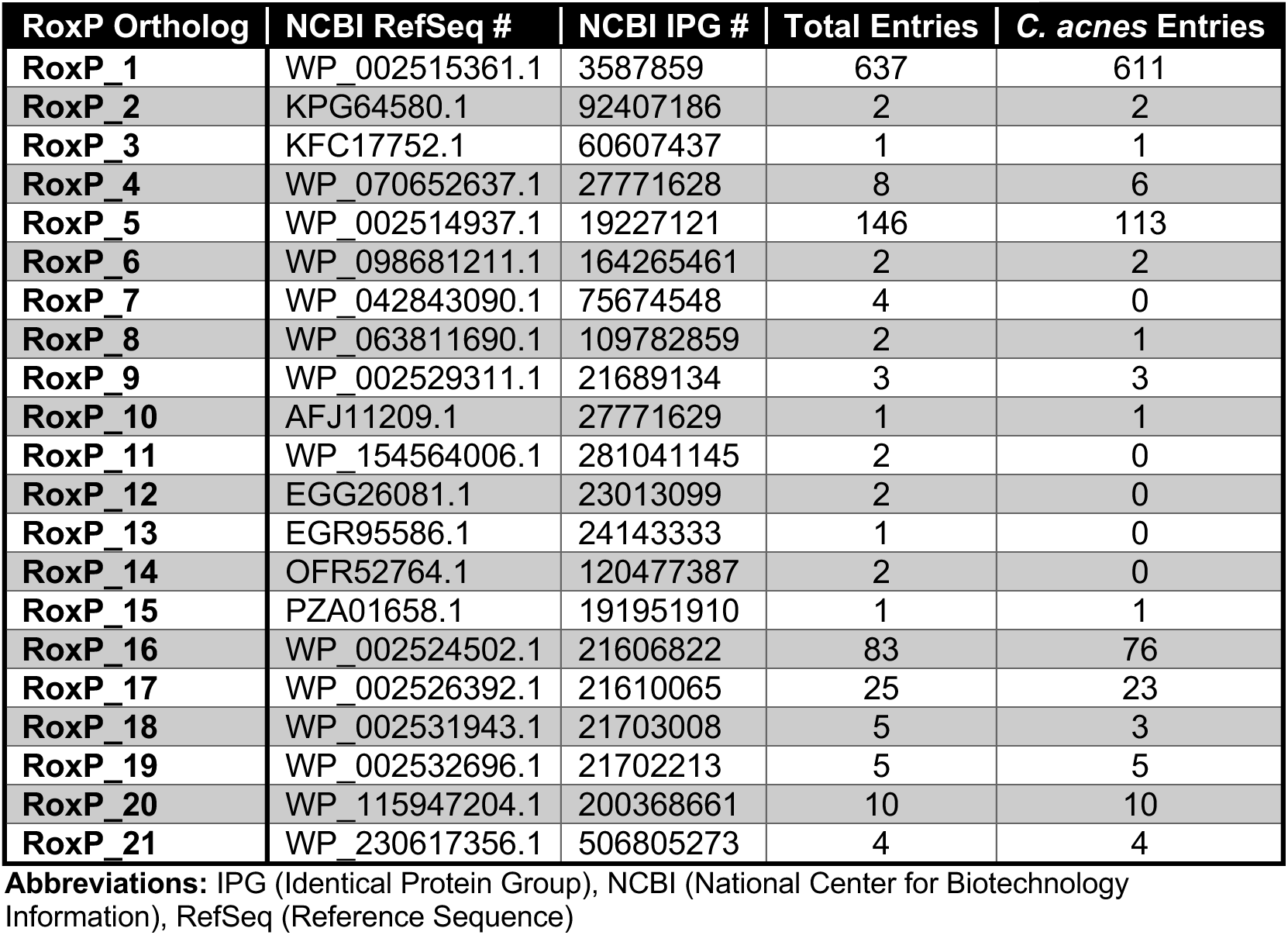
Identified RoxP Protein Sequences.

| **RoxP Ortholog** | **NCBI RefSeq #** | **NCBI IPG #** | **Total Entries** | ***C. acnes* Entries** |
| --- | --- | --- | --- | --- |
| **RoxP_1** | WP_002515361.1 | 3587859 | 637 | 611 |
| **RoxP_2** | KPG64580.1 | 92407186 | 2 | 2 |
| **RoxP_3** | KFC17752.1 | 60607437 | 1 | 1 |
| **RoxP_4** | WP_070652637.1 | 27771628 | 8 | 6 |
| **RoxP_5** | WP_002514937.1 | 19227121 | 146 | 113 |
| **RoxP_6** | WP_098681211.1 | 164265461 | 2 | 2 |
| **RoxP_7** | WP_042843090.1 | 75674548 | 4 | 0 |
| **RoxP_8** | WP_063811690.1 | 109782859 | 2 | 1 |
| **RoxP_9** | WP_002529311.1 | 21689134 | 3 | 3 |
| **RoxP_10** | AFJ11209.1 | 27771629 | 1 | 1 |
| **RoxP_11** | WP_154564006.1 | 281041145 | 2 | 0 |
| **RoxP_12** | EGG26081.1 | 23013099 | 2 | 0 |
| **RoxP_13** | EGR95586.1 | 24143333 | 1 | 0 |
| **RoxP_14** | OFR52764.1 | 120477387 | 2 | 0 |
| **RoxP_15** | PZA01658.1 | 191951910 | 1 | 1 |
| **RoxP_16** | WP_002524502.1 | 21606822 | 83 | 76 |
| **RoxP_17** | WP_002526392.1 | 21610065 | 25 | 23 |
| **RoxP_18** | WP_002531943.1 | 21703008 | 5 | 3 |
| **RoxP_19** | WP_002532696.1 | 21702213 | 5 | 5 |
| **RoxP_20** | WP_115947204.1 | 200368661 | 10 | 10 |
| **RoxP_21** | WP_230617356.1 | 506805273 | 4 | 4 |

**Abbreviations:** IPG (Identical Protein Group), NCBI (National Center for Biotechnology Information), RefSeq (Reference Sequence)

**SI Table 2:**
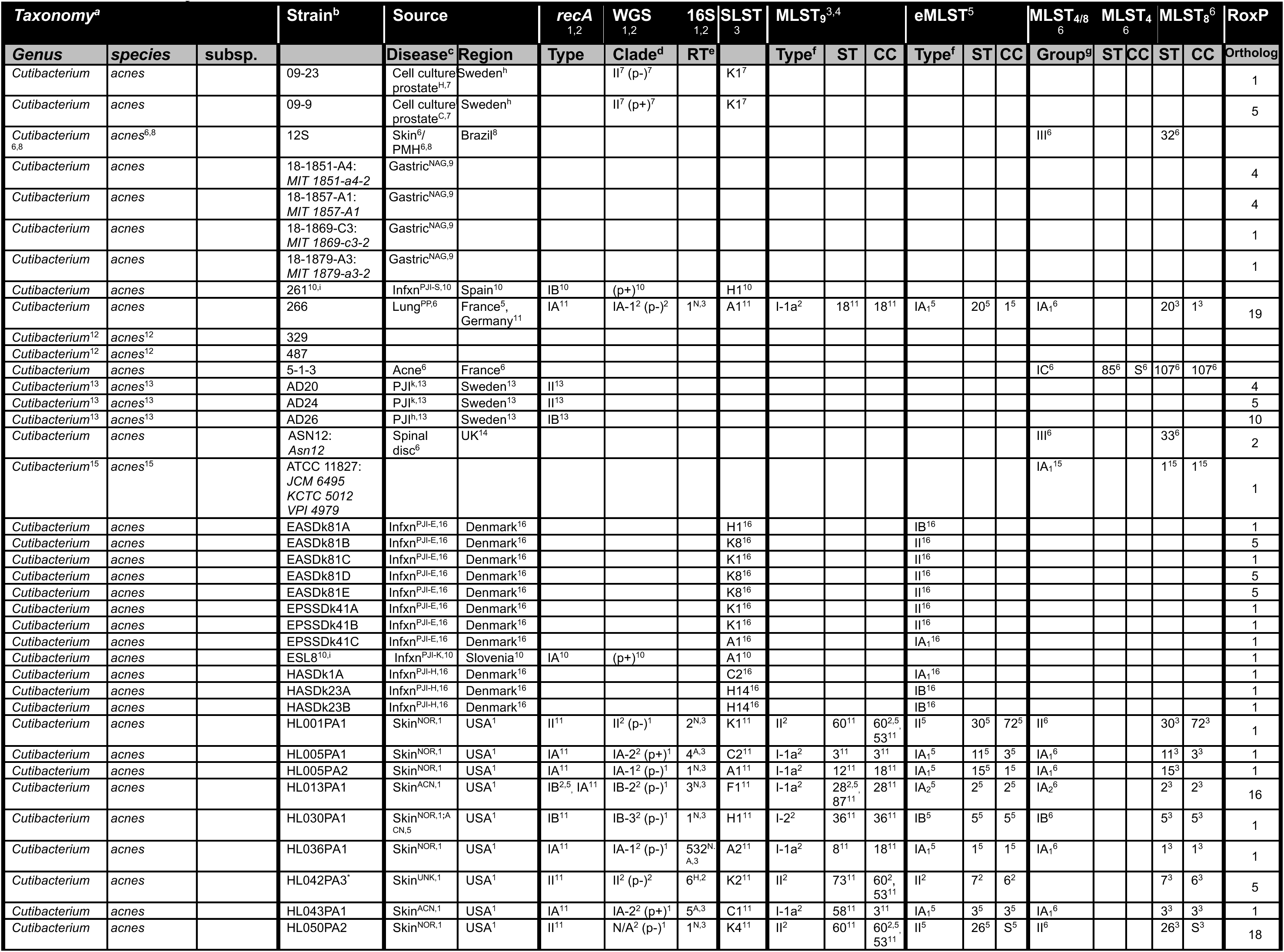
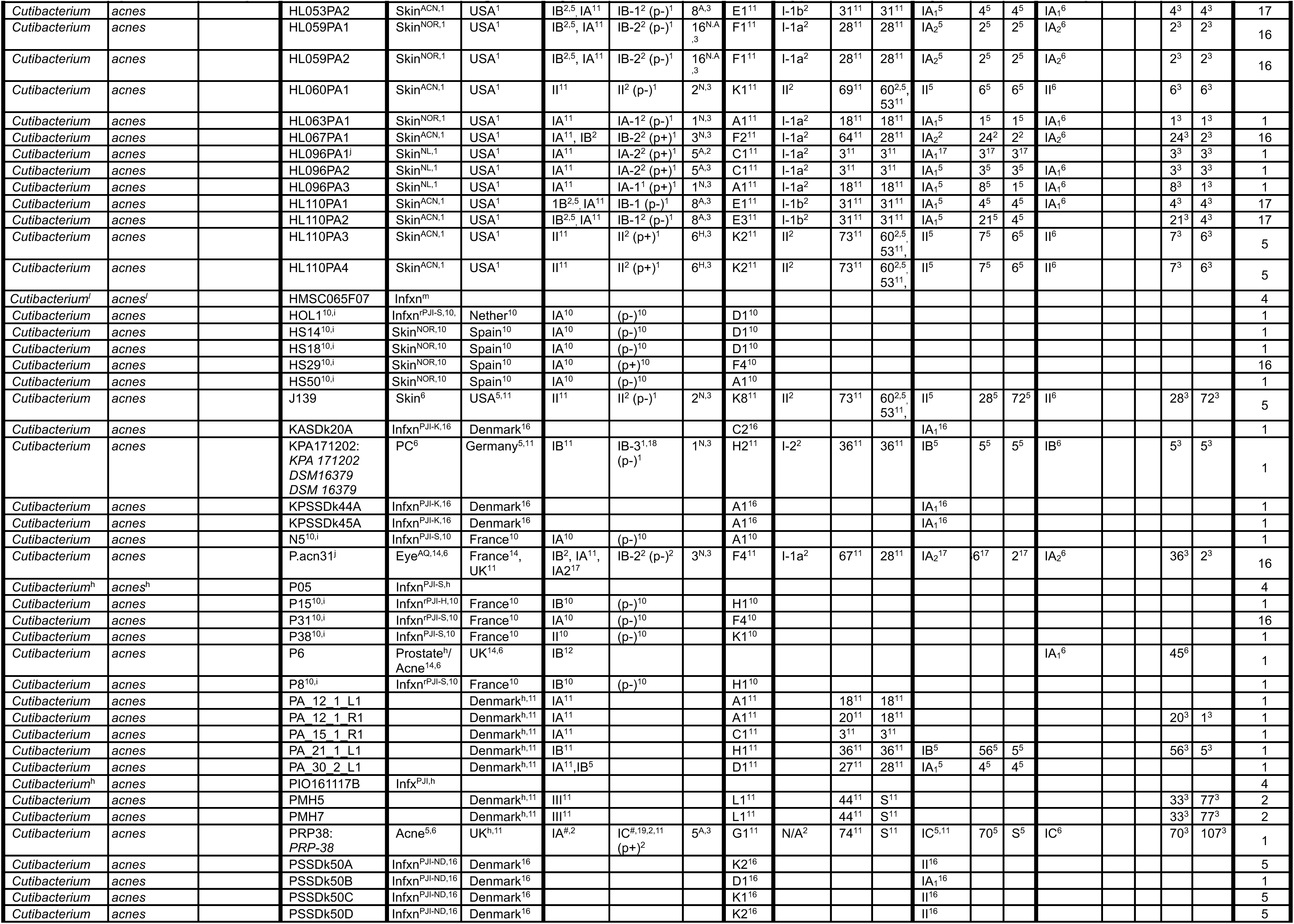
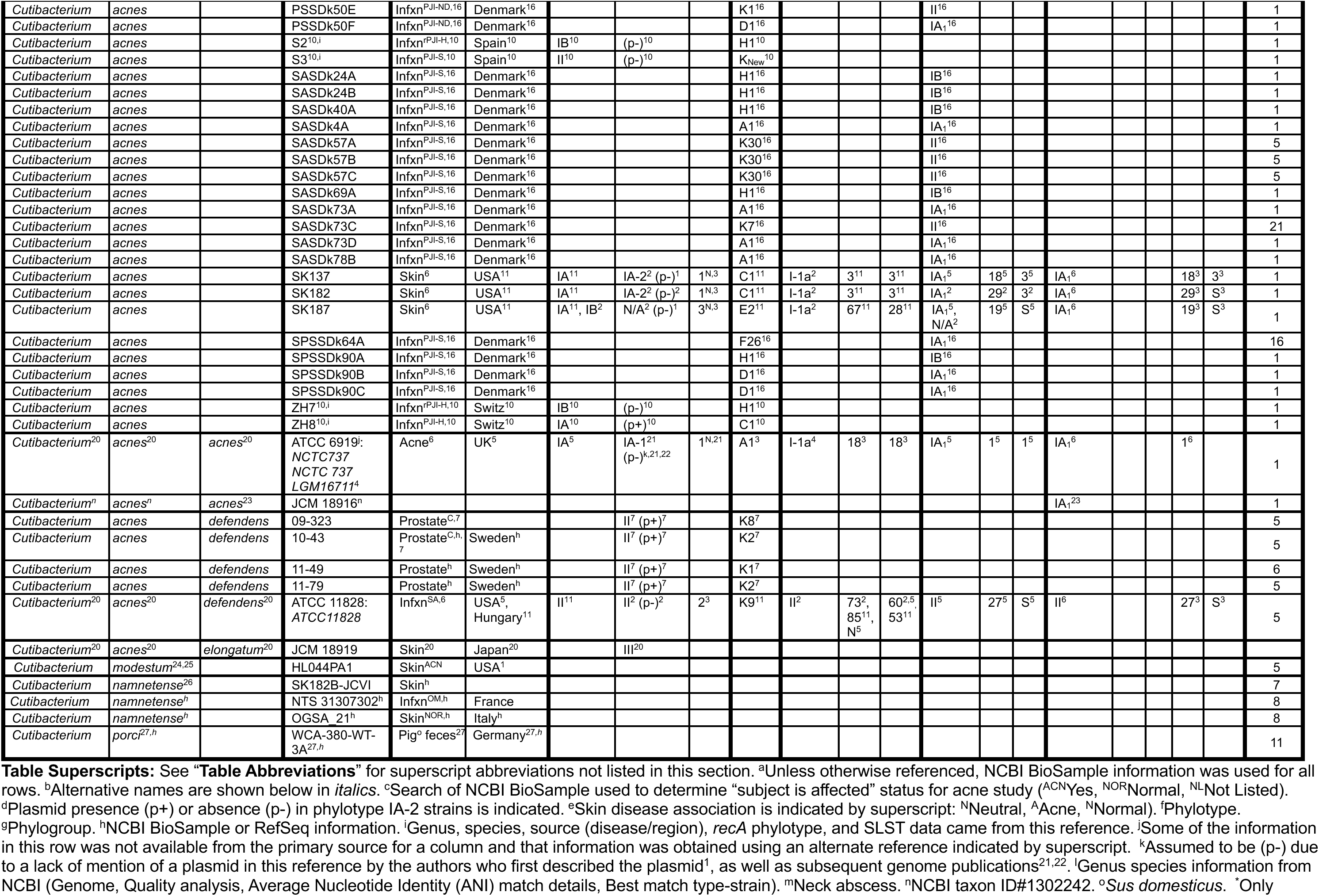
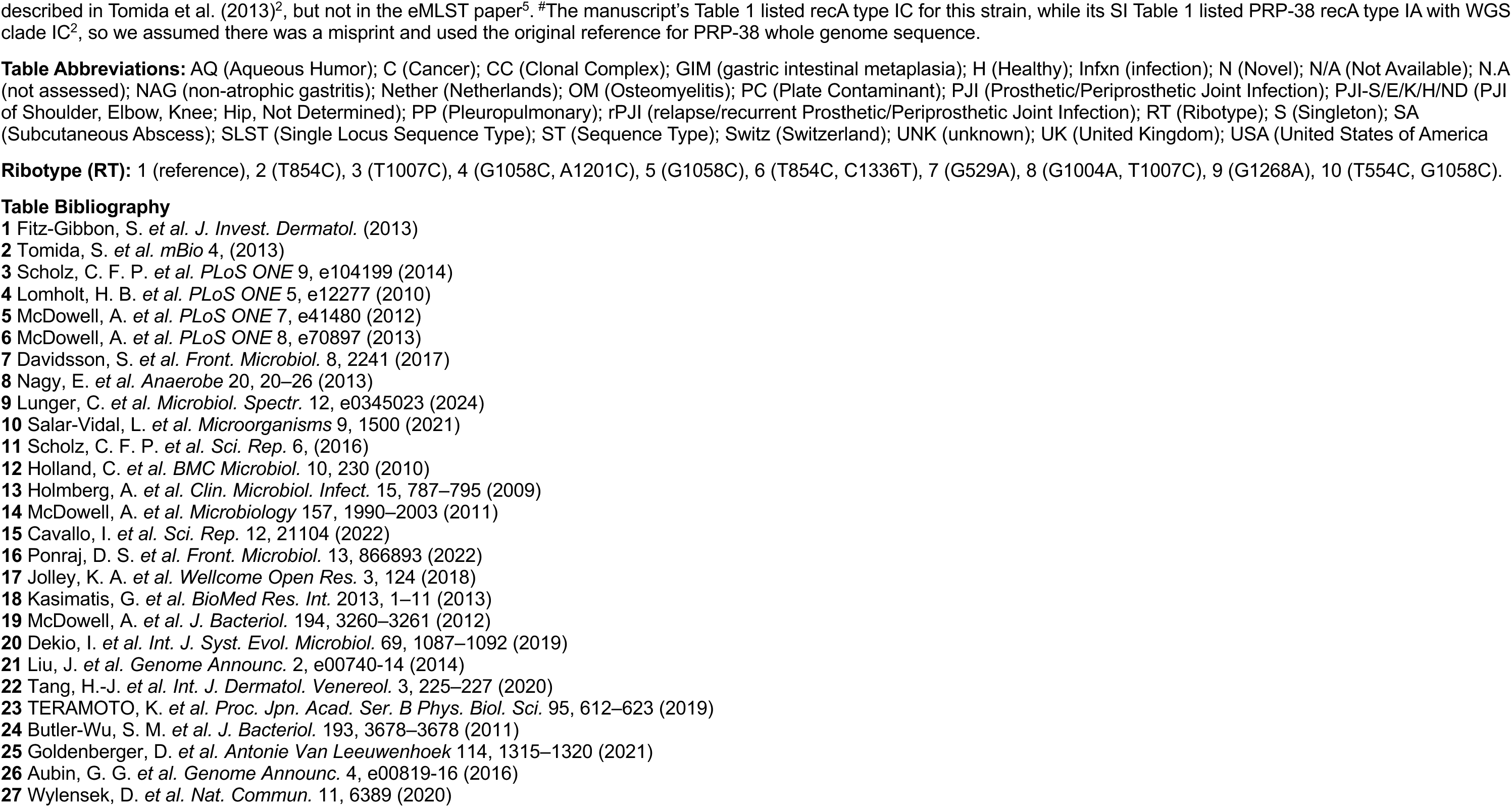
Summary of Cutibacterium Strain Information.

**SI Table 3:**
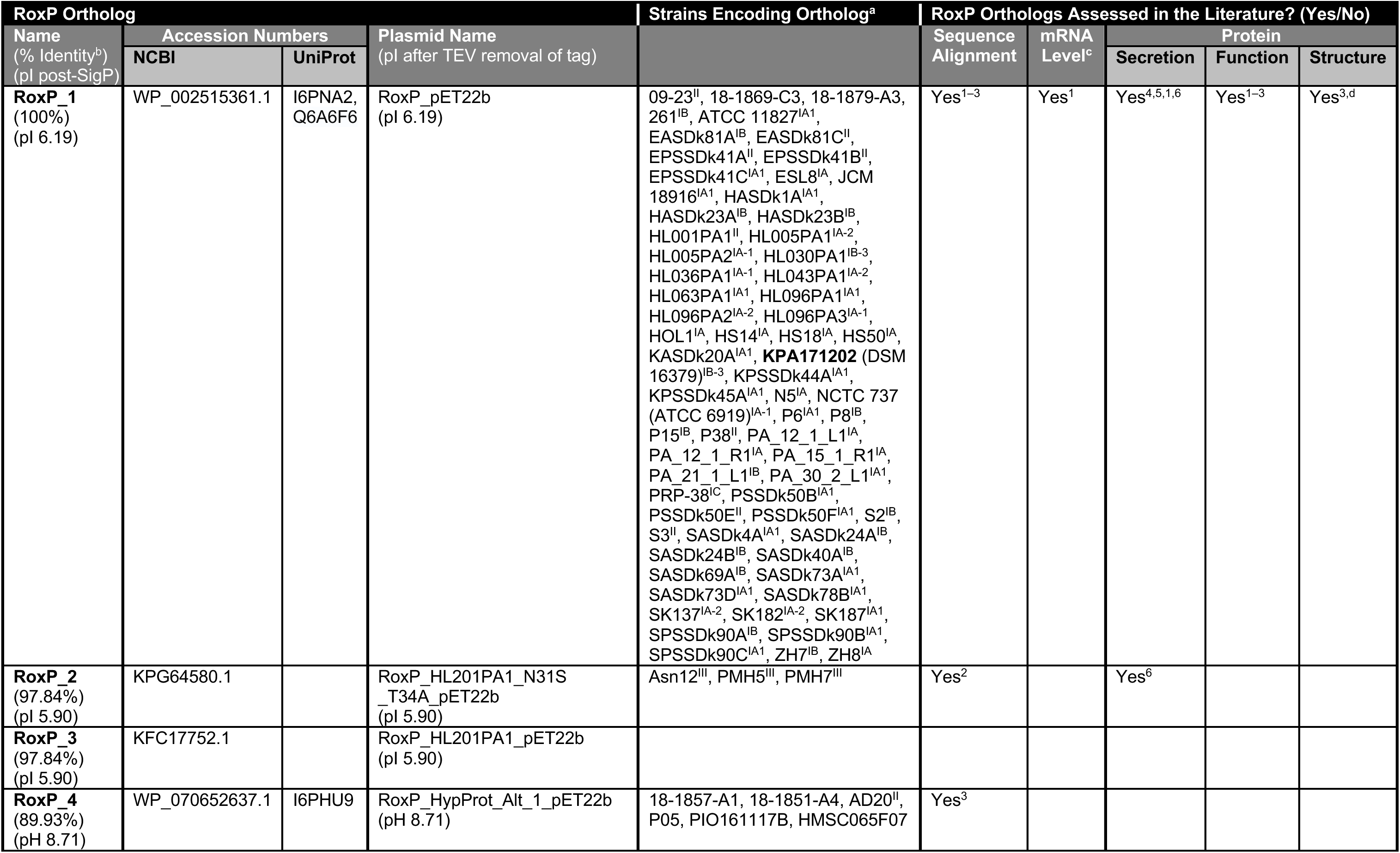
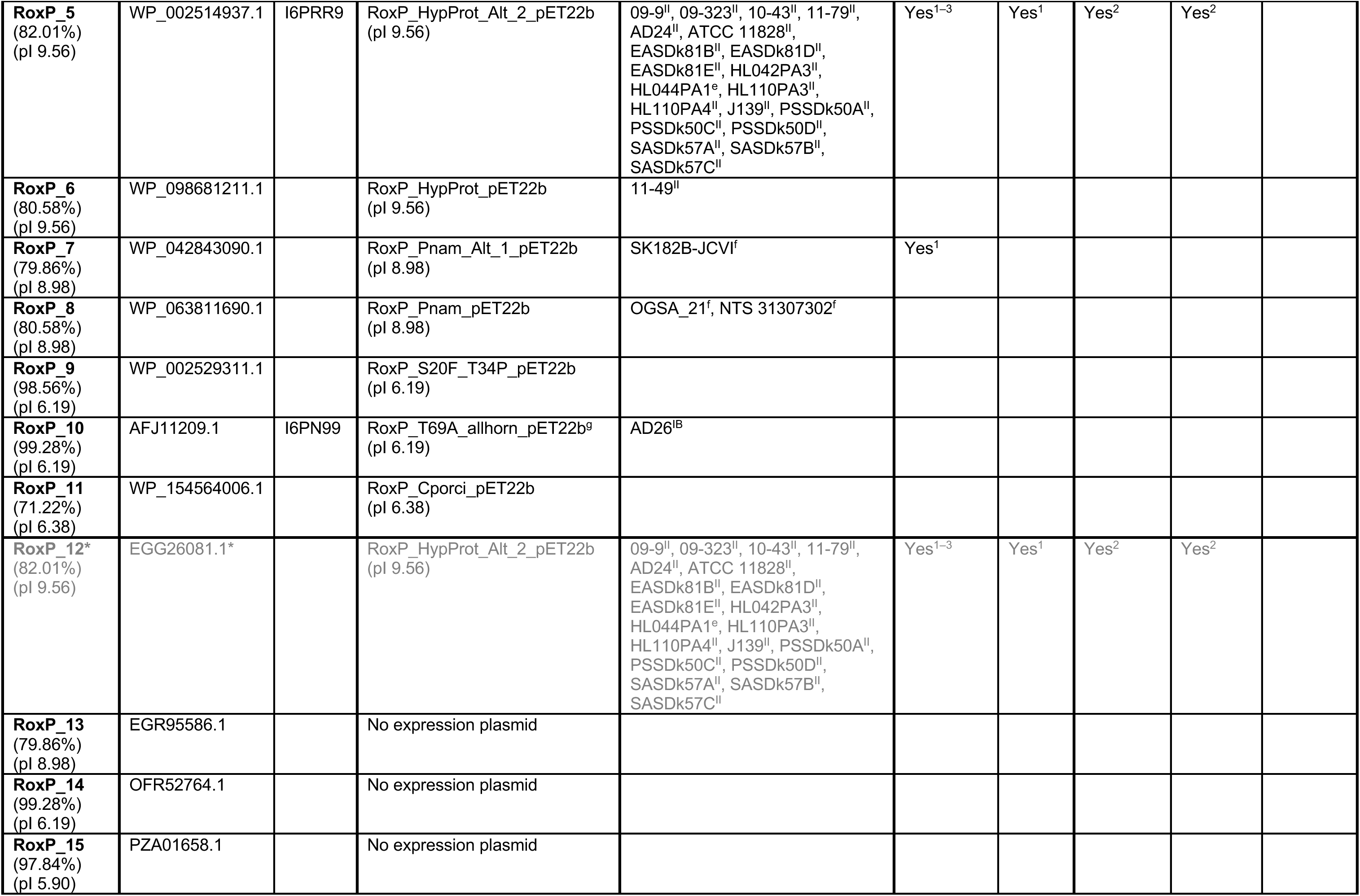
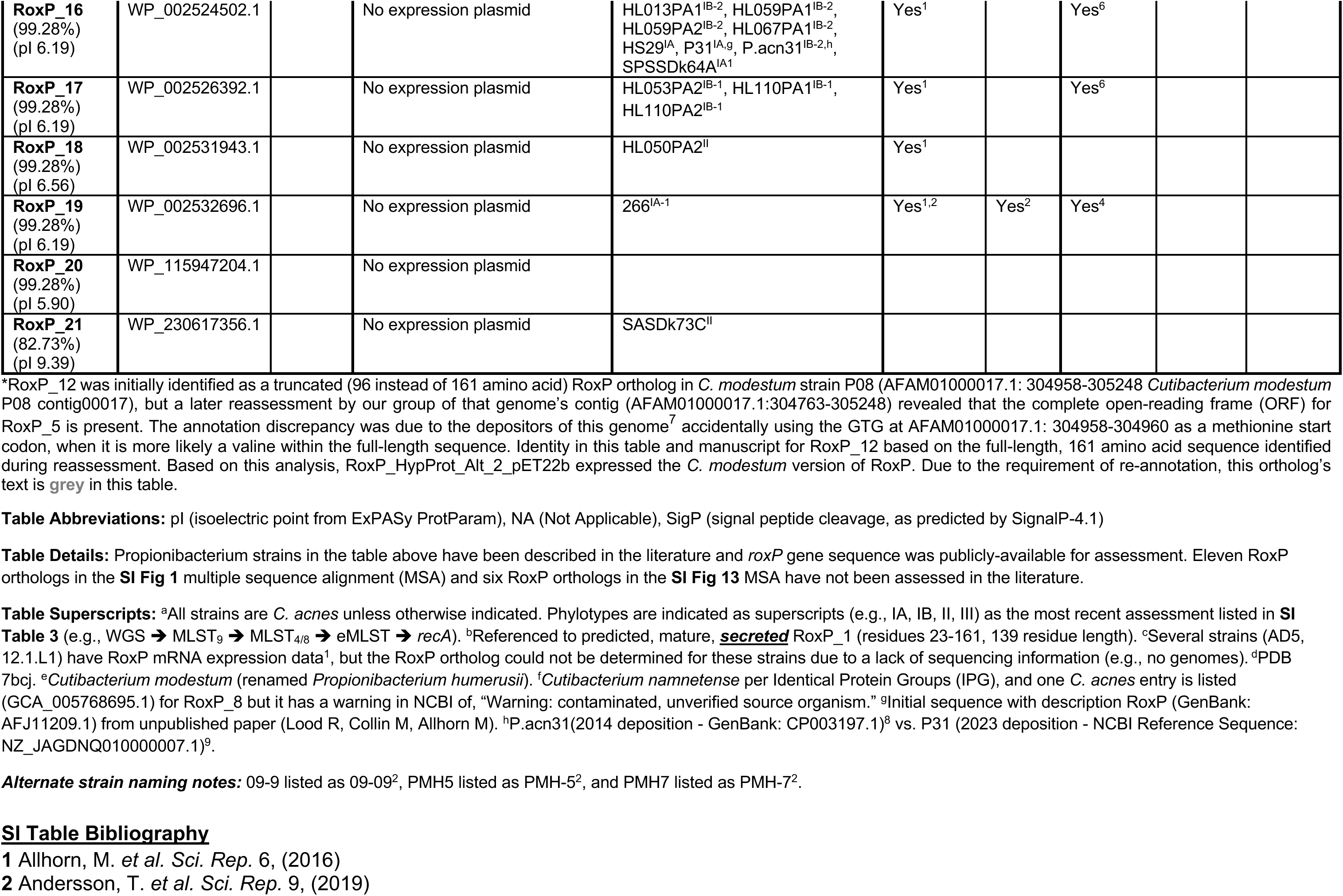
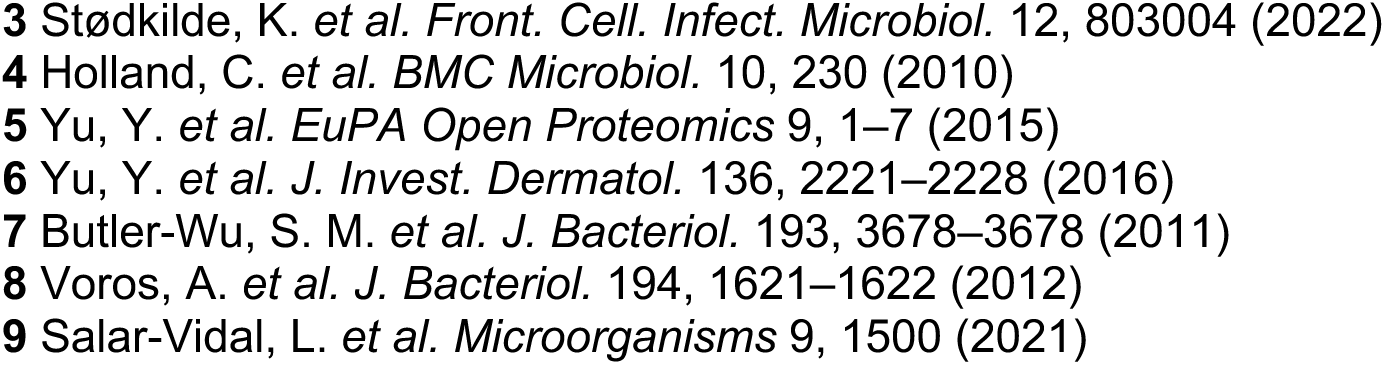
Summary of RoxP Orthologs Assessed in the Literature or This Study.

**SI Table 4:**
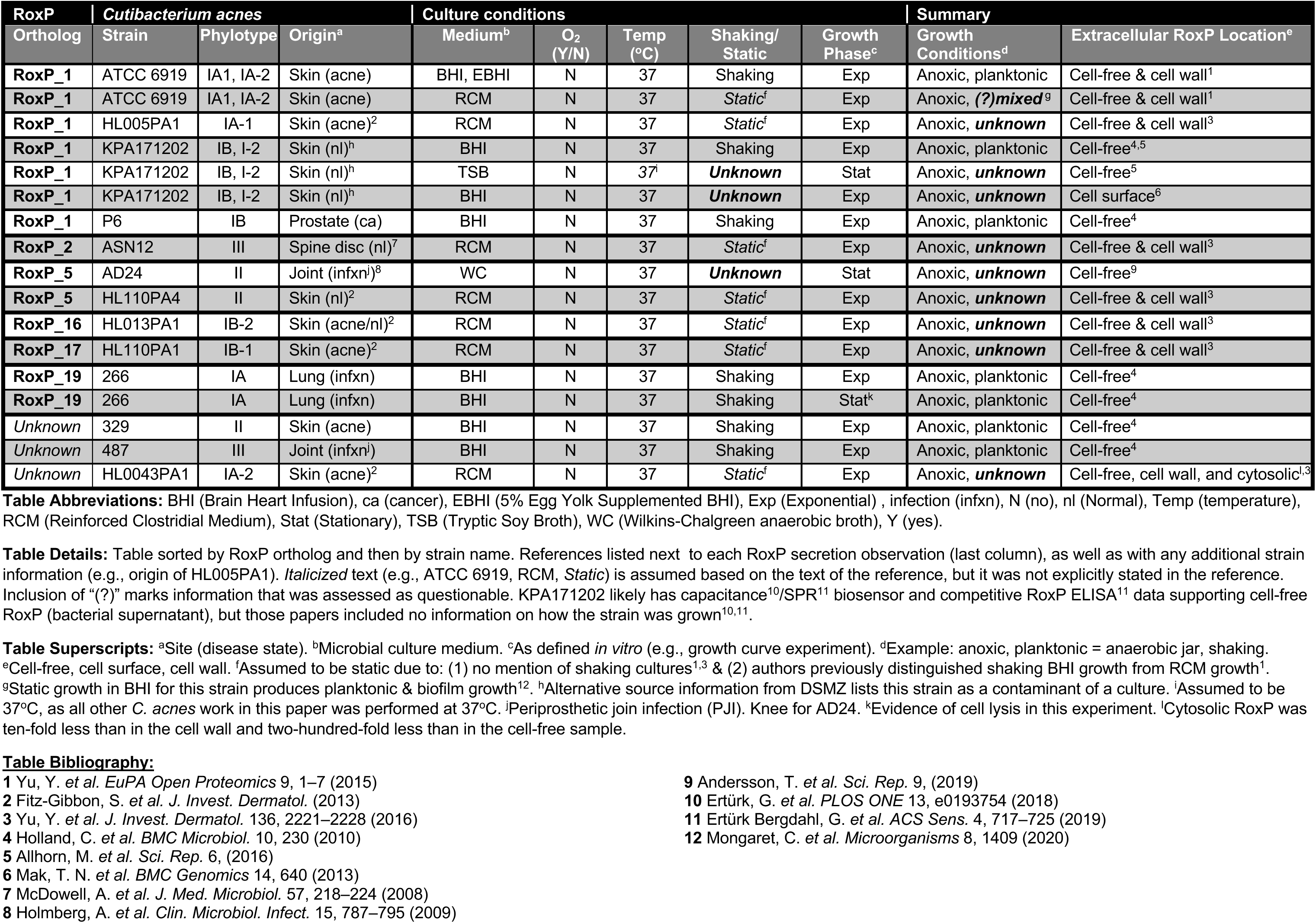
RoxP Secretion Assessment in the Literature.

**SI Table 5.**
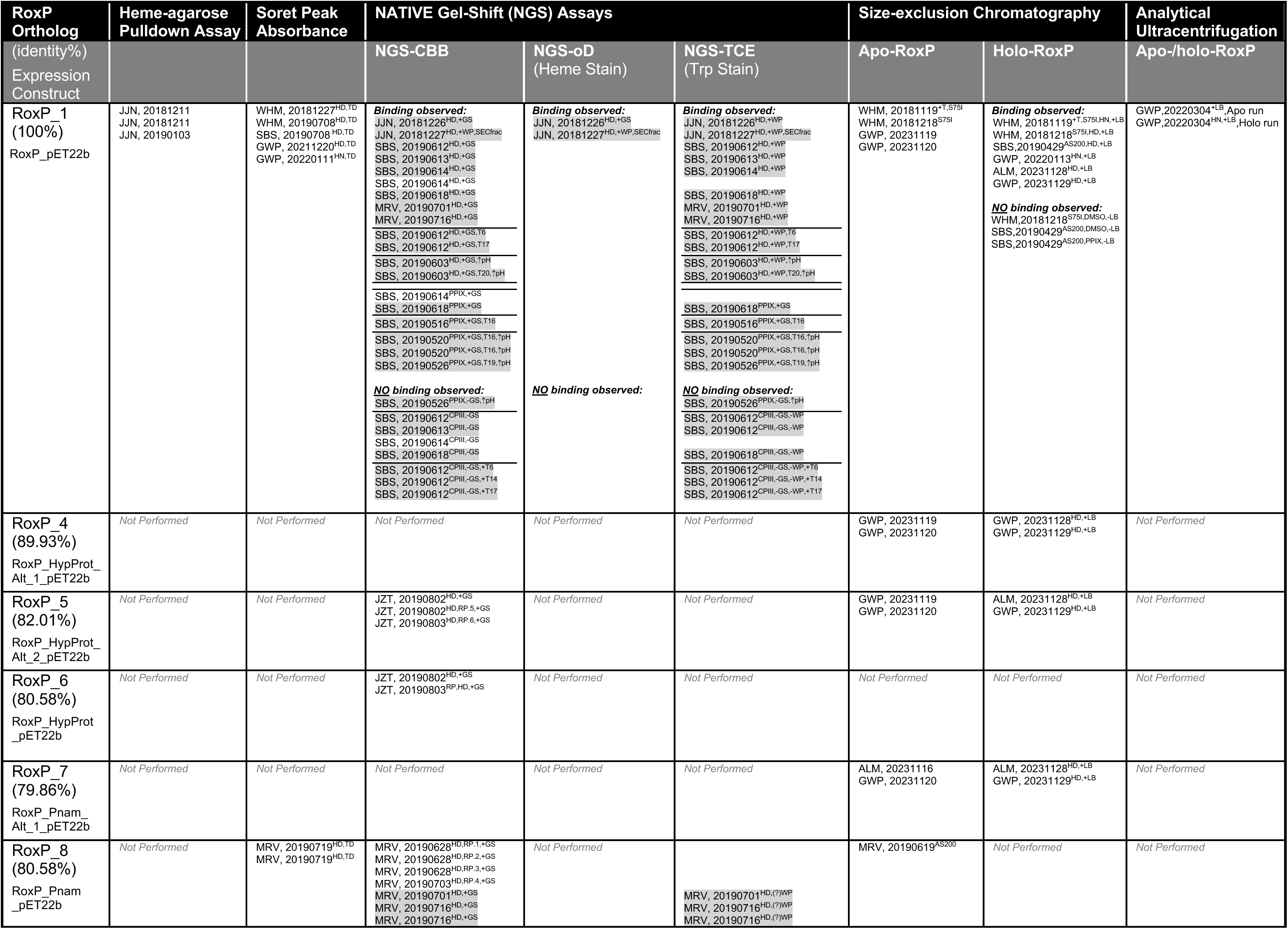
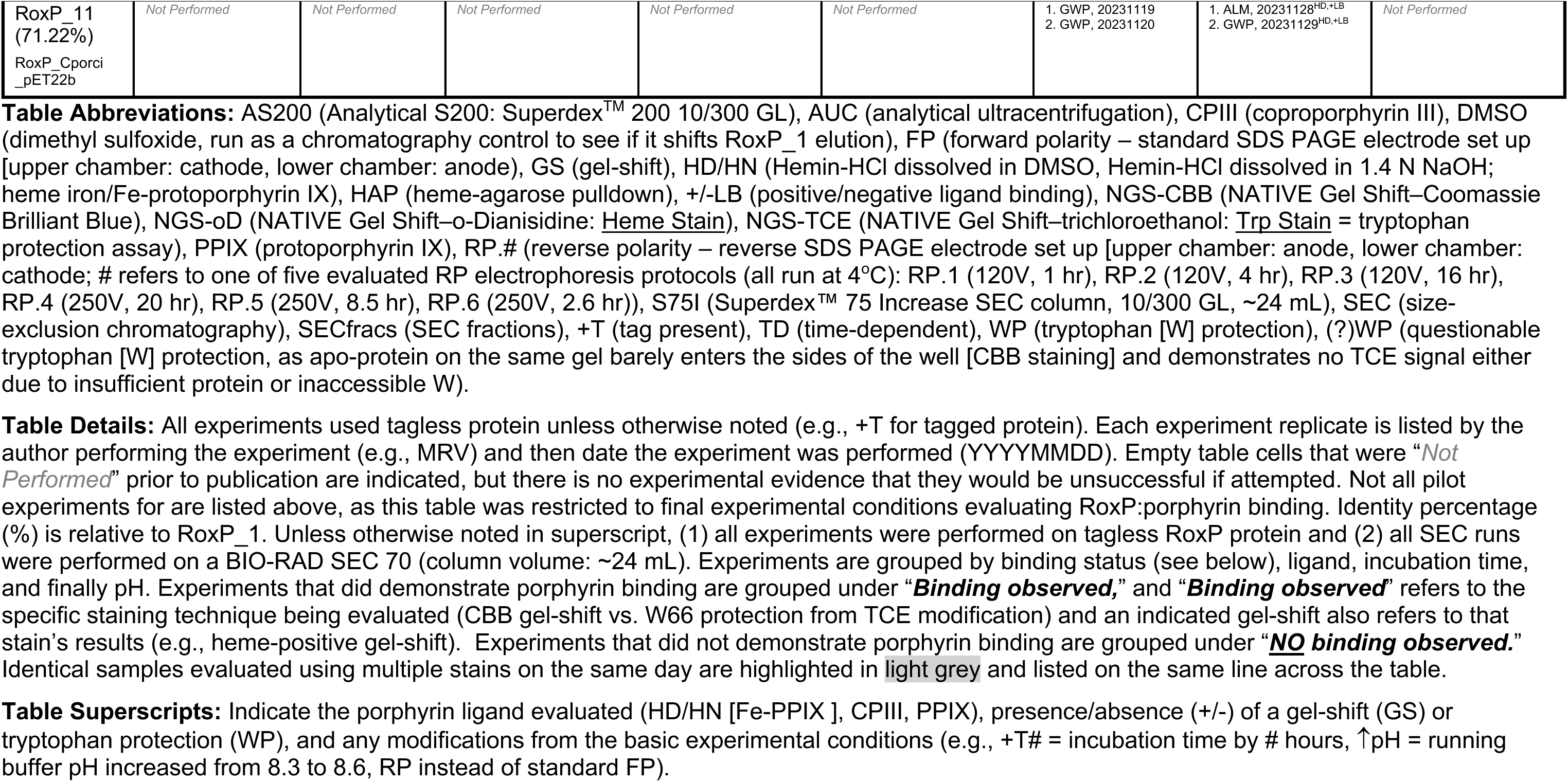
Summary of RoxP:Porphyrin Binding Assessments.

**SI Table 6.** Inter-residue Distance Between W66 and Potential Heme Ligands.

| **RoxP_1**<br>**Residue<sup>b,c</sup>** | **Ortholog Variant(s)<sup>a</sup>**<br>**Potential Heme Axial Ligand<sup>b</sup>** | **Ortholog Variant(s)<sup>a</sup>**<br>**Not a Heme Axial Ligand<sup>b</sup>** | **Distance to**<br>**RoxP_1 W66 (Å)** |
| --- | --- | --- | --- |
| H87 | --- | --- | 7.6 |
| H63 | --- | RoxP_7,8 H63R | 13.3 |
| Y116 | --- | --- | 14.8 |
| H98 | --- | --- | 14.9 |
| K74 | --- | RoxP_4-8 K74Q, RoxP_11 K74T | 17.5 |
| K61 | --- | RoxP_11 K61Q | 17.6 |
| H115 | --- | --- | 20.4 |
| Y49 | RoxP_11 Y49H | RoxP_4-6 Y49R | 25.8 |

<sup>a</sup> Limited to those RoxP orthologs that were demonstrated to bind heme (SI Table 5).
<sup>b</sup> Potential heme axial ligands include cysteine (C), histidine (H), lysine (K), methionine (M), and tyrosine (Y).
<sup>c</sup> Limited to those potential heme axial ligands<sup>b</sup> within 30 Å of RoxP\_1 W66.

**SI Table 7a.** Summary of Anti-RoxP Binding of RoxP Orthologs.

| Antibody | Ortholog/Construct Variants That Affect <sup>a</sup> Antibody Binding |  | RoxP Orthologs |  |
| --- | --- | --- | --- | --- |
|  | Shared Across Antibodies | Antibody-specific | Null Binders <sup>b,c</sup> | Binders <sup>c,d</sup> |
| **2H2F3** | T34A, G89D/E | P91A | RoxP_5-8 | RoxP_1,10>>3>>>2>4>>11 |
| **3E10E3** | A28S/T, A29T, S30A, T34A, G89D/E, S100N | S100D/G, E101K | RoxP_5-8 | RoxP_1,10,11>>>3,4>2 |
| **4D8H1** | A28S/T, A29T, S30A, S31N, T36K, G89E, S100N | *G12D* <sup>e</sup>, *Q14R* <sup>e</sup>, S20I/P, M133A/T, D135N, L138A/I/K | RoxP_5,7,8,11 | RoxP_1,10>2>4>3>>6 <sup>f</sup> |
| **5F3D11** | S31N, T34A, T36K, G89D/E, S100N | N-terminal tag <sup>g</sup>, T69A | RoxP_5-8,11 | RoxP_1>10>>2,4>3 <sup>h</sup> |

<sup>a</sup> Impeded antibody binding unless noted otherwise in *italics*.
<sup>b</sup> Assessment was based on Fig 6f experiment (RoxP coating concentration: 10.0 µg/mL; detection reagent: SA-Poly-HRP)
<sup>c</sup> Binding was defined as ≥5 standard deviation (SD) above the average of the buffer-only control for an antibody.
<sup>d</sup> Assessment was based on Fig 6f, SI Fig 15a (RoxP coating concentration: 1.0 µg/mL; detection reagent: SA-HRP), and SI Fig 15b (RoxP coating concentration: 1.0 µg/mL; detection reagent: SA-Poly-HRP).
<sup>e</sup> Appeared to *improve binding* to antibody.
<sup>f</sup> SI Fig15a was used for order of RoxP\_2-4 based on its greater dynamic range with 1.0 µg/mL RoxP coating concentration and SA-HRP detection reagent.
Alternative affinity ranking based on SI Fig 15b: RoxP\_1,3,4,10>2>>6.
<sup>g</sup> Presence of N-terminal tag (20 amino acids) appeared to impede antibody binding (SI Fig 15a,b).
<sup>h</sup> SI Fig15b was used for order of RoxP\_2-4 based on its greater dynamic range with 1.0 µg/mL RoxP coating concentration and SA-Poly-HRP detection reagent. Alternative affinity ranking based on SI Fig 15a: RoxP\_1>10>>2>4>3.

**SI Table 7b.**
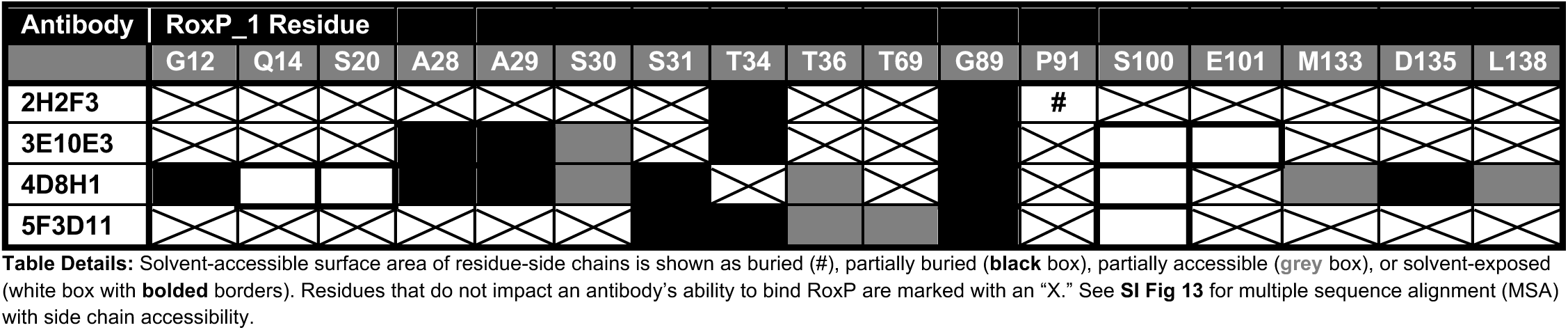
Solvent Accessibility of RoxP Variant Positions Implicated in Anti-RoxP Antibody.

**SI Table 8:**
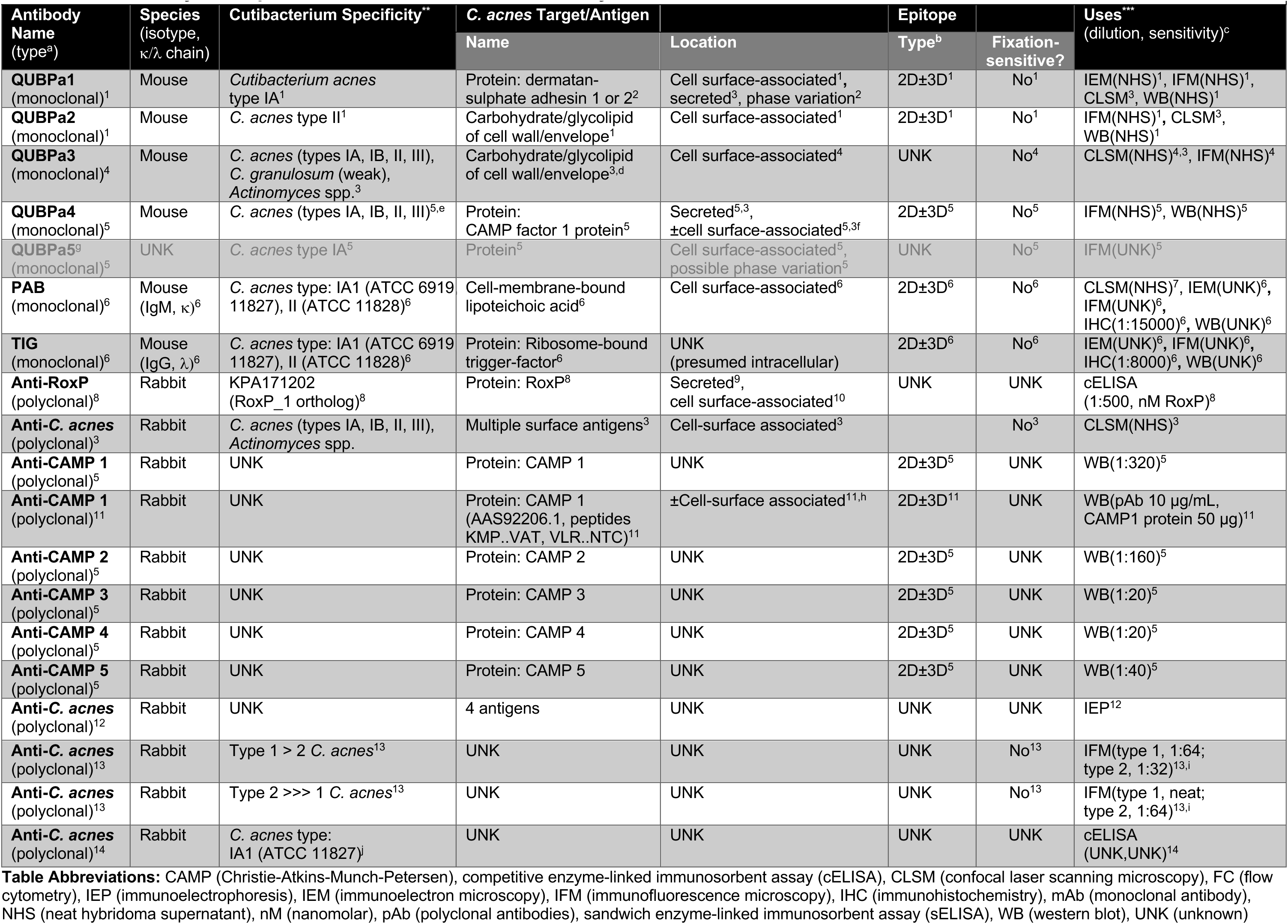
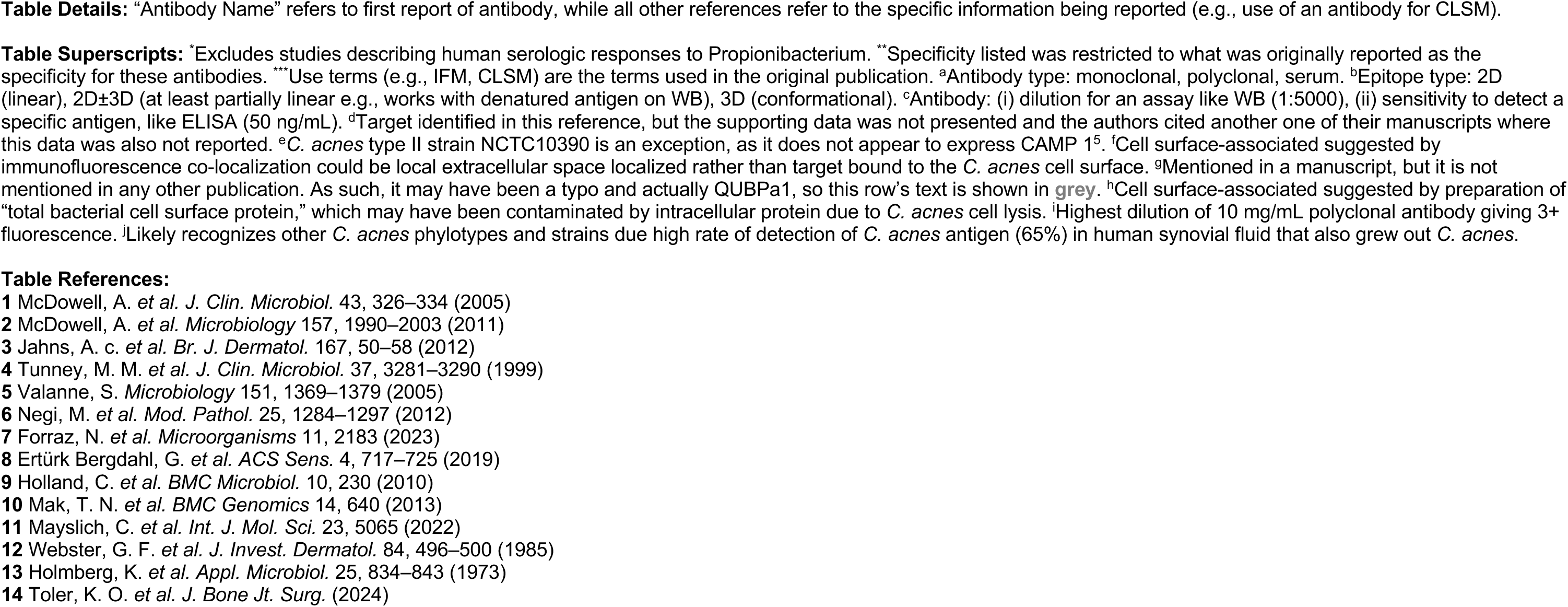
Summary of Propionibacterium Antibodies Previously Described in the Literature*.

**SI Table 9:**
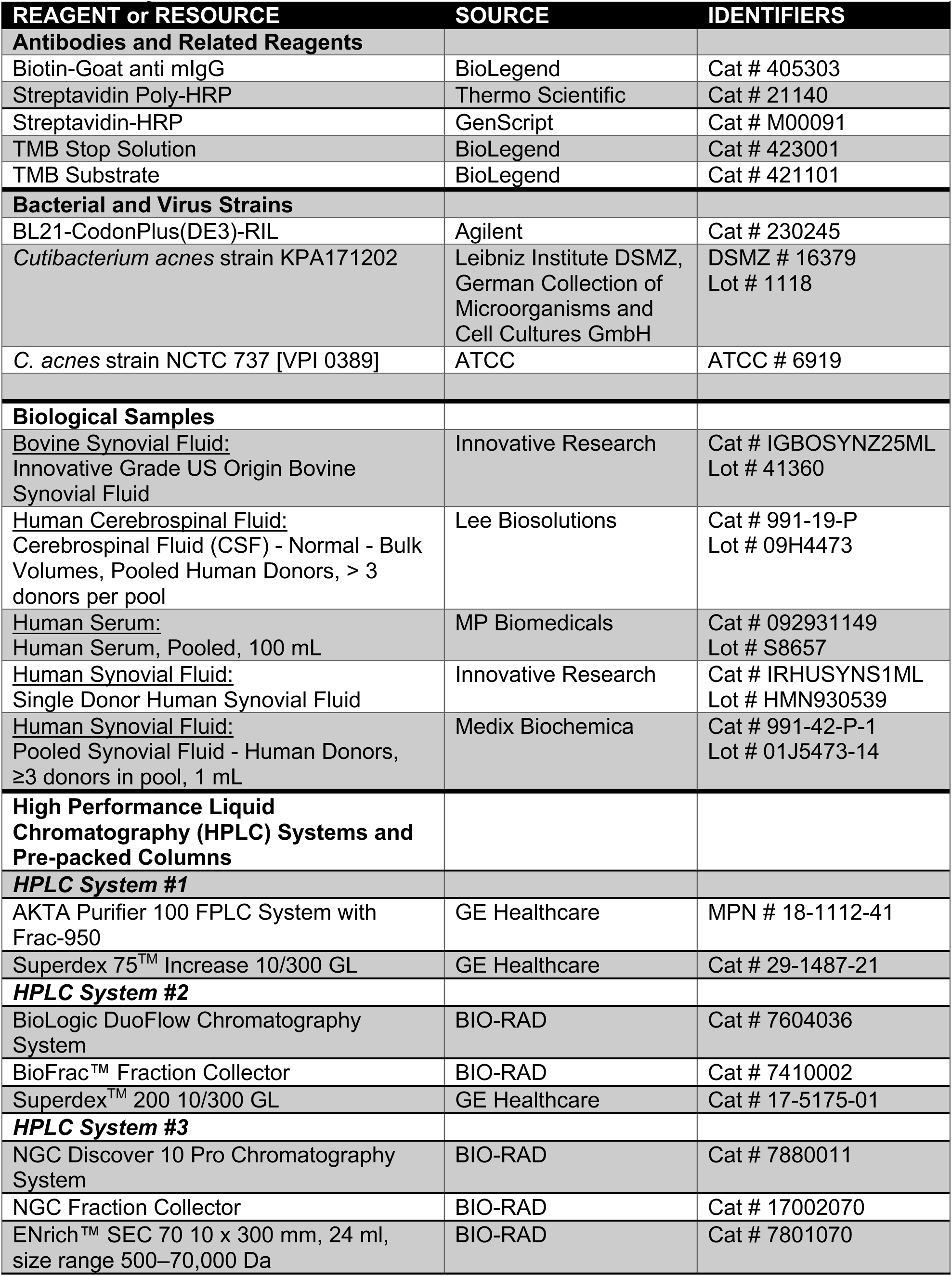
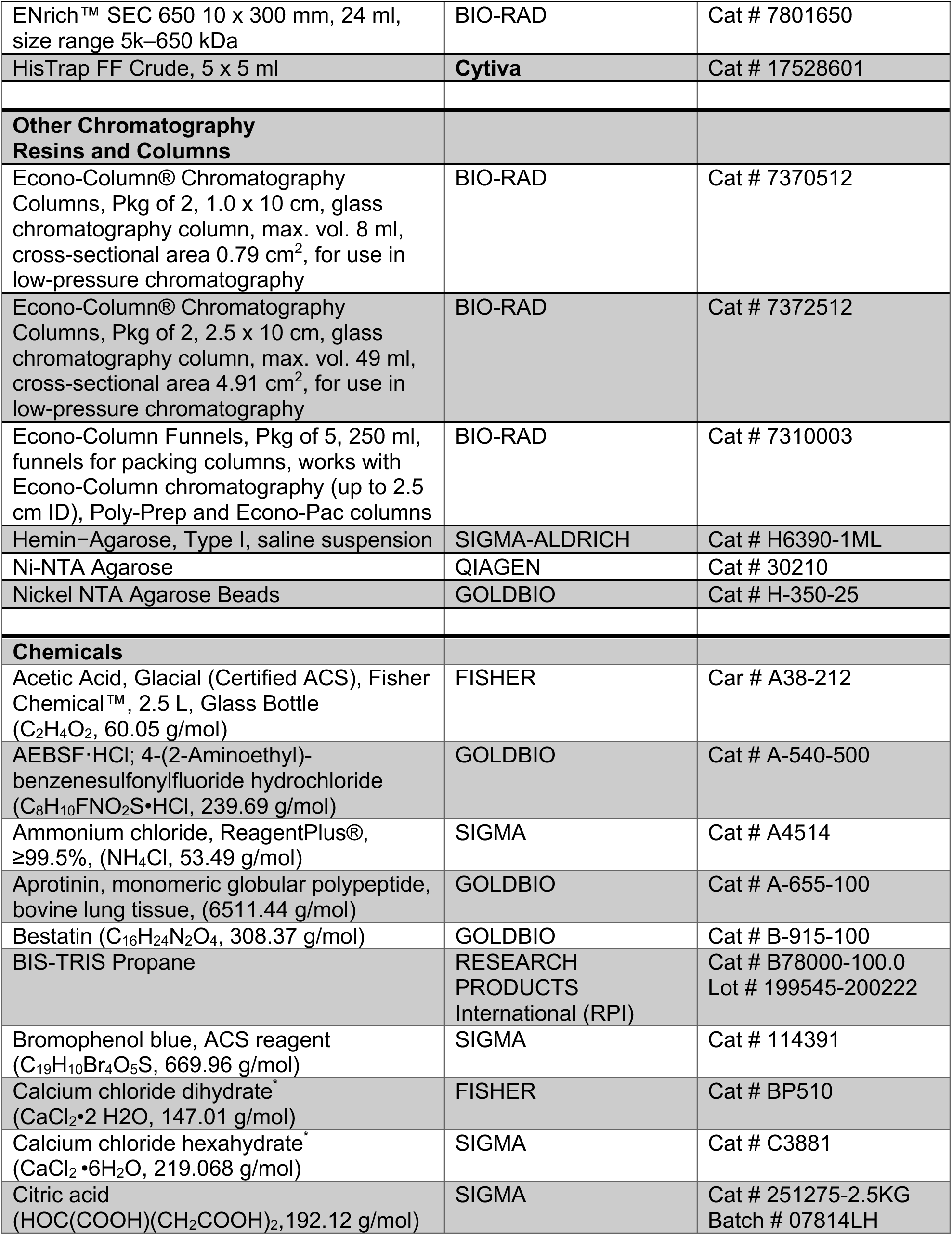
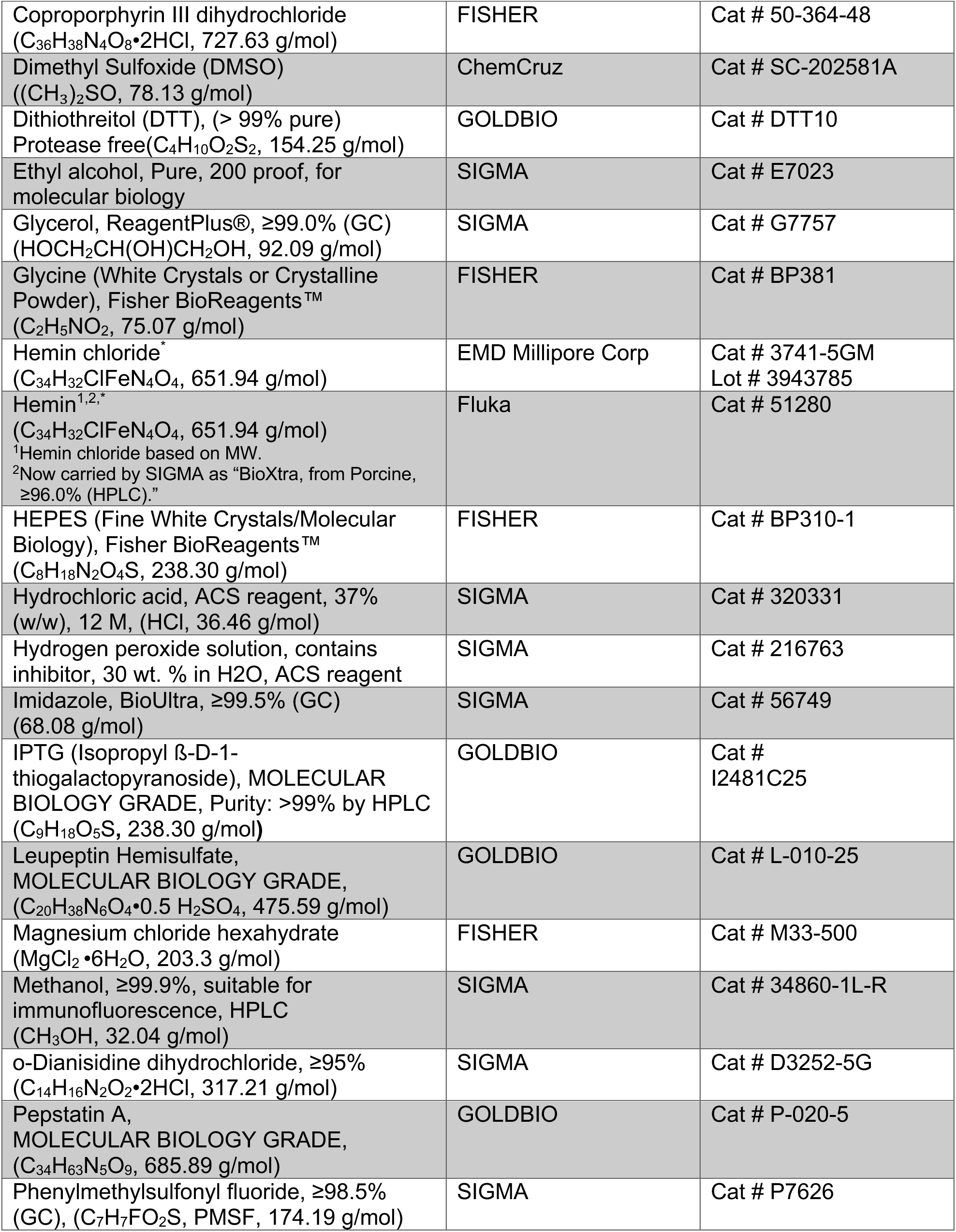
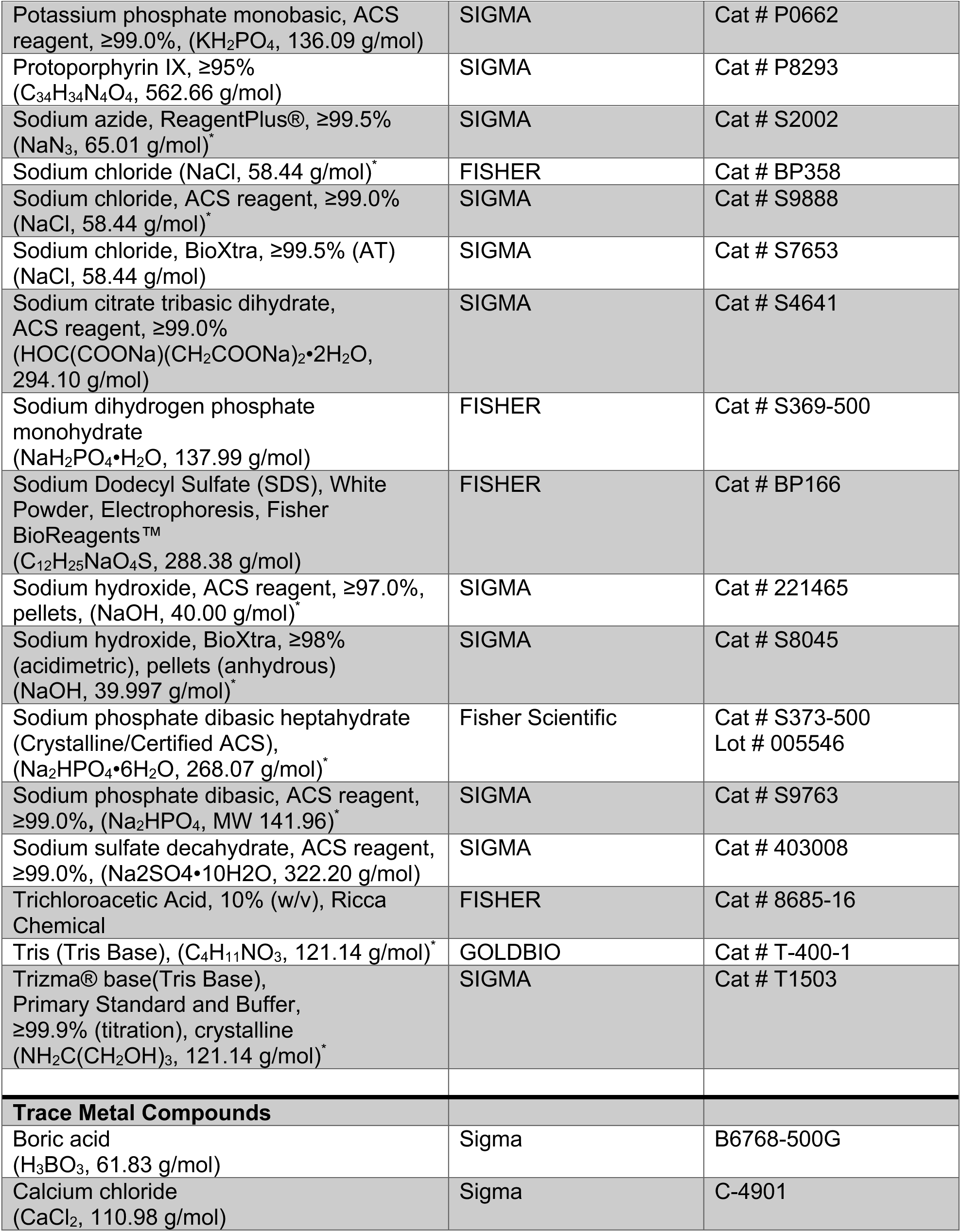
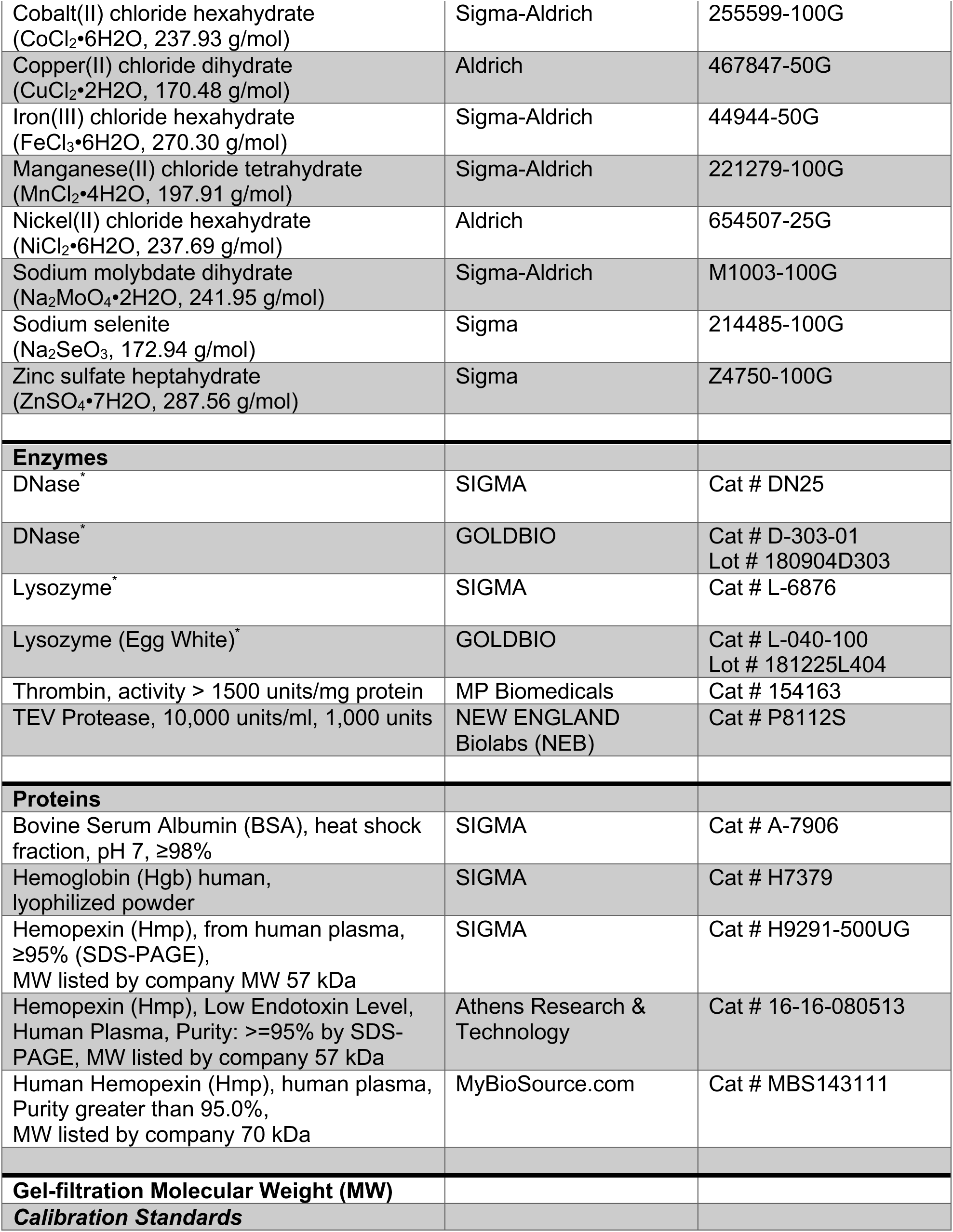
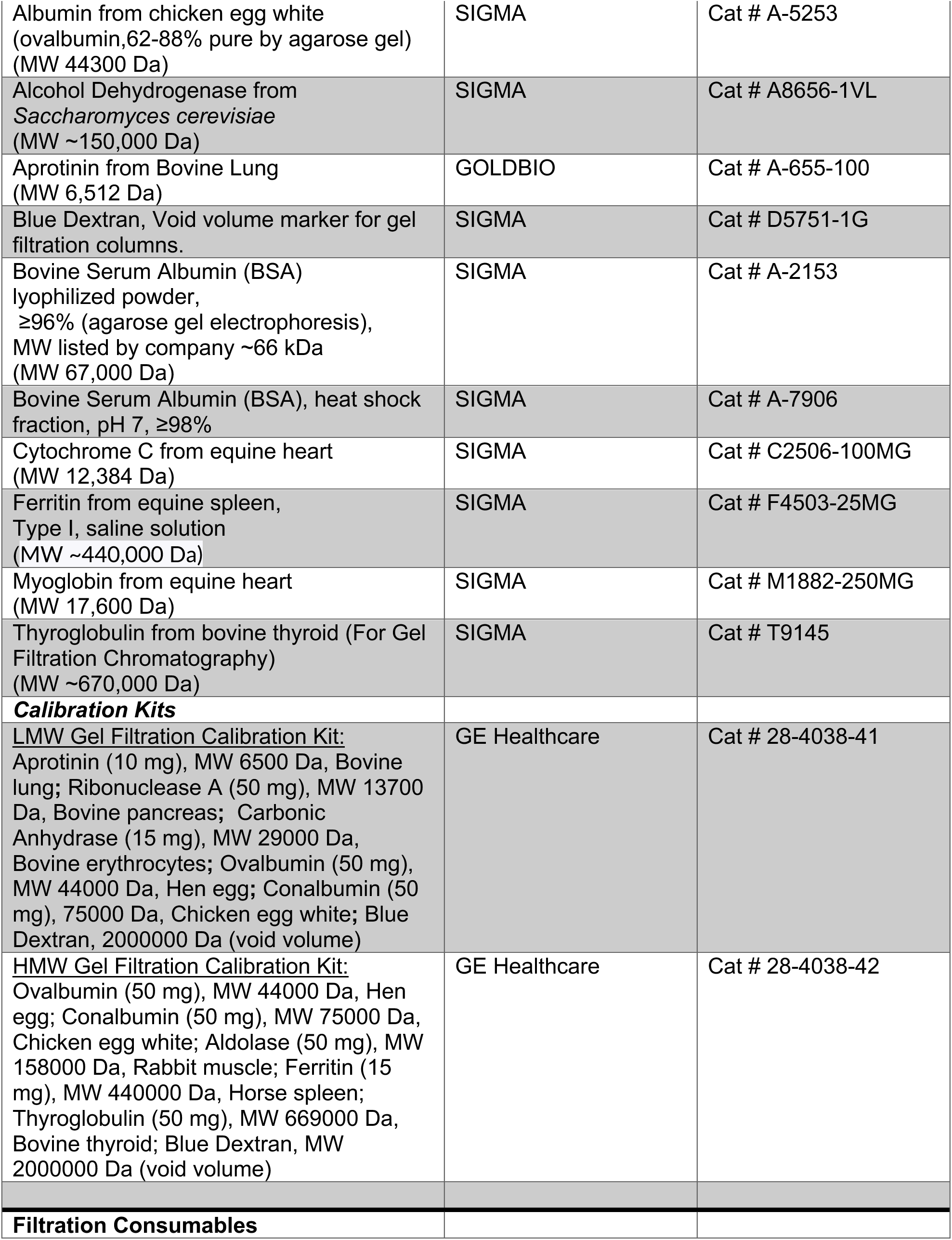
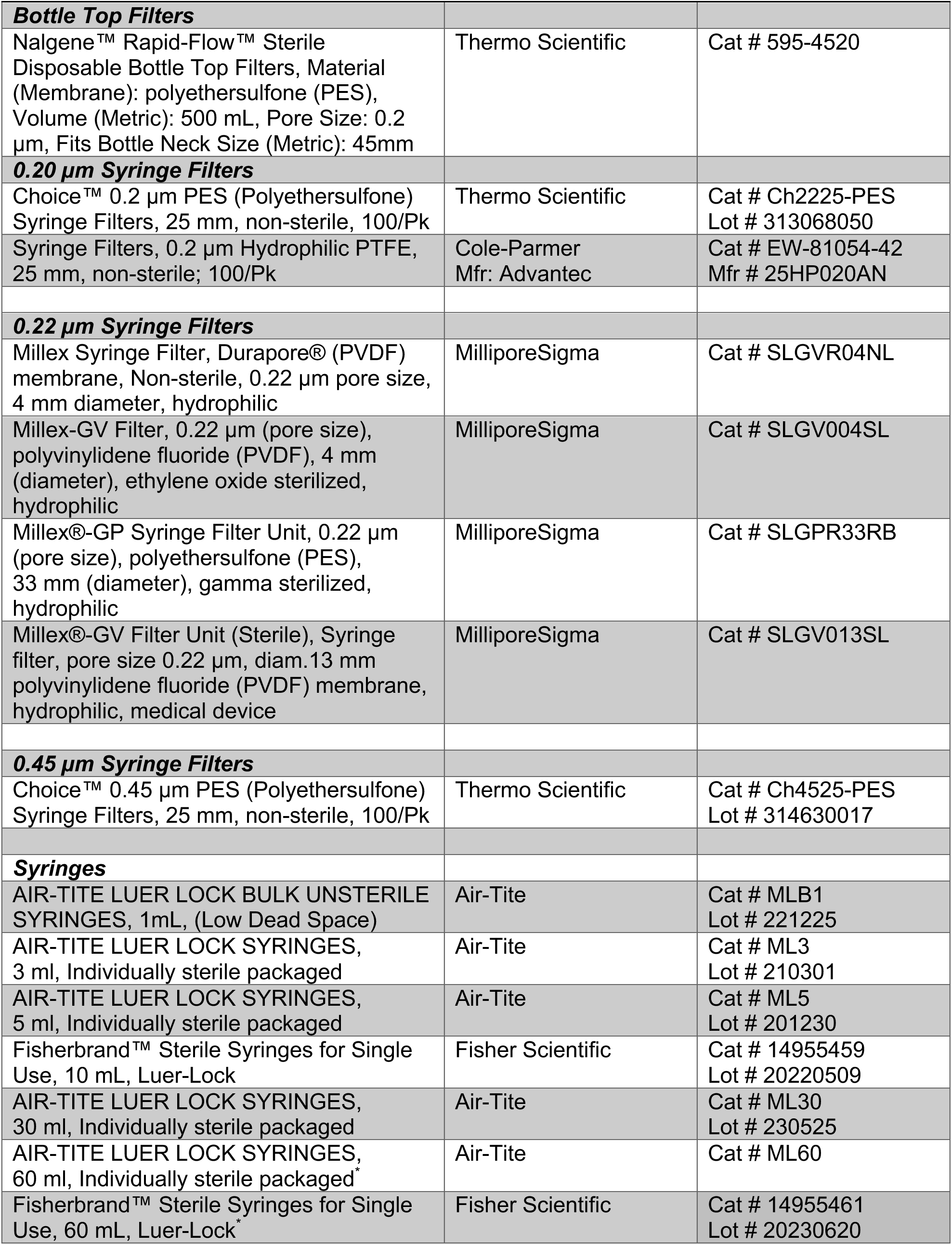
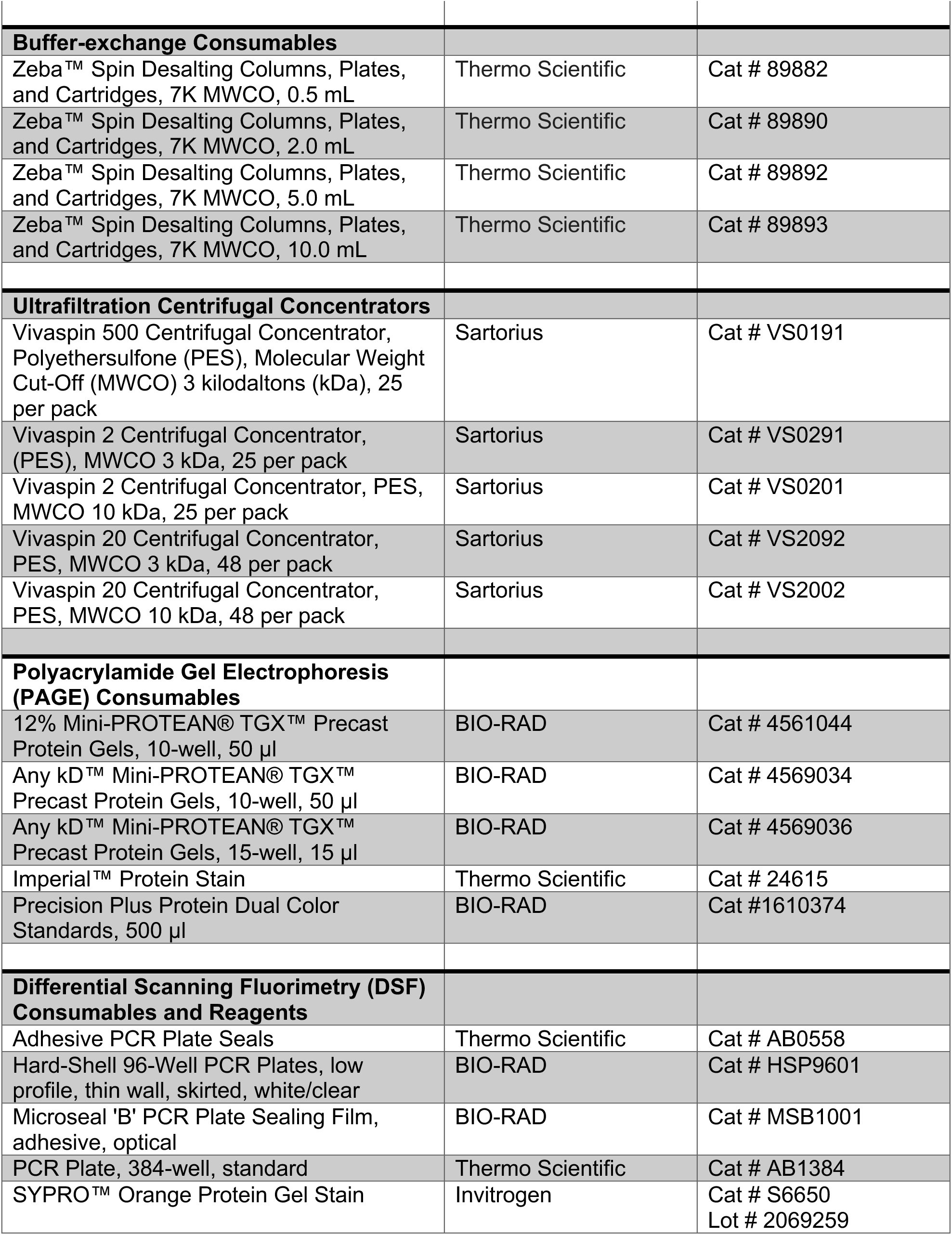
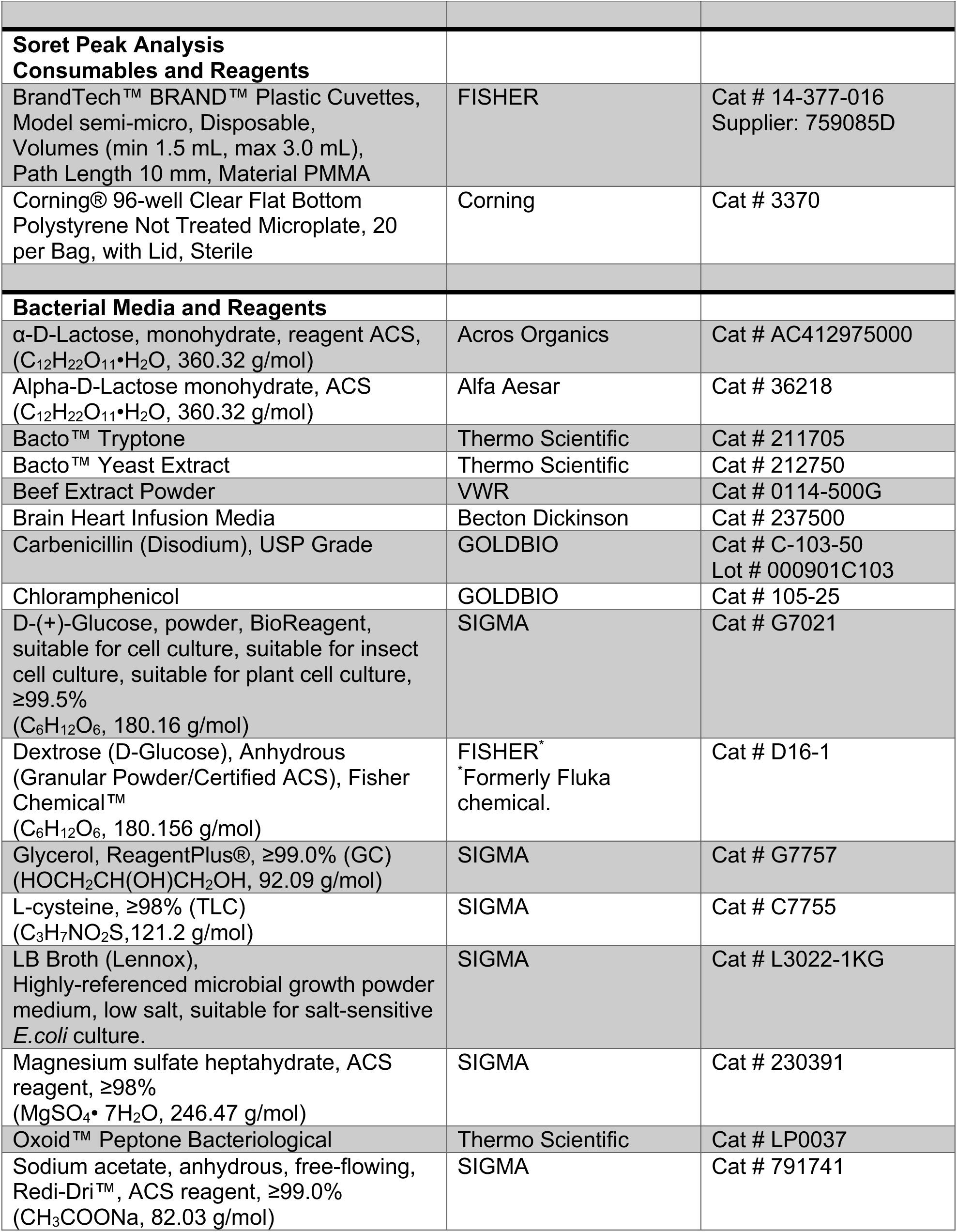
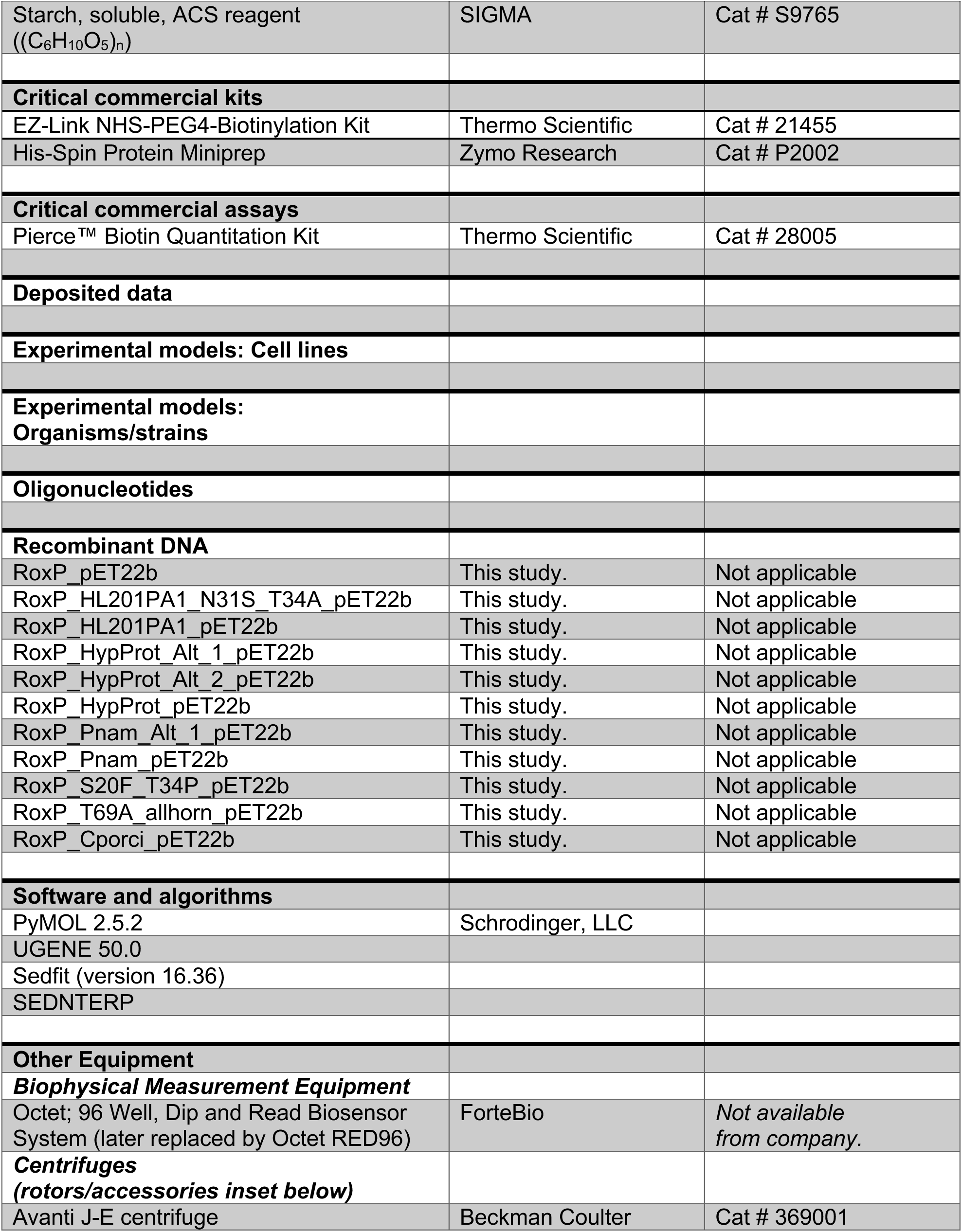
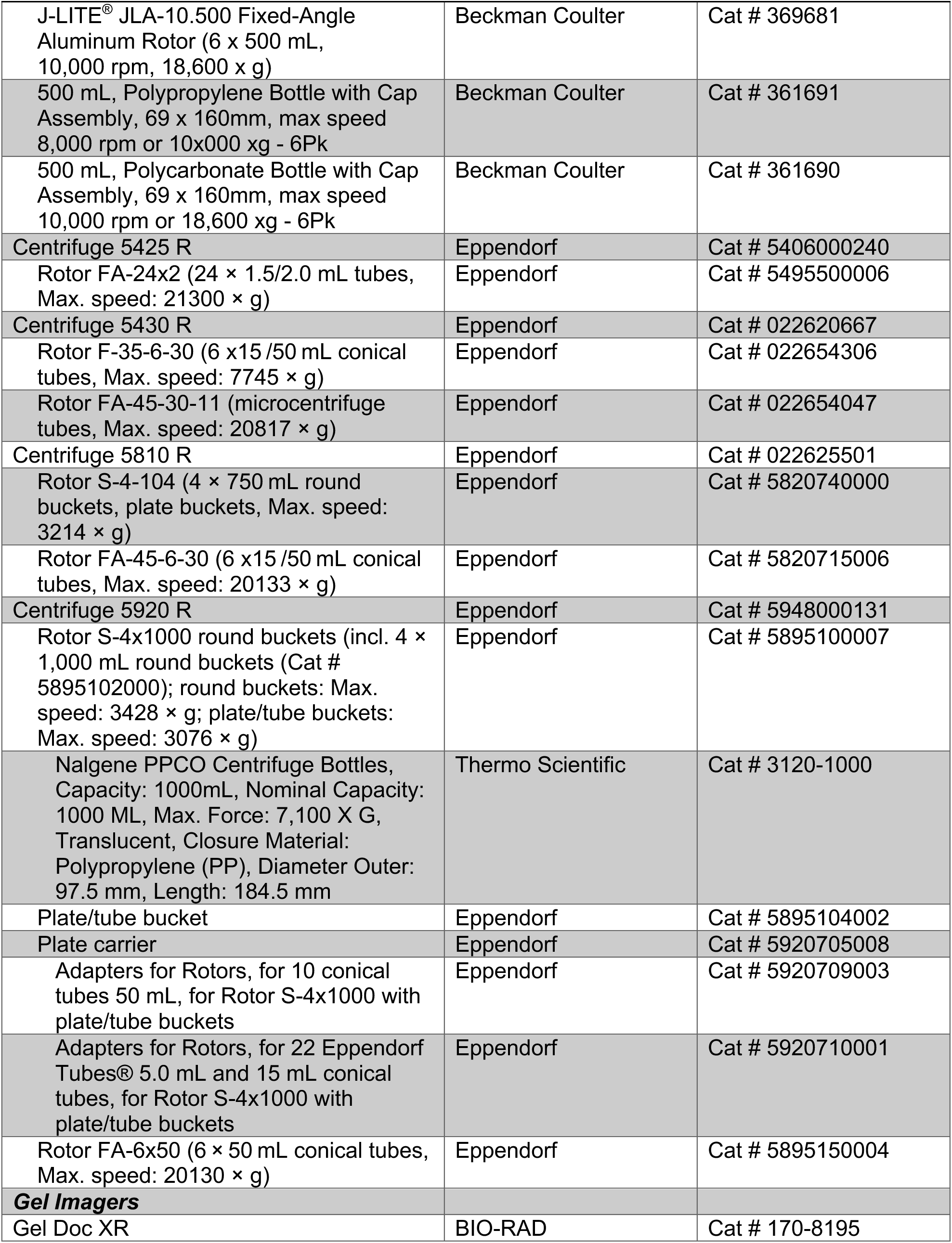
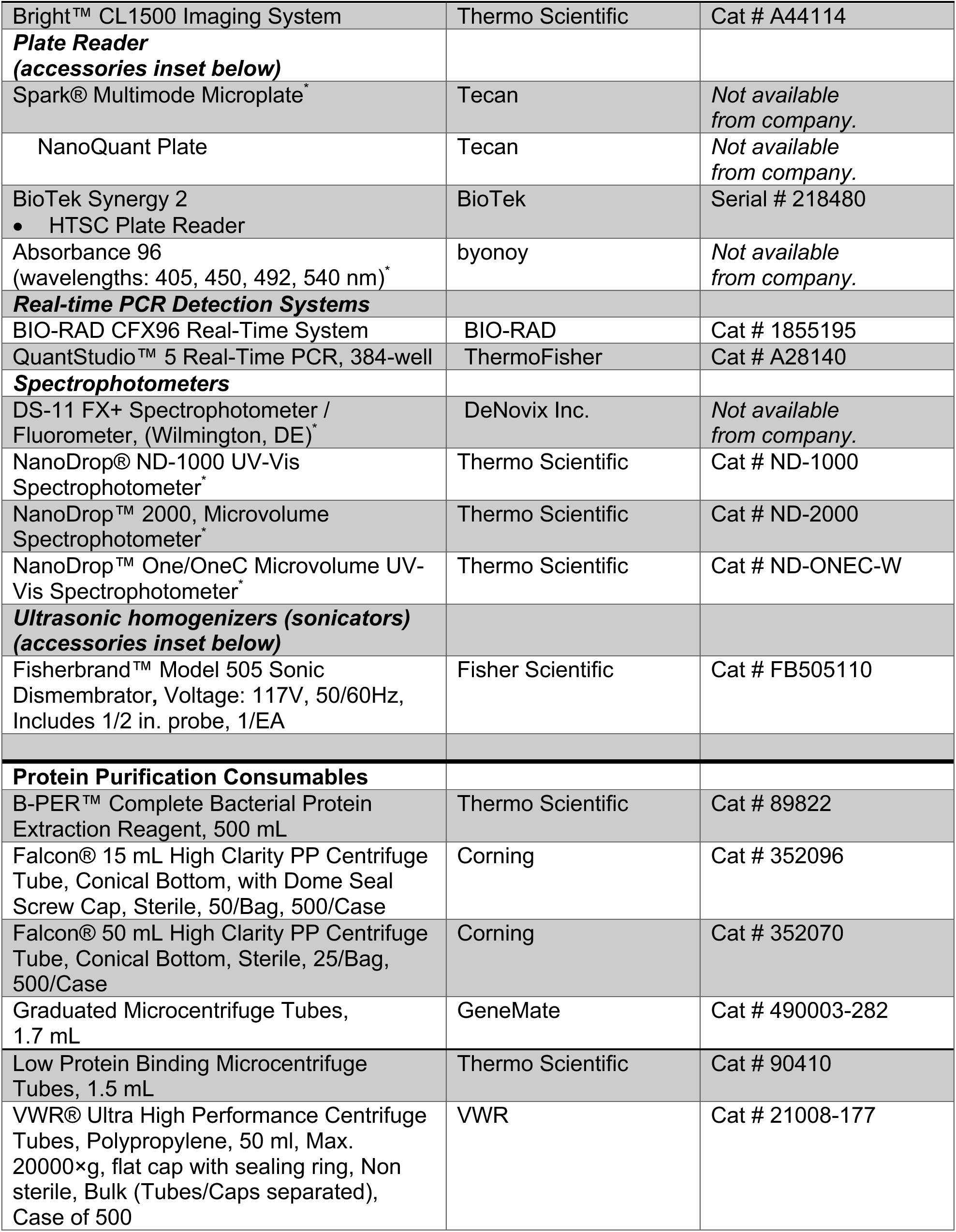
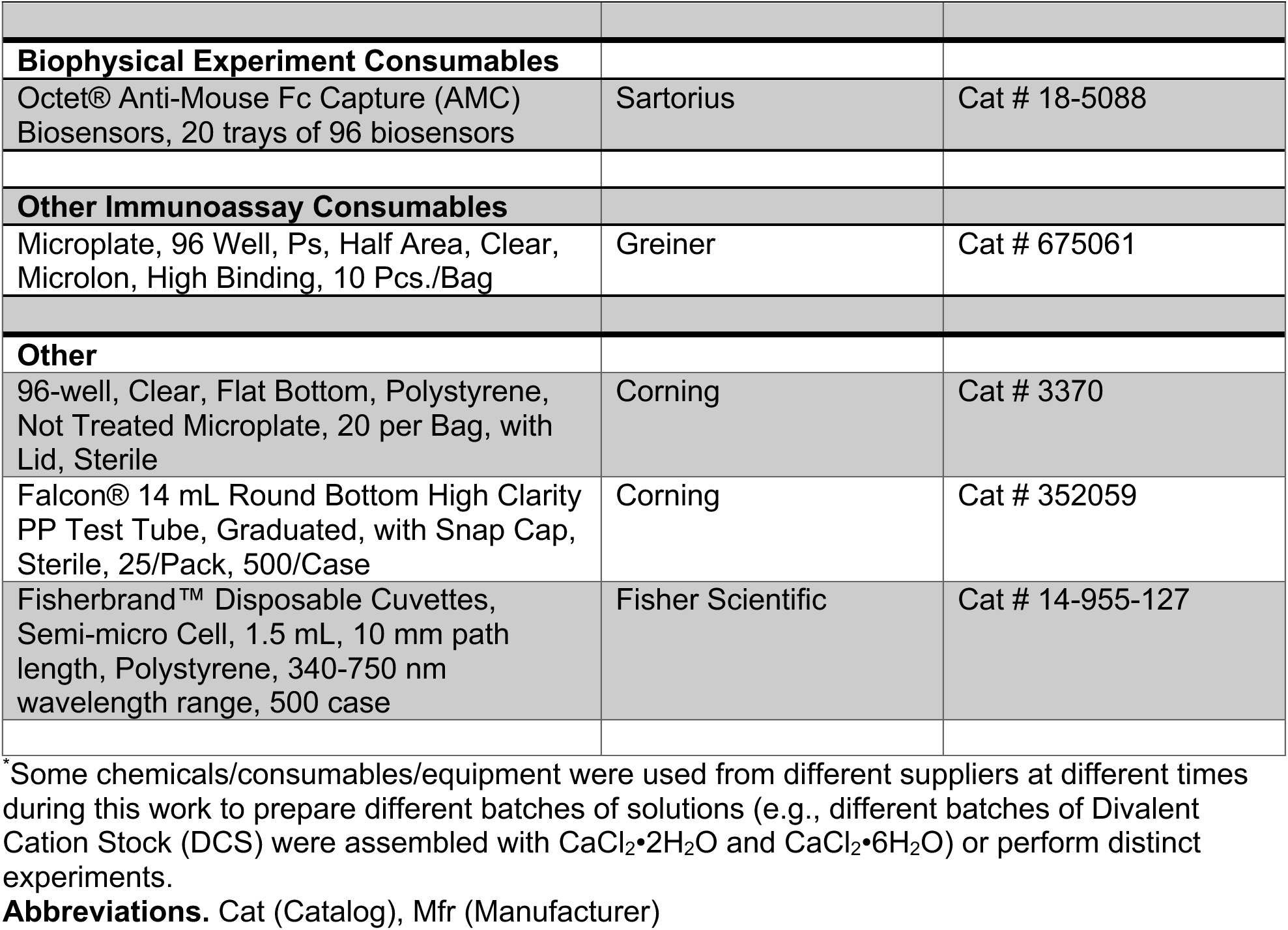
Key Resources Table.

**SI Table 10.** Effect of Hematin on pH in HSB samples.

| Reagent Concentrations |  |  | pH |
| --- | --- | --- | --- |
| Hematin (mM) | NaOH (mN) | Tris pH 7.0 (mM) |  |
| 9.04000 | 0.00000 | 0.00000 | Not Available (NA)* |
| 4.52000 | 28.00000 | 27.80000 | NA* |
| 0.90400 | 0.00000 | 0.00000 | 6.5-7.5 |
| 0.45000 | 2.80000 | 2.78000 | 6.5-7.5 |
| 0.09040 | 0.00000 | 0.00000 | 7.0-7.5 |
| 0.04520 | 0.28000 | 0.27800 | 7.0-7.5 |
| 0.00904 | 0.00000 | 0.00000 | 7.0 |
| 0.00452 | 0.02800 | 0.02780 | 7.0 |

*\*Heme darkened all three-color panels too much to be interpreted.*

**SI Table 11.** Assembly of DSF pH Screen.

| **1.0 M Citric**<br>**Acid (mL)** | **1.0 M BIS-TRIS**<br>**Propane (mL)** | **3.0 M NaCl**<br>**(mL)** | **Water**<br>**(mL)** | **Expected**<br>**pH** | **Measured**<br>**pH** |
| --- | --- | --- | --- | --- | --- |
| 1.8 | 0.2 | 1.0 | 17.0 | 2.5 | **2.30** |
| 1.7 | 0.3 | 1.0 | 17.0 | 2.7 | **2.54** |
| 1.6 | 0.4 | 1.0 | 17.0 | 2.9 | **2.80** |
| 1.5 | 0.5 | 1.0 | 17.0 | 3.2 | **3.01** |
| 1.4 | 0.6 | 1.0 | 17.0 | 3.4 | **3.31** |
| 1.3 | 0.7 | 1.0 | 17.0 | 3.8 | **3.63** |
| 1.2 | 0.8 | 1.0 | 17.0 | 4.1 | **3.97** |
| 1.1 | 0.9 | 1.0 | 17.0 | 4.5 | **4.36** |
| 1.0 | 1.0 | 1.0 | 17.0 | 5.0 | **4.81** |
| 0.9 | 1.1 | 1.0 | 17.0 | 5.5 | **5.39** |
| 0.8 | 1.2 | 1.0 | 17.0 | 6.4 | **6.29** |
| 0.7 | 1.3 | 1.0 | 17.0 | 7.1 | **7.03** |
| 0.6 | 1.4 | 1.0 | 17.0 | 7.6 | **7.70** |
| 0.5 | 1.5 | 1.0 | 17.0 | 8.2 | **8.41** |
| 0.4 | 1.6 | 1.0 | 17.0 | 8.8 | **9.04** |
| 0.3 | 1.7 | 1.0 | 17.0 | 9.1 | **9.45** |
| 0.2 | 1.8 | 1.0 | 17.0 | 9.5 | **9.81** |

## Notes

### Competing Interest Statement

The authors have declared no competing interest.

