## Supplemental Files 1-12 for "*Cutibacterium* adaptation to life on humans provides a novel biomarker of *C. acnes* infections": 20240820_SI_File_05_AntiRoxPmAb_VarDom_Seq_v7.docx

**2H2F3 - Antibody Variable Domain Sequencing of Hybridoma Clone**

Mouse IgG2b,K

**LEGEND**

Framework Region (FR)

Complementarity-determining region (CDR)

FR Positively Charged Residues: H, K, R

CDR Positively Charged Residues: **H**, **K**, **R**

Aliphatic index: relative volume occupied by aliphatic side chains (Ala, Val, Iso, Leu)

Grand average of hydropathicity (GRAVY): <0 hydrophilic, >0 hydrophobic

**Heavy chain: Amino acid sequence (140 aa)**

*MGWSCIIFFLVATATGVHS*QVQLQQSGAELVRPGVSVKISCKGSGYTFT**DYAMH**WVKQSHAKSLEWIG**IISTYYGNASYNQKFKG**KATMTVDKSSSTAYMELARLTSEDSAIYYCAR**VPYYGSSYAMDY**WGQGTSVTVSS

**Light chain: Amino acid sequence (131 aa)**

*MKLPVRLLVLMFWIPASSS*DVLMTQTPLSLPVSLGDQASISC**RSSQNIVHSDGNTYLE**WYLQKPGQSPKLLIY**KVSNRFS**GVPDRFSGSGSGTDFTLKISRVEAEDLGVYYC**FQGSHVPWT**FGGGTKLEIK

All CDRs:

**DYAMH, IISTYYGNASYNQKFKG, VPYYGSSYAMDY, RSSQNIVHSDGNTYLE, KVSNRFS, FQGSHVPWT**

*ExPASy ProtParam Assessment of All CDRs:*

Theoretical pI: 8.20

Arg (R) residues: 2

His (H) residues: 3

Lys (K) residues: 3

Trp (W) residues: 1

Aliphatic index: 45.76

GRAVY: -0.735

*ExPASy ProtParam Assessment of Post-signal Peptide Cleavage Variable Domains:*

Theoretical pI: 8.55

**3E10E3 - Antibody Variable Domain Sequencing of Hybridoma Clone**

Mouse IgG2b,K

**LEGEND**

Framework Region (FR)

Complementarity-determining region (CDR)

FR Positively Charged Residues: H, K, R

CDR Positively Charged Residues: **H**, **K**, **R**

Aliphatic index: relative volume occupied by aliphatic side chains (Ala, Val, Iso, Leu)

Grand average of hydropathicity (GRAVY): <0 hydrophilic, >0 hydrophobic

**Heavy chain: Amino acid sequence (136 aa)**

*MGRLTSSFLLLIVPAYVLS*QVTLKESGPGILQPSQTLSLTCSFSGFSLS**TYGIGVG**WIRQPSGKGLEWLA**HIWWNDNKYYNTALKS**RLTISKDTSNNQVFLKIASVDTADTATYYCAR**SLPYFDY**WGQGTTLTVSS

**Light chain: Amino acid sequence (127 aa)**

*MSVPTQVLGLLLLWLTGARC*DIQMTQSPASLSASVGETVTITC**RASENIYSYLA**WYQQKQGKSPQFLVY**NAKTLAE**GVPSRFSGSGSGTQFSLKINSLQPEDFGSYYC**QHHYGIPRT**FGGGTKLEIK

All CDRs:

**TYGIGVG, HIWWNDNKYYNTALKS, SLPYFDY, RASENIYSYLA, NAKTLAE, QHHYGIPRT**

*ExPASy ProtParam Assessment of All CDRs:*

Theoretical pI: 7.96

Arg (R) residues: 2

His (H) residues: 3

Lys (K) residues: 3

Trp (W) residues: 2

Aliphatic index: 68.60

GRAVY: -0.686

*ExPASy ProtParam Assessment of Post-signal Peptide Cleavage Variable Domains:*

Theoretical pI: 8.79

*ExPASy ProtParam Assessment of Heavy Chain CDR2:*

Theoretical pI: 8.44

**4D8H1- Antibody Variable Domain Sequencing of Hybridoma Clone**

Mouse IgG2b,λ

**LEGEND**

Framework Region (FR)

Complementarity-determining region (CDR)

FR Positively Charged Residues: H, K, R

CDR Positively Charged Residues: **H**, **K**, **R**

Aliphatic index: relative volume occupied by aliphatic side chains (Ala, Val, Iso, Leu)

Grand average of hydropathicity (GRAVY): <0 hydrophilic, >0 hydrophobic

**Heavy chain: Amino acid sequence (134 aa)**

*MDFGLIFFIVALLKGVQC*EVKLLESGGGLVQPGGSLKLSCAASGFDFS**RYWMS**WVRQAPGKGLEWIG**EINPDSSTINYTPSLKD**KFIISRDNAKNTLYLQMSKVRSEDTALYYCAR**YYSRFAY**WGQGTLVTVSA

**Light chain: Amino acid sequence (128 aa)**

*MAWISLILSLLALSSGAIS*QAVVTQESALTTSPGETVTLTC**RSSTGAVTTRNYAN**WVQEKPDHLFTGLIG**DTNNRAP**GVPARFSGSLIGDKAALTITGAQTEDEAIYFC**ALWYSNHWV**FGGGTKLTVL

All CDRs:

**RYWMS, EINPDSSTINYTPSLKD, YYSRFAY, RSSTGAVTTRNYAN, DTNNRAP, ALWYSNHWV**

*ExPASy ProtParam Assessment of All CDRs:*

Theoretical pI: 9.16

Arg (R) residues: 5

His (H) residues: 1

Lys (K) residues: 1

Trp (W) residues: 3

Aliphatic index: 44.75

GRAVY: -0.966

*ExPASy ProtParam Assessment of Post-signal Peptide Cleavage Variable Domains:*

Theoretical pI: 7.87

**5F3D11 - Antibody Variable Domain Sequencing of Hybridoma Clone**

Mouse IgG2b,K

**LEGEND**

Framework Region (FR)

Complementarity-determining region (CDR)

FR Positively Charged Residues: H, K, R

CDR Positively Charged Residues: **H**, **K**, **R**

Aliphatic index: relative volume occupied by aliphatic side chains (Ala, Val, Iso, Leu)

Grand average of hydropathicity (GRAVY): <0 hydrophilic, >0 hydrophobic

**Heavy chain: Amino acid sequence (140 aa)**

*MDRLTSSFLLLIVPVYVLS*QVTLKESGPGILQPSQTLSLTCSFSGFSLS**TFGMGVS**WIRQPSGKGLEWLA**HIYWDDDKHYNPSLKS**RLTISKDTSNNQVFLKITTVDTADTATYYCAR**RAWGDYDAFAY**WGQGTLVTVSA

**Light chain: Amino acid sequence (131 aa)**

*MKLPVRLLVLMFWIPASSS*DVVMTQTPLSLPVSLGDQASISC**RSSQSLVHSNGNTYLH**WYLQKPGQSPKLLIY**KVSNRFS**GVPDRFSGSGSGTDFTLKISRVEAEDLGVYFC**SQSTHVPPT**FGGGTKLEIK

All CDRs:

**TFGMGVS, HIYWDDDKHYNPSLKS, RAWGDYDAFAY, RSSQSLVHSNGNTYLH, KVSNRFS, SQSTHVPPT**

*ExPASy ProtParam Assessment of All CDRs:*

Theoretical pI: 8.07

Arg (R) residues: 3

His (H) residues: 5

Lys (K) residues: 3

Trp (W) residues: 2

Aliphatic index: 45.76

GRAVY: -0.873

*ExPASy ProtParam Assessment of Post-signal Peptide Cleavage Variable Domains:*

Theoretical pI: 8.32

*ExPASy ProtParam Assessment of Heavy Chain CDR2:*

Theoretical pI: 5.99
